# Exploring heteroaromatic rings as a replacement for the labile amide of antiplasmodial pantothenamides

**DOI:** 10.1101/2020.10.02.324079

**Authors:** Jinming Guan, Christina Spry, Erick T. Tjhin, Penghui Yang, Tanakorn Kittikool, Vanessa M. Howieson, Harriet Ling, Lora Starrs, Gaetan Burgio, Kevin J. Saliba, Karine Auclair

## Abstract

The *Plasmodium* parasites that cause malaria are adept at developing resistance to antimalarial drugs, necessitating the search for new antiplasmodials. Although several amide analogs of pantothenate (pantothenamides) show potent antiplasmodial activity, hydrolysis by pantetheinases (or vanins) present in blood rapidly inactivates them. We report herein the facile synthesis and biological activity of a small library of pantothenamide analogs in which the labile amide group is replaced with a variety of heteroaromatic rings. Several of the new analogs display antiplasmodial activity in the nanomolar range against *P. falciparum* and/or *P. knowlesi* in the presence of pantetheinase. A previously reported triazole and an isoxazole derivative presented here were further characterized and found to possess high selectivity indices, medium or high Caco-2 permeability, and medium or low microsomal clearance *in vitro*. Although we show here that the two compounds fail to suppress proliferation of *P. berghei in vivo*, pharmacokinetic and contact time data presented provide a benchmark for the compound profile required to achieve antiplasmodial activity in mice and should facilitate lead optimization.

## 1. INTRODUCTION

There were 228 million cases of malaria worldwide in 2018, leading to 405,000 deaths.^1^ Malaria is caused by unicellular protozoan *Plasmodium* parasites, of which six species infect humans: *P. falciparum*, *P. vivax, P. malariae*, *P. ovale curtisi, P. ovale wallikeri* and *P. knowlesi*.^2^ Resistance to the currently marketed antimalarial drugs, including standard artemisinin-combination therapies, is spreading, making the discovery of new antimalarials ever more crucial.^3^ New structural classes of antiplasmodials with a novel mechanism of action are particularly desirable to minimize cross-resistance. The antibacterial and antifungal activities of pantothenamides (amides of pantothenate; Figure 1) were recognized in the 1970’s;^4^ whereas their antiplasmodial activity was demonstrated more recently.^5^ The unique mechanism of action of pantothenamides involves metabolic activation by the coenzyme A (CoA) biosynthetic pathway, which otherwise uses pantothenate as a substrate. The resultant antimetabolites are proposed to interfere with a subsequent CoA biosynthetic step or downstream CoA-utilizing enzymes, thereby inhibiting parasite proliferation.^6–9^ Pantothenamides are easily accessible synthetic targets, potent, and typically non-cytotoxic.^10^ Their clinical use is however hindered by poor stability in human serum; pantothenamides are rapidly hydrolyzed to pantothenate and the corresponding amine by pantetheinases (also known as vanins), which are present in serum (Figure 1).^5, 11^

**Figure 1.**
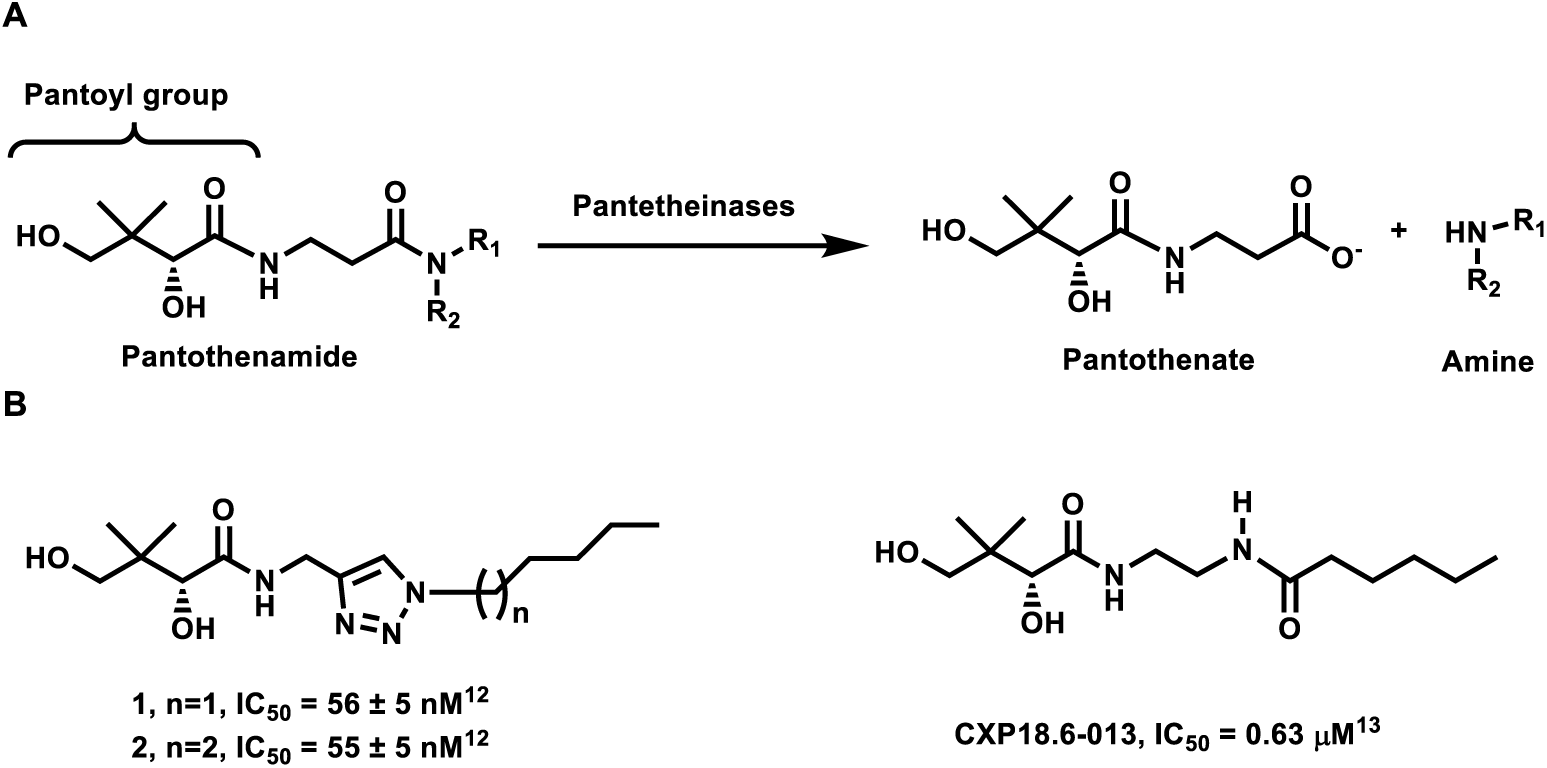
(A) Hydrolysis of pantothenamides by serum pantetheinases to yield pantothenate and an amine. The pantoyl moiety is indicated in the bracket. R_1_ is typically H and R_2_ is commonly an organic moiety. (B) Structure of compounds **1, 2** and **CXP18.6-013** with their respective 50% inhibitory concentrations (IC_50_ values) measured against intraerythrocytic *P. falciparum* in the presence of pantetheinase.^12–13^

We and others have previously explored several strategies to improve the biological activity of pantothenamides in the presence of pantetheinases. One approach consists of designing pantetheinase inhibitors to be used in combination with pantothenamides.^11, 14^ Another strategy is to modify pantothenamides to protect them from pantetheinase-mediated hydrolysis.^9, 12–13, 15–21^ Such alterations have been reported at the secondary hydroxyl group,^17, 20^ the geminal-dimethyl group,^15, 17, 19^ the β-alanine moiety,^9, 16, 18^ and the labile amide itself,^12–13, 21^ sometimes at the cost of activity. Our team has reported a series of analogs in which the labile amide group is replaced with a 1,2,3-triazole ring that translates into enhanced stability and potency.^21^ Our previous exploration of structure-activity relationships (SARs) focused on the triazole *N*-substituent, the length of the carbon linker between the pantoyl group and the triazole ring, and the relative position of the two substituents on the ring. It was found that the triazole 1,4-substitution pattern with a simple methylene linker between the triazole and pantoyl moieties is preferred for optimal antiplasmodial activity, and yields compounds with low nanomolar antiplasmodial activity, *e.g.* **1** and **2**, which are proposed to mimic compounds with an inverted amide group (Figure 1).^12, 21^ We report herein the synthesis and antiplasmodial activity of novel pantothenamide analogs containing diverse heteroaromatic rings instead of the labile amide moiety, several of which show similar potency to triazole compound **1**. In addition, we report selectivity indices, time-of-kill, Caco-2 cell permeability, metabolic stability, and the first *in vivo* evaluation, of heteroaromatic ring-modified pantothenamides.

## 2. RESULTS

### 2.1 Synthesis

Heteroaromatic rings are attractive groups due to their hydrolytic stability, their hydrogen-bond donors and acceptors (as found in amides), and their ubiquity in approved drugs.^22^ The specific target compounds (**3a-v**, Schemes 1-5) were selected based on synthetic accessibility and diversity. A feature common to **1**, **2** and our synthetic targets is the pantoyl moiety. We therefore envisaged a synthetic scheme with the corresponding heteroaromatic ring-containing amine undergoing a condensation with D-pantolactone (Scheme 1). The desired primary amines were prepared either directly or via the *tert*-butyloxycarbonyl (Boc) protected amine.

**Scheme 1.**
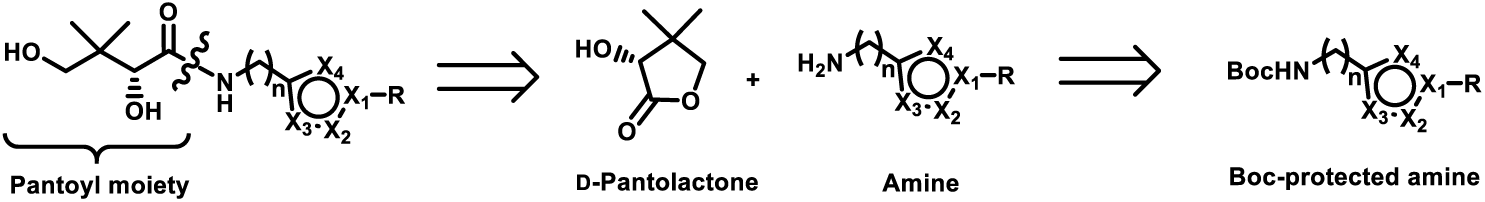
Retrosynthetic approach to the target heteroaromatic pantothenamide analogs **3a-v**. X_1_, X_2_, X_3_ and X_4_ are C, N, O or S atoms; R is a butyl, pentyl, hexyl, or phenethyl group; and n = 1 or 2.

**Scheme 2.**
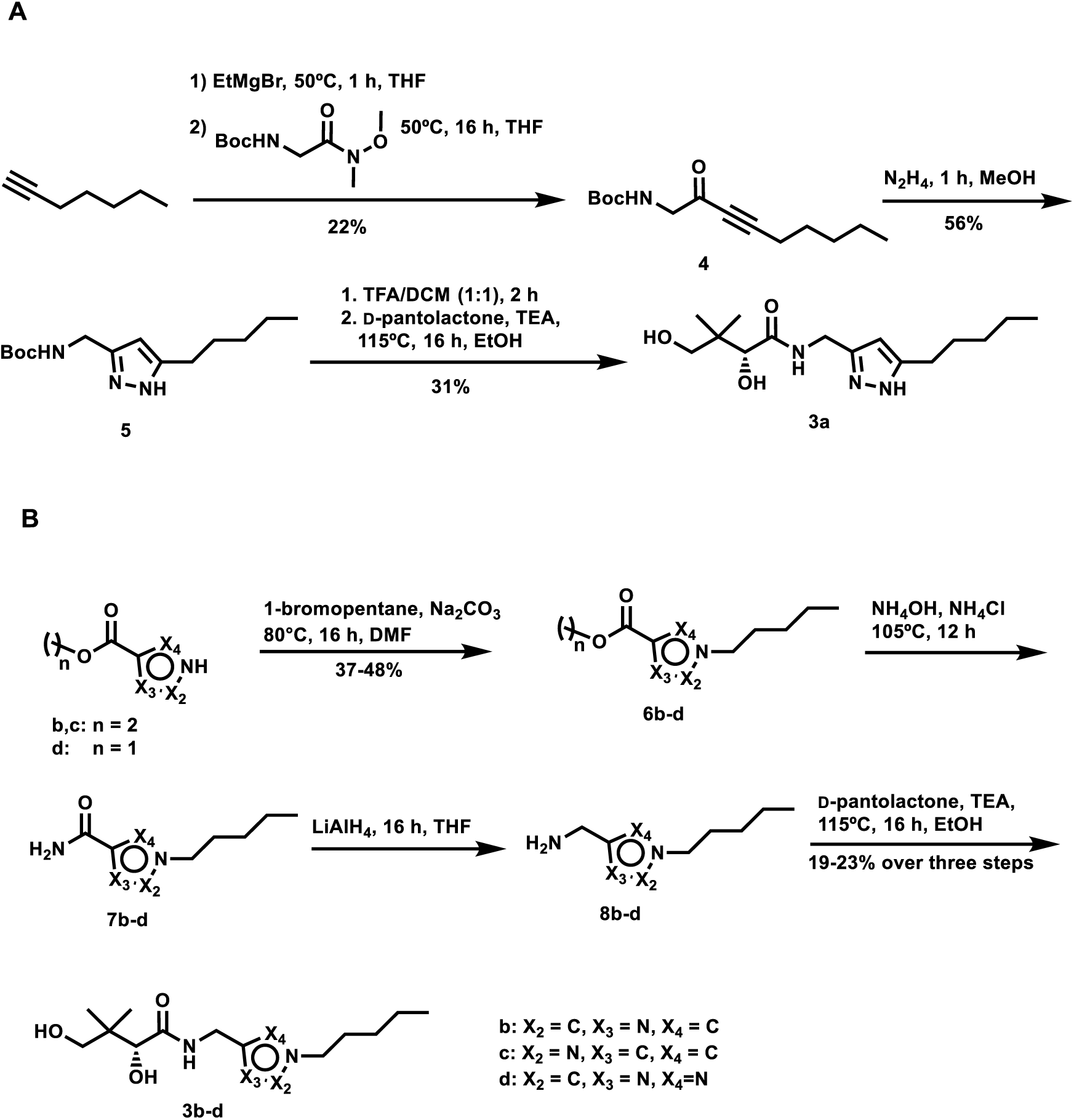
Synthesis of compound **3a** (**A**) and compounds **3b-d** (**B**). DCM: dichloromethane; DMF: *N,N*-dimethylformamide; TEA: triethylamine; TFA: trifluoroacetic acid; THF: tetrahydrofuran.

**Scheme 3.**
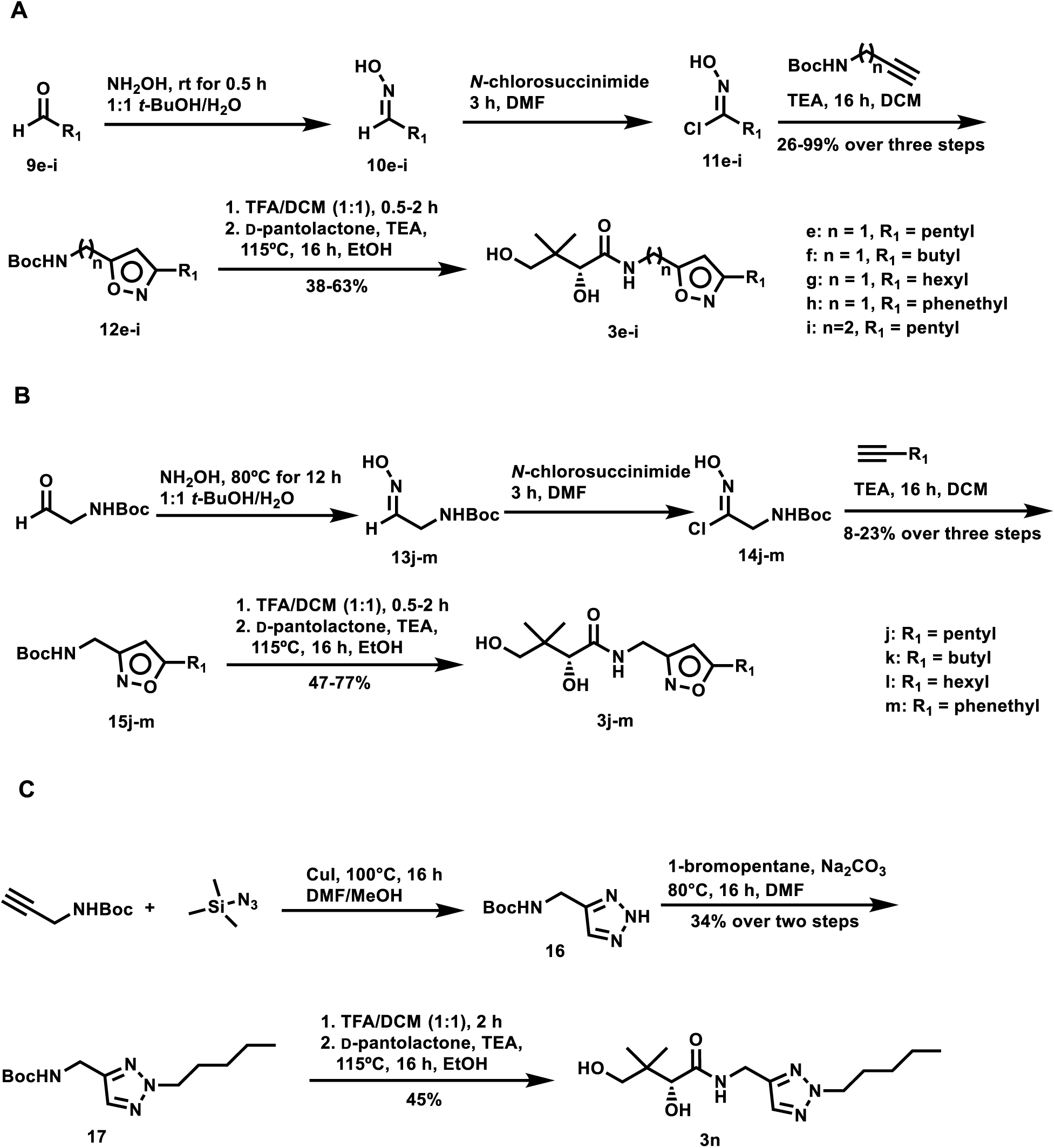
Synthesis of compounds **3e-i** (**A**), compounds **3j-m** (**B**), and compound **3n** (**C**). DCM: dichloromethane; DMF: *N,N*-dimethylformamide; TEA: triethylamine; TFA: trifluoroacetic acid.

**Scheme 4.**
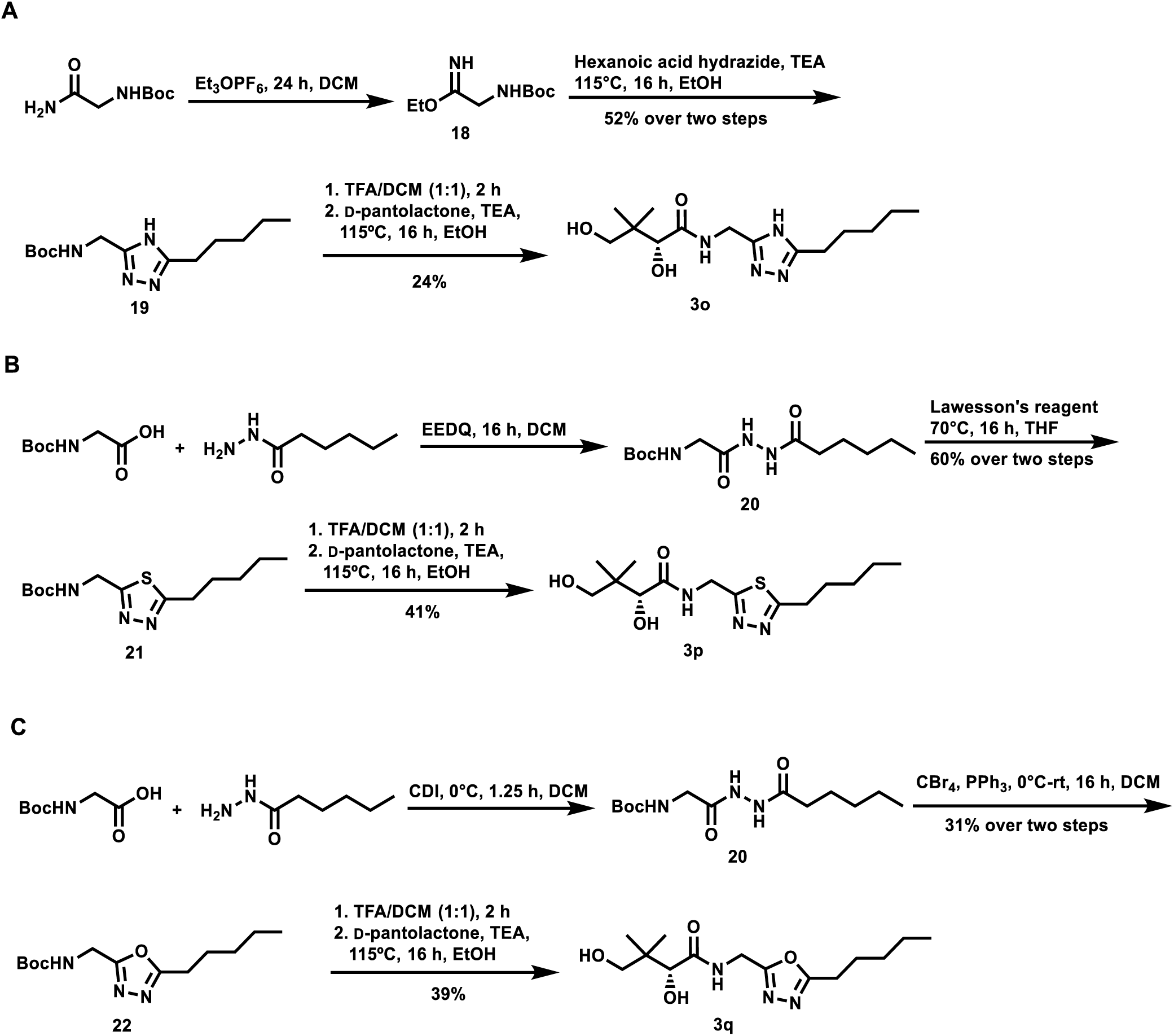
Synthesis of compound **3o** (**A**), compound **3p** (**B**), and compound **3q** (**C**). CDI: carbonyldiimidazole; DCM, dichloromethane; EEDQ: 2-ethoxy-1-ethoxycarbonyl-1,2-dihydroquinoline; TEA: triethylamine; TFA: trifluoroacetic acid; THF, tetrahydrofuran.

**Scheme 5.**
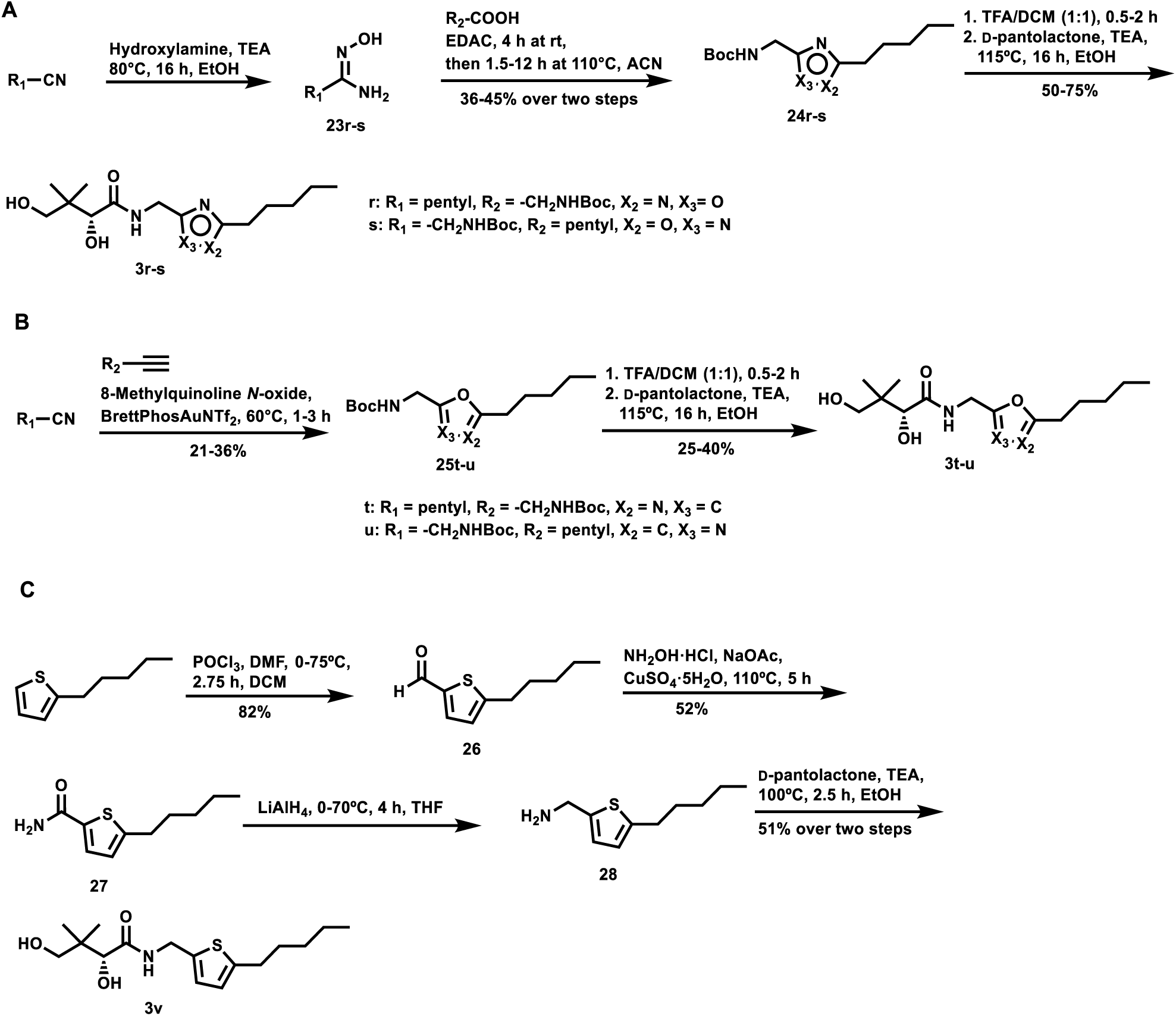
Synthesis of compounds **3r-s** (**A**), compounds **3t-u** (**B**), and compound **3v** (**C**). DCM, dichloromethane; DMF, *N,N*-dimethylformamide; EDAC: *N*-(3-dimethylaminopropyl)-*N’*-ethylcarbodiimide hydrochloride; TEA: triethylamine; TFA: trifluoroacetic acid.

The pyrazole derivative **3a** was assembled from 1-heptyne (Scheme 2A). Deprotonation of the terminal alkyne using ethylmagnesium bromide, followed by attack on the required Weinreb amide yielded compound **4**, which was condensed with hydrazine to afford the pyrazole **5**. Finally, Boc-deprotection produced the amine used to open the D-pantolactone ring, yielding the desired **3a**.

Compounds **3b-d** were prepared from commercial esters already containing the desired heteroaromatic ring (Scheme 2B). The pentyl group was introduced at the ring N-H with 1-bromopentane. The resulting compounds **6b-d** were then converted to the terminal amides **7b-d** with ammonium hydroxide and ammonium chloride, before reduction with lithium aluminum hydride to yield amines **8b-d**. The reaction of each of these amines with D-pantolactone generated **3b-d**.

Synthesis of compounds **3e-i** (Scheme 3A) first entailed producing oximes **10e-i** by condensation of commercial aldehydes **9e-i** with hydroxylamine. The oximes **10e-i** were next chlorinated using *N*-chlorosuccinimide to form **11e-i**, followed by cyclization with the desired alkyne at room temperature. Boc-deprotection of isoxazoles **12e-i** before addition to D-pantolactone afforded the desired **3e-i**. Compounds **3j-m** were produced in a similar manner, but from *N*-Boc-2-aminoacetaldehyde and using various alkynes (Scheme 3B).

To access compound **3n** (Scheme 3C), *N*-Boc-propargylamine and trimethylsilyl azide were cyclized to the 1,2,3-triazole **16**. Compound **16** was next extended with 1-bromopentane, before Boc-deprotection and condensation with D-pantolactone to produce **3n**.

To synthesize compound **3o** (Scheme 4A), *N*-Boc-glycinamide was first converted to imidate **18** in the presence of triethyloxonium hexafluorophosphate. Compound **18** was next cyclized with hexanoic acid hydrazide to generate the 1,2,4-triazole **19**. Finally, Boc-deprotection followed by D-pantolactone ring-opening afforded compound **3o**.

1,3,4-Thiadiazole **3p** was synthesized in three steps (Scheme 4B). First, *N*-Boc-glycine was coupled to hexanoic acid hydrazide in the presence of 2-ethoxy-1-ethoxycarbonyl-1,2-dihydroquinoline (EEDQ) to yield compound **20**, before thiation with Lawesson’s reagent and cyclization to generate the 1,3,4-thiadiazole **21**.^23^ Boc-deprotection and D-pantolactone ring opening yielded **3p**. The 1,3,4-oxadiazole **3q** was similarly prepared (Scheme 3C). After coupling *N*-Boc-glycine and hexanoic acid hydrazide to yield **20**, direct cyclization to the 1,3,4-oxadiazole **22** was facilitated by CBr_4_ and PPh_3_. Finally, Boc-deprotection of **22**, followed by condensation with D-pantolactone, afforded **3q**.

Compounds **3r** and **3s**, which differ only by swapped heteroaromatic N and O atoms (Scheme 5A), were accessed by first reacting the desired nitrile with hydroxylamine. Next, condensation of the resulting amino oximes **23r-s** with the desired carboxylic acid in the presence of a carbodiimide, generated the 1,2,4-oxadiazoles **24r-s**. Boc-deprotection, followed by D-pantolactone ring-opening yielded **3r-s**.

Oxazoles **3t-u** were prepared in three steps (Scheme 5B). The first step was the [2+2+1] annulation of a terminal alkyne, a nitrile, and an oxygen atom from the oxidant 8-methylquinoline *N*-oxide, catalyzed by BrettPhosAuNTf_2_.^24^ This was followed by Boc-deprotection and D-pantolactone ring opening to yield **3t-u**.

The thiophene **3v** was synthesized in four successive steps starting from 2-pentyl thiophene (Scheme 5C). Formylation via the Vilsmeier-Haack reaction, amidation, and reduction of the amide to the amine, followed by ring-opening of D-pantolactone afforded the target compound **3v**.

### 2.2 Antiplasmodial activity *in vitro*

Compounds **3a-v** were evaluated for their antiplasmodial activity against intraerythrocytic *P. falciparum*. Isoxazoles **3e-f**, **3j-l** and thiadiazole **3p** were found to inhibit proliferation of *P. falciparum* with IC_50_ values in the nanomolar range, with compound **3k** showing comparable activity to lead compound **1** (Table 1).

**Table 1.**
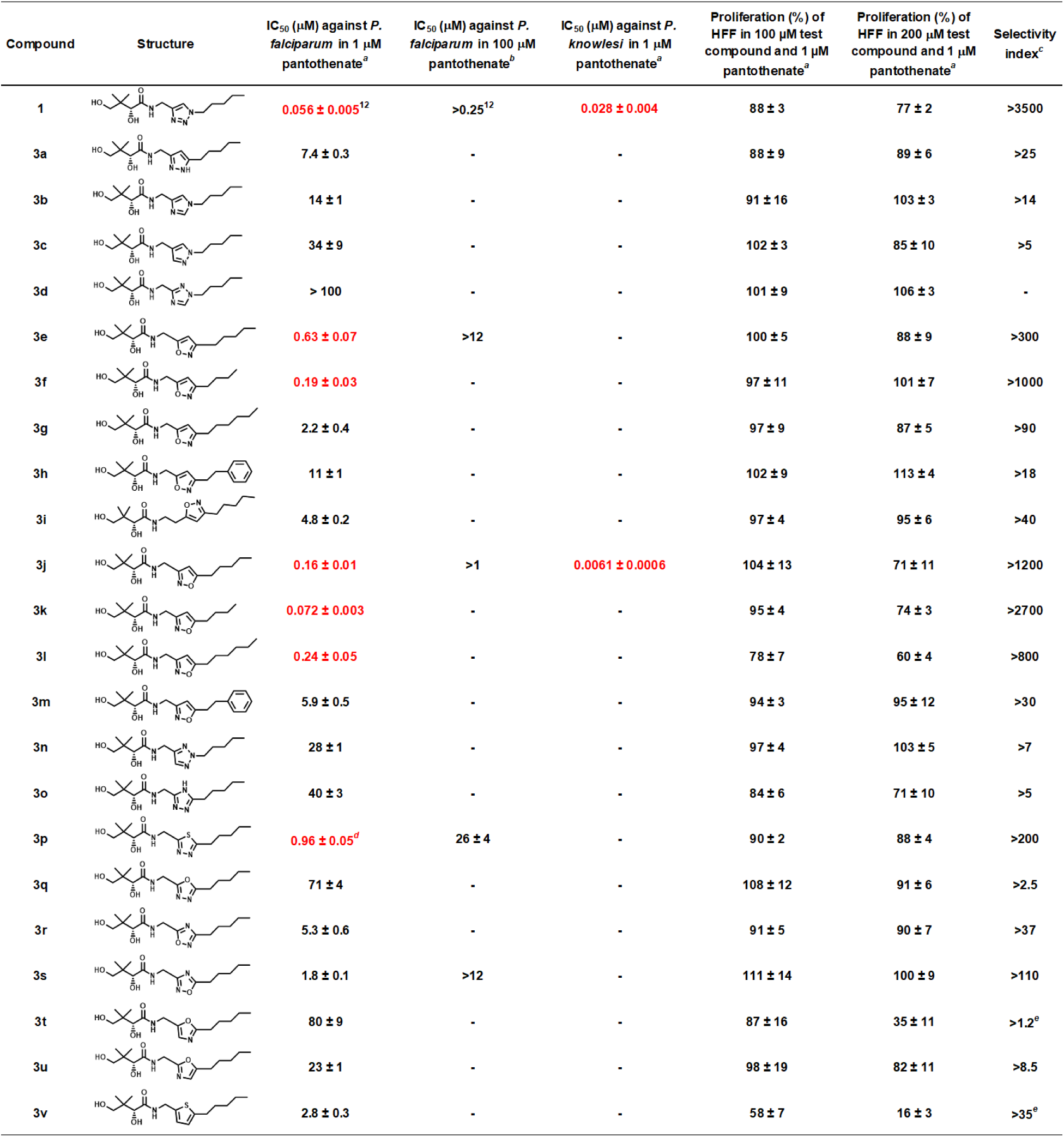
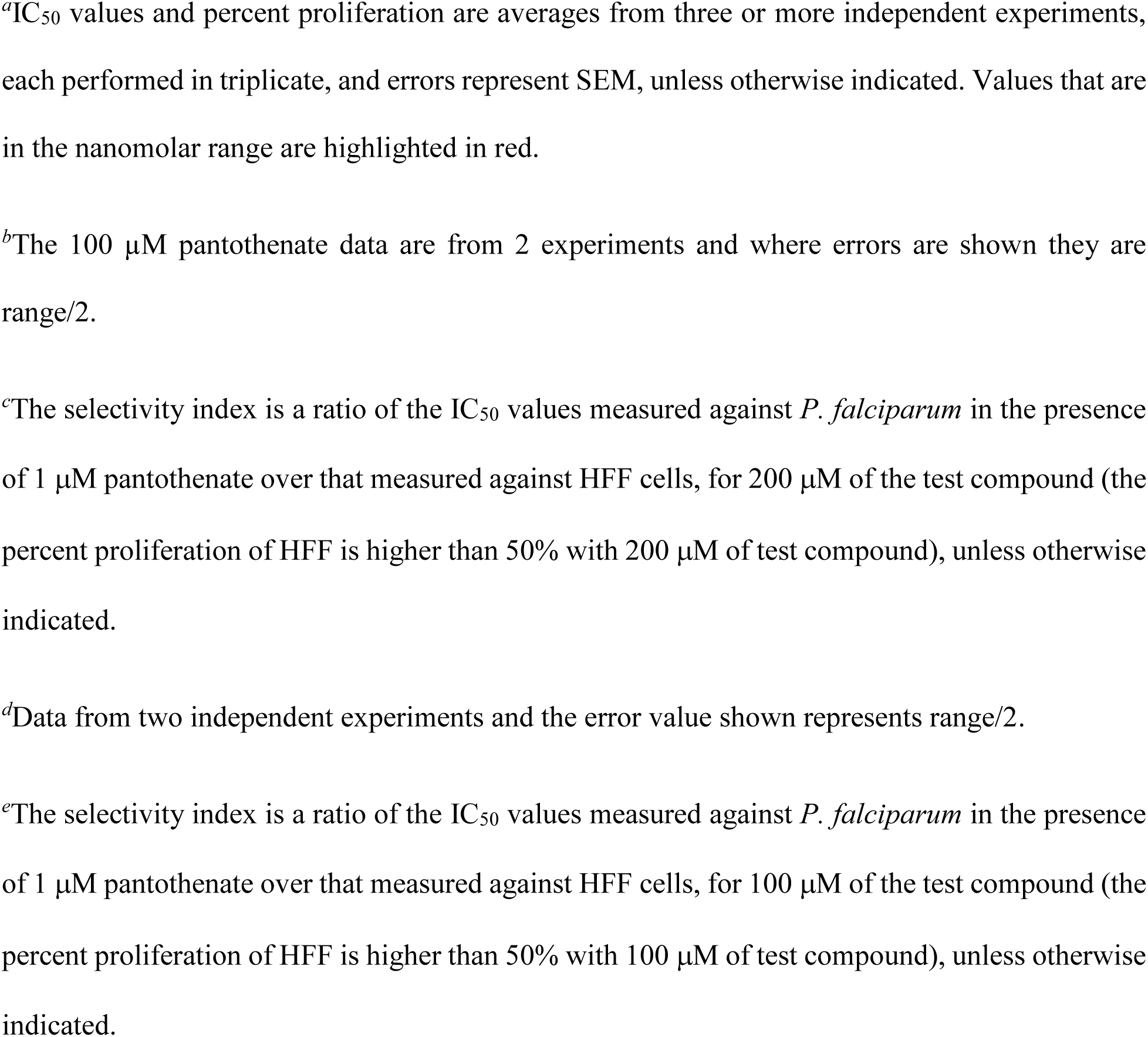
Effect of compounds **3a-v** on proliferation of intraerythrocytic *P. falciparum*, *P. knowlesi,* and human foreskin fibroblast (HFF) cells in growth medium containing pantetheinases and 1 or 100 μM added pantothenate.

| Compound | Structure | IC<sub>50</sub> (μM) against *P. falciparum* in 1 μM pantothenate<sup>a</sup> | IC<sub>50</sub> (μM) against *P. falciparum* in 100 μM pantothenate<sup>b</sup> | IC<sub>50</sub> (μM) against *P. knowlesi* in 1 μM pantothenate<sup>a</sup> | Proliferation (%) of HFF in 100 μM test compound and 1 μM pantothenate<sup>a</sup> | Proliferation (%) of HFF in 200 μM test compound and 1 μM pantothenate<sup>a</sup> | Selectivity index<sup>c</sup> |
| --- | --- | --- | --- | --- | --- | --- | --- |
| **1** | | 0.056 ± 0.005<sup>12</sup> | >0.25<sup>12</sup> | 0.028 ± 0.004 | 88 ± 3 | 77 ± 2 | >3500 |
| **3a** | | 7.4 ± 0.3 | - | - | 88 ± 9 | 89 ± 6 | >25 |
| **3b** | | 14 ± 1 | - | - | 91 ± 16 | 103 ± 3 | >14 |
| **3c** | | 34 ± 9 | - | - | 102 ± 3 | 85 ± 10 | >5 |
| **3d** | | > 100 | - | - | 101 ± 9 | 106 ± 3 | - |
| **3e** | | 0.63 ± 0.07 | >12 | - | 100 ± 5 | 88 ± 9 | >300 |
| **3f** | | 0.19 ± 0.03 | - | - | 97 ± 11 | 101 ± 7 | >1000 |
| **3g** | | 2.2 ± 0.4 | - | - | 97 ± 9 | 87 ± 5 | >90 |
| **3h** | | 11 ± 1 | - | - | 102 ± 9 | 113 ± 4 | >18 |
| **3i** | | 4.8 ± 0.2 | - | - | 97 ± 4 | 95 ± 6 | >40 |
| **3j** | | 0.16 ± 0.01 | >1 | 0.0061 ± 0.0006 | 104 ± 13 | 71 ± 11 | >1200 |
| **3k** | | 0.072 ± 0.003 | - | - | 95 ± 4 | 74 ± 3 | >2700 |
| **3l** | | 0.24 ± 0.05 | - | - | 78 ± 7 | 60 ± 4 | >800 |
| **3m** | | 5.9 ± 0.5 | - | - | 94 ± 3 | 95 ± 12 | >30 |
| **3n** | | 28 ± 1 | - | - | 97 ± 4 | 103 ± 5 | >7 |
| **3o** | | 40 ± 3 | - | - | 84 ± 6 | 71 ± 10 | >5 |
| **3p** | | 0.96 ± 0.05<sup>d</sup> | 26 ± 4 | - | 90 ± 2 | 88 ± 4 | >200 |
| **3q** | | 71 ± 4 | - | - | 108 ± 12 | 91 ± 6 | >2.5 |
| **3r** | | 5.3 ± 0.6 | - | - | 91 ± 5 | 90 ± 7 | >37 |
| **3s** | | 1.8 ± 0.1 | >12 | - | 111 ± 14 | 100 ± 9 | >110 |
| **3t** | | 80 ± 9 | - | - | 87 ± 16 | 35 ± 11 | >1.2<sup>e</sup> |
| **3u** | | 23 ± 1 | - | - | 98 ± 19 | 82 ± 11 | >8.5 |
| **3v** | | 2.8 ± 0.3 | - | - | 58 ± 7 | 16 ± 3 | >35<sup>e</sup> |

<sup>a</sup>IC<sub>50</sub> values and percent proliferation are averages from three or more independent experiments, each performed in triplicate, and errors represent SEM, unless otherwise indicated. Values that are in the nanomolar range are highlighted in red.
<sup>b</sup>The 100 µM pantothenate data are from 2 experiments and where errors are shown they are range/2.
<sup>c</sup>The selectivity index is a ratio of the IC<sub>50</sub> values measured against *P. falciparum* in the presence of 1 µM pantothenate over that measured against HFF cells, for 200 µM of the test compound (the percent proliferation of HFF is higher than 50% with 200 µM of test compound), unless otherwise indicated.
<sup>d</sup>Data from two independent experiments and the error value shown represents range/2.
<sup>e</sup>The selectivity index is a ratio of the IC<sub>50</sub> values measured against *P. falciparum* in the presence of 1 µM pantothenate over that measured against HFF cells, for 100 µM of the test compound (the percent proliferation of HFF is higher than 50% with 100 µM of test compound), unless otherwise indicated.

Interestingly, individual replacement of each of the ring N atoms of compound **1** (positions X_1_, X_2_ or X_3_, Scheme 1) with a C atom leads to a significant decrease in activity (see **3a-c**), with the largest effect associated with replacement at the X_3_ position. Consistent with this observation, 1,3,4-thiadiazole **3p** also shows higher potency than thiophene **3v**. Furthermore, as evidenced by the higher activity of **3e-f** and **3j** relative to **3a**, replacing the ring N atom with an O atom is beneficial at both the X_2_ and X_3_ positions. Overall the preferred atoms are O > N > C for both positions, correlating with electronegativity and size. By contrast, **3e**, **3j**, **3a** and **3b** are more potent than **3r**, **3s**, **3o**, and **3d**, respectively, implying that C is preferred over N at the X_4_ position. Additionally, the activity trend of **3p > 3a > 3o > 3q** suggests a preference for S > C > N > O at position X_4_. The low tolerance for an O atom at this position is also evidenced by the high IC_50_ values measured for compounds **3t-u**. Interestingly, the order of atom preference at X_4_ seems to correlate inversely with the electronegativity of the atom.^25^

The substituent at X_1_ was found to also influence activity, as revealed by the SARs observed for the **3e-h** series, and the **3j-m** series. It is apparent that a four-carbon chain is preferred over a pentyl or a hexyl group. This contrasts with the trend previously observed for the corresponding triazole pantothenamide analogs for which there is a very slight preference for five- and six-carbon chains over four.^12^ The diverging trend may arise because of the smaller ring size of the triazole (requiring a longer substituent to compensate), compared to the isoxazole. Similar to what was previously reported for triazole pantothenamide analogs,^21^ alkyl chains are preferred over the phenethyl group associated with high potency in unmodified pantothenamides,^5^ with phenethyl analogs **3h** and **3m** showing lower activity than **3e-g** and **3j-l**, respectively. A one-carbon linker between the pantoyl and isoxazole moieties is also favored over a two-carbon linker, at least within derivatives containing a 5-carbon chain (**3e** shows improved activity relative to **3i**), analogous to the optimal spacing observed in the triazole series. This is consistent with our previous hypothesis that compounds with a one-carbon linker better mimic pantothenamides with the labile amide bond inverted, while those containing a two-carbon linker simulate standard pantothenamides.^21^

Previously, we have shown that the mechanism of action of compound **1** against *P. falciparum* involves bioactivation by pantothenate kinase (PanK), phosphopantetheine adenylyltransferase (PPAT), and dephospho-CoA kinase (DPCK) to form a CoA derivative that is hypothesized to inhibit CoA-utilizing enzymes.^7^ To investigate whether the newly discovered antiplasmodial isoxazole and thiadiazole derivatives exert their activity via a similar mechanism, the antiplasmodial activity of a selection of compounds (**3e**, **3j** and **3p)** was also evaluated in the presence of excess pantothenate (100 µM). Similar to compound **1**, a > 6-fold increase in the IC_50_ values of all three compounds was observed upon the addition of excess pantothenate (Table 1), consistent with the isoxazoles and thiadiazole derivatives also competing with pantothenate as substrates of PanK.

Two compounds – **1** and **3j** – were additionally evaluated for their ability to inhibit proliferation of *P. knowlesi*, a recognized cause of human malaria in South-East Asia.^26–27^ *P. knowlesi* is the closest relative of *P. vivax* (a parasite responsible for 50% of malaria cases in South-East Asia and the predominant malaria parasite in the Americas) that is amenable to *in vitro* continuous culturing.^1, 28^ Both compounds were found to inhibit proliferation of *P. knowlesi* at nanomolar concentrations. Interestingly, in contrast with the findings for *P. falciparum*, **3j** showed superior activity compared to **1** against this species (Table 1).

The selectivity of compounds **3a-v** was investigated by measuring proliferation of human foreskin fibroblast (HFF) cells (a low-passage cell line) in the presence of each compound. None of the compounds inhibited proliferation of the cells by more than 50% at 100 µM and only compounds **3t** and **3v** showed greater than 50% inhibition at a concentration of 200 µM (Table 1). The selectivity indices of the best parasite inhibitors are therefore well above 200 (Table 1).

### 2.3 Absorption and metabolism *in vitro*

Having demonstrated that compounds **1** and **3j** show selective nanomolar antiplasmodial activity against *P. falciparum* and *P. knowlesi*, we next investigated whether they are likely to be absorbed from the intestine by measuring their apparent permeability (P_app_) across confluent, differentiated Caco-2 cell monolayers.^29^ Compound **3j** permeated Caco-2 cell monolayers in the apical-to-basolateral (A→B) direction, with a P_app_ higher than that determined for propranolol – a highly permeable reference compound (Table 2).^29^ Compound **1** showed moderate permeability, with a P_app_ approximately 4-fold lower than that determined for compound **3j** (Table 2). Neither compound **1** nor **3j** was found to compromise the integrity of the Caco-2 monolayer, as evidenced by the exclusion of lucifer yellow^30^, a non-permeable reference compound.

**Table 2.** Apparent permeability (P_app_) of compounds **1** and **3j**, as well as reference compounds across Caco-2 cell monolayers assessed in the apical to basolateral (A→B) direction.

| Compound | A→B $P_{app}^a$ ($10^{-6}$ cm/s) |
| --- | --- |
| Lucifer yellow | < 1 |
| Propranolol | $49 \pm 2.3$ |
| **1** | $13 \pm 1.8$ |
| **3j** | $55 \pm 0.6$ |

<sup>a</sup>Values reported are the mean ± standard deviation of measurements from three transwells.

To explore the susceptibility of compounds **1** and **3j** to hepatic metabolism, stability of the compounds in the presence of liver microsomes was measured. Minimal degradation of compound **1** was observed during incubation with mouse or human microsomes (<15% over 60 min), corresponding to an *in vitro* half-life (t_1/2_) of > 255 min (Table 3) and a low *in vitro* intrinsic clearance (CL_int, *in vitro*_; < 7 µL/min/mg protein). In the presence of mouse microsomes, some NADPH-independent degradation (20-30% degradation over 60 min) and NADH-dependent degradation of compound **3j** was observed, with an estimated t_1/2_ of 33 min. The compound was, however, more stable in human than mouse microsomes. A t_1/2_ of 152 min was measured in the presence of human microsomes (CL_int, *in vitro*_ = 11 µL/min/mg protein).

**Table 3.** Stability of compounds **1** and **3j** in mouse and human liver microsomes *in vitro*.

| Compound | Microsome species | $t_{1/2}$<br>(min) | $CL_{\text{int, in vitro}}$<br>($\mu\text{L}/\text{min}/\text{mg protein}$) |
| --- | --- | --- | --- |
| **1** | Mouse | >255 | <7 |
|  | Human | >255 | <7 |
| **3j** | Mouse | 33 | 53 |
|  | Human | 152 | 11 |

### 2.4 Antiplasmodial activity *in vivo*

Encouraged by the permeability and stability of compounds **1** and **3j**, the *in vivo* antiplasmodial activity of these compounds was investigated using a mouse model of malaria. Female BALB/c mice were infected with *P. berghei,* and when the average parasitemia reached ∼1%, they were administered intraperitoneally with 100 mg/kg/day of compound **1** or **3j**, or the corresponding solvent control, for two days. Unfortunately, neither of the compounds were found to inhibit proliferation of *P. berghei* in mice, with the average parasitemia increasing at a similar rate in all treatment groups (Figure 2A).

**Figure 2.**
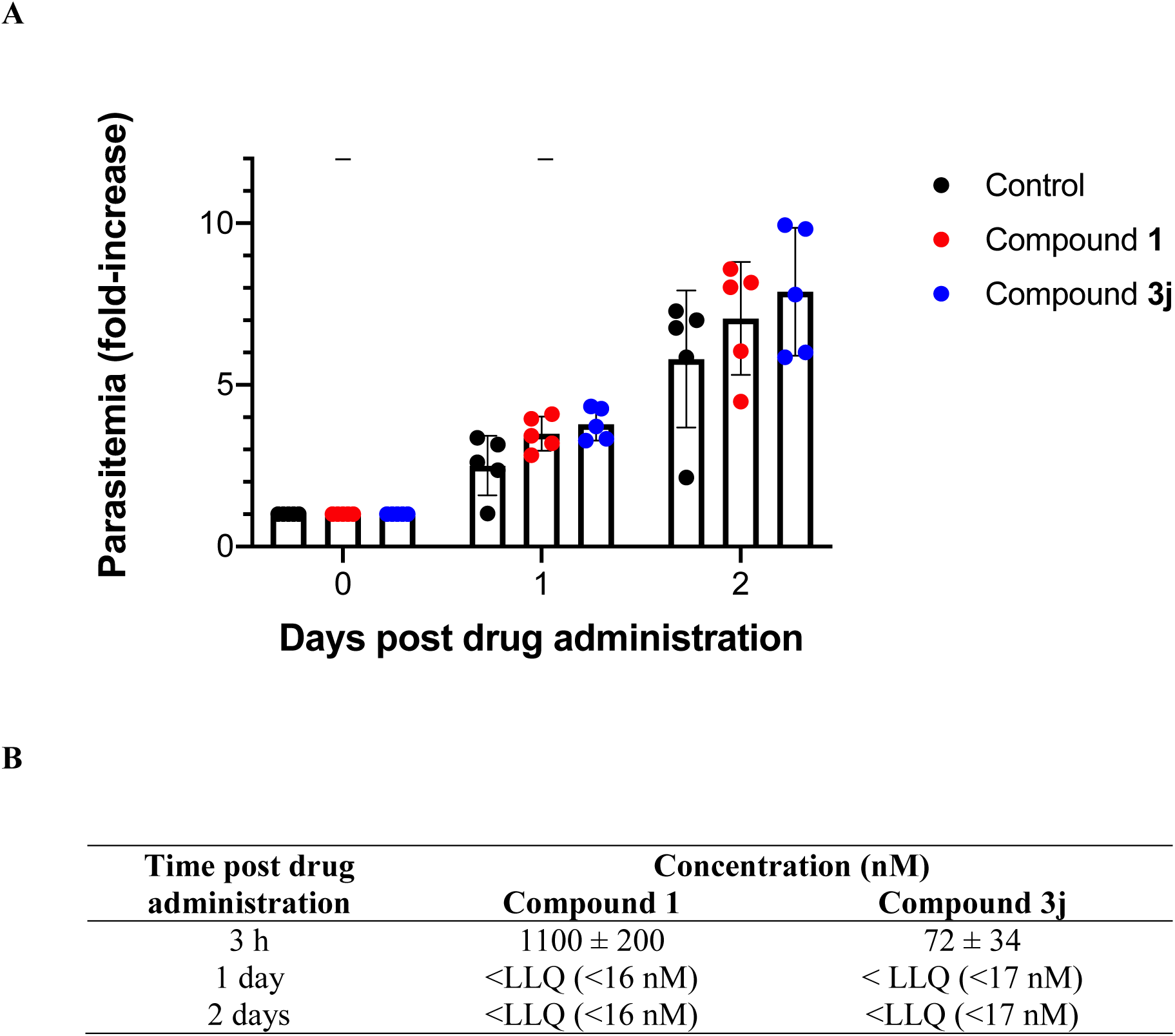
Effect of compounds **1** and **3j** on the proliferation of *P. berghei* in female BALB/c mice **(A**), and whole-blood concentrations of each compound (**B**), following IP administration with 100 mg/kg/day for two days. Groups of five *P. berghei*-infected mice were administered with solvent (black circles), compound **1** (red circles) or compound **3j** (blue circles) at day 0 and day 1 (as indicated by the arrows in A). Bars represent the average fold-increase in parasitemia in each treatment group and error bars denote standard deviations. Blood samples were collected 3 h after the first dose, and on day 1 blood samples were collected immediately before the second dose. The whole-blood concentrations measured in each of the five mice in each treatment group are averaged in panel B. Errors represent standard deviations. <LLQ: concentrations were below the limit of quantitation of 16 or 17 nM (for compounds **1** and **3j**, respectively).

Three and twenty-four hours after each dose of compound was administered to mice, blood samples were collected and the whole-blood concentration of the test compound was determined in an attempt to shed light on the reasons for the lack of antiplasmodial activity *in vivo*. Three hours after the first dose, compound **1** was detected in whole-blood at a concentration of 1.1 ± 0.2 µM (20 times the IC_50_ value against *P. falciparum* reported in Table 1), but within 24 h of drug administration, this decreased to below the limit of quantitation (16 nM) (Figure 2B). Twenty-four hours after the second dose, the whole-blood concentration remained below the limit of quantitation. Fifteen times less compound **3j** (72 ± 34 nM; Figure 2B) was detected in blood 3 h after the first dose relative to that detected of compound **1**, a finding consistent with the higher *in vitro* intrinsic clearance observed in the microsome stability assays. In contrast with compound **1**, the concentration of compound **3j** detected at 3 h did not exceed the IC_50_ value determined against *P. falciparum in vitro* (Table 1). As for compound **1**, the concentration of **3j** in whole blood 24 h after drug administration was below the limit of quantitation (Figure 2B).

To investigate whether the observed exposures were likely to be sufficient for an antiplasmodial effect or could instead provide an explanation for why the antiplasmodial activity in mice was not greater, we determined the contact time necessary for compounds **1** and **3j** to inhibit proliferation of *P. falciparum in vitro*. Ring-stage *P. falciparum*-infected erythrocytes – the same stage of the parasite that was initially exposed to the compounds in the *in vivo P. berghei* experiments – were exposed for pre-determined periods to compounds **1** and **3j** before the compounds were removed and proliferation was monitored over a total of 96 h. As shown in Figure 3, a 10-h exposure to the compounds was insufficient for nanomolar antiplasmodial activity. At least 24 h of contact was needed to observe an antiplasmodial effect, and a full effect required a 48 h exposure to the compounds. These results, together with the whole-blood concentrations detected 3 h and 24 h post drug administration, are consistent with compounds **1** and **3j** not being available at a sufficiently high concentrations for long enough to irreversibly inhibit parasite proliferation under the conditions of the *in vivo* experiment.

**Figure 3.**
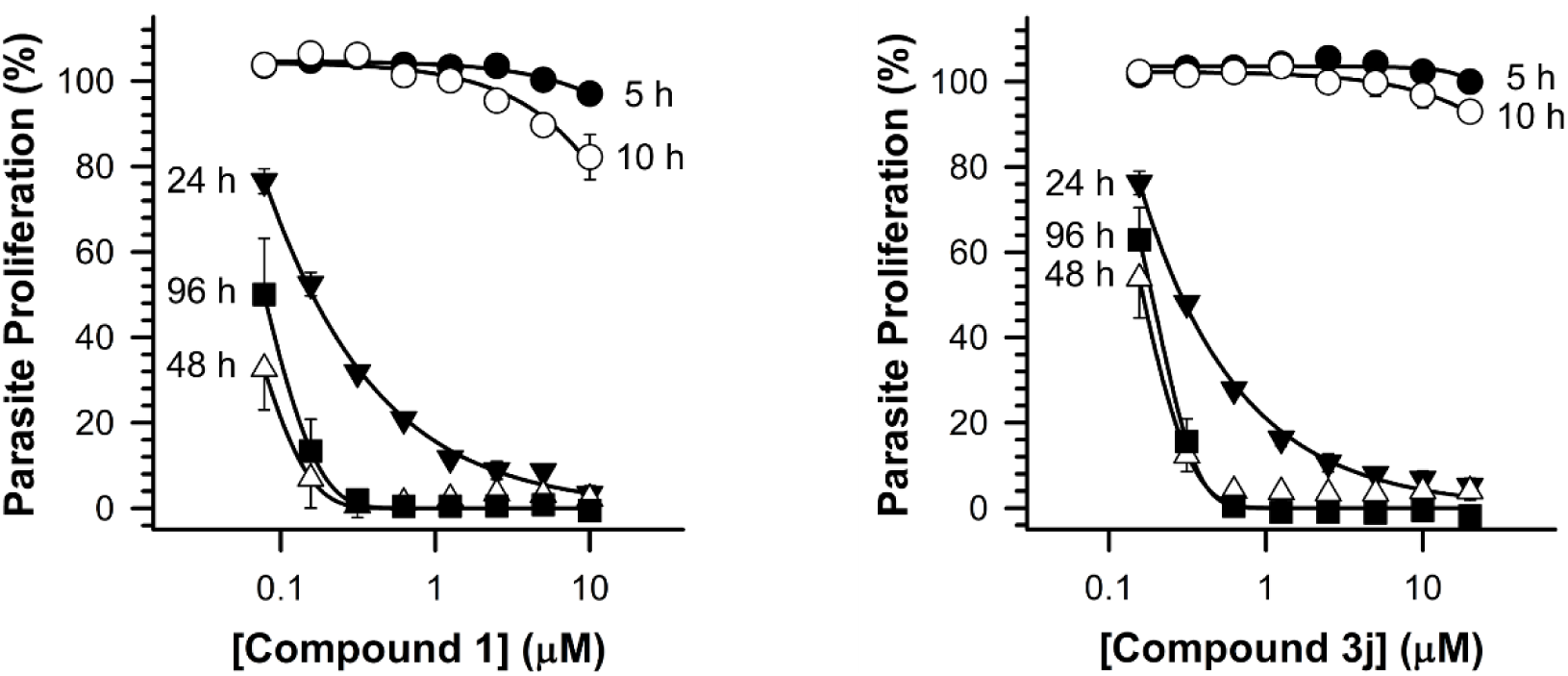
Effect of exposure time on the antiplasmodial activity of **1** and **3j** *in vitro*. *P. falciparum-*infected erythrocytes were incubated in the presence of compound **1** or **3j** for 5 h (closed circles), 10 h (open circles), 24 h (closed triangles), 48 h (open triangles) or 96 h (closed squares) before the compounds were removed and parasite proliferation was monitored over a total of 96 h. Values are averages from three independent experiments that were initiated with “triple-synchronized” ring stage parasites, each performed in triplicate. Error bars represent SEM and are not visible when smaller than the symbols.

Schalkwijk *et al*. (2019) previously showed that pantothenamide analogs with an inverted amide bond (*e.g.* **CXP18.6-013**, Figure 1) are bioactivated to the corresponding CoA analogs by uninfected erythrocytes, which renders erythrocytes inhospitable to *P. falciparum*.^8^ They proposed that this phenomenon may lead to accumulation of the compounds within erythrocytes (with the CoA analogs being less permeable to membranes than pantothenamides) and could explain why parasite elimination in mice continues after the concentration of the inverted pantothenamide is undetectable in plasma. To explore further the reasons for the lack of antiplasmodial activity of compounds **1** and **3j** in mice, we examined whether or not pre-exposure to compounds **1** and **3j** similarly affects the ability of erythrocytes to support proliferation of *P. falciparum*. Uninfected erythrocytes were incubated with compounds **1**, **3j**, chloroquine (an antimalarial shown previously to accumulate in uninfected erythrocytes),^8, 31^ dihydroartemisinin (an antimalarial that does not accumulate in uninfected erythrocytes),^8^ or the corresponding solvent control. The pre-treatment was performed as described before,^8^ prior to mixing with enriched *P. falciparum*-infected erythrocytes (parasitemia >95%). Proliferation of *P. falciparum* in erythrocytes pre-treated with 100 µM chloroquine was inhibited in some experiments. However, variability between erythrocyte batches was observed and the averaged inhibition was not statistically significant (Figure 4, 95% confidence intervals, CI, include 100). Proliferation of parasites in erythrocytes treated with solvent (0.1%, v/v, dimethyl sulfoxide, DMSO), dihydroartemisinin (25 µM), compound **1** (100 µM), or compound **3j** (100 µM), was unaffected (Figure 4). These data are consistent with exposure of uninfected erythrocytes to compounds **1** and **3j** not contributing to their antiplasmodial activity, unlike what was proposed for the inverted pantothenamides. Hence, antiplasmodial activity of these compounds is unlikely to continue after they have been cleared.

**Figure 4.**
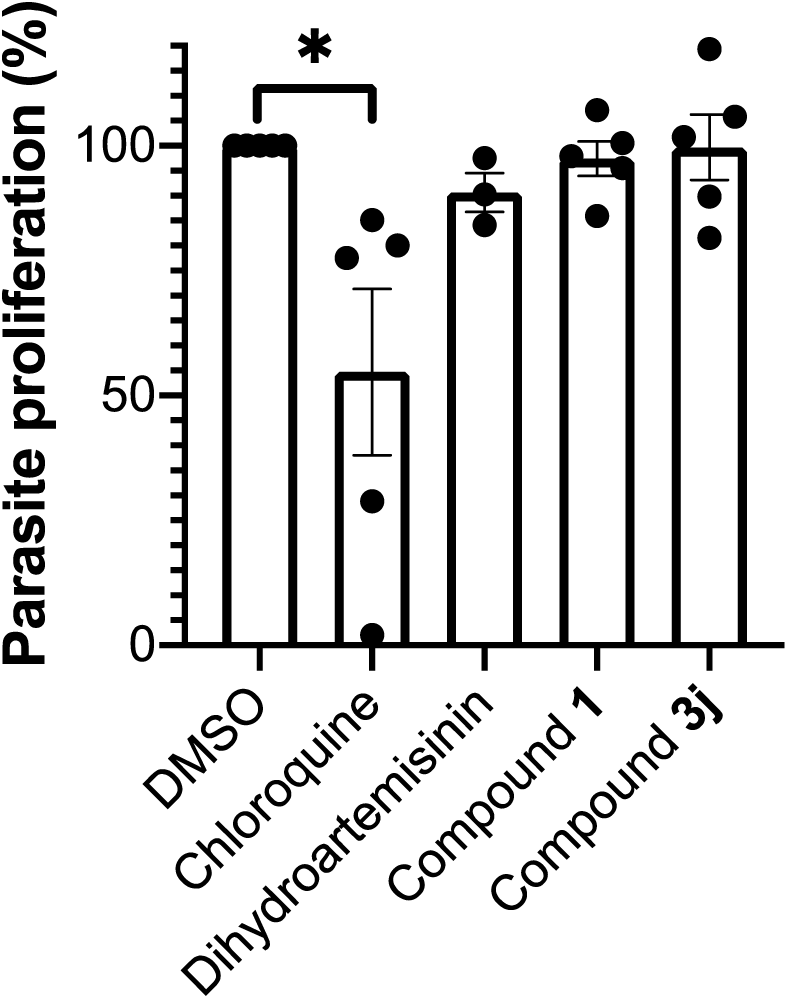
Effect of pre-exposure of erythrocytes to compounds **1**, **3j**, or reference compounds on the proliferation of *P. falciparum*. Bars show the average proliferation measured in three to five independent experiments (using blood from three different donors: black circles, donor 1; grey symbols, donor 2; white symbols, donor 3) each performed in duplicate. Error bars represent the standard error of the mean. Proliferation in erythrocytes pre-exposed to test or reference compounds is shown as a percentage of proliferation in erythrocytes exposed to 0.1% (v/v) DMSO, which was present in all treatments. Although proliferation was reduced following chloroquine pre-exposure of erythrocytes from certain donors, the averaged decrease was not statistically significant (95% CI: 8.5 – 100.9). Pre-exposure to dihydroartemisinin or compounds **1** or **3j** did not result in a reduction in parasite proliferation (95% CI: dihydroartemisinin = 73.9 – 107.4, compound **1**: 87.8 – 107.0, compound **3j** = 81.5 – 117.8).

## 3. DISCUSSION

A key goal of this study was to explore the range of heteroaromatic rings that could be used to replace the labile amide bond of pantothenamides and thereby stabilize these compounds in serum. Not all heteroaromatic rings present in compounds **3a-v** are good substitutes for the labile amide group. For example some, such as the oxadiazoles harbored by compounds **3q-s**, are missing a hydrogen atom that can mimic the amide N-H. It is interesting that among the new compounds reported here, the most potent antiplasmodials, isoxazoles **3e-f**,**j-l** and thiadiazole **3p**, may simulate pantothenamide analogs with an inverted amide bond.^13^ As shown in Figure 5, we propose that N-2 of **3e** and O-1 of **3j** may mimic the carbonyl oxygen of **CXP18.6-013**, while H-4 of both **3e** and **3j** may replace the N-H of **CXP18.6-013**. Compound **3p** is somewhat different; while its ring N-3 may imitate the carbonyl oxygen of **CXP18.6-013**, it does not have a hydrogen atom on the thiadiazole ring. The low-lying σ* orbital of the C-S bond is however known to interact with electron donors like amide N-H bonds do.^32^ It is therefore possible that this σ* orbital in **3p** may emulate the N-H of **CXP18.6-013**. Our postulate also correlates well with the findings that 1) a N atom is preferred at position X_1_ (numbering of Scheme 1), 2) N or O atoms are preferred at positions X_2_ and X_3_, and 3) C and S atom are preferred at position X_4_.

**Figure 5.**
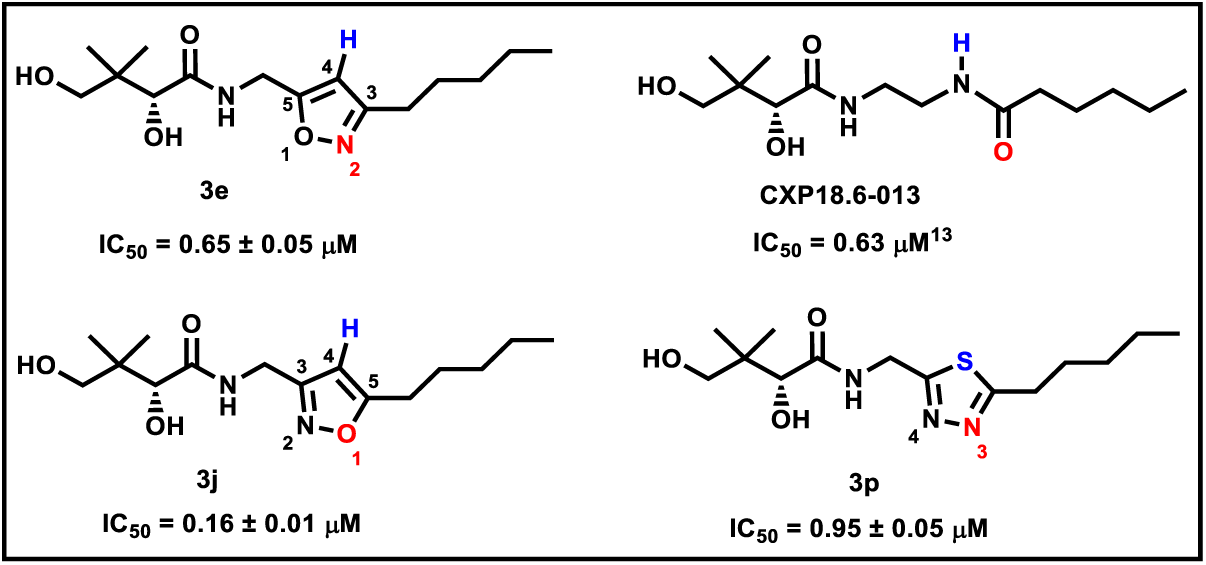
The isoxazole rings of **3e** and **3j** and the thiadiazole ring of **3p** may mimic the inverted amide bond of pantothenamide **CXP18.6-013**. Key electron donors are shown in red, while key electron acceptors are highlighted in blue.

The antiplasmodial activity of pantothenamides with an inverted amide has been optimized by replacing the pentyl substituent with various groups, of which a phenethyl moiety was especially promising.^13^ In contrast, our current results with isoxazoles, and those previously reported for triazole-modified pantothenamides,^12, 21^ reveal that replacement of the pentyl chain with a phenethyl moiety (as in compounds **3h** and **3m**) is detrimental to antiplasmodial activity. Here, the only substituent that proved advantageous over the pentyl chain was a linear butyl chain (as in compounds **3f** and **3k**), which interestingly was associated with marginally lower activity than the pentyl moiety in pantothenamides with an inverted amide,^13^ and comparable activity in the case of 1,2,3-triazole-modified pantothenamides.^12^ One possible explanation for the divergent SAR is that the antimetabolites of the different pantothenamide analogs may bind to their target(s) in slightly different orientations, or even act on distinct targets. Nonetheless, a more expansive exploration of additional substituents that have enhanced antiplasmodial activity in other pantothenamide analog series is warranted in the context of both triazole- and isoxazole-modified pantothenamides.

This study is the first to demonstrate nanomolar antiplasmodial activity of pantothenamide analogs against *Plasmodium* species infecting humans other than *P. falciparum*. The sensitivity of *P. knowlesi* to compounds **1** and **3j** suggests that proliferation of the closely related parasite *P. vivax* is likely to also be inhibited by these compounds. Interestingly, while compound **1** is more potent than **3j** against *P. falciparum*, the opposite is true for *P. knowlesi*. Such variations between the drug susceptibility of *P. knowlesi* and *P. falciparum* have previously been reported for other drug classes.^33^ The difference observed in this study could reflect an inherent difference in the biology of the two parasites, or may instead be a consequence of the non-identical conditions under which *P. knowlesi* and *P. falciparum* were cultured *in vitro* – *i.e. P. knowlesi* in the presence of both human serum and the serum substitute Albumax, while *P. falciparum* just in the presence of Albumax.

The potency and selectivity of the known triazole-modified pantothenamide **1**^12^ and of the potent isoxazoles reported here warranted an investigation into the absorption, distribution, metabolism and excretion (ADME) properties of these compounds. . The Caco-2 permeability (A→B), and stability to *in vitro* metabolic clearance of triazole **1** and isoxazole **3j** compare favorably to other modified pantothenamides, where direct comparisons are possible.^8–9^ Nonetheless, both **1** and **3j** failed to suppress *P. berghei* proliferation in mice. At 3 h post-administration, the whole-blood concentration of compound **3j**, was less than half the IC_50_ value measured against *P. falciparum* over 96 h, and this limited exposure of *P. berghei* is likely to explain the lack of activity of this compound *in vivo*. By contrast, compound **1** exceeded the IC_99_ concentration at 3 h post-administration. This difference in the whole-blood concentration of the compounds, which was consistent with the slower microsomal clearance observed for compound **1** *in vitro*, highlights the impact of simply changing the triazole ring to an isoxazole ring on the ADME properties. Contact time experiments revealed that exposure to compound **1** for 24 h was required for sub-micromolar inhibition of *P. falciparum* proliferation (and 48 h – a complete intraerythrocytic *P. falciparum* cycle – was required for full effect). By comparison, whole-blood serum measurements showed that the concentration of compound **1** had dropped below the limit of quantification by 24 h post-administration (one intraerythrocytic *P. berghei* cycle). Hence, insufficient exposure may also have contributed to the failure of compound **1**. An additional contributing factor to be considered is the high concentration of pantothenate in mouse serum, which can compete for metabolic transformation. This was recently determined to be approximately 2 µM,^8^ 6-fold higher than the concentration present in human serum, and 2-fold higher than that present in the *in vitro* proliferation assays performed to measure antiplasmodial activity.

Here, we did not observe a reduced capacity of erythrocytes pre-treated with compounds **1** and **3j** to support *P. falciparum* proliferation, as reported for another series of pantothenamide analogs.^8^ It will be important to not only understand why, but also to gauge the contribution of erythrocyte accumulation of pantothenamide analog antimetabolites to *in vivo* efficacy. Evaluating the Caco-2 permeability and microsome stability of some of the other potent pantothenamide analogs presented here (*e.g.* compounds **3f** and **3p**) will also be important for selecting/designing the best pre-clinical candidates.

## 4. CONCLUSION

Triazole, isoxazole and thiadiazole rings are among the most commonly used scaffolds in drugs.^22^ We report herein that, as established for the triazole ring,^18, 21^ replacement of the labile amide bond of pantothenamides with either an isoxazole or a thiadiazole ring considerably enhances potency towards *P. falciparum* in the presence pantetheinase. This is also the first report of heteroaromatic ring-containing pantothenamide analogs with activity against *P. knowlesi*. Furthermore, as for the triazole-modified pantothenamides, the isoxazoles and thiadiazoles possess high selectivity indices. Favorable ADME properties were determined for the triazole and isoxazole investigated, with isoxazole **3j** showing high Caco-2 permeability, consistent with good bioavailability, and triazole **1** showing low microsomal clearance. Although we observed here that compounds **1** and **3j** fail to suppress proliferation of *P. berghei in vivo*, the pharmacokinetic analyses and contact time experiments provide a benchmark for the compound profile required to achieve antiplasmodial activity in mice, which should facilitate lead optimization. In addition to isoxazole **3j**, several novel compounds reported here show potent activity against *P. falciparum* and provide additional leads for the development of novel antimalarials. Their straightforward synthesis further increases the appeal.

## EXPERIMENTAL SECTION

### Biology

#### Materials

All pantothenamide analogs tested were initially dissolved to a concentration of 200 mM in DMSO. The final concentration of DMSO in *in vitro* proliferation assays never exceeded 0.1% (v/v). Reagents were from Sigma-Aldrich, unless otherwise specified. Gentamicin and Albumax II were purchased from Thermo Fisher Scientific.

### Cell culture

*P. falciparum* (3D7 strain) parasites were maintained *in vitro* essentially as described previously,^12^ in human O^-^ erythrocytes suspended in HEPES- and Glutamax^TM^-supplemented RPMI 1640 (Thermo Fisher Scientific, order number 72400120) to which 11 mM D-glucose, 200 µM hypoxanthine, 24 mg/L gentamicin and 0.6% (w/v) Albumax II had been added. Parasites were kept synchronous by regular sorbitol synchronization of predominantly ring-stage cultures.^34–35^

*P. knowlesi* parasites (A1.H1 clone) were maintained under the same conditions as *P. falciparum*, except that the culture medium was additionally supplemented with 10% (v/v) pooled human serum and the cultures were maintained at a lower hematocrit (2%), as described elsewhere.^28^

HFF cells were maintained *in vitro* in DMEM supplemented with 10% (v/v) bovine calf serum, 50 units/mL penicillin, 50 µg/mL streptomycin, 10 µg/mL gentamicin, 0.2 mM L-glutamine, and 0.25 µg/mL amphotericin B, as described elsewhere.^36^

### *In vitro* antiplasmodial activity assays

*In vitro* antiplasmodial activity against *P. falciparum* and *P. knowlesi* was assessed over 96 and 72 h, respectively, using a modification^5^of the malaria SYBR Green I-based fluorescence assay reported elsewhere.^37^ *P. falciparum* assays were performed starting with ring-stage *P. falciparum*-infected erythrocytes at a parasitemia and hematocrit of 0.5 and 1%, respectively, as previously described.^5^ *P. knowlesi* assays were initiated with asynchronous *P. knowlesi*-infected erythrocytes, also at a parasitemia and hematocrit of 0.5 and 1%, respectively. Assays were performed using the same medium used to culture each of the parasites. Where specified, 100 µM sodium pantothenate (Yick-Vic Chemicals and Pharmaceuticals, Hong Kong) was added to the medium. Infected erythrocytes incubated in the presence of chloroquine (0.25-25 µM for *P. falciparum* assays and 2.5 µM for *P. knowlesi* assays) served as zero proliferation controls. At the end of the 72 or 96 h incubation, assay plates were frozen and stored at -20°C. Plates were subsequently thawed, the lysed cells in each well mixed with SYBR Safe DNA gel stain in 20 mM Tris, pH 7.5, containing 5 mM ethylenediaminetetraacetic acid (EDTA), 0.008% (w/v) saponin, 0.08% (v/v) Triton X-100. The fluorescence in each well was measured as previously described.^5^

### *In vitro* HFF cell proliferation assays

Activity against HFF cells was assessed over a period of 3-4 days using a SYBR safe-based assay, as previously reported.^9, 17^ To facilitate the comparison of antiplasmodial activity and antiproliferative effects on HFF cells, HFF cell assays were performed in RPMI 1640 (rather than DMEM) supplemented with 10% (v/v) bovine calf serum, 50 units/mL penicillin, 50 µg/mL streptomycin, 10 µg/mL gentamicin, 0.2 mM L-glutamine, and 0.25 µg/mL amphotericin B.

### Caco-2 permeability

Caco-2 permeability was assessed as described previously.^9, 38^ The concentration of compounds **1** and **3j** in donor solutions were confirmed to be within 5% of that intended (20 µM). At the end of the permeability assay the mass balance was calculated according to the following equation: Mass balance (%) = (Mass_final-donor_ + Mass_final-acceptor_)/Mass_initial_ × 100. Mass balances of 104 ± 3% (mean ± standard deviation) and 105 ± 1%, were determined for compound **1** and **3j**, respectively.

### *In vitro* microsomal stability

Stability to degradation by mouse and human liver microsomes was assessed *in vitro* in the presence and absence of an NADPH-regenerating system as described elsewhere.^9, 39^

### *In vivo* antiplasmodial activity assays

The antiplasmodial activity of compounds **1** and **3j** was assessed in *P. berghei*-infected female BALB/c mice. To initiate experiments, stock suspensions of *P. berghei* ANKA-infected erythrocytes (stored at -80°C) were rapidly warmed to 37°C, before injection (250 µL) into the intraperitoneal cavity of ‘donor’ SJL/J mice. Once the parasitaemia reached between 1-10%, the donor mice were ethically culled and the infected erythrocytes harvested from one mouse via cardiac puncture. The parasitized erythrocytes collected were immediately diluted to 5 × 10^5^/mL in Krebs-Ringer buffered saline solution, containing 0.2% (w/v) glucose, as previously described.^40^ Fifteen female BALB/c mice, between 8 and 12 weeks of age, were then infected intraperitoneally with 1 × 10^5^ *P. berghei*-infected erythrocytes (in 200 µL of Krebs-Ringer buffered saline solution). Tail bleeds were performed daily and the parasitemia of the collected blood analyzed by giemsa-stained thin blood smear or flow cytometry (as described below). When the parasitemia of all 15 mice reached between 0.4-1.5%, five mice were administered (via intraperitoneal injection) 100 mg/kg of compound **1** (dissolved in 150 mM NaCl containing 20.9%, v/v, DMSO), another five mice were administered 100 mg/kg **3j** (dissolved as for compound **1**), and the final five mice were administered a corresponding volume of the vehicle (160 µL per 20 g mouse).

For determination of the whole-blood concentration of the test compounds, 3 h after treatment, mice were tail bled again, this time with 10 µL of blood collected using a Mitra^®^ microsampling device (Neoteryx). The microsampling devices were allowed to dry overnight and processed as described below. Approximately 24 h after the first dose, blood was collected for determination of parasitemia and whole-blood inhibitor concentration (as already described) and immediately thereafter, all mice were administered a second dose. Blood collection was repeated again 48 h after the first dose and the mice were ethically culled at 72 h post-administration of the first dose.

Flow cytometry was used to assess mouse parasitemia according to a modification of a previously reported method.^41^ Immediately after collection, the 5 µL samples of tail blood were mixed with 20 µL of ice-cold staining solution (MT-Ringer Complete; 154 mM NaCl, 5.6 mM KCl, 1 mM MgCl_2_, 2.2 mM CaCl_2_, 20 mM HEPES, 10 mM glucose, 0.5%, w/v, bovine serum albumin, 4 mM EDTA, pH 7.4, 0.22 μm filter sterilized) containing 12 μM JC-1 (Life Technologies, Carlsbad, CA), 5 μM Hoechst 33342 (Sigma-Aldrich, St Louis, MO), 1 μg/mL anti-CD45 APC eFluor 780 (clone 30-F11) (eBioscience, San Diego, CA), 1 μg/mL anti-CD71 APC (clone R17217) (eBioscience, San Diego, CA) and 2 μg/mL anti-TER119 PE-Cyanine7 (BioLegend, San Diego, CA). The cells were stained on ice for 30 min, before being washed (twice) with 500 µL of MT-Ringer Complete (800 × *g*, 5 min) and finally resuspended in 1 mL MT-Ringer Complete. All samples were analyzed using a BD LSRII Fortessa Flow Cytometer operated using the BD FACS Diva software (BD Bioscience, Franklin Lakes, NJ), with 1,000,000 total events collected per sample. The final gating strategy was based on FSC/SSC properties; single cell gating was applied using FSC(area):FSC(height), before the mature erythrocyte population (CD45^-^/CD71^low^/TER119^+^) was selected and gated for the presence of Hoechst and JC-1 (indicative of nuclear and mitochondrial staining, respectively), with CD45^-^/CD71^low^/TER119^+^/Hoechst^+^/JC1^+^ cells defined as mature, infected erythrocytes. The CD45^-^/CD71^high^/TER119^+^ immature erythrocytic population was also examined for parasitaemia via Hoechst^+^/JC1^+^ staining. Final population analyses were performed using FlowJo software (BD Bioscience, Franklin Lakes, NJ).

Analytes were extracted from the mouse blood samples (now dried) collected with microsampling devices, using 80% (v/v) acetonitrile in water. The blood-filled absorbent tip from each device was transferred to a microcentrifuge tube to which 80% (v/v) acetonitrile (100 µL) and the internal standard diazepam (10 µL of 5 µg/mL in 50% (v/v) acetonitrile in water) was added. Each tube was then vortexed, sonicated for 5 min, and mixed on an orbital platform (operating at 500 rpm) for 1 h, prior to centrifugation (10,000 × g, 3 min). An aliquot of the supernatant (3 µL) was subsequently analyzed by LC-MS performed on a Waters Xevo TQS micro coupled to a Waters Acquity UPLC operating in positive electrospray ionization multiple-reaction monitoring mode. Analytes were separated on a Supelco Ascentis Express RP Amide column (50 × 2.1 mm, 2.7 µm) using a 4 min gradient of acetonitrile:water with 0.05% (v/v) formic acid, at a flow rate of 0.4 mL/min. To enable quantitation of the amounts of compound **1** and **3h** detected in the blood of mice administered with the compounds, a standard curve was generated by spiking 20 µL samples of mouse blood (in quadruplicate) with each compound at ten different concentrations, ranging from 5 ng/mL – 20 µg/mL. A 10 µL aliquot of each sample was then collected using a Mitra® microsampling device. Devices were dried overnight and the absorbed analytes extracted and analyzed as for the experimental samples.

### Onset of action experiments

The length of exposure to compounds **1** and **3j** required for irreversible inhibition of *P. falciparum* proliferation was investigated using a modification of a previously reported method.^42^ Test compound (at one of ten concentrations tested) or 0.5 µM chloroquine was added to erythrocytes infected with ring-stage parasites suspended in parasite culture medium (at 0.5% parasitemia and 1% hematocrit) in T25 culture flasks. A T25 flask containing a suspension of infected erythrocytes at the same parasitemia and hematocrit was prepared in the absence of test compound or chloroquine to serve as a control for uninhibited parasite proliferation. Aliquots (200 µL) of each flask were immediately transferred in triplicate to the wells of a 96-well microtitre plate to serve as the 96 h incubation time point. All T25 flasks and plates were incubated stationarily at 37°C under an atmosphere of 96% N_2_, 3% CO_2_ and 1% O_2_. After 5 h, the cells from each flask were resuspended and a 1 mL aliquot transferred to a microcentrifuge tube and centrifuged (10,000 × *g*, 30 s) before the supernatant was removed. The infected erythrocytes were thereafter washed three times in 1 mL of inhibitor-free culture medium before being resuspended in 1 mL of inhibitor-free culture medium or, in the case of the chloroquine-treated control, medium containing 0.5 µM chloroquine. Aliquots (200 µL) of the washed and resuspended cells were then transferred in triplicate to the wells of a 96-well microtitre plate and the plate incubated as before. This procedure was then repeated 10, 24 and 48 h after the assay was initiated and all infected erythrocyte suspensions were incubated for a total of 96 h. The contents of the plates were then frozen at - 20°C prior to thawing and processing as described for the SYBR Safe-based *in vitro* antiplasmodial activity assays.

Onset of action experiments were performed with parasites synchronized by the triple synchronization method described previously.^43^ Briefly, 48 h prior to experiments, *P. falciparum* cultures were sorbitol synchronized as described above. Sorbitol synchronization was then repeated 12 h later, and immediately prior to the assay.

### Erythrocyte pre-loading experiments

The effect of preincubating uninfected erythrocytes with compounds **1** and **3j** on proliferation of *P. falciparum* was investigated using a modification of a method previously described.^8^ Uninfected erythrocytes were incubated (at a hematocrit of 5%) in *P. falciparum* culture medium to which a test compound (at 100 µM) or reference compound (chloroquine at 100 µM or dihydroartemisinin at 25 µM) had been added. The incubation was performed in T25 culture flasks, at 37°C (with shaking) under an atmosphere of 96% N_2_, 3% CO_2_, 1% O_2_. A flask of erythrocytes to which no test/reference compound was added, but that was otherwise treated under identical conditions, served as a “no inhibitor” control. The concentration of DMSO in all flasks was 0.1% (v/v). After 3 h, the cells were transferred to centrifuge tubes and collected by centrifugation (500 × *g*, 5 min). The cell pellets were subsequently washed with 10 mL culture medium (500 × g, 5 min) and then with 1 mL culture medium three times (each time centrifuging at 10 000 × *g* for 2 min) to remove unaccumulated test/reference compound. Finally, the pelleted erythrocytes were suspended in culture medium (at a hematocrit of 5%) without test/reference compound and incubated in T25 culture flasks, as before, for 24 h.

After 24 h, the erythrocytes were again transferred to centrifuge tubes, collected by centrifugation (500 × *g*, 5 min), and then washed with 1 mL culture medium (10 000 × *g*, 2 min). During the incubation of uninfected erythrocytes, *P. falciparum*-infected erythrocytes were enriched using a Miltenyi Biotec VarioMACS magnet, using a modification of a previously described method.^44^ Briefly, ∼30 mL of a predominantly trophozoite/schizont-stage *P. falciparum* culture (4% hematocrit, 5-10% parasitemia) was passed through an LS column (Miltenyi Biotec; previously rinsed with 2 mL culture medium) mounted in the magnet. The *P. falciparum*-infected erythrocytes retained on the column were subsequently washed with culture medium (∼30 mL), before the column was removed from the magnet and the infected erythrocytes eluted with 5 mL medium. The process was repeated (typically 3 times) until a sufficient quantity of *P. falciparum*-infected erythrocytes was obtained. The *P. falciparum*-infected erythrocytes, which were confirmed by Giemsa-stained smear to be at a parasitemia > 95%, were thereafter harvested by centrifugation at 500 × *g* (5 min), washed with 1 mL culture medium (1000 × *g*, 3 min) and the supernatant removed. To generate suspensions of *P. falciparum*-infected erythrocytes at a parasitemia and hematocrit of 1% and 2%, respectively, the washed and pelleted uninfected erythrocytes were diluted in culture medium and mixed with the pelleted > 95% parasitemia trophozoite/schizont stage *P. falciparum*-infected parasites.

Aliquots (100 µL) of each of the suspensions were mixed in triplicate with 100 µL culture medium or 100 µL culture medium containing 0.5 µM chloroquine, in the wells of a 96-well microtitre plate. This resulted in *P. falciparum*-infected erythrocytes at a final parasitemia and hematocrit of 1%, in the presence or absence of 0.25 µM chloroquine, in a final volume of 200 µL. Subsequently, plates were incubated for 72 h at 37°C in an atmosphere of 96% N_2_, 3% CO_2_ and 1% O_2_, before being processed (without freezing) as described for the SYBR Safe-based *in vitro* antiplasmodial activity assays. Initially, the average fluorescence measured across the triplicate wells for each cell suspension in the presence of chloroquine was subtracted from the fluorescence measurements obtained for each cell suspension in the absence of chloroquine. These “background-adjusted” measurements were subsequently expressed as a percentage of the background-adjusted measurements obtained for the wells containing parasites grown in the “no inhibitor control” erythrocytes. The confidence intervals associated with the mean % proliferation determined for *P. falciparum* parasites in each pre-treated erythrocyte suspension were calculated using GraphPad.

## Chemistry

### Materials

All reagents and solvents were purchased either from Chem Impex International Inc. or Sigma-Aldrich Canada, and used without further purification. Dry solvents were obtained from an Innovative Tech Pure Solve MD-7 Solvent purification system. MilliQ-quality water was used whenever water is mentioned. Unless noted otherwise, all reactions were performed under a nitrogen atmosphere. Flash chromatography was performed on RediSep Rf Gold Silica Flash Chromatography Columns from Teledyne ISCO. Analysis by thin layer chromatography (TLC) was performed with 60 Å silica gel F-254 TLC plates from Silicycle. High resolution mass spectrometry (HRMS) spectra were acquired at the McGill University Mass Spectral Facility on an EXACTIVE TM Plus Orbitrap Mass Spectrometer from Thermo Fisher Scientific or a MaXis Impact HD Mass Spectrometer from Bruker. The NMR chemical shifts (δ) are reported in parts per million (ppm) and are referenced to the internal standard tetramethylsilane. Compound purity was determined by reversed-phase HPLC analyses using an Agilent 6120 Quadrupole LCMS system.

### General protocol 1 for synthesis of compounds 3a and 3e-u

The synthesis of compounds **3a** and **3e-u** followed the previously reported procedure with modifications.^12^ The Boc-protected heteroaromatic compound (0.19-0.82 mmol, 1.0 eq) was dissolved in a mixture of DCM and trifluoroacetic acid (1:1, v/v, 2-4 mL). The reaction was stirred at room temperature for 30 min-2 h, or until completion of the deprotection as judged by TLC. An aqueous NaOH solution (1 M, 10-20 mL) was used to quench the reaction, making sure that the final pH was higher than 9. The product was extracted in EtOAc (4 × 10 mL). The combined organic layers were dried over anhydrous sodium sulfate. The solid was removed by filtration and the solvent was evaporated *in vacuo* to afford the crude free amine, which was used directly in the next step.

In a pressure vessel (5 mL), the free amine dissolved in anhydrous methanol or ethanol (2 mL) was mixed with D-pantolactone (0.75-3.27 mmol, 4.0 eq) and triethylamine (0.75-3.27 mmol, 4.0 eq). The reaction was heated to 115°C for 12-16 h in an oil bath. The reaction mixture was directly loaded on a silica gel column and purification was achieved with a gradient of 0-100% EtOAc in hexanes, followed by 0-50% methanol in EtOAc.

### General protocol 2 for synthesis of compounds 3b-d

The synthesis of compounds **3b-d** followed the previously reported procedure with modifications.^12, 45–46^ Compound **6b**, **6c** or **6d** (0.95 mmol, 1.0 eq) was added to an aqueous solution (14 mL) of ammonium hydroxide (100.66 mmol, 105.8 eq), before addition of ammonium chloride (0.26 mmol, 0.3 eq). The mixture was heated to 105°C for 16 h. The reaction mixture was cooled to room temperature and extracted with EtOAc (4 × 10 mL). The organic layers were dried over anhydrous sodium sulfate and concentrated *in vacuo* to get the crude amide.

The crude amide was dissolved in THF (10 mL) and lithium aluminum hydride (4.75 mmol, 5.0 eq) in THF (2 mL) was slowly added. The mixture was stirred at room temperature for 16 h, before quenching with water (0.18 mL), a 15% aqueous NaOH solution (0.18 mL) and water (10 mL) sequentially. The product was extracted with EtOAc (4 × 10 mL). The combined organic layers were dried over anhydrous sodium sulfate and concentrated *in vacuo* to get the crude amine.

To the crude amine were added D-pantolactone (3.80 mmol, 4.0 eq), triethylamine (3.80 mmol, 4.0 eq) and ethanol (2 mL). The reaction was stirred at 115°C for 12 h. The reaction mixture was purified by flash chromatography on silica gel using a gradient of 0-100% EtOAc in hexanes, followed by 0-50% methanol in EtOAc.

A solution of compound **27** (0.78 mmol, 1.0 eq) in dried THF was added dropwise to a suspension of lithium aluminum hydride (2.30 mmol, 3.0 eq) in dried THF over 3 min at 0°C. The mixture was then refluxed for 4 h. Water (3 mL) was added and the mixture was stirred for 3 min at room temperature. The product was extracted with EtOAc (4 × 10 mL), dried with sodium sulfate and evaporated to afford amine **28**. To the amine in EtOH (2 mL) was added D-pantolactone (1.17 mmol, 1.5 eq) followed by triethylamine (1.19 mmol, 1.5 eq). The mixture was then heated in a microwave reactor at 100°C for 2.5 h. Subsequently, the reaction mixture was concentrated *in vacuo* and the residue was purified by flash chromatography on silica gel using a gradient of 0-100% EtOAc in hexanes, followed by 0-50% methanol in EtOAc.

### Synthesis of compound 4

The synthesis of compound **4** followed the previously reported procedure with modifications.^47–48^ 1-Heptyne (2.75 mmol, 1.2 eq) was added dropwise to a round-bottomed flask, which contained ethylmagnesium bromide (2.75 mmol, 1.2 eq) in THF (4 mL) at 50°C. The reaction mixture was stirred at 50°C for 30 min, before *N*-(*tert*-Butoxycarbonyl)glycine *N’*-methoxy-*N’*-methylamide (2.29 mmol, 1.0 eq) in THF (4 mL) was added. The resulting mixture was stirred at 50°C for 16 h, before being cooled and poured into water (20 mL). The product was extracted with EtOAc (4 × 20 mL). The combined organic layers were dried over anhydrous sodium sulfate and concentrated *in vacuo*. Purification was achieved on silica gel using a gradient of 0-100% EtOAc in hexanes.

### Synthesis of compound 5

The synthesis of compound **5** followed the previously reported procedure with modifications.^49^ To compound **4** (0.87 mmol, 1.0 eq) dissolved in methanol (3 mL) was added hydrazine monohydrate (2.95 mmol, 3.4 eq). The reaction was stirred at room temperature for 1 h. The product was purified by flash chromatography on silica gel using a gradient of 0-100% EtOAc in hexanes.

### General protocol 3 for synthesis of compounds 6b-d

The synthesis of compounds **6b-d** followed the previously reported procedure with modifications.^50^ To the desired commercial heteroaromatic compound (2.10 mmol, 1.0 eq) in dry DMF (4.5 mL) were added 1-bromopentane (2.31 mmol, 1.1 eq) and sodium carbonate (4.20 mmol, 2.0 eq) sequentially. The mixture was stirred at 80°C for 16 h, before being diluted in EtOAc (40 mL) and washed with water (15 mL). The organic layer was dried over anhydrous sodium sulfate and concentrated *in vacuo*. Purification was achieved on silica gel using a gradient of 0-100% EtOAc in hexanes.

### General protocol 4 for synthesis of compounds 12e-i and 15j-m

The synthesis of compounds **12e-i** and **15j-m** followed the previously reported procedure with modifications.^51^ The desired commercial aldehyde (1.20-2.00 mmol, 1.0 eq) was added to hydroxylamine (1.25-2.10 mmol, 1.1 eq) in *t*-BuOH/H2O (1:1, v/v, 4 mL). After stirring for 30 min to 12 h at room temperature or 80°C, TLC analysis indicated that oxime formation was complete. The product was extracted with EtOAc (4 × 10 mL). The combined organic layers were dried over anhydrous sodium sulfate and concentrated *in vacuo* to get the crude oxime.

To a stirred solution of oxime in DMF (2 mL), was added *N*-chlorosuccinimide (1.55-2.60 mmol, 1.3 eq). The reaction mixture was stirred at room temperature for 3 h. The pale green solution was then diluted in brine (10 mL), before extraction of the product was in diethyl ether (3 × 10 mL). The combined organic layers were dried with anhydrous sodium sulfate and concentrated *in vacuo* to afford the crude **11e-i** or **14j-m**.

To a stirred solution of the desired alkyne (0.90-1.29 mmol, 0.7 eq) in DCM (2 mL), triethylamine (1.10-1.84 mmol, 0.9 eq) was added. A solution of **11e-i** or **14j-m** in DCM (2 mL) was added dropwise. The reaction was stirred at room temperature for 16 h. The reaction mixture was washed with brine (10 mL), dried over anhydrous sodium sulfate and concentrated *in vacuo*. Purification was achieved on silica gel using a gradient of 0-100% EtOAc in hexanes.

### Synthesis of compound 17

The synthesis of compound **17** followed the previously reported procedure with modifications.^50, 52^ To a solution of Boc-protected propargylamine (1.93 mmol, 1.0 eq) in DMF (2 mL) and methanol (0.45 mL) were added CuI (0.19 mmol, 0.1 eq) and trimethylsilyl azide (2.90 mmol, 1.5 eq). After heating at 100°C for 16 h, the reaction was cooled to room temperature and filtered through Celite. This crude reaction mixture was used directly in the following step.

To the crude residue were added sodium carbonate (3.87 mmol, 2.0 eq) and 1-bromopentane (2.13 mmol, 1.1 eq) sequentially. The reaction was heated to 80°C for 16 h. The reaction mixture was cooled to room temperature and filtered through Celite. The mixture was then loaded on silica gel directly and purified with a gradient of 0-100% EtOAc in hexanes.

### Synthesis of compound 19

The synthesis of compound **19** followed the previously reported procedure with modifications.^53–55^ To *N*-Boc-glycinamide (2.18 mmol, 1.0 eq) in DCM (5 mL) was added triethyloxonium hexafluorophosphate (2.40 mmol. 1.1 eq) in DCM (2 mL). After stirring at room temperature for 24 h, the reaction mixture was poured into an ice-cold aqueous sodium carbonate solution (1 M, 10 mL). The product was extracted in DCM (4 × 10 mL). The combined organic layers were dried over anhydrous sodium sulfate and concentrated *in vacuo*. The crude residue was used directly for the following step.

To hexanoic acid hydrazide (2.18 mmol, 1.0 eq) were added triethylamine (2.18 mmol, 1.0 eq) and an ethanolic solution (2 mL) of the residue from the previous step. The reaction mixture was stirred at 115°C for 16 h. The mixture was loaded on silica gel directly and purified with a gradient of 0-100% EtOAc in hexanes.

### Synthesis of compound 21

The synthesis of compound **21** followed the previously reported procedure with modifications.^23^ To *N*-Boc glycine (1.14 mmol, 1.0 eq) dissolved in DCM (3 mL), was added *N*-ethoxycarbonyl-2-ethoxy-1,2-dihydroquinoline (1.14 mmol, 1.0 eq), followed by hexanoic acid hydrazide (1.37 mmol, 1.2 eq). The mixture was stirred at room temperature for 16 h. The mixture was filtered and concentrated *in vacuo* to provide the desired coupling product **20**, which was used directly in the next step.

To the crude residue of **20** in dry THF (5 mL) was added Lawesson’s reagent (1.26 mmol, 1.1 eq). The reaction mixture was stirred at 70°C for 16 h, before concentration *in vacuo*. The concentrated crude product was loaded on silica gel directly. Purification was achieved using a gradient of 0-100% EtOAc in hexanes.

### Synthesis of compound 22

The synthesis of compound **22** followed the previously reported procedure with modifications.^56^ At 0°C, carbonyldiimidazole (0.92 mmol, 1.1 eq) in DCM (1 mL) was mixed with *N*-Boc glycine (0.88 mmol, 1.0 eq) in DCM (4 mL). The mixture was stirred at 0°C for 30 minutes. Hexanoic acid hydrazide (0.88 mmol, 1.0 eq) in DCM (1 mL) was next added. After stirring at 0°C for 45 minutes, CBr_4_ (1.76 mmol, 2.0 eq) and PPh_3_ (1.76 mmol, 2.0 eq) in DCM (2 mL) were added at 0°C sequentially. The reaction mixture was stirred for 16 h while allowing the mixture to warm up to the room temperature. The reaction mixture was loaded on silica gel directly and purified with a gradient of 0-100% EtOAc in hexanes.

### General protocol 5 for synthesis of compounds 24r-s

The synthesis of compounds **24r-s** followed the previously reported procedure with modifications.^57–58^ To a stirring solution of the desired commercial nitrile (1.92-2.06 mmol, 1.0 eq) in EtOH (2 mL) were added hydroxylamine (9.60-0.29 mmol, 5.0 eq) and triethylamine (2.11-2.26 mmol, 1.1 eq). The reaction was heated to 80°C for 16 h. The reaction mixture was diluted in EtOAc (20 mL) and washed with water (10 mL). The organic layer was dried over anhydrous sodium sulfate and concentrated *in vacuo*, and used directly for the next step.

To a stirring solution of 1-ethyl-3-(3-dimethylaminopropyl)carbodiimide (1.92-2.06 mmol, 1.0 eq) in dry acetonitrile (2-5 mL) was added the desired carboxylic acid (1.92-2.06 mmol, 1.0 eq). The mixture was stirred at room temperature for 20 minutes, before adding the crude product from the first step in DMF (2 mL). The reaction mixture was stirred at room temperature for 4 h, and then heated at 110°C for 1.5-12 h. The reaction residue was loaded on silica gel directly and purified with a gradient of 0-100% EtOAc in hexanes.

### General protocol 6 for synthesis of compounds 25t-u

The synthesis of compounds **25t-u** followed the previously reported procedure with modifications.^24^ The desired commercial alkyne (0.38-0.73 mmol, 1.0 eq), nitrile (0.73-2.28 mmol, 1.0-6.0 eq), 8-methylquinoline *N*-oxide (0.49-0.94 mmol, 1.3 eq) and the catalyst BrettPhosAuNTf_2_ (0.02-0.04 mmol, 0.05 eq) were added to a pressure vessel (5 mL). The reaction was heated at 60°C for 1-3 h in a microwave reactor. The cloudy mixture was loaded on silica gel directly and purified with a gradient of 0-100% EtOAc in hexanes.

### Synthesis of compound 26

DMF (3.30 mmol, 1.3 eq) in DCM (3 mL) was cooled to 0°C, and POCl_3_ (3.20 mmol, 1.3 eq) was added dropwise. The mixture was stirred for 30 min at 0°C. Next, 2-pentyl thiophene (2.50 mmol, 1.0 eq) dissolved in DCM (5 mL) was added to the mixture dropwise over 45 min at 0°C. After warming to room temperature, the mixture was heated to 70°C and stirred for 1.5 h. Upon cooling to room temperature, the mixture was poured into iced water and neutralized with sodium bicarbonate. The mixture was extracted with diethyl ether (4 × 10 mL) and dried with sodium sulfate. After evaporation, the reaction residue was loaded on silica gel directly and purified with a gradient of 0-100% EtOAc in hexanes, followed by 0-50% methanol in EtOAc.

### Synthesis of compound 27

To a mixture of compound **26** (1.50 mmol, 1.0 eq), NH_2_OH·HCl (1.50 mmol, 1.0 eq) and NaOAc (1.65 mmol, 1.10 eq) in a dried round-bottomed flask was added CuSO_4_.5H_2_O (0.08 mmol, 0.05 eq). The mixture was heated at 110°C on an oil bath with thorough stirring for 5 h (TLC monitoring revealed that the reaction was complete). After cooling the reaction mixture to room temperature, water (5 mL) was added to quench the reaction followed by extraction with EtOAc (3 × 10 mL). The combined extracts were washed with brine (2 × 5 mL), dried (anhydrous sodium sulfate) and concentrated under reduced pressure. The crude product was loaded on silica gel directly and purified with a gradient of 0-100% EtOAc in hexanes, followed by 0-50% methanol in EtOAc.

### Compound characterization

#### (*R*)-2,4-Dihydroxy-3,3-dimethyl-*N*-((5-pentyl-1H-pyrazol-3-yl)methyl)butanamide (3a)

Compound **3a** was prepared from **5** (0.48 mmol) using general protocol 1. Yield: 31%, *R*_f_ = 0.50 (10% MeOH in EtOAc). Purity is 99%, R_t_ = 4.87 min with method A (see supporting information); R_t_ = 5.27 min with method B. ^1^H NMR (500 MHz, CDCl_3_) δ 7.55 (m, 1H), 5.93 (s, 1H), 4.52 (dd, *J* = 15.4, 6.2 Hz, 1H), 4.32 (dd, *J* = 15.5, 5.0 Hz, 1H), 4.03 (s, 1H), 3.47 (m, 2H), 2.55 (t, *J* = 7.7 Hz, 2H), 1.58 (m, 2H), 1.37–1.26 (m, 4H), 0.97 (s, 3H), 0.91 (s, 3H), 0.87 (t, *J* = 6.7 Hz, 3H). ^13^C NMR (126 MHz, CDCl_3_) δ 174.18, 147.71, 147.20, 102.27, 77.28 (based on HSQC and HMBC), 70.51, 39.38, 36.06, 31.40, 28.82, 26.23, 22.36, 21.49, 20.67, 13.96. HRMS for C_15_H_27_N_3_O_3_Na [M+Na]^+^ calcd. 320.1945, found 320.1937.

#### (*R*)-2,4-Dihydroxy-3,3-dimethyl-*N*-((1-pentyl-1H-imidazol-4-yl)methyl)butanamide (3b)

Compound **3b** was prepared from **6b** (0.95 mmol) using general protocol 2. Yield: 19%, *R*_f_ = 0.24 (10% MeOH in EtOAc). Purity is 99%, R_t_ = 3.27 min with method A; R_t_ = 6.35 min with method B. ^1^H NMR (500 MHz, CDCl_3_) δ 7.62 (m, 1H), 7.36 (d, *J* = 1.1 Hz, 1H), 6.83 (s, 1H), 4.40 (dd, *J* = 15.0, 6.1 Hz, 1H), 4.32 (dd, *J* = 14.9, 5.5 Hz, 1H), 4.05 (s, 1H), 3.86 (t, *J* = 7.2 Hz, 2H), 3.49 (d, *J* = 11.4 Hz, 1H), 3.46 (d, *J* = 11.4 Hz, 1H), 1.75 (m, 2H), 1.33 (m, 2H), 1.25 (m, 2H), 1.05 (s, 3H), 0.93 (s, 3H), 0.89 (t, *J* = 7.2 Hz, 3H). ^13^C NMR (126 MHz, CDCl_3_) δ 173.55, 138.48, 136.54, 116.36, 77.50, 70.80, 47.31, 39.70, 36.18, 30.57, 28.63, 22.46, 22.12, 20.28, 13.86. HRMS for C_15_H_28_N_3_O_3_ [M+H]^+^ calcd. 298.21252, found 298.21185.

#### (*R*)-2,4-Dihydroxy-3,3-dimethyl-*N*-((1-pentyl-1H-pyrazol-4-yl)methyl)butanamide (3c)

Compound **3c** was prepared from **6c** (0.95 mmol) using general protocol 2. Yield: 23%, *R*_f_ = 0.59 (10% MeOH in EtOAc). Purity is 99%, R_t_ = 3.30 min with method A; R_t_ = 5.96 min with method B. ^1^H NMR (400 MHz, CDCl_3_) δ 7.40 (s, 1H), 7.36 (s, 1H), 6.95 (m, 1H), 4.33 (d, *J* = 5.7 Hz, 2H), 4.07–4.04 (m, 3H), 3.52 (d, *J* = 11.2 Hz, 1H), 3.49 (d, *J* = 11.2 Hz, 1H), 1.83 (m, 2H), 1.39–1.20 (m, 4H), 1.02 (s, 3H), 0.93–0.86 (m, 6H). ^13^C NMR (126 MHz, CDCl_3_) δ 172.56, 138.37, 128.30, 117.85, 77.72, 71.42, 52.31, 39.47, 33.55, 30.08, 28.74, 22.21, 21.51, 20.10, 13.92. HRMS for C_15_H_27_N_3_O_3_Na [M+Na]^+^ calcd. 320.19446, found 320.19396.

#### (*R*)-2,4-Dihydroxy-3,3-dimethyl-*N*-((1-pentyl-1H-1,2,4-triazol-3-yl)methyl)butanamide (3d)

Compound **3d** was prepared from **6d** (1.14 mmol) using general protocol 2. Yield: 4%, *R*_f_ = 0.62 (20% MeOH in EtOAc). Purity is 94%, R_t_ = 3.66 min with method A; R_t_ = 2.14 min with method B. ^1^H NMR (500 MHz, CDCl_3_) δ 7.99 (s, 1H), 7.54 (bs, 1H), 4.56 (m, 2H), 4.09–4.05 (m, 3H), 3.9 (d, *J* = 11.3 Hz, 1H), 3.47 (d, *J* = 11.3 Hz, 1H), 1.84 (m, 2H), 1.37–1.21 (m, 4H), 1.03 (s, 3H), 0.95 (s, 3H), 0.88 (t, *J* = 7.2 Hz, 3H). ^13^C NMR (126 MHz, CDCl_3_) δ 173.72, 160.98, 143.42, 77.64, 70.54, 49.92, 39.51, 36.62, 29.31, 28.55, 22.07, 21.90, 20.78, 13.83. HRMS for C_14_H_26_N_4_O_3_Na [M+Na]^+^ calcd. 321.1897, found 321.1895.

#### (*R*)-2,4-Dihydroxy-3,3-dimethyl-*N*-((3-pentylisoxazol-5-yl)methyl)butanamide (3e)

Compound **3e** was prepared from **12e** (0.35 mmol) using general protocol 1. Yield: 44%, *R*_f_ = 0.39 (100% EtOAc). Purity is 97%, R_t_ = 4.56 min with method A; R_t_ = 6.66 min with method B. ^1^H NMR (500 MHz, CDCl_3_) δ 7.36 (m, 1H), 6.04 (s, 1H), 4.55 (d, *J* = 6.1 Hz, 2H), 4.09 (s, 1H), 3.53 (d, *J* = 11.1 Hz, 1H), 3.50 (d, *J* = 11.1 Hz, 1H), 2.62 (m, 2H), 1.62 (m, 2H), 1.38–1.28 (m, 4H), 1.02 (s, 3H), 0.92 (s, 3H), 0.89 (t, *J* = 7.0 Hz, 3H). ^13^C NMR (126 MHz, CDCl_3_) δ 173.19, 168.10, 164.44, 101.92, 77.82, 71.47, 39.32, 34.77, 31.30, 27.86, 25.95, 22.30, 21.13, 20.35, 13.92. HRMS for C_15_H_25_N_2_O_4_ [M+H]^+^ calcd. 297.18198, found 297.18244.

#### (*R*)-*N*-((3-Butylisoxazol-5-yl)methyl)-2,4-dihydroxy-3,3-dimethylbutanamide (3f)

Compound **3f** was prepared from **12f** (0.32 mmol) using general protocol 1. Yield: 38%, *R*_f_ = 0.35 (100% EtOAc). ^1^H NMR (400 MHz, CDCl_3_) δ 7.44(bs, 1H), 6.04 (s, 1H), 4.54 (d, *J* = 6.0 Hz, 2H), 4.08 (s, 1H), 3.52 (d, *J* = 11.5, 1H), 3.48 (d, *J* = 11.5, 1H), 3.22 (bs, 2H), 2.61 (t, *J* = 7.7Hz, 2H), 1.56-1.61 (m, 2H), 1.30-1.40 (m, 2H), 0.99 (s, 3H), 0.89-0.93 (m, 6H). ^13^C NMR (126 MHz, CDCl_3_) δ 173.3, 168.1, 164.4, 101.9, 77.7, 71.4, 39.3, 34.8, 30.2, 25.6, 22.2, 21.1, 20.4, 13.7. HRMS for C_14_H_23_N_2_O_4_ [M-H]^-^ calcd. 283.1663, found 283.1662.

#### (*R*)-*N*-((3-Hexylisoxazol-5-yl)methyl)-2,4-dihydroxy-3,3-dimethylbutanamide (3g)

Compound **3g** was prepared from **12g** (1.09 mmol) using general protocol 1. Yield: 63%, *R*_f_ = 0.39 (100% EtOAc). ^1^H NMR (500 MHz, CDCl_3_) δ 7.45 (t, *J* = 5.9Hz, 1H), 6.06 (s, 1H), 4.56 (d, *J* = 6.0 Hz, 2H), 4.10 (s, 1H), 3.54 (d, *J* = 11.2, 1H), 3.50 (d, *J* = 11.2, 1H), 2.63 (t, *J* = 7.7Hz, 2H), 1.60-1.66 (m, 2H), 1.29-1.39 (m, 6H), 1.02 (s, 3H), 0.94 (s, 3H), 0.89 (t, *J* = 6.9Hz, 3H). ^13^C NMR (126 MHz, CDCl_3_) δ 173.4, 168.1, 164.4, 101.9, 77.7, 71.4, 39.3, 34.8, 31.4, 28.8, 28.1, 26.0, 22.5, 21.1, 20.4, 14.0. HRMS for C_16_H_28_N_2_O_4_Na [M+Na]^+^ calcd. 335.1941, found 335.1943.

#### (*R*)-2,4-Dihydroxy-3,3-dimethyl-*N*-((3-phenethylisoxazol-5-yl)methyl)butanamide (3h)

Compound **3h** was prepared from **12h** (0.40 mmol) using general protocol 1. Yield: 54%, *R*_f_ = 0.43 (100% EtOAc). ^1^H NMR (500 MHz, CDCl_3_) δ 7.21-7.33 (m, 6H), 6.00 (s, 1H), 4.57 (d, *J* = 2.2 Hz, 1H), 4.56 (d, *J* = 2.3 Hz, 1H), 4.12 (s, 1H), 3.56 (d, *J* = 11.1 Hz, 1H), 3.52 (d, *J* = 11.1 Hz, 1H), 2.99 (m, 4H), 1.04 (s, 3H), 0.95 (s, 3H). ^13^C NMR (126 MHz, CDCl_3_) δ 173.1, 168.2, 163.5, 140.4, 128.5, 128.3, 126.4, 102.0, 77.8, 71.5, 39.3, 34.8, 34.2, 27.8, 21.1, 20.4. HRMS for C_18_H_24_N_2_O_4_Na [M+Na]^+^ calcd. 355.1628, found 355.1630.

#### (*R*)-2,4-Dihydroxy-3,3-dimethyl-*N*-(2-(3-pentylisoxazol-5-yl)ethyl)butanamide (3i)

Compound **3i** was prepared from **12i** (1.15 mmol) using general protocol 1. Yield: 47%, *R*_f_ = 0.39 (100% EtOAc). ^1^H NMR (500 MHz, CDCl_3_) δ 7.06 (bs, 1H), 5.96 (s, 1H), 4.05 (s, 1H), 3.67-3.72 (m, 1H), 3.60-3.66 (m, 1H), 3.53 (d, *J* = 11.4, 1H), 3.50 (d, *J* = 11.2, 1H), 3.01 (t, *J* = 6.6 Hz, 2H), 2.63 (t, *J* = 7.1 Hz, 2H), 1.62-1.68 (m, 4H), 1.34-1.37 (m, 4H), 1.02 (s, 3H), 0.90-0.93 (m, 6H). ^13^C NMR (126 MHz, CDCl_3_) δ 173.4, 170.0, 164.4, 101.7, 77.6, 71.3, 39.2, 37.0, 31.3, 27.9, 27.1, 26.0, 22.3, 21.3, 20.2, 13.9. HRMS for C_16_H_28_N_2_O_4_Na [M+Na]^+^ calcd. 335.1941, found 335.1943.

#### (*R*)-2,4-Dihydroxy-3,3-dimethyl-*N*-((5-pentylisoxazol-3-yl)methyl)butanamide (3j)

Compound **3j** was prepared from **15j** (0.43 mmol) using general protocol 1. Yield: 74%, *R*_f_ = 0.50 (100% EtOAc). Purity is 99%, R_t_ = 12.48 min with method C (see supporting information); R_t_ = 13.57 min with method D. ^1^H NMR (500 MHz, CDCl_3_) δ 7.41 (m, 1H), 5.95 (s, 1H), 4.53 (dd, *J* = 15.9, 6.1 Hz, 1H), 4.47 (dd, *J* = 15.8, 5.9 Hz, 1H), 4.07 (s, 1H), 3.50 (m, 2H), 2.69 (t, *J* = 7.6 Hz, 2H), 1.66 (m, 2H), 1.37–1.28 (m, 4H), 1.01 (s, 3H), 0.94 (s, 3H), 0.89 (t, *J* = 7.0 Hz, 3H). ^13^C NMR (126 MHz, CDCl_3_) δ 174.67, 173.48, 160.98, 100.01, 77.81, 71.11, 39.38, 35.02, 31.17, 27.10, 26.66, 22.24, 21.39, 20.55, 13.88. HRMS for C_15_H_26_N_2_O_4_Na [M+Na]^+^ calcd. 321.17848, found 321.17918.

#### (*R*)-*N*-((5-Butylisoxazol-3-yl)methyl)-2,4-dihydroxy-3,3-dimethylbutanamide (3k)

Compound **3k** was prepared from **15k** (0.16 mmol) using general protocol 1. Yield: 77%, *R*_f_ = 0.45 (100% EtOAc). ^1^H NMR (500 MHz, CDCl_3_) δ 7.41 (bs, 1H), 5.98 (s, 1H), 4.56 (dd, *J* = 6.1, 15.9 Hz, 1H), 4.50 (dd, *J* = 5.85, 15.9 Hz, 1H), 4.11 (s, 1H), 3.54 (s, 2H), 3.20 (bs, 2H), 2.73 (t, *J* = 7.6 Hz, 2H), 1.64-1.70 (m, 2H), 1.35-1.43 (m, 2H), 1.05 (s, 3H), 0.97 (s, 3H), 0.95 (t, *J* = 7.35 Hz, 3H). ^13^C NMR (126 MHz, CDCl_3_) δ 174.7, 173.4, 161.0, 100.0, 77.9, 71.1, 39.4, 35.0, 29.5, 26.4, 22.1, 21.4, 20.6, 13.6. HRMS for C_14_H_24_N_2_O_4_Na [M+Na]^+^ calcd. 307.1628, found 307.1622.

#### (*R*)-2,4-Dihydroxy-3,3-dimethyl-*N*-((5-pentylisoxazol-3-yl)methyl)butanamide (3l)

Compound **3l** was prepared from **15l** (0.30 mmol) using general protocol 1. Yield: 77%, *R*_f_ = 0.48 (100% EtOAc). ^1^H NMR (500 MHz, CDCl_3_) δ 7.35 (bs, 1H), 5.98 (s, 1H), 4.59 (dd, *J* = 6.1, 15.9 Hz, 1H), 4.51 (dd, *J* = 5.9, 15.9 Hz, 1H), 4.11 (s, 1H), 3.56 (s, 2H), 2.73(t, *J* = 7.6 Hz, 2H), 1.66-1.72 (m, 2H), 1.30-1.40 (m, 6H), 1.07 (s, 3H), 0.99 (s, 3H), 0.91 (t, *J* = 6.9 Hz, 3H). ^13^C NMR (126 MHz, CDCl_3_) δ 174.7, 173.2, 161.0, 100.0, 80.0, 71.2, 39.4, 35.1, 31.4, 28.7, 27.4, 26.7, 22.5, 21.5, 20.6, 14.0. HRMS for C_16_H_28_N_2_O_4_Na [M+Na]^+^ calcd. 335.1941, found 335.1952.

#### (*R*)-2,4-Dihydroxy-3,3-dimethyl-*N*-((5-phenethylisoxazol-3-yl)methyl)butanamide (3m)

Compound **3m** was prepared from **15m** (0.55 mmol) using general protocol 1. Yield: 21%, *R*_f_ = 0.50 (100% EtOAc). ^1^H NMR (500 MHz, CDCl_3_) δ 7.29-7.33 (m, 3H), 7.23-7.26 (m, 1H), 7.18-7.20 (m, 2H), 5.94 (s, 1H), 4.58 (dd, *J* = 6.1, 15.9 Hz, 1H), 4.49 (dd, *J* = 5.9, 15.9 Hz, 1H), 4.11 (d, *J* = 5.2 Hz, 1H), 3.72 (d, *J* = 5.2 Hz, 1H), 3.55 (d, *J* = 4.9 Hz, 1H), 3.00-3.09 (m, 5H), 1.07 (s, 3H), 0.98 (s, 3H). ^13^C NMR (126 MHz, CDCl_3_) δ 173.34, 173.09, 161.01, 139.85, 128.60, 128.24, 126.52, 100.47, 78.06, 71.24, 39.43, 35.06, 33.46, 28.46, 21.57, 20.60. HRMS for C_18_H_24_N_2_O_4_Na [M+Na]^+^ calcd. 335.1628, found 335.1627.

#### (*R*)-2,4-Dihydroxy-3,3-dimethyl-*N*-((2-pentyl-2H-1,2,3-triazol-4-yl)methyl)butanamide (3n)

Compound **3n** was prepared from **17** (0.78 mmol) using general protocol 1. Yield: 45%, *R*_f_ = 0.65 (10% MeOH in EtOAc). Purity is 99%, R_t_ = 3.49 min with method A; R_t_ = 2.31 min with method B. ^1^H NMR (500 MHz, CDCl_3_) δ 7.50 (s, 1H), 7.18 (bs, 1H), 4.59 (dd, *J* = 15.4, 6.0 Hz, 1H), 4.51 (dd, *J* = 15.4, 5.7 Hz, 1H), 4.36 (t, *J* = 7.2 Hz, 2H), 4.08 (d, *J* = 4.7 Hz, 1H), 3.53 (m, 2H), 1.93 (m, 2H), 1.39–1.23 (m, 4H), 1.04 (s, 3H), 0.94 (s, 3H), 0.89 (t, *J* = 7.2 Hz, 3H). ^13^C NMR (126 MHz, CDCl_3_) δ 172.75, 144.59, 132.41, 77.91, 71.33, 54.99, 39.47, 34.44, 29.41, 28.62, 22.12, 21.44, 20.38, 13.88. HRMS for C_14_H_26_N_4_O_3_Na [M+Na]^+^ calcd. 321.1897, found 321.1899.

#### (*R*)-2,4-Dihydroxy-3,3-dimethyl-*N*-((5-pentyl-4H-1,2,4-triazol-3-yl)methyl)butanamide (3o)

Compound **3o** was prepared from **19** (0.74 mmol) using general protocol 1. Yield: 24%, *R*_f_ = 0.54 (20% MeOH in EtOAc). Purity is 99%, R_t_ = 3.73 min with method A; R_t_ = 3.892 min with method B. ^1^H NMR (500 MHz, CD_3_OD) δ 4.47 (m, 2H), 3.96 (s, 1H), 3.46 (d, *J* = 11.0 Hz, 1H), 3.43 (d, *J* = 11.0 Hz, 1H), 2.73 (m, 2H), 1.72 (m, 2H), 1.41–1.27 (m, 4H), 0.95 (s, 3H), 0.94 (s, 3H), 0.91 (t, *J* = 7.1 Hz, 3H). ^13^C NMR (126 MHz, CD_3_OD) δ 176.20, 161.52, 159.42, 77.43, 70.07, 40.50, 37.10, 32.32, 28.73, 27.06, 23.31, 21.53, 21.00, 14.25. HRMS for C_14_H_25_N_4_O_3_ [M-H]^-^ calcd. 297.19321, found 297.19359.

#### (*R*)-2,4-Dihydroxy-3,3-dimethyl-*N*-((5-pentyl-1,3,4-thiadiazol-2-yl)methyl)butanamide (3p)

Compound **3p** was prepared from **21** (0.69 mmol) using general protocol 1. Yield: 41%, *R*_f_ = 0.16 (100% EtOAc). Purity is 92%, R_t_ = 12.25 min with method A; R_t_ = 2.14 min with method B. ^1^H NMR (500 MHz, CDCl_3_) δ 7.66 (m, 1H), 4.88 (dd, *J* = 15.9, 6.3 Hz, 1H), 4.81 (dd, *J* = 15.9, 6.1 Hz, 1H), 4.12 (s, 1H), 3.54 (m, 2H), 3.06 (t, *J* = 7.6, 2H), 1.78 (m, 2H), 1.42–1.30 (m, 4H), 1.04 (s, 3H), 0.98 (s, 3H), 0.90 (t, *J* = 7.1 Hz, 3H). ^13^C NMR (126 MHz, CDCl_3_) δ 173.51, 172.51, 166.77, 78.10, 71.45, 39.59, 38.07, 31.23, 30.21, 29.80, 22.38, 21.40, 20.91, 14.02. HRMS for C_14_H_25_N_3_O_3_SNa [M+Na]^+^ calcd. 338.1509, found 338.1509.

#### (*R*)-2,4-Dihydroxy-3,3-dimethyl-*N*-((5-pentyl-1,3,4-oxadiazol-2-yl)methyl)butanamide (3q)

Compound **3q** was prepared from **22** (0.26 mmol) using general protocol 1. Yield: 39%, *R*_f_ = 0.47 (10% MeOH in EtOAc). Purity is 96%, R_t_ = 4.82 min with method A; R_t_ = 9.52 min with method B. ^1^H NMR (500 MHz, CDCl_3_) δ 7.73 (t, *J* = 5.8 Hz, 1H), 4.66 (m, 2H), 4.11 (s, 1H), 3.54 (d, *J* = 11.1 Hz, 1H), 3.46 (d, *J* = 11.1 Hz, 1H), 2.79 (t, *J* = 7.6 Hz, 2H), 1.75 (m, 2H), 1.38–1.29 (m, 4H), 0.99 (s, 3H), 0.97 (s, 3H), 0.89 (t, *J* = 6.9 Hz, 3H). ^13^C NMR (126 MHz, CDCl_3_) δ 174.12, 168.05, 163.98, 77.68, 70.91, 39.47, 34.09, 31.18, 26.11, 25.36, 22.25, 21.21, 21.04, 13.93. HRMS for C_14_H_25_N_3_O_4_Na [M+Na]^+^ calcd. 322.17373, found 322.17248.

#### (*R*)-2,4-Dihydroxy-3,3-dimethyl-*N*-((3-pentyl-1,2,4-oxadiazol-5-yl)methyl)butanamide (3r)

Compound **3r** was prepared from **24r** (0.41 mmol) using general protocol 1. Yield: 50%, *R*_f_ = 0.67 (10% MeOH in EtOAc). Purity is 94%, R_t_ = 3.19 min with method A; R_t_ = 3.92 min with method B. ^1^H NMR (500 MHz, CDCl_3_) δ 7.62 (m, 1H), 4.75 (dd, *J* = 17.3, 6.3 Hz, 1H), 4.63 (dd, *J* = 17.3, 5.8 Hz, 1H), 4.10 (s, 1H), 3.50 (m, 2H), 2.68 (m, 2H), 1.71 (m, 2H), 1.36–1.29 (m, 4H), 1.03 (s, 3H), 0.97 (s, 3H), 0.88 (t, *J* = 6.9 Hz, 3H). ^13^C NMR (126 MHz, CDCl_3_) δ 175.89, 173.90, 170.77, 77.89, 71.07, 39.53, 35.34, 31.30, 26.61, 25.92, 22.34, 21.28, 20.96, 13.99. HRMS for C_14_H_24_N_3_O_4_ [M-H]^-^ calcd. 298.17723, found 298.17721.

#### (*R*)-2,4-Dihydroxy-3,3-dimethyl-*N*-((5-pentyl-1,2,4-oxadiazol-3-yl)methyl)butanamide (3s)

Compound **3s** was prepared from **24s** (0.82 mmol) using general protocol 1. Yield: 75%, *R*_f_ = 0.62 (10% MeOH in EtOAc). Purity is 99%, R_t_ = 3.78 min with method A; R_t_ = 6.45 min with method B. ^1^H NMR (500 MHz, CDCl_3_) δ 7.39 (m, 1H), 4.67 (dd, *J* = 16.7, 6.2 Hz, 1H), 4.56 (dd, *J* = 16.7, 5.7 Hz, 1H), 4.10 (s, 1H), 3.55 (m, 2H), 2.86 (t, *J* = 7.6 Hz, 2H), 1.80 (m, 2H), 1.39–1.31 (m, 4H), 1.07 (s, 3H), 0.98 (s, 3H), 0.90 (t, *J* = 7.1 Hz, 3H). ^13^C NMR (126 MHz, CDCl_3_) δ 180.97, 173.41, 167.73, 78.12, 71.24, 39.63, 35.18, 31.23, 26.64, 26.30, 22.28, 21.71, 20.81, 13.96. HRMS for C_14_H_25_N_3_O_4_Na [M+Na]^+^ calcd. 322.17373, found 322.17430.

#### (*R*)-2,4-Dihydroxy-3,3-dimethyl-*N*-((2-pentyloxazol-5-yl)methyl)butanamide (3t)

Compound **3t** was prepared from **25t** (0.34 mmol) using general protocol 1. Yield: 40%, *R*_f_ = 0.50 (10% MeOH in EtOAc). Purity is 97%, R_t_ = 3.76 min with method A; R_t_ = 4.76 min with method B. ^1^H NMR (500 MHz, CDCl_3_) δ 7.19 (m, 1H), 6.79 (s, 1H), 4.50 (dd, *J* = 15.8, 6.0 Hz, 1H), 4.44 (dd, *J* = 15.8, 5.8 Hz, 1H), 4.07 (s, 1H), 3.52 (d, *J* = 11.1 Hz, 1H), 3.49 (d, *J* = 11.1 Hz, 1H), 2.70 (t, *J* = 7.6 Hz, 2H), 1.72 (m, 2H), 1.35–1.29 (m, 4H), 1.01 (s, 3H), 0.91 (s, 3H), 0.88 (t, *J* = 7.0 Hz, 3H). ^13^C NMR (126 MHz, CDCl_3_) δ 173.14, 165.48, 148.04, 124.00, 77.94, 71.59, 39.42, 33.81, 31.36, 28.23, 26.73, 22.38, 21.26, 20.46, 14.03. HRMS for C_15_H_25_N_2_O_4_ [M-H]^-^ calcd. 297.18198, found 297.18170.

#### (*R*)-2,4-Dihydroxy-3,3-dimethyl-*N*-((5-pentyloxazol-2-yl)methyl)butanamide (3u)

Compound **3u** was prepared from **25u** (0.25 mmol) using general protocol 1. Yield: 25%, *R*_f_ = 0.50 (8% MeOH in EtOAc). Purity is 98%, R_t_ = 3.93 min with method A; R_t_ = 6.75 min with method B. ^1^H NMR (400 MHz, CDCl_3_) δ 7.56 (m, 1H), 6.61 (s, 1H), 4.63 (dd, *J* = 16.8, 6.4 Hz, 1H), 4.48 (dd, *J* = 16.8, 5.5 Hz, 1H), 4.07 (s, 1H), 3.51 (d, *J* = 11.2 Hz, 1H), 3.48 (d, *J* = 11.2 Hz, 1H), 2.59 (t, *J* = 7.6, 2H), 1.60 (m, 2H), 1.36–1.27 (m, 4H), 1.04 (s, 3H), 0.99 (s, 3H), 0.89 (t, *J* = 6.9 Hz, 3H). ^13^C NMR (101 MHz, CDCl_3_) δ 173.87, 159.97, 154.15, 121.75, 78.10, 70.72, 39.63, 36.30, 31.32, 27.24, 25.53, 22.41, 21.99, 21.21, 14.05. HRMS for C_15_H_26_N_2_O_4_Na [M+Na]^+^ calcd. 321.1785, found 321.1783.

#### (*R*)-2,4-Dihydroxy-3,3-dimethyl-*N*-((5-pentylthiophen-2-yl)methyl)butanamide (3v)

Compound **3v** was prepared from **27** (0.78 mmol) as described. Yield: 51%. ^1^H NMR (500 MHz, CDCl_3_) δ 7.08 (bs, 1H, -NH), 6.78 (d, *J* = 3.35 Hz, 1H), 6.61 (d, *J* = 3.35Hz, 1H), 4.53-4.64 (m, 2H), 4.09 (s, 1H), 3.66 (m, 1H), 3.58 (d, *J* = 11.15 Hz, 1H), 3.51 (d, *J* = 11.15 Hz, 1H), 2.77 (t, *J* = 7.70 Hz, 2H), 1.65-1.68 (m, 2H), 1.33-1.36 (m, 4H), 1.06 (s, 3H), 0.95 (s, 3H), 0.91 (t, *J* = 6.95 Hz, 3H). ^13^C NMR (126 MHz, CDCl3) δ 172.47, 146.22, 137.71, 125.79, 123.58, 77.73, 71.45, 39.52, 38.18, 31.33, 31.25, 30.14, 22.40, 21.45, 20.14, 14.00. HRMS for C_16_H_27_NNaO_3_S [M+Na]^+^ calcd. 336.1604, found 336.1611.

#### *tert*-Butyl (2-oxonon-3-yn-1-yl)carbamate (4)

Compound **4** was prepared from *N*-(*tert*-butoxycarbonyl)glycine *N’*-methoxy-*N’*-methylamide (2.29 mmol) as described. Yield: 22%. *R*_f_ = 0.62 (30% EtOAc in hexanes). ^1^H NMR (500 MHz, CDCl_3_) δ 5.15 (bs, 1H), 4.11 (d, *J* = 5.0 Hz, 2H), 2.38 (t, *J* = 7.1 Hz, 2H), 1.59 (m, 2H), 1.45 (s, 9H), 1.42–1.29 (m, 4H), 0.91 (t, *J* = 7.2 Hz, 3H). ^13^C NMR (126 MHz, CDCl_3_) δ 183.10, 155.41, 97.87, 80.01, 78.79, 52.28, 30.96, 28.30, 27.24, 22.07, 19.03, 13.86. HRMS for C_14_H_23_NO_3_Na [M+Na]^+^ calcd. 276.1570, found 276.1573.

#### *tert*-Butyl ((5-pentyl-1*H*-pyrazol-3-yl)methyl)carbamate (5)

Compound **5** was prepared from **4** (0.87 mmol) as described. Yield: 56%. *R*_f_ = 0.23 (50% EtOAc in hexanes). ^1^H NMR (500 MHz, CDCl_3_) δ 5.96 (s, 1H), 5.15 (bs, 1H), 4.28 (d, *J* = 5.7 Hz, 2H), 2.60 (t, *J* = 7.9 Hz, 2H), 1.63 (m, 2H), 1.45 (s, 9H), 1.35–1.29 (m, 4H), 0.89 (t, *J* = 6.8 Hz, 3H). ^13^C NMR (126 MHz, CDCl_3_) δ 156.19, 147.79, 147.79 (based on HSQC and HMBC), 102.23, 79.62, 37.68, 31.43, 28.89, 28.41, 26.45, 22.40, 13.97. HRMS for C_14_H_25_N_3_O_2_Na [M+Na]^+^ calcd. 290.1839, found 290.1843.

#### Ethyl 1-pentyl-1*H*-imidazole-4-carboxylate (6b)

Compound **6b** was prepared from ethyl imidazole-4-carboxylate (2.10 mmol) using general protocol 3. Yield: 37%. *R*_f_ = 0.32 (100% EtOAc). ^1^H NMR (500 MHz, CDCl_3_) δ 7.59 (s, 1H), 7.46 (s, 1H), 4.35 (q, *J* = 7.1 Hz, 2H), 3.94 (t, *J* = 7.1 Hz, 2H), 1.79 (m, 2H), 1.37 (t, *J* = 7.1 Hz, 3H), 1.35–1.22 (m, 4H), 0.88 (t, *J* = 7.2 Hz, 3H). ^13^C NMR (126 MHz, CDCl_3_) δ 162.51, 137.81, 134.13, 124.94, 60.50, 47.56, 30.55, 28.52, 22.08, 14.42, 13.82. HRMS for C_11_H_18_N_2_O_2_Na [M+Na]^+^ calcd. 233.1260, found 233.1262.

#### Ethyl 1-pentyl-1*H*-pyrazole-4-carboxylate (6c)

Compound **6c** was prepared from ethyl 4-pyrazolecarboxylate (2.10 mmol) using general protocol 3. Yield: 48%, *R*_f_ = 0.55 (30% EtOAc in hexanes). ^1^H NMR (500 MHz, CDCl_3_) δ 7.89 (s, 1H), 7.86 (s, 1H), 4.28 (q, *J* = 7.1 Hz, 2H), 4.11 (t, *J* = 7.2 Hz, 2H), 1.87 (m, 2H), 1.37–1.31 (m, 5H), 1.26 (m, 2H), 0.89 (t, *J* = 7.2 Hz, 3H). ^13^C NMR (126 MHz, CDCl_3_) δ 163.12, 140.89, 132.30, 114.85, 60.11, 52.67, 29.76, 28.61, 22.15, 14.40, 13.87. HRMS for C_11_H_18_N_2_O_2_Na [M+Na]^+^ calcd. 233.1260, found 233.1268.

#### Methyl 1-pentyl-1*H*-1,2,4-triazole-3-carboxylate (6d)

Compound **6d** was prepared from methyl 1,2,4-triazole-3-carboxylate (3.93 mmol) using general protocol 3. Yield: 55%, *R*_f_ = 0.15 (50% EtOAc in hexanes). ^1^H NMR (500 MHz, CDCl_3_) δ 8.12 (s, 1H), 4.22 (t, *J* = 7.3 Hz, 2H), 3.98 (s, 3H), 1.92 (m, 2H), 1.39–1.26 (m, 4H), 0.88 (t, *J* = 7.2 Hz, 3H). ^13^C NMR (126 MHz, CDCl_3_) δ 160.20, 154.80, 144.29, 52.74, 50.65, 29.42, 28.49, 22.04, 13.81. HRMS for C_9_H_15_N_3_O_2_Na [M+Na]^+^ calcd. 220.1056, found 220.1059.

#### *tert*-Butyl ((3-pentylisoxazol-5-yl)methyl)carbamate (12e)

Compound **12e** was prepared from *N*-Boc-propargylamine (1.29 mmol) using general protocol 4. Yield: 99%, *R*_f_ = 0.32 (20% EtOAc in hexanes). ^1^H NMR (500 MHz, CDCl_3_) δ 6.01 (s, 1H), 4.97 (bs, 1H), 4.38 (d, *J* = 5.6 Hz, 2H), 2.62 (t, *J* = 7.6 Hz, 2H), 1.64 (m, 2H), 1.45 (s, 9H), 1.36–1.31 (m, 4H), 0.90 (t, *J* = 7.0 Hz, 3H). ^13^C NMR (126 MHz, CDCl_3_) δ 169.08, 164.29, 155.47, 101.45, 80.23, 36.62, 31.33, 28.32, 27.92, 25.97, 22.32, 13.93. HRMS for C_14_H_24_N_2_O_3_Na [M+Na]^+^ calcd. 291.1679, found 291.1679.

#### *tert*-Butyl ((3-butylisoxazol-5-yl)methyl)carbamate (12f)

Compound **12f** was prepared from *N*-Boc-propargylamine (1.29 mmol) using general protocol 4. Yield: 26%, *R*_f_ = 0.30 (20% EtOAc in hexanes). ^1^H NMR (500 MHz, CDCl_3_) δ 6.03 (s, 1H), 5.01 (bs, 1H), 4.41 (d, *J* = 5.2 Hz, 2H), 2.65 (t, *J* = 7.5Hz, 2H), 1.61-1.67 (m, 2H), 1.47 (s, 9H), 1.37-1.42 (m, 2H), 0.95 (t, *J* = 7.3 Hz, 3H). ^13^C NMR (126 MHz, CDCl_3_) δ 169.1, 164.3, 155.5, 101.4, 90.2, 35.6, 30.3, 28.3, 25.7, 22.3, 13.7. HRMS for C_13_H_22_N_2_O_3_Na [M+Na]^+^ calcd. 277.1523, found 277.1516.

#### *tert*-Butyl ((3-hexylisoxazol-5-yl)methyl)carbamate (12g)

Compound **12g** was prepared from *N*-Boc-propargylamine (1.50 mmol) using general protocol 4. Yield: 72%, *R*_f_ = 0.30 (20% EtOAc in hexanes). ^1^H NMR (500 MHz, CDCl_3_) δ 6.00 (s, 1H), 5.05 (bs, 1H), 4.36 (d, *J* = 5.8 Hz, 2H), 2.60 (t, *J* = 7.7Hz, 2H), 1.57-1.65 (m, 2H), 1.43 (s, 9H), 1.27-1.34 (m, 6H), 0.86 (t, *J* = 7.1 Hz, 3H). ^13^C NMR (126 MHz, CDCl_3_) δ 169.1, 164.3, 155.5, 101.4, 80.2, 36.6, 31.5, 28.8, 28.3, 28.2, 26.0, 22.5, 14.0. HRMS for C_15_H_26_N_2_O_3_Na [M+Na]^+^ calcd. 305.1835, found 305.1829.

#### *tert*-Butyl ((3-phenethylisoxazol-5-yl)methyl)carbamate (12h)

Compound **12h** was prepared from *N*-Boc-propargylamine (1.50 mmol) using general protocol 4. Yield: 27%, *R*_f_ = 0.35 (20% EtOAc in hexanes). ^1^H NMR (500 MHz, CDCl_3_) δ 7.33-7.30 (m, 2H), 7.26-7.21 (m, 3H), 5.99 (s, 1H), 4.99 (bs, 1H), 4.40 (d, *J* = 5.6 Hz, 2H), 3.01-2.97 (m, 4H), 1.48 (s, 9H). ^13^C NMR (126 MHz, CDCl_3_) δ 169.3, 163.4, 155.5, 140.6, 128.5, 128.3, 126.3, 101.6, 80.3, 36.6, 34.3, 28.3, 27.9. HRMS for C_17_H_22_N_2_O_3_Na [M+Na]^+^ calcd. 325.1523, found 325.1520.

#### *tert*-Butyl (2-(3-pentylisoxazol-5-yl)ethyl)carbamate (12i)

Compound **12i** was prepared from *N*-Boc-3-butynylamine (2.20 mmol) using general protocol 4. Yield: 72%, *R*_f_ = 0.33 (20% EtOAc in hexanes). ^1^H NMR (500 MHz, CDCl_3_) δ 5.92 (s, 1H), 4.75 (bs, 1H), 3.50-3.46 (m, 2H), 2.95 (t, *J* = 6.6 Hz, 2H), 2.64 (t, *J* = 7.6 Hz, 2H), 1.63-1.69 (m, 2H), 1.46 (s, 9H), 1.34-1.37 (m, 4H), 0.92 (t, *J* = 7.0 Hz, 3H). ^13^C NMR (126 MHz, CDCl_3_) δ 170.2, 164.3, 155.8, 101.5, 79.6, 38.5, 31.4, 28.4, 28.0, 27.6, 26.0, 22.3, 13.9. HRMS for C_15_H_26_N_2_O_3_Na [M+Na]^+^ calcd. 305.1836, found 305.1832.

#### *tert*-Butyl ((5-pentylisoxazol-3-yl)methyl)carbamate (15j)

Compound **15j** was prepared from commercial 1-heptyne (0.90 mmol) using general protocol 4. Yield: 23%, *R*_f_ = 0.77 (50% EtOAc in hexanes). ^1^H NMR (500 MHz, CDCl_3_) δ 5.96 (s, 1H), 4.99 (bs, 1H), 4.34 (d, *J* = 5.8 Hz, 2H), 2.70 (t, *J* = 7.6 Hz, 2H), 1.68 (m, 2H), 1.46 (s, 9H), 1.35–1.32 (m, 4H), 0.90 (t, *J* = 7.0 Hz, 3H). ^13^C NMR (126 MHz, CDCl_3_) δ 174.30, 161.57, 155.78, 99.86, 79.93, 36.69, 31.20, 28.35, 27.14, 26.68, 22.27, 13.90. HRMS for C_14_H_24_N_2_O_3_Na [M+Na]^+^ calcd. 291.16791, found 291.16834.

#### *tert*-Butyl ((5-butylisoxazol-3-yl)methyl)carbamate (15k)

Compound **15k** was prepared from commercial 1-hexyne (2.50 mmol) using general protocol 4. Yield: 8%, *R*_f_ = 0.63 (50% EtOAc in hexanes). ^1^H NMR (500 MHz, CDCl_3_) δ 5.98 (s, 1H), 5.01 (bs, 1H), 4.37 (d, *J* = 5.6 Hz, 2H), 2.74 (t, *J* = 7.1 Hz, 2H), 1.65-1.71 (m, 2H), 1.48 (s, 9H), 1.37-1.44 (m, 2H), 0.96 (t, *J* = 7.4 Hz, 3H). ^13^C NMR (126 MHz, CDCl_3_) δ 174.3, 161.6, 155.8, 99.9, 79.9, 36.7, 29.5, 29.4, 26.4, 22.2, 13.7. HRMS for C_13_H_22_N_2_O_3_Na [M+Na]^+^ calcd. 277.1523, found 277.1526.

#### *tert*-Butyl ((5-hexylisoxazol-3-yl)methyl)carbamate (15l)

Compound **15l** was prepared from commercial 1-octyne (4.09 mmol) using general protocol 4. Yield: 11%, *R*_f_ = 0.65 (50% EtOAc in hexanes). ^1^H NMR (500 MHz, CDCl_3_) δ 5.98 (s, 1H), 5.02 (bs, 1H), 4.37 (d, *J* = 5.3 Hz, 2H), 2.73 (t, *J* = 7.5 Hz, 2H), 1.66-1.72 (m, 2H), 1.48 (s, 9H), 1.35-1.40 (m, 2H), 1.30-1.33 (m, 4H), 0.91 (t, *J* = 6.9 Hz, 3H). ^13^C NMR (126 MHz, CDCl_3_) δ 174.3, 161.6, 155.8, 99.9, 79.9, 36.7, 31.4, 28.7, 28.4, 27.4, 26.7, 22.5, 14.0. HRMS for C_15_H_26_N_2_O_3_Na [M+Na]^+^ calcd. 305.1836; found 305.1836.

#### *tert*-Butyl ((5-phenethylisoxazol-3-yl)methyl)carbamate (15m)

Compound **15m** was prepared from commercial 4-phenyl-1-butyne (6.00 mmol) using general protocol 4. Yield: 10%, *R*_f_ = 0.66 (50% EtOAc in hexanes). ^1^H NMR (500 MHz, CDCl_3_) δ 7.28-7.32 (m, 2H), 7.19-7.25 (m, 3H), 5.96 (s, 1H), 5.11 (bs, 1H), 4.35 (d, *J* = 5.8 Hz, 2H), 3.00-3.07 (m, 4H), 1.48 (s, 9H). ^13^C NMR (126 MHz, CDCl_3_) δ 172.93, 161.70, 155.81, 139.99, 128.57, 128.26, 126.47, 100.42, 79.90, 36.63, 33.52, 28.51, 28.36. HRMS for C_17_H_22_N_2_O_3_Na [M+Na]^+^ calcd. 325.1522, found 325.1522.

#### *tert*-Butyl ((2-pentyl-2*H*-1,2,3-triazol-4-yl)methyl)carbamate (17)

Compound **17** was prepared from *N*-Boc-propargylamine (1.93 mmol) as described. Yield: 34%, *R*_f_ = 0.45 (30% EtOAc in hexanes). ^1^H NMR (500 MHz, CDCl_3_) δ 7.49 (s, 1H), 4.94 (bs, 1H), 4.38–4.34 (m, 4H), 1.93 (m, 2H), 1.45 (s, 9H), 1.38–1.23 (m, 4H), 0.89 (t, *J* = 7.2 Hz, 3H). ^13^C NMR (126 MHz, CDCl_3_) δ 155.71, 145.38, 132.33, 79.74, 54.92, 36.16, 29.43, 28.64, 28.37, 22.13, 13.88. HRMS for C_13_H_24_N_4_O_2_Na [M+Na]^+^ calcd. 291.1791, found 291.1799.

#### *tert*-Butyl ((5-pentyl-4*H*-1,2,4-triazol-3-yl)methyl)carbamate (19)

Compound **19** was prepared from *N*-Boc-glycinamide (2.18 mmol) as described. Yield: 52%, *R*_f_ = 0.15 (60% EtOAc in hexanes). ^1^H NMR (500 MHz, CDCl_3_) δ 5.42 (bs, 1H), 4.40 (d, *J* = 5.7 Hz, 2H), 2.74 (t, *J* = 7.7 Hz, 2H), 1.75 (m, 2H), 1.44 (s, 9H), 1.37–1.32 (m, 4H), 0.89 (t, *J* = 7.1 Hz, 3H). ^13^C NMR (126 MHz, CDCl_3_) δ 160.79, 158.20, 156.50, 80.20, 37.67, 31.56, 28.47, 27.88, 27.36, 22.44, 14.05. HRMS for C_13_H_25_N_4_O_2_ [M+H]^+^ calcd. 269.19720, found 269.19651.

#### *tert*-Butyl ((5-pentyl-1,3,4-thiadiazol-2-yl)methyl)carbamate (21)

Compound **21** was prepared from *N*-Boc glycine (1.14 mmol) as described. Yield: 60%, *R*_f_ = 0.38 (50% EtOAc in hexanes). ^1^H NMR (500 MHz, CDCl_3_) δ 5.27 (bs, 1H), 4.67 (d, *J* = 6.1 Hz, 2H), 3.06 (t, *J* = 7.6, 2H), 1.78 (m, 2H), 1.46 (s, 9H), 1.41–1.32 (m, 4H), 0.90 (t, *J* = 7.1 Hz, 3H). ^13^C NMR (126 MHz, CDCl_3_) δ 172.11, 167.87, 155.71, 80.65, 39.91, 31.26, 30.24, 29.86, 28.46, 22.39, 14.02. HRMS for C_13_H_23_N_3_O_2_SNa [M+Na]^+^ calcd. 308.1403, found 308.1401.

#### *tert*-Butyl ((5-pentyl-1,3,4-oxadiazol-2-yl)methyl)carbamate (22)

Compound **22** was prepared from *N*-Boc glycine (0.88 mmol) as described. Yield: 31%, *R*_f_ = 0.41 (50% EtOAc in hexanes). ^1^H NMR (500 MHz, CDCl_3_) δ 5.10 (bs, 1H), 4.52 (d, *J* = 5.1 Hz, 2H), 2.82 (t, *J* = 7.6, 2H), 1.78 (m, 2H), 1.46 (s, 9H), 1.39–1.33 (m, 4H), 0.90 (t, *J* = 7.1 Hz, 3H). ^13^C NMR (126 MHz, CDCl_3_) δ 167.89, 163.98, 155.53, 80.70, 36.01, 31.25, 28.43, 26.24, 25.44, 22.31, 13.98. HRMS for C_13_H_23_N_3_O_3_Na [M+Na]^+^ calcd. 292.16316, found 292.16219.

#### *tert*-Butyl ((3-pentyl-1,2,4-oxadiazol-5-yl)methyl)carbamate (24r)

Compound **24r** was prepared from hexanenitrile (2.06 mmol) using general protocol 5. Yield: 36%, *R*_f_ = 0.57 (40% EtOAc in hexanes). ^1^H NMR (500 MHz, CDCl_3_) δ 5.13 (bs, 1H), 4.55 (d, *J* = 5.3 Hz, 2H), 2.71 (t, *J* = 7.7, 2H), 1.75 (m, 2H), 1.46 (s, 9H), 1.37–1.32 (m, 4H), 0.90 (t, *J* = 7.1 Hz, 3H). ^13^C NMR (126 MHz, CDCl_3_) δ 176.18, 170.89, 155.50, 80.81, 37.29, 31.39, 28.42, 26.75, 26.04, 22.41, 14.05. HRMS for C_13_H_23_N_3_O_3_Na [M+Na]^+^ calcd. 292.1632, found 292.1639.

#### *tert*-Butyl ((5-pentyl-1,2,4-oxadiazol-3-yl)methyl)carbamate (24s)

Compound **24s** was prepared from *N*-(tert-butoxycarbonyl)-2-aminoacetonitrile (1.92 mmol) using general protocol 5. Yield: 45%, *R*_f_ = 0.70 (50% EtOAc in hexanes). ^1^H NMR (500 MHz, CDCl_3_) δ 5.12 (bs, 1H), 4.44 (d, *J* = 5.1 Hz, 2H), 2.86 (t, *J* = 7.6 Hz, 2H), 1.80 (m, 2H), 1.45 (s, 9H), 1.39–1.31 (m, 4H), 0.90 (t, *J* = 7.1 Hz, 3H). ^13^C NMR (126 MHz, CDCl_3_) δ 180.72, 168.00, 155.67, 80.27, 37.00, 31.25, 28.46, 26.65, 26.34, 22.28, 13.96. HRMS for C_13_H_23_N_3_O_3_Na [M+Na]^+^ calcd. 292.1632, found 292.1641.

#### *tert*-Butyl ((2-pentyloxazol-5-yl)methyl)carbamate (25t)

Compound **25t** was prepared from *N*-Boc-propargylamine (0.38 mmol) using general protocol 6. Yield: 36%, *R*_f_ = 0.39 (40% EtOAc in hexanes). ^1^H NMR (400 MHz, CDCl_3_) δ 6.80 (s, 1H), 4.81 (bs, 1H), 4.30 (d, *J* = 5.2 Hz, 2H), 2.71 (t, *J* = 7.6 Hz, 2H), 1.74 (m, 2H), 1.45 (s, 9H), 1.37–1.31 (m, 4H), 0.89 (t, *J* = 7.0 Hz, 3H). ^13^C NMR (126 MHz, CDCl_3_) δ 165.15, 155.56, 148.55, 123.94, 80.15, 35.70, 31.43, 28.49, 28.30, 26.77, 22.42, 14.06. HRMS for C_14_H_24_N_2_O_3_Na [M+Na]^+^ calcd. 291.1679, found 291.1666.

#### *tert*-Butyl ((5-pentyloxazol-2-yl)methyl)carbamate (25u)

Compound **25u** was prepared from 1-heptyne (0.72 mmol) using general protocol 6. Yield: 21%, *R*_f_ = 0.56 (50% EtOAc in hexanes). ^1^H NMR (500 MHz, CDCl_3_) δ 6.64 (s, 1H), 5.11 (bs, 1H), 4.39 (d, *J* = 5.0 Hz, 2H), 2.60 (t, *J* = 7.4 Hz, 2H), 1.61 (m, 2H), 1.46 (s, 9H), 1.36–1.30 (m, 4H), 0.90 (t, *J* = 7.0 Hz, 3H). ^13^C NMR (126 MHz, CDCl_3_) δ 159.94, 155.70, 153.78, 122.09, 80.15, 38.27, 31.36, 28.47, 27.29, 25.58, 22.44, 14.08. HRMS for C_14_H_24_N_2_O_3_Na [M+Na]^+^ calcd. 291.1679, found 291.1669.

#### 5-Pentylthiophene-2-carbaldehyde (26)

Compound **26** was prepared from 2-pentyl thiophene (2.50 mmol) as described. Yield: 82%. The characterization data is as previously reported.^59 1^H NMR (500 MHz, CDCl_3_) δ 9.84 (s, 1H), 7.63 (d, *J* = 3.75 Hz, 1H), 6.93 (d, *J* = 3.75 Hz, 1H), 2.89 (t, *J* = 7.70 Hz, 2H), 1.71-1.77 (m, 2H), 1.36-1.39 (m, 4H), 0.93 (t, *J* = 7.05 Hz, 3H). ^13^C NMR (126 MHz, CDCl_3_) δ 182.66, 157.80, 141.60, 137.02, 125.83, 31.17, 30.96, 30.82, 22.33, 13.94.

#### 5-Pentylthiophene-2-carboxamide (27)

Compound **27** was prepared from **26** (1.50 mmol) as described. Yield: 52%. ^1^H NMR (500 MHz, CDCl_3_) δ 7.39 (d, *J* = 3.7 Hz, 1H), 6.80 (d, *J* = 3.7 Hz, 1H), 5.66 (bs, 2H), 2.84 (t, *J* = 7.6Hz, 2H), 1.68-1.74 (m, 2H), 1.35-1.39 (m, 4H), 0.92 (t, *J* = 7.1 Hz, 3H). ^13^C NMR (126 MHz, CDCl_3_) δ 163.66, 152.59, 134.59, 129.52, 125.05, 31.16, 31.13, 30.36, 22.36, 13.96. HRMS for C_10_H_16_NOS [M+H]^+^ calcd. 198.0947, found 198.0952.

## ASSOCIATED CONTENT

### Supporting Information

The following files are available free of charge.

S1 Biological results, purity studies and NMR spectrum (file type, word) S2 SMILE (file type, excel)

## AUTHOR INFORMATION

### Author Contributions

The manuscript was drafted by JG and CS, and all authors contributed to editing and finalizing the document. We confirm that all authors have approved submission of the final version of this manuscript.

### Notes

The authors declare no competing financial interest. Animal experiments were approved by the Australian National University Animal Experimentation Ethics Committee (Protocol number A2017/50).

## ACKNOWLEDGMENT

The China Scholarship Council is thanked for providing a scholarship to J.G. The Australian Government provided scholarships for E.T.T. and V.M.H., whereas C.S. was funded by an NHMRC Overseas Biomedical Fellowship (1016357). We thank the Centre for Drug Candidate Optimisation, Monash Institute of Pharmaceutical Sciences, Monash University, for ADME analyses. We are also grateful to the Canberra branch of the Australian Red Cross Blood Service for providing red blood cells and human serum.

## ABBREVIATIONS

ADME: absorption, distribution metabolism and excretion
CDI: carbonyldiimidazole
CoA: coenzyme A
DCM: dichloromethane
DMF: N,N-dimethylformamide
DHA: dihydroartemisin
EDAC: N-(3-dimethylaminopropyl)-N’-ethylcarbodiimide hydrochloride
EEDQ: 2-ethoxy-1-ethoxycarbonyl-1,2-dihydroquinoline
HFF: human foreskin fibroblast;
SAR: structure-activity relationship
TEA: triethylamine
TFA: trifluoroacetic acid
THF: tetrahydrofuran

