## Supplemental Methods, Figures and Tables for "Exploring heteroaromatic rings as a replacement for the labile amide of antiplasmodial pantothenamides"

1. Biological Results S2
2. Purity determination S6
3. NMR spectra S8

### 1. Biological results

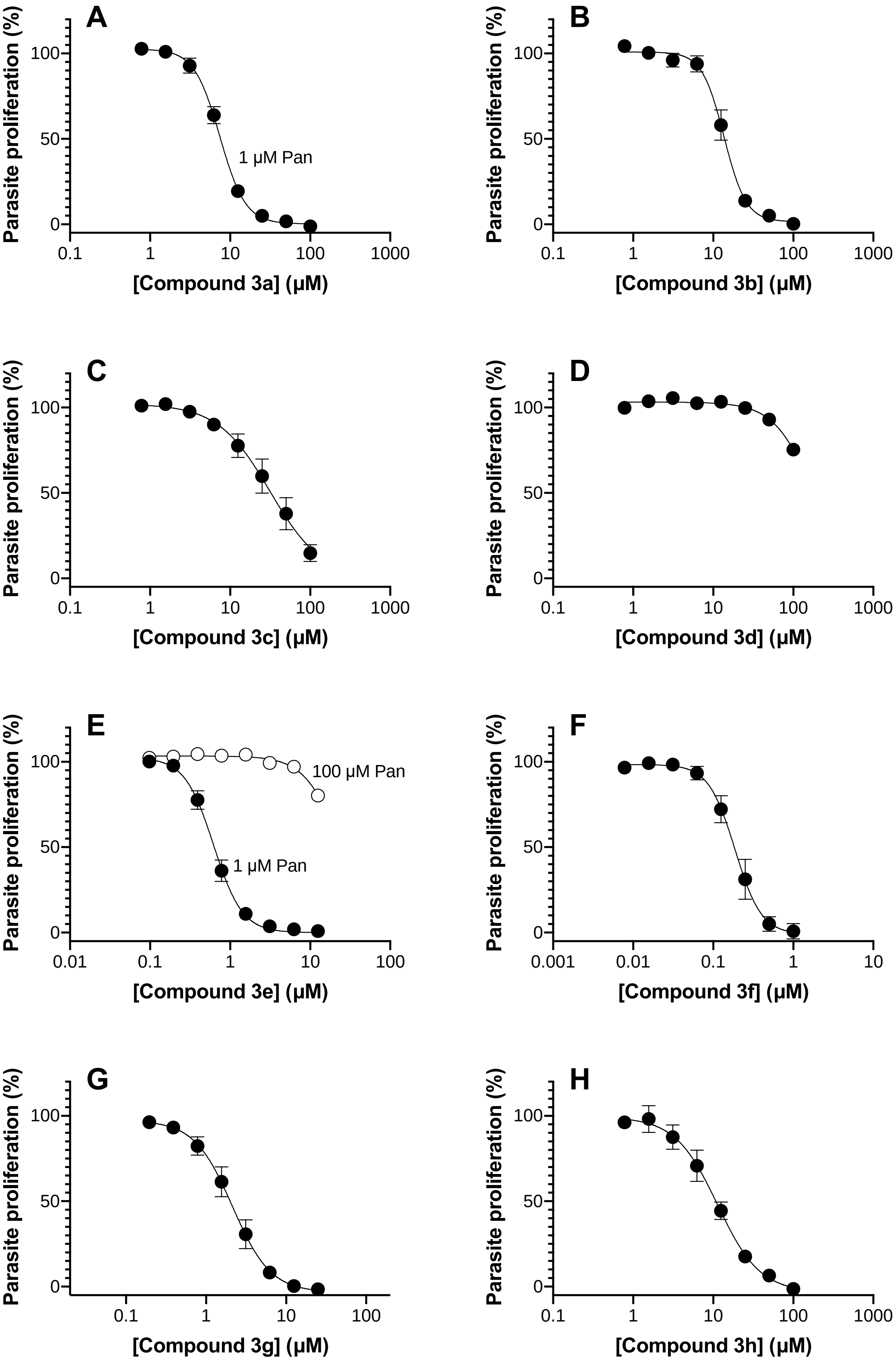

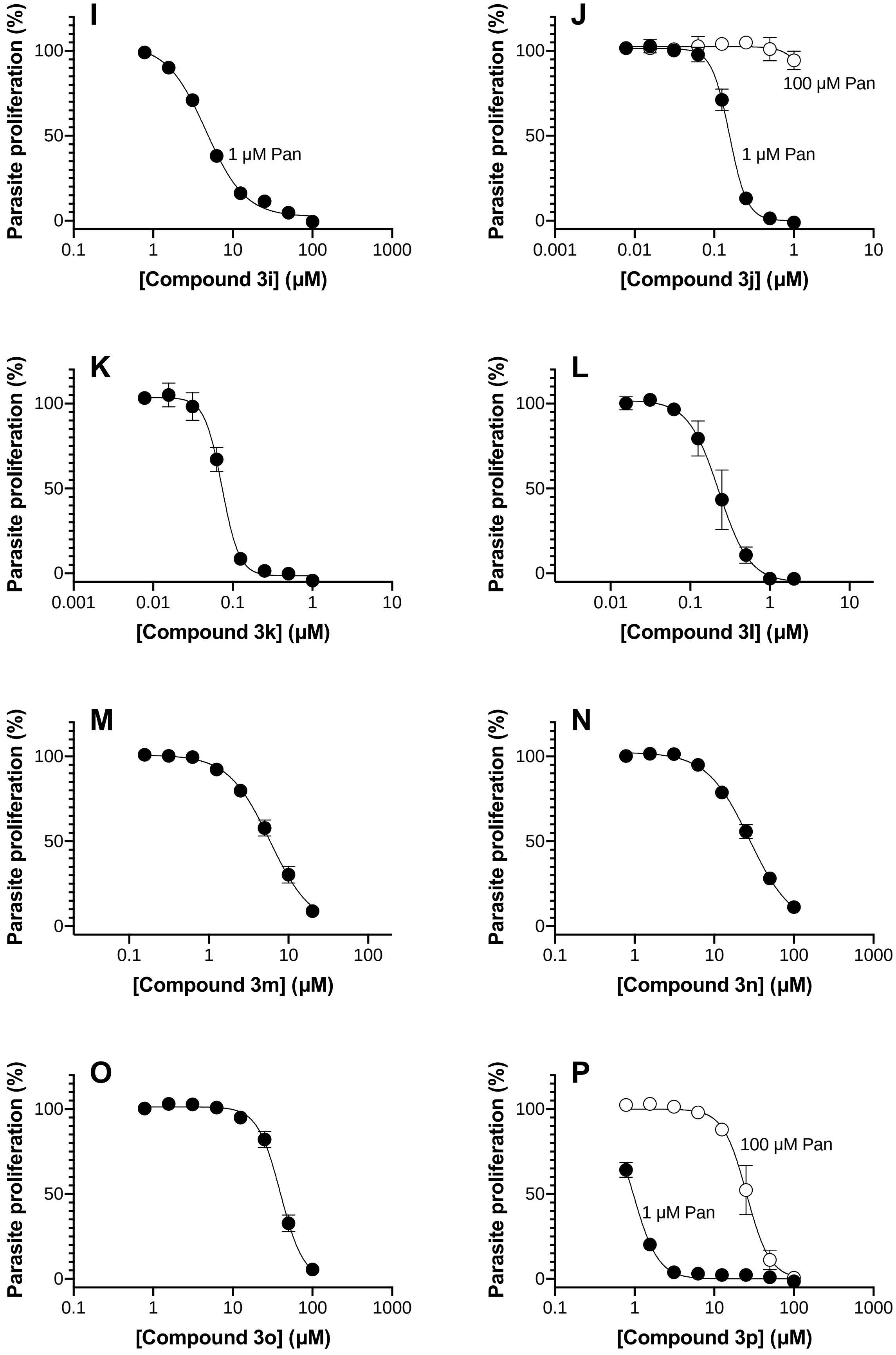
I
Parasite proliferation (%)
100
50
0
1 µM Pan
0.1
1
10
100
1000
[Compound 3i] (µM)
J
Parasite proliferation (%)
100
50
0
100 µM Pan
1 µM Pan
0.001
0.01
0.1
1
10
[Compound 3j] (µM)
K
Parasite proliferation (%)
100
50
0
0.001
0.01
0.1
1
10
[Compound 3k] (µM)
L
Parasite proliferation (%)
100
50
0
0.01
0.1
1
10
[Compound 3l] (µM)
M
Parasite proliferation (%)
100
50
0
0.1
1
10
100
[Compound 3m] (µM)
N
Parasite proliferation (%)
100
50
0
0.1
1
10
100
1000
[Compound 3n] (µM)
O
Parasite proliferation (%)
100
50
0
0.1
1
10
100
1000
[Compound 3o] (µM)
P
Parasite proliferation (%)
100
50
0
100 µM Pan
1 µM Pan
0.1
1
10
100
1000
[Compound 3p] (µM)

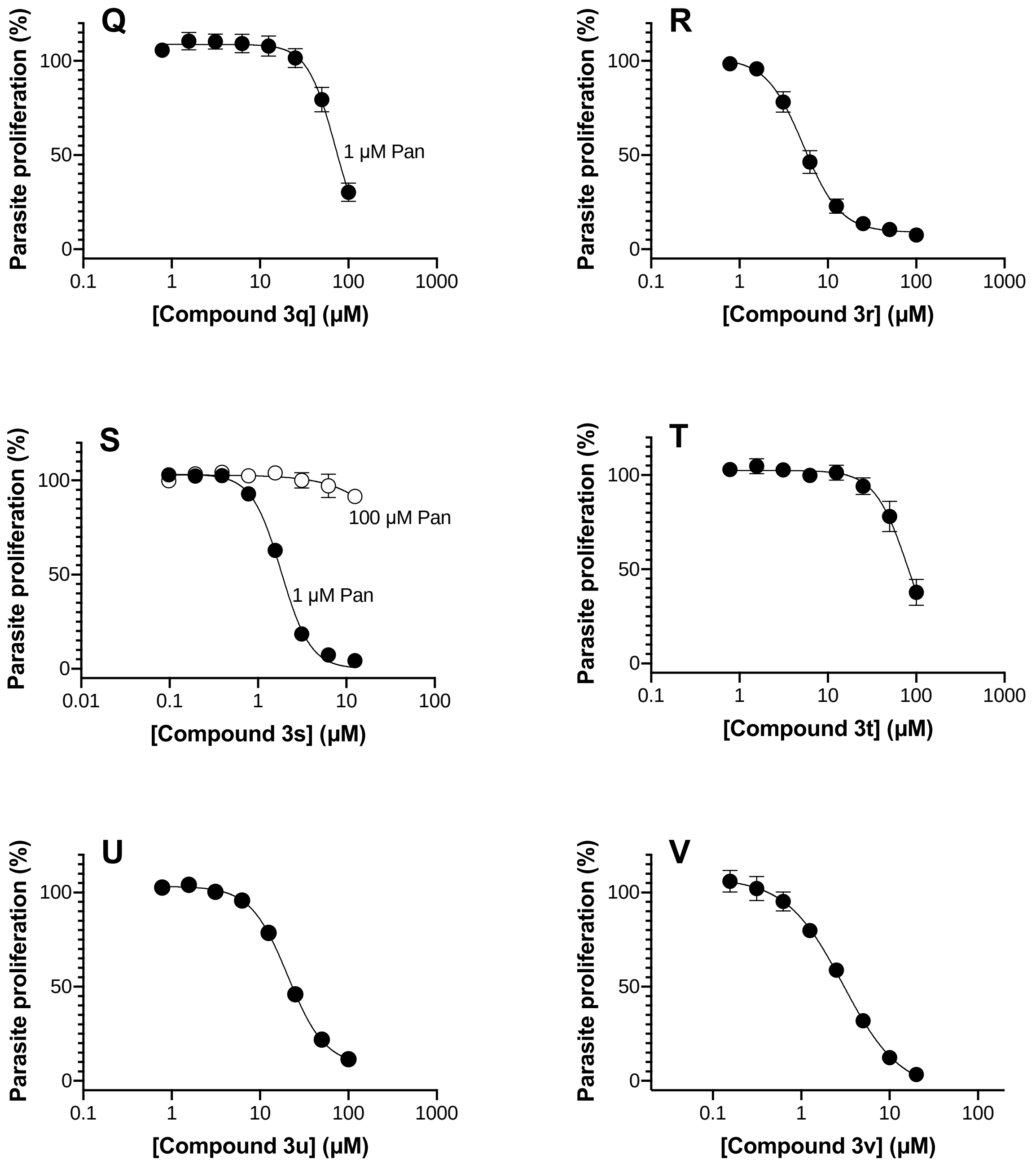


**Figure S1.** Antiplasmodial activity of analogs of triazole **1** against *P. falciparum in vitro* in the presence of 1 µM (closed symbols) or 100 µM (open symbols) pantothenate (Pan). Data generated in the presence of 1 µM pantothenate are averages from three or more experiments and error bars represent SEM, except in the case of compound **3p**, where the data are from two experiments and error bars represent range/2. Data generated in the presence of 100 µM pantothenate are averages from two independent experiments and error bars represent range/2. Where not visible, error bars are smaller than the symbol.

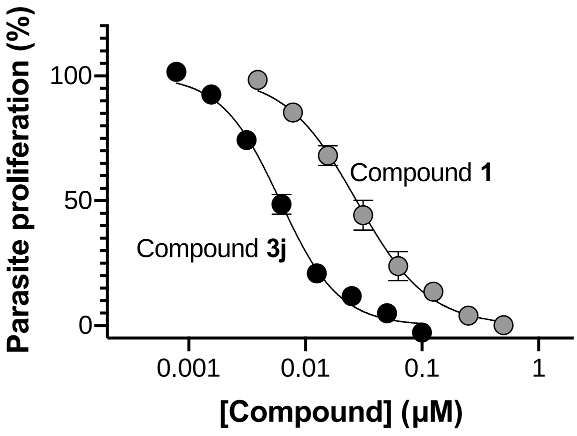


**Figure S2.** Antiplasmodial activity of compounds **1** (grey circles) and **3j** (black circles) against *P. knowlesi in vitro* in the presence of 1 μM pantothenate. Data are averages from three experiments and error bars represent SEM. Where not visible, error bars are smaller than the symbol.

### 2. Purity determination

Purity of the final products is reported using two independent reversed-phase HPLC analytical methods within Methods A-D. The samples were prepared by dissolving each compound in HPLC-grade acetonitrile to a concentration of 1 mg/mL.

Method A
Flow rate: 0.5 mL/min
Injection volume: 15-30 µL
Column: Agilent ZORBAX Eclipse XDB-C8 150 × 4.60 mm, 5 µm
Detection: 214 or 220 nm

| Time (min) | Water (%) | Acetonitrile (%) |
|---|---|---|
| 0 | 99 | 1 |
| 10 | 50 | 50 |
| 12 | 1 | 99 |
| 13 | 1 | 99 |
| 15 | 99 | 1 |

Method B
Flow rate: 0.5 mL/min
Injection volume: 15-30 µL
Column: Agilent ZORBAX RX-C18 150 × 4.60 mm, 5 µm
Detection: 214 or 220 nm

| Time (min) | Water (%) | Acetonitrile (%) |
|---|---|---|
| 0 | 99 | 1 |
| 10 | 1 | 99 |
| 20 | 1 | 99 |
| 23 | 99 | 1 |
| 25 | 99 | 1 |

Method C
Flow rate: 0.5 mL/min
Injection volume: 15 µL
Column: Phenomenex Synergi Hydro-RP 80 Å 250 × 4.60 mm, 4 µm
Detection: 214 nm

| Time (min) | Water (%) | Acetonitrile (%) |
|---|---|---|
| 0 | 99 | 1 |
| 10 | 1 | 99 |
| 20 | 1 | 99 |
| 23 | 99 | 1 |
| 25 | 99 | 1 |

Method D
Flow rate: 0.5 mL/min
Inject volume: 15 µL
Column: Phenomenex Luna C18 100 Å 250 × 4.60 mm, 5 µm
Detection: 214 nm

| Time (min) | Water (%) | Acetonitrile (%) |
|---|---|---|
| 0 | 99 | 1 |
| 10 | 1 | 99 |
| 20 | 1 | 99 |
| 23 | 99 | 1 |
| 25 | 99 | 1 |

### 3. NMR spectra

#### $^1$H NMR of compound 3a

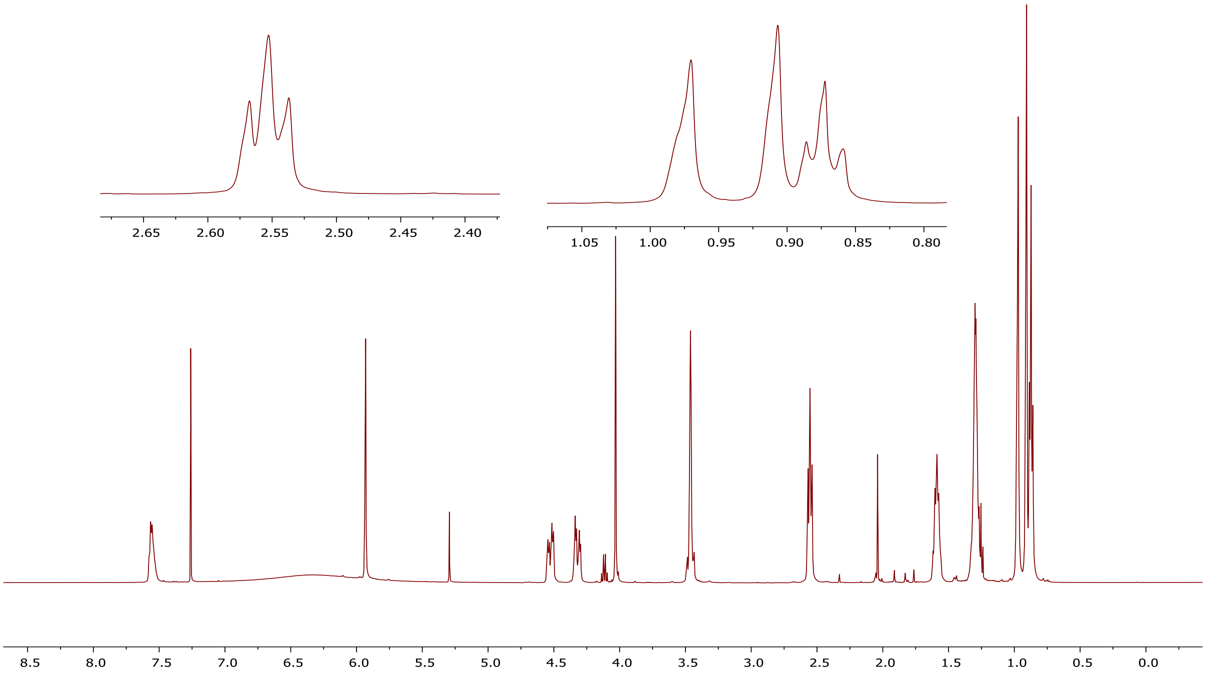


#### $^{13}$C NMR of compound 3a

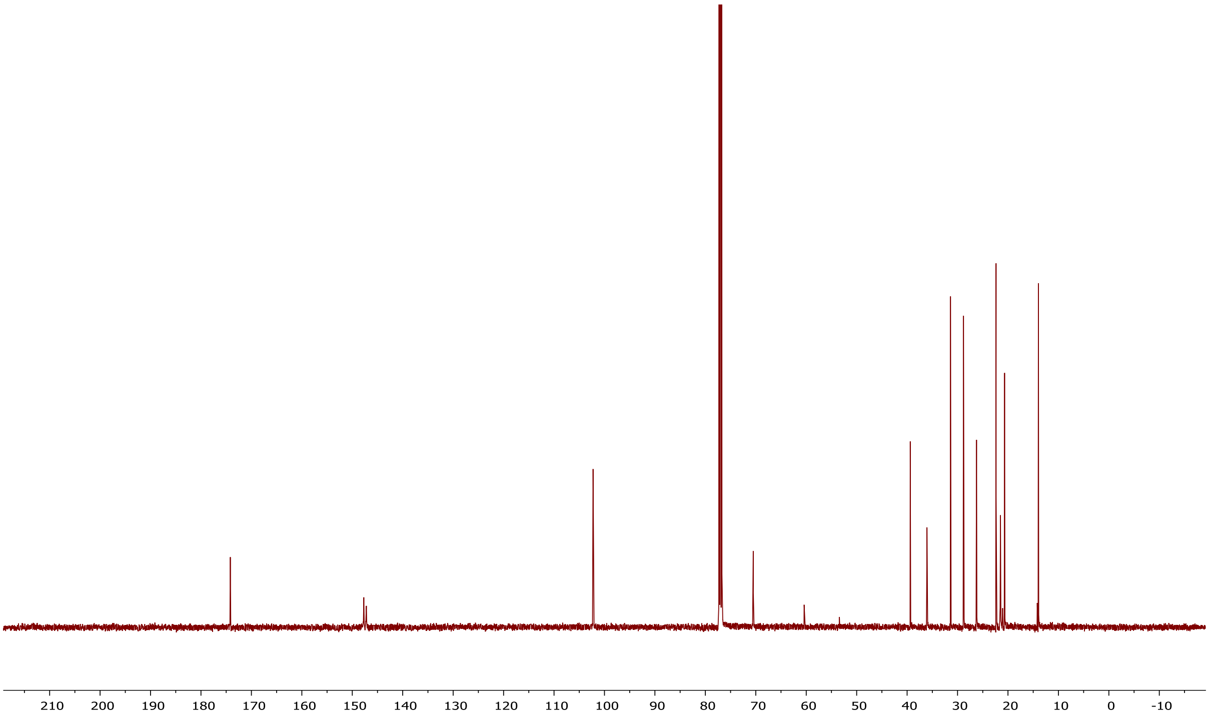

[1]H NMR of compound 3b

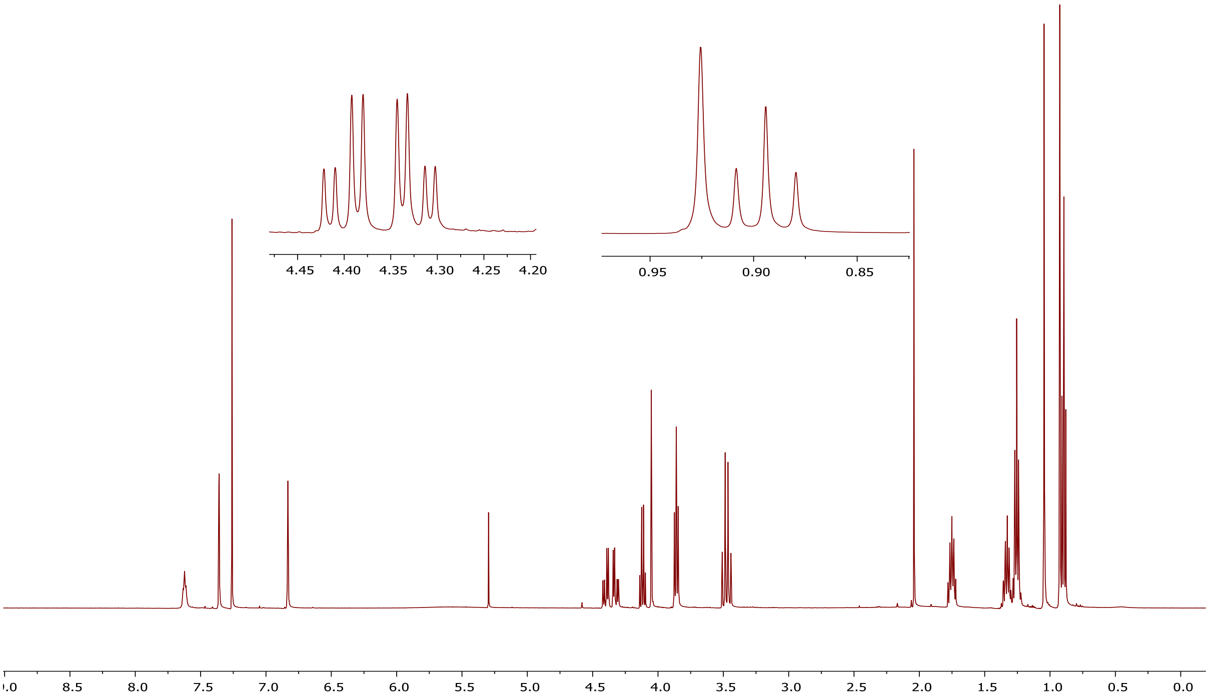


[13]C NMR of compound 3b

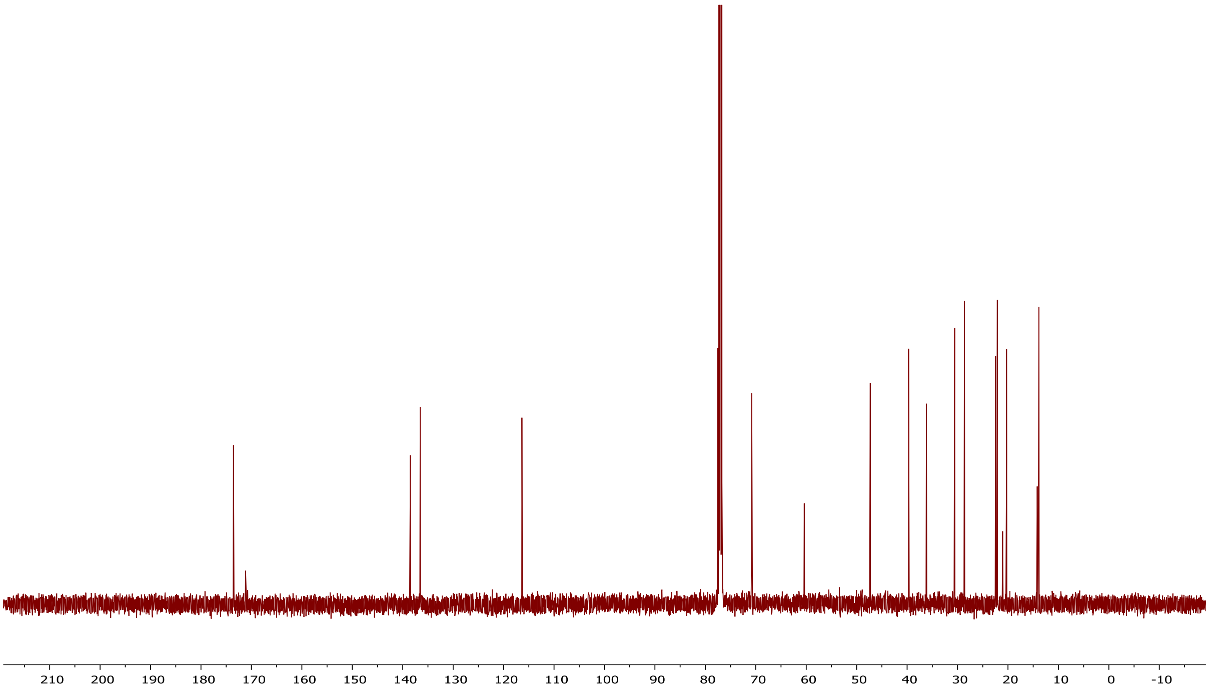

### $^{1}$H NMR of compound 3c

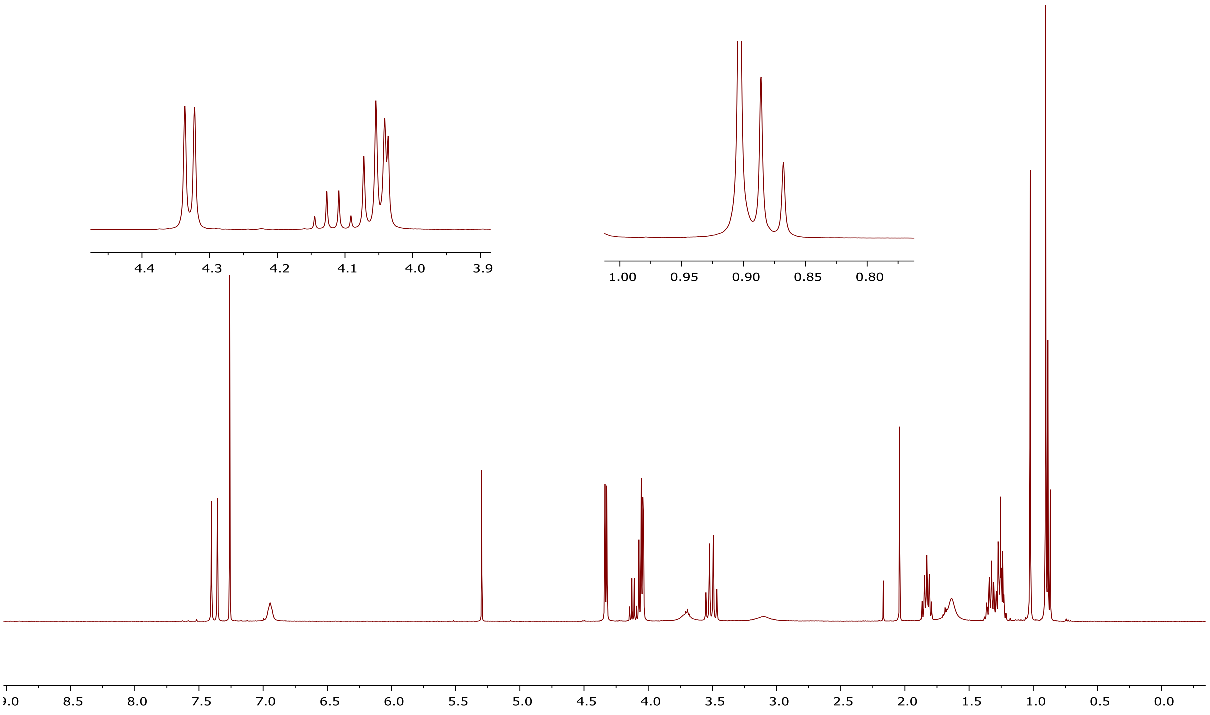


### $^{13}$C NMR of compound 3c

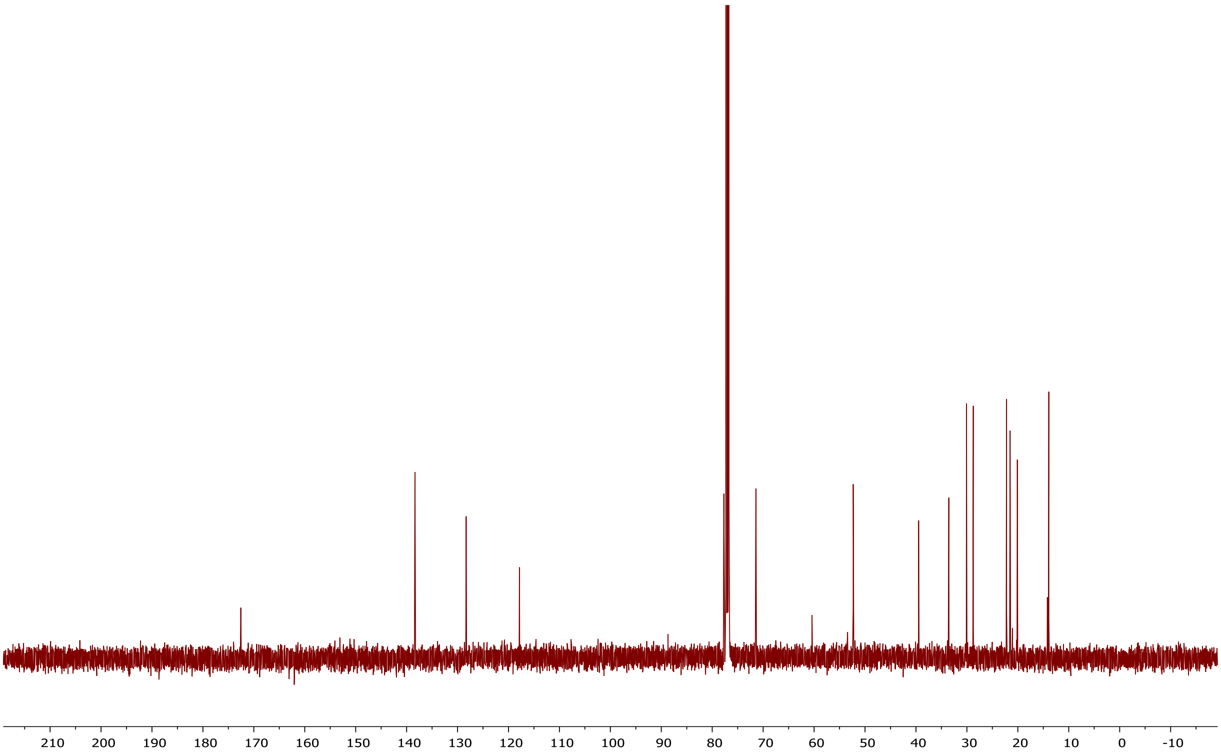

### $^{1}H$ NMR of compound 3d

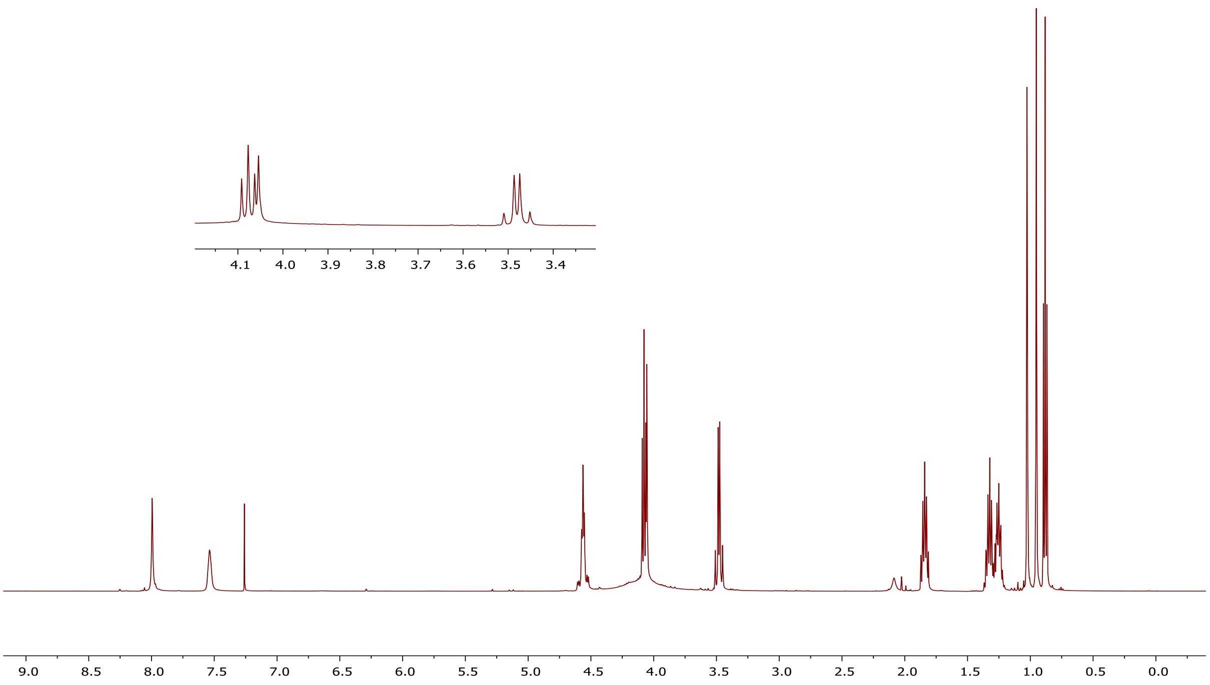


### $^{13}C$ NMR of compound 3d

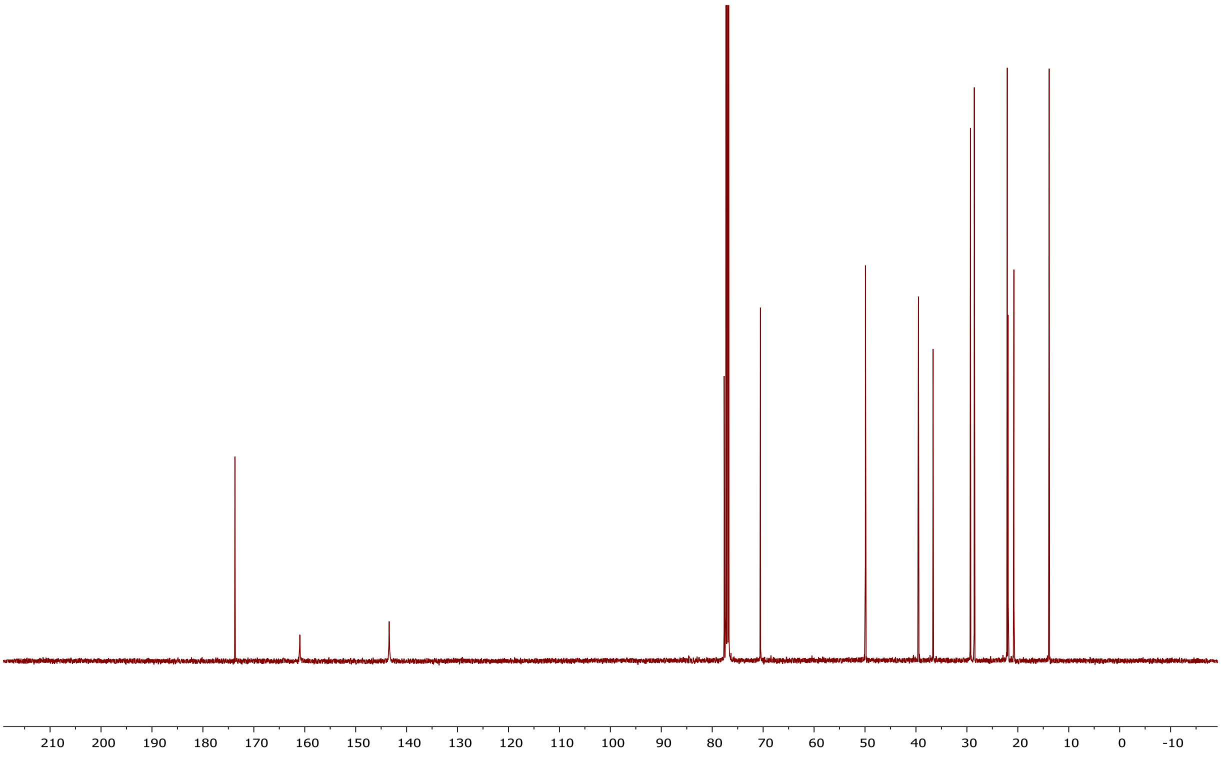

### [1]H NMR of compound 3e

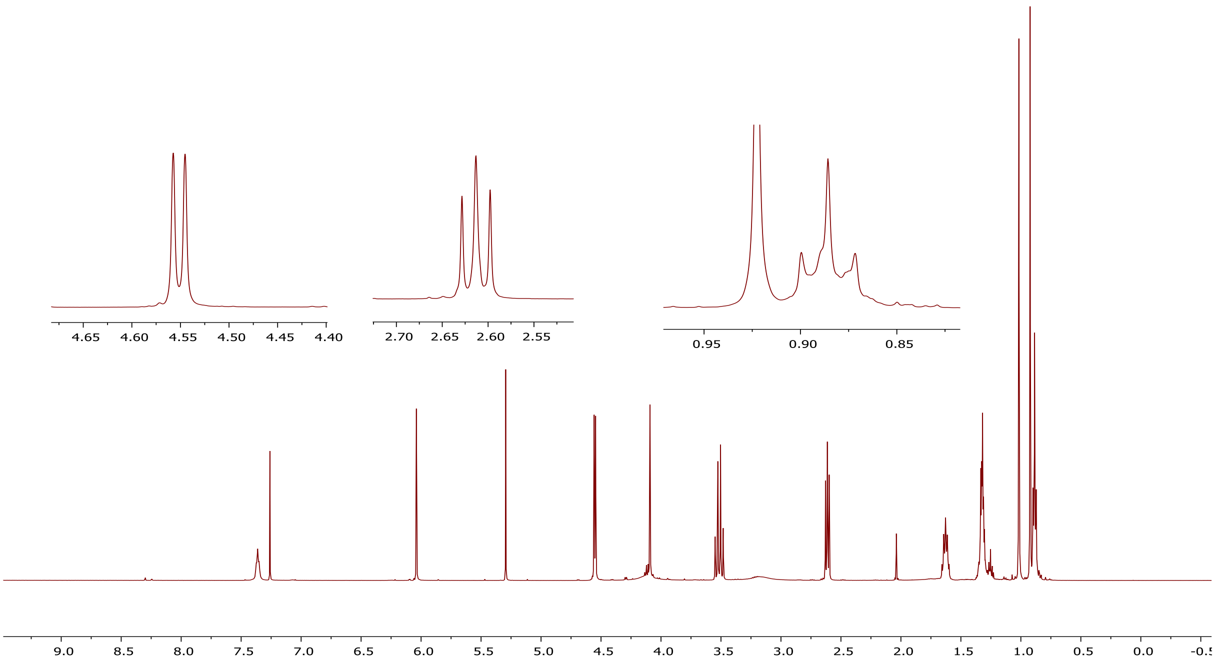


### [13]C NMR of compound 3e

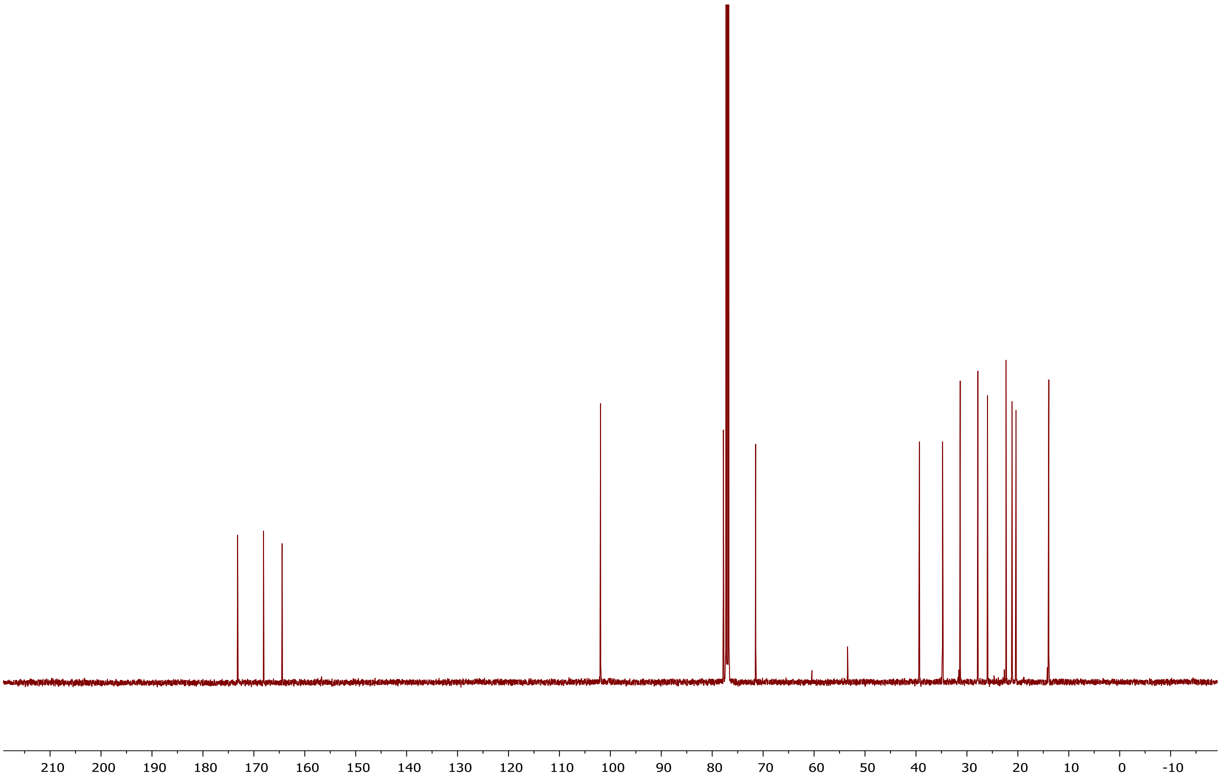

### [1]H NMR of compound 3f

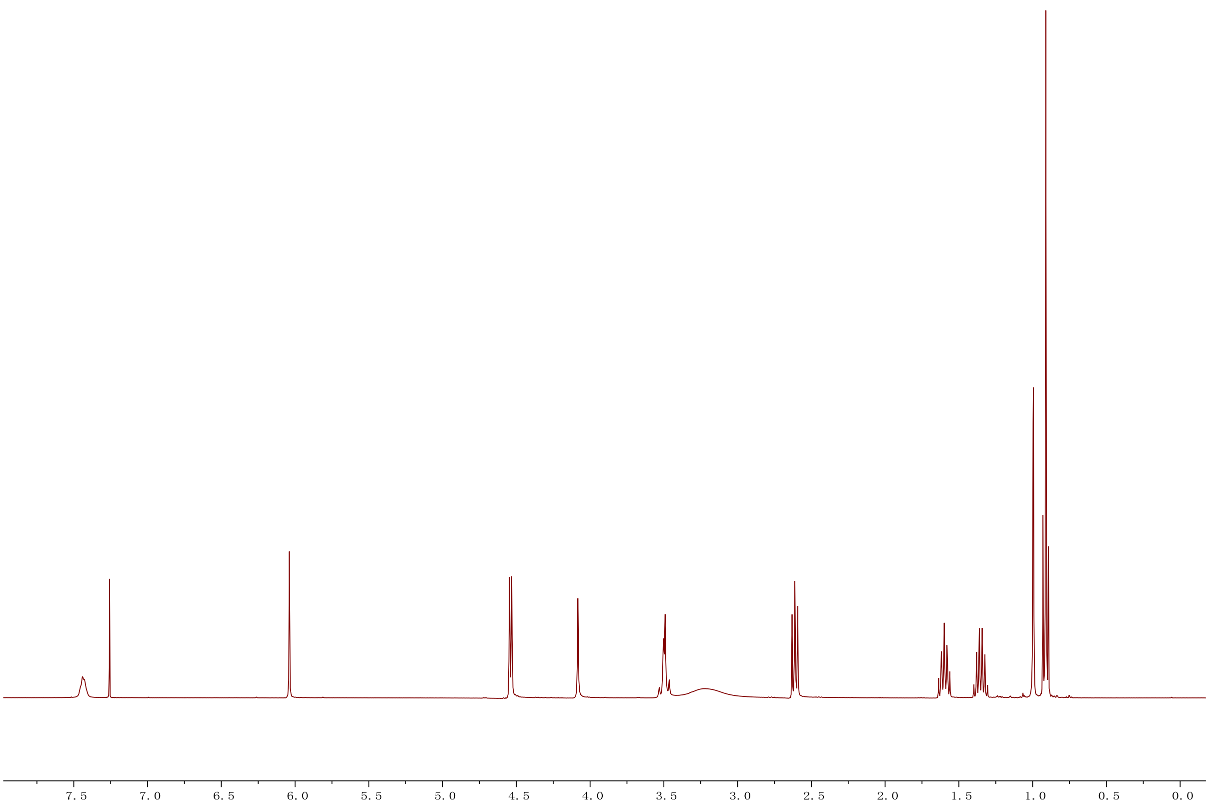


### [13]C NMR of compound 3f

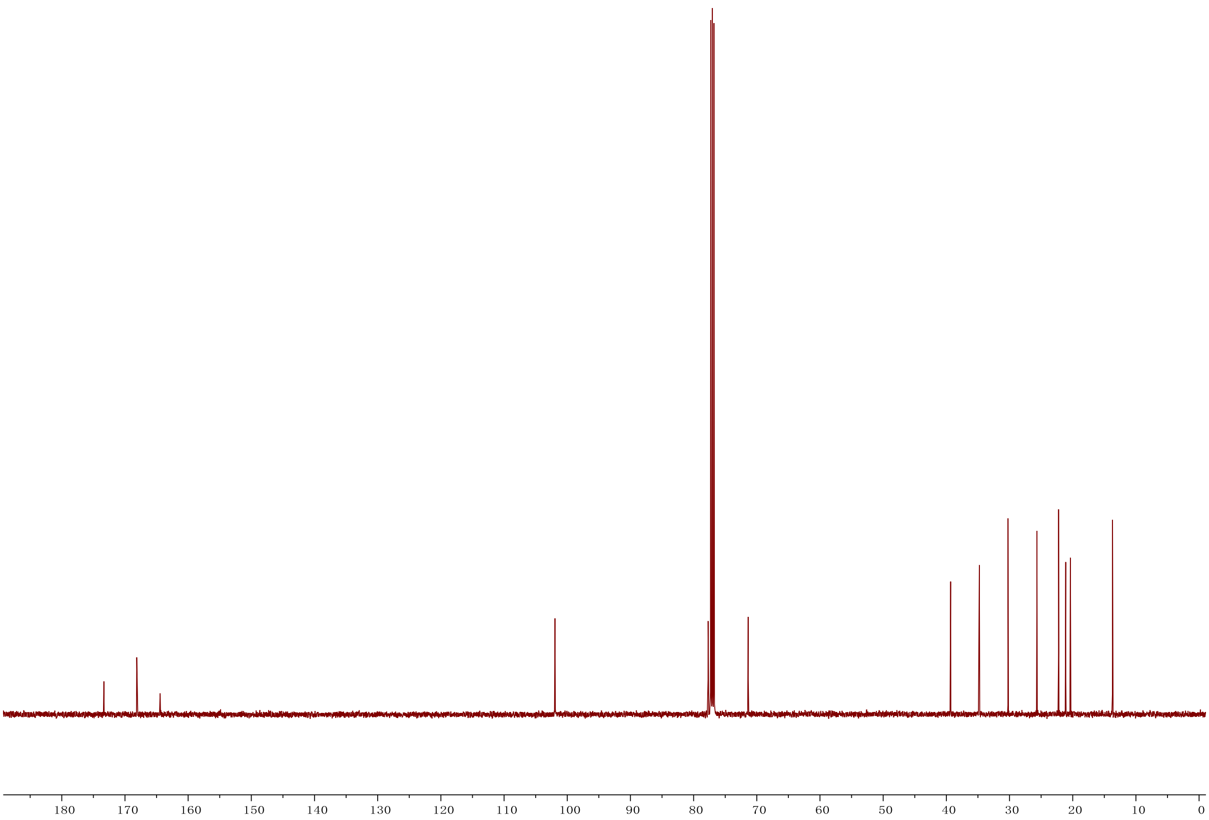

### [1]H NMR of compound 3g

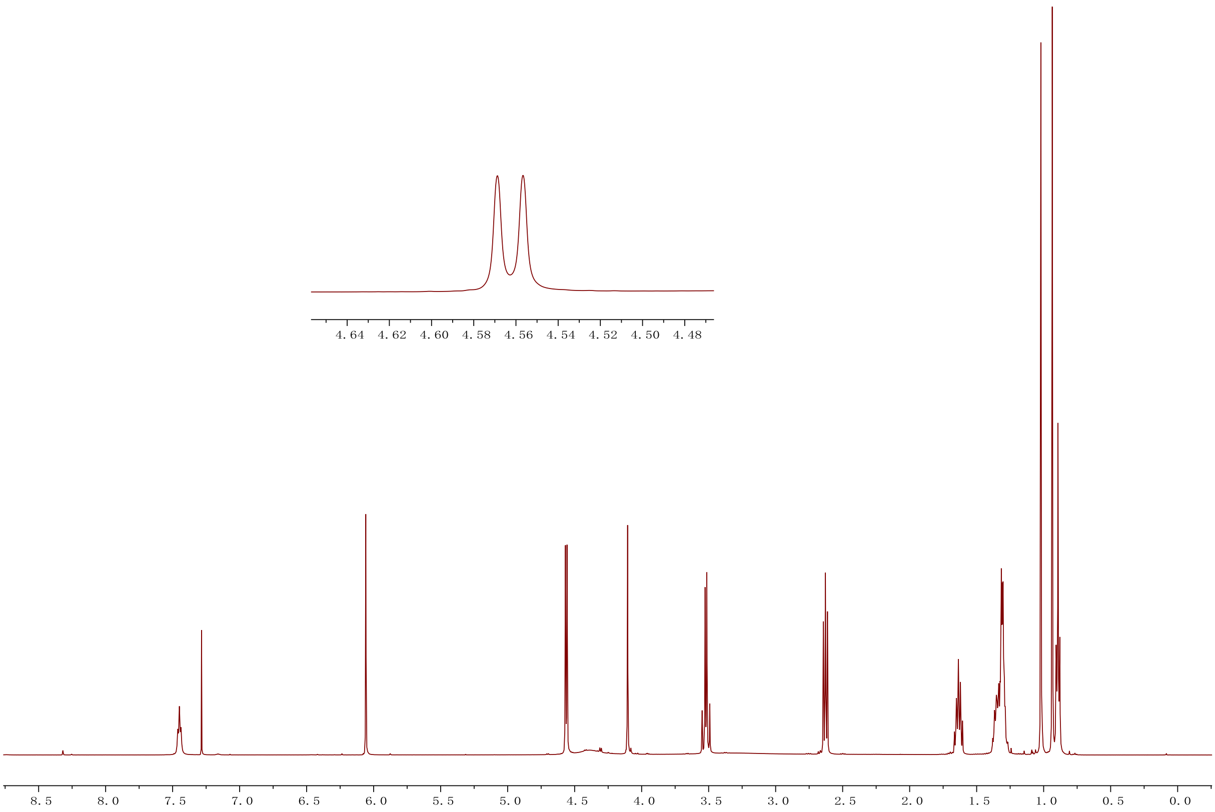


### [13]C NMR of compound 3g

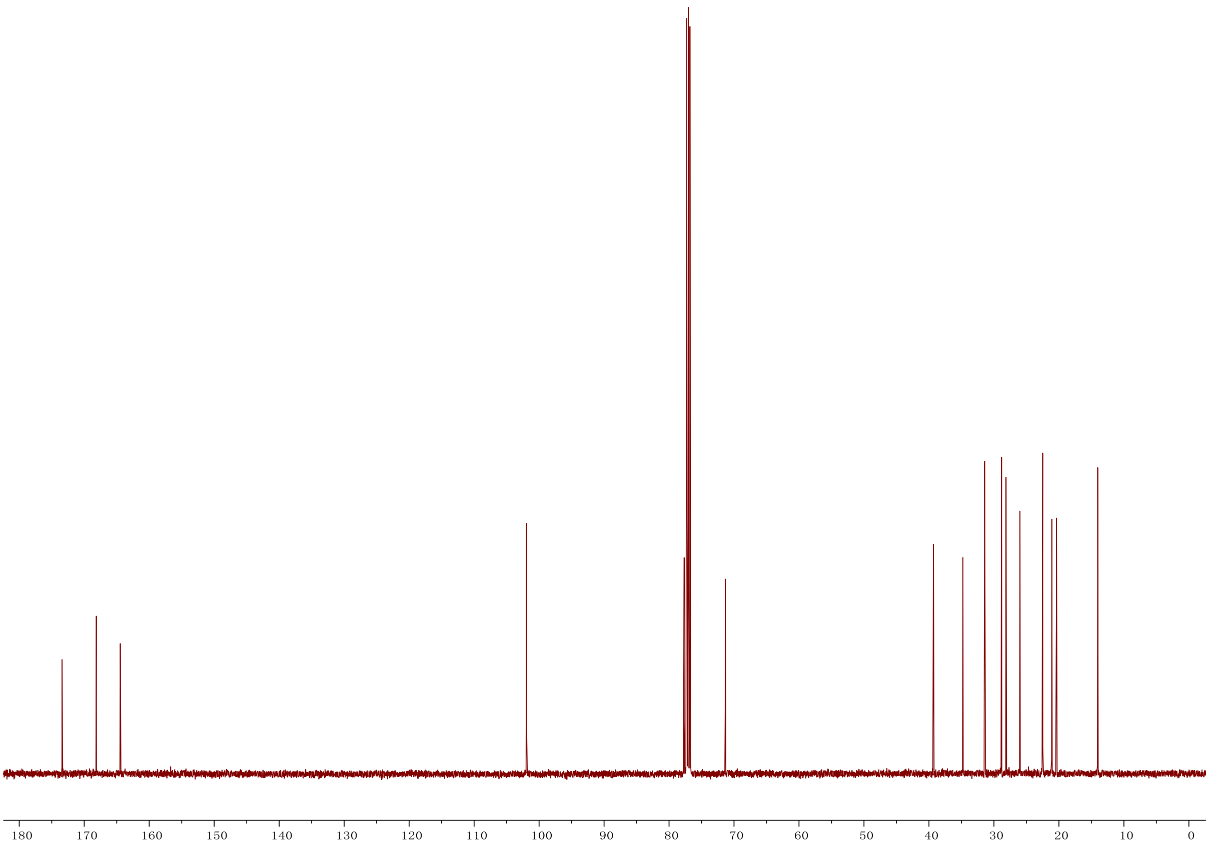

**[1]H NMR of compound 3h**

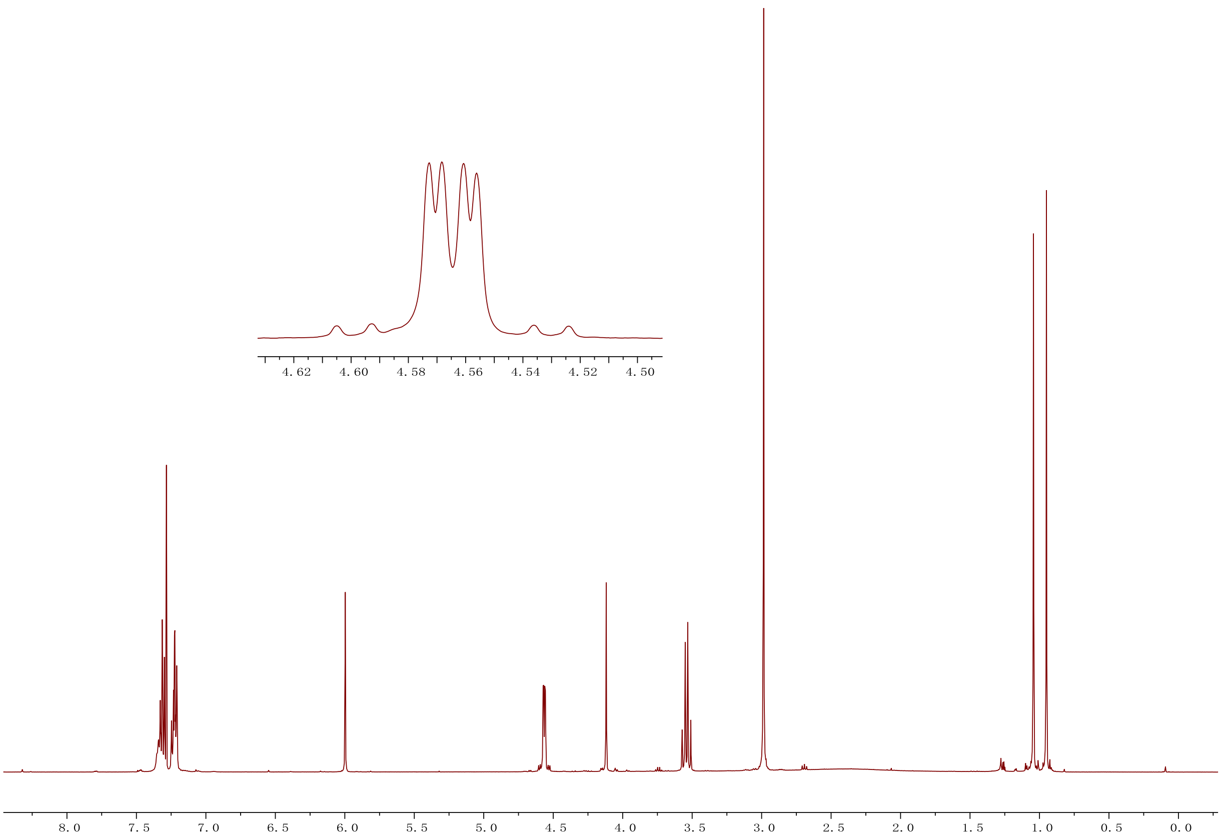


**[13]C NMR of compound 3h**

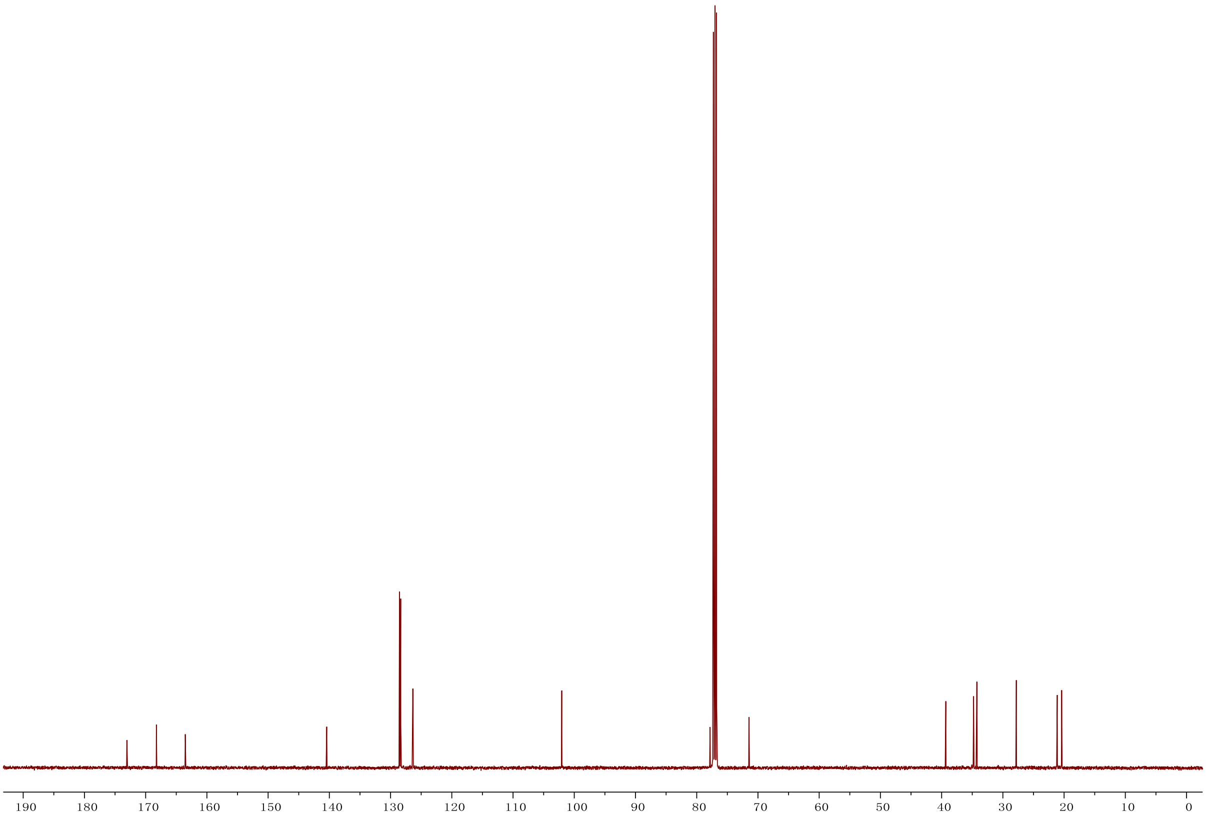

$^1$H NMR of compound 3i

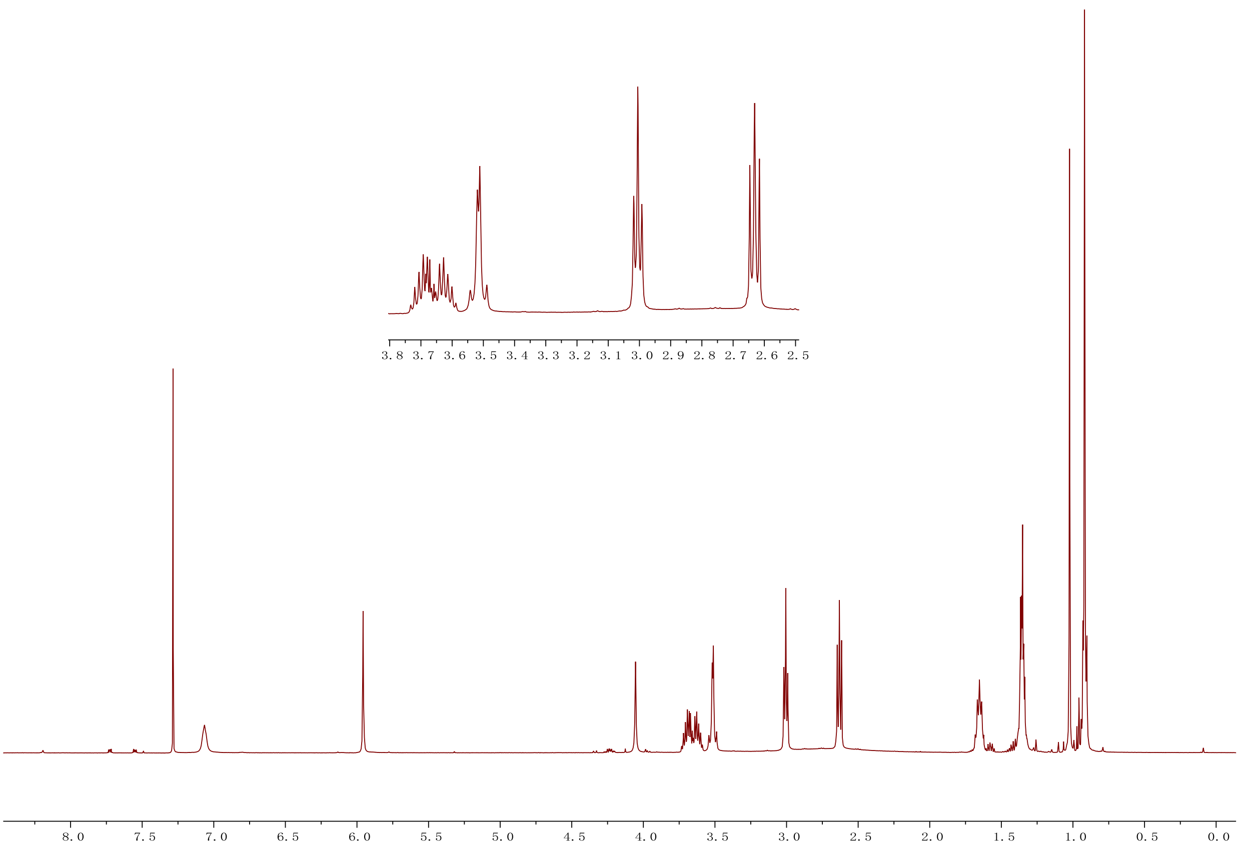


$^{13}$C NMR of compound 3i

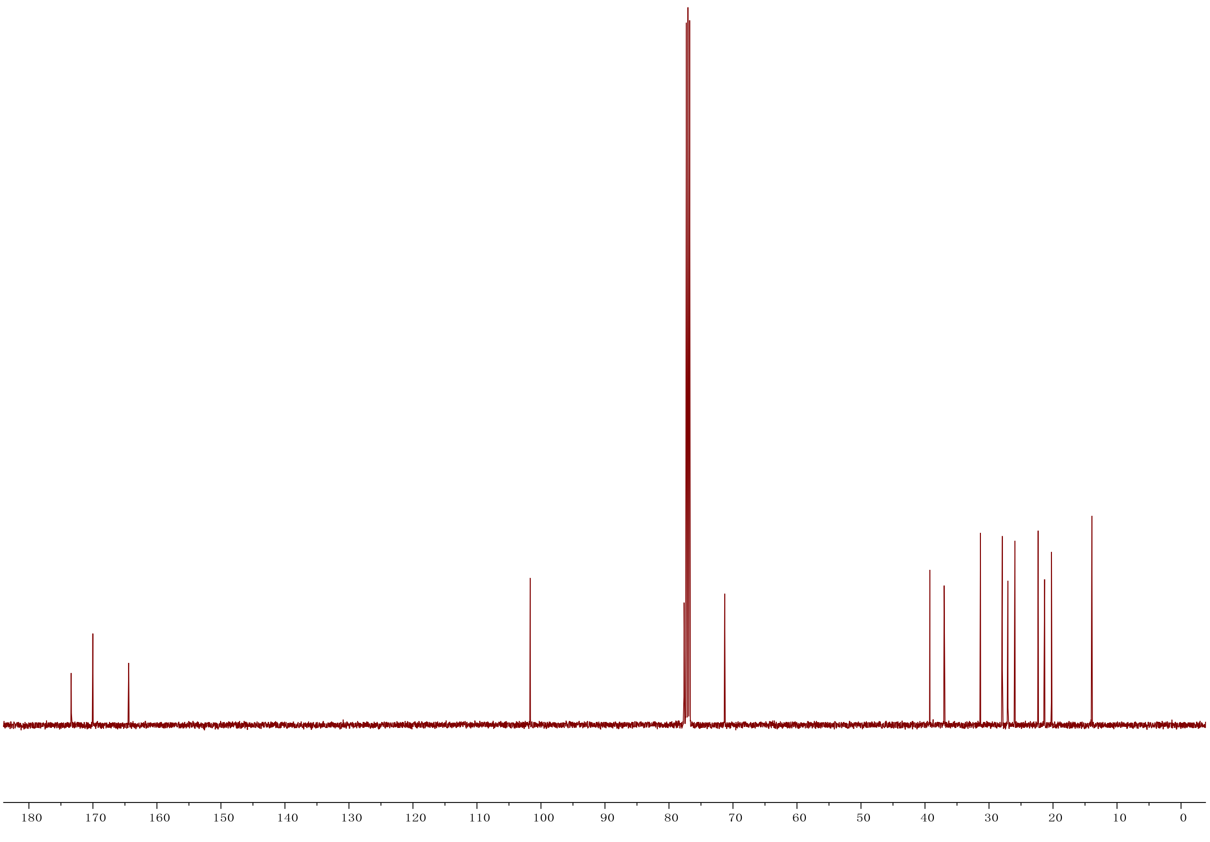

### [1]H NMR of compound 3j

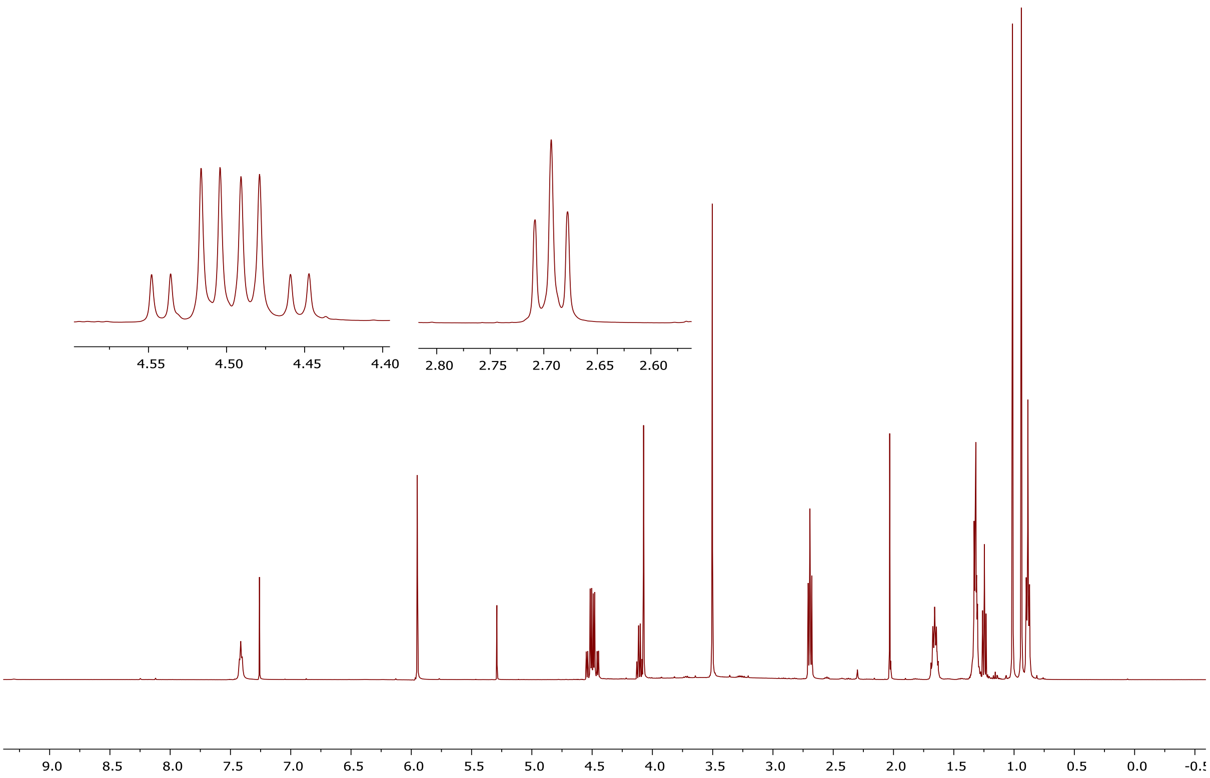


### [13]C NMR of compound 3j

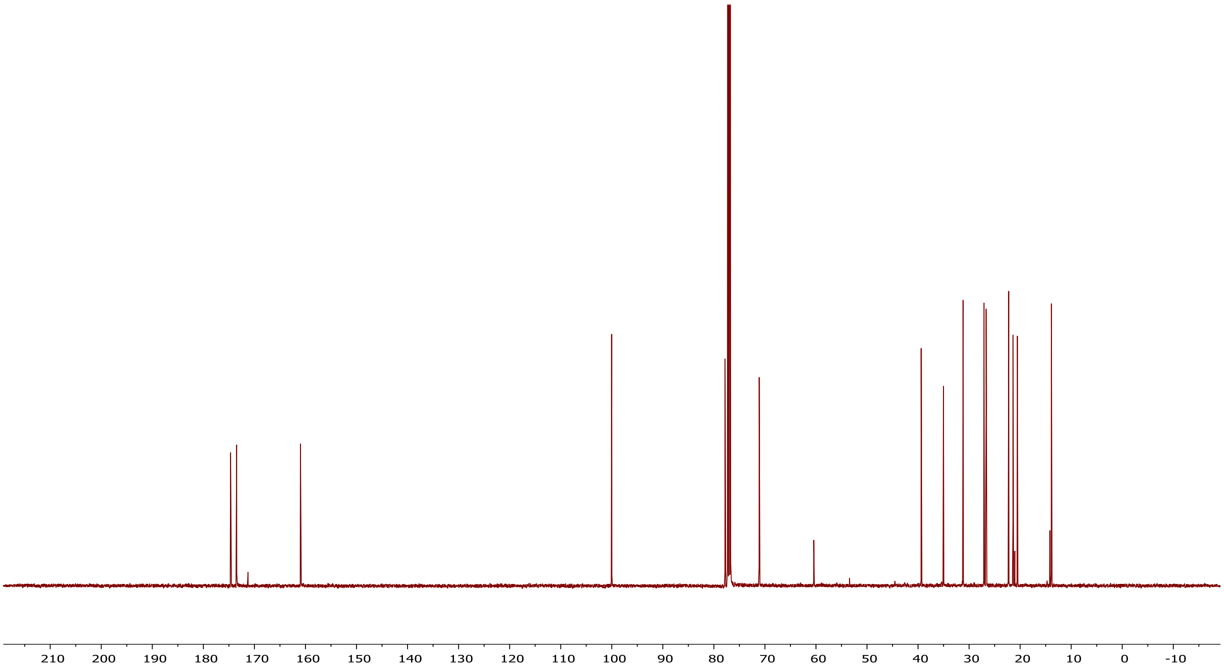

### $^{1}$H NMR of compound 3k

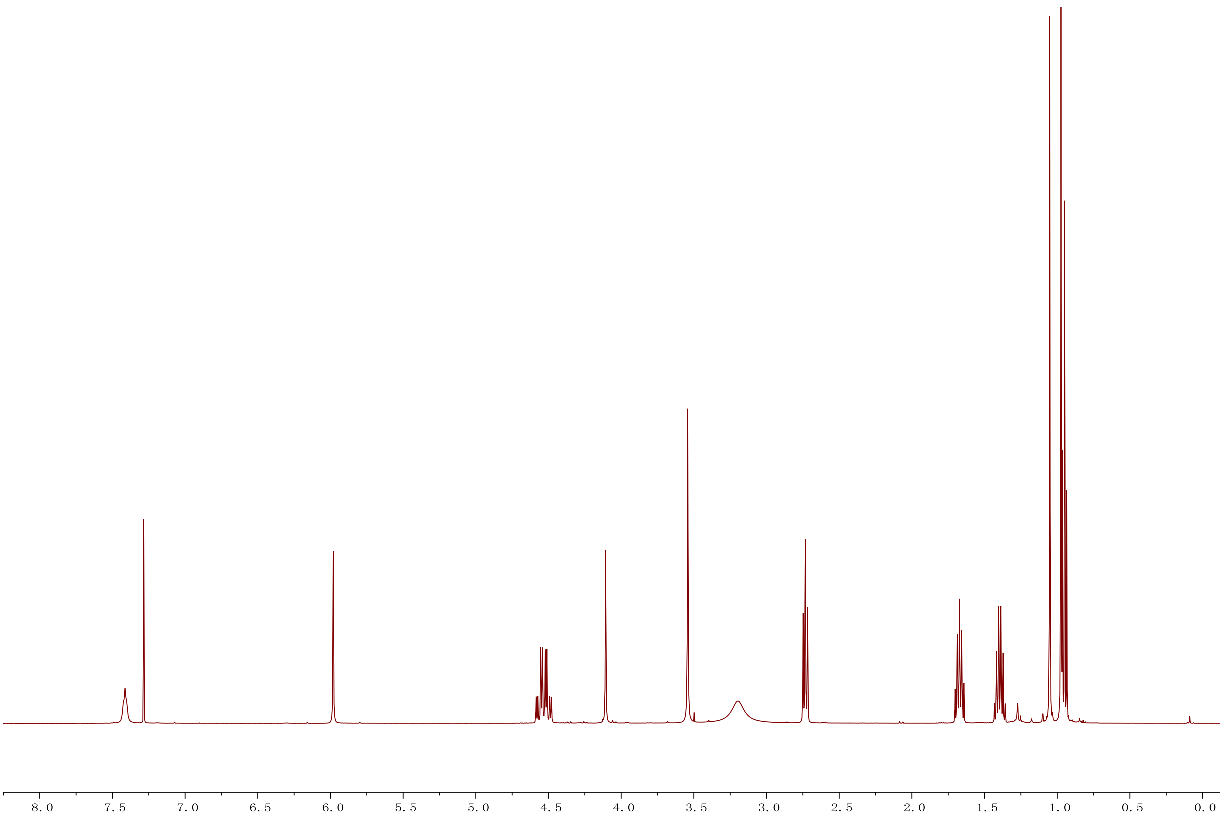


### $^{13}$C NMR of compound 3k

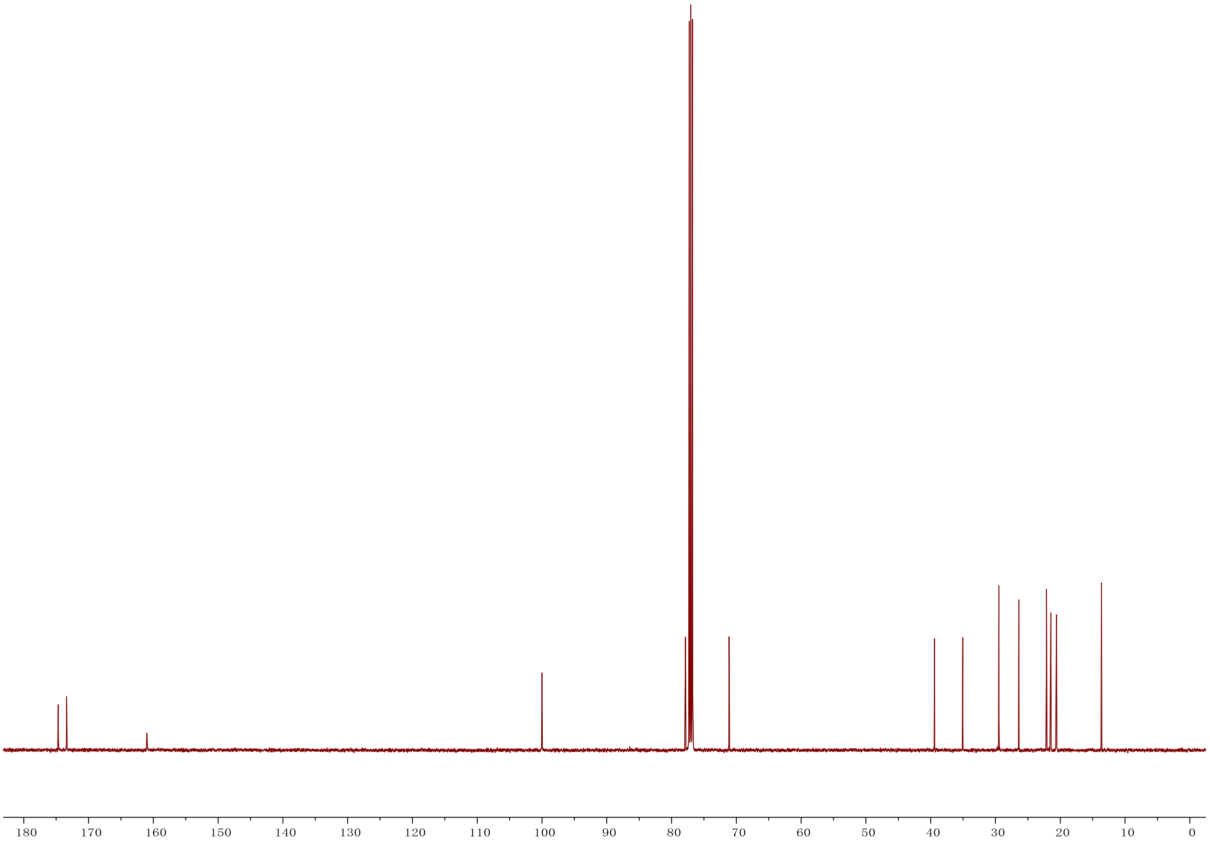

**[1]H NMR of compound 3l**

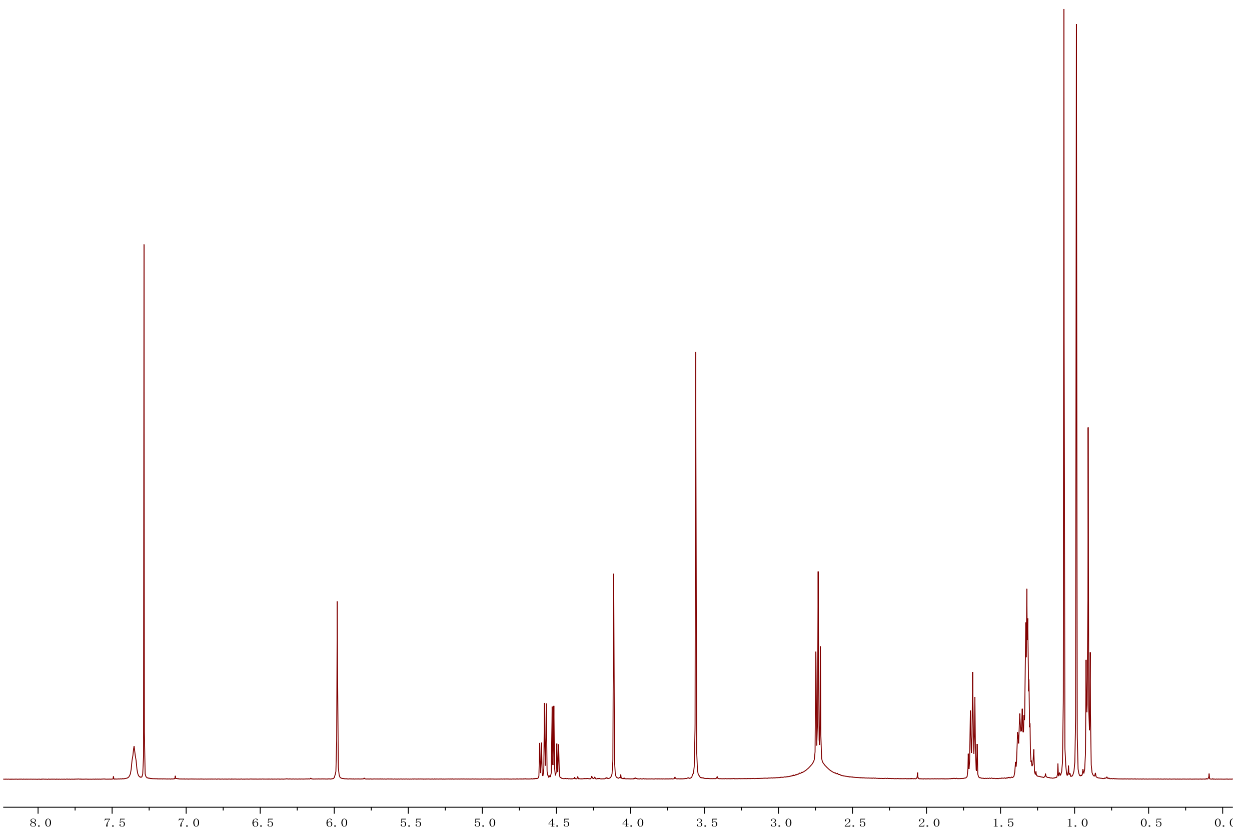


**[13]C NMR of compound 3l**

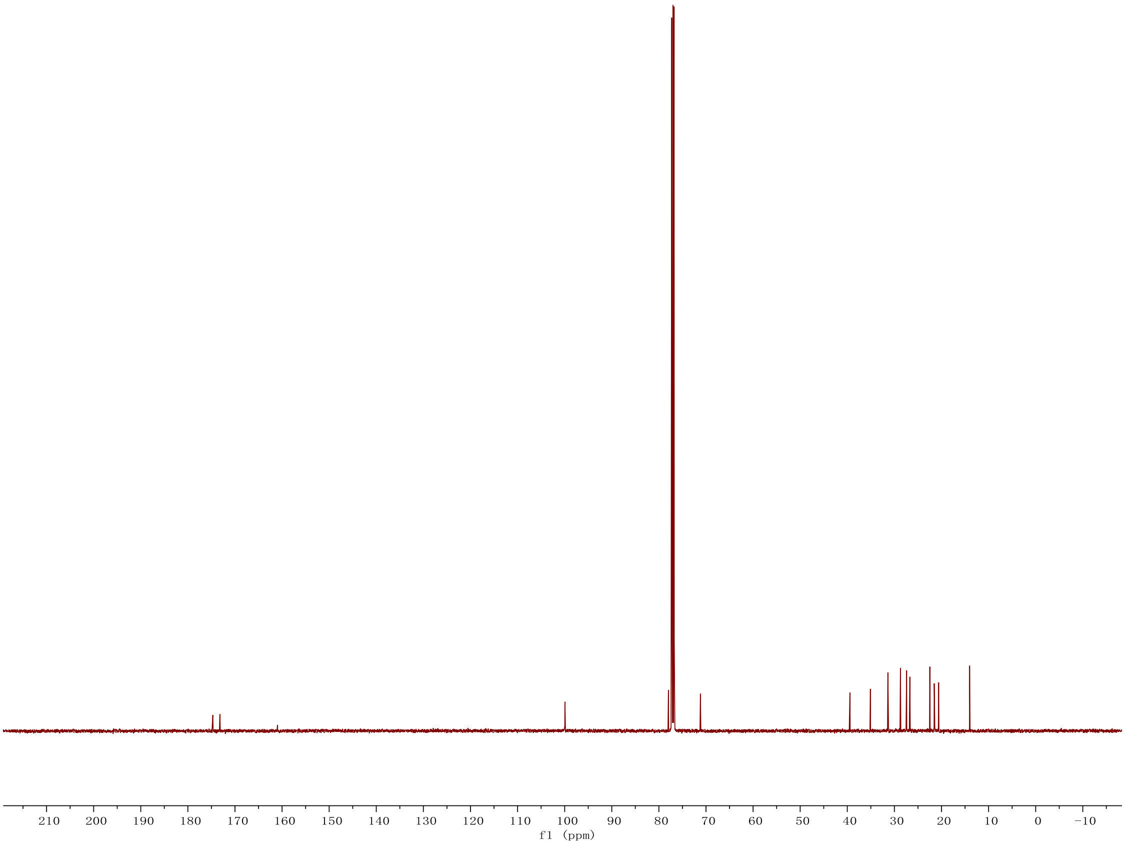

### [1]H NMR of compound 3m

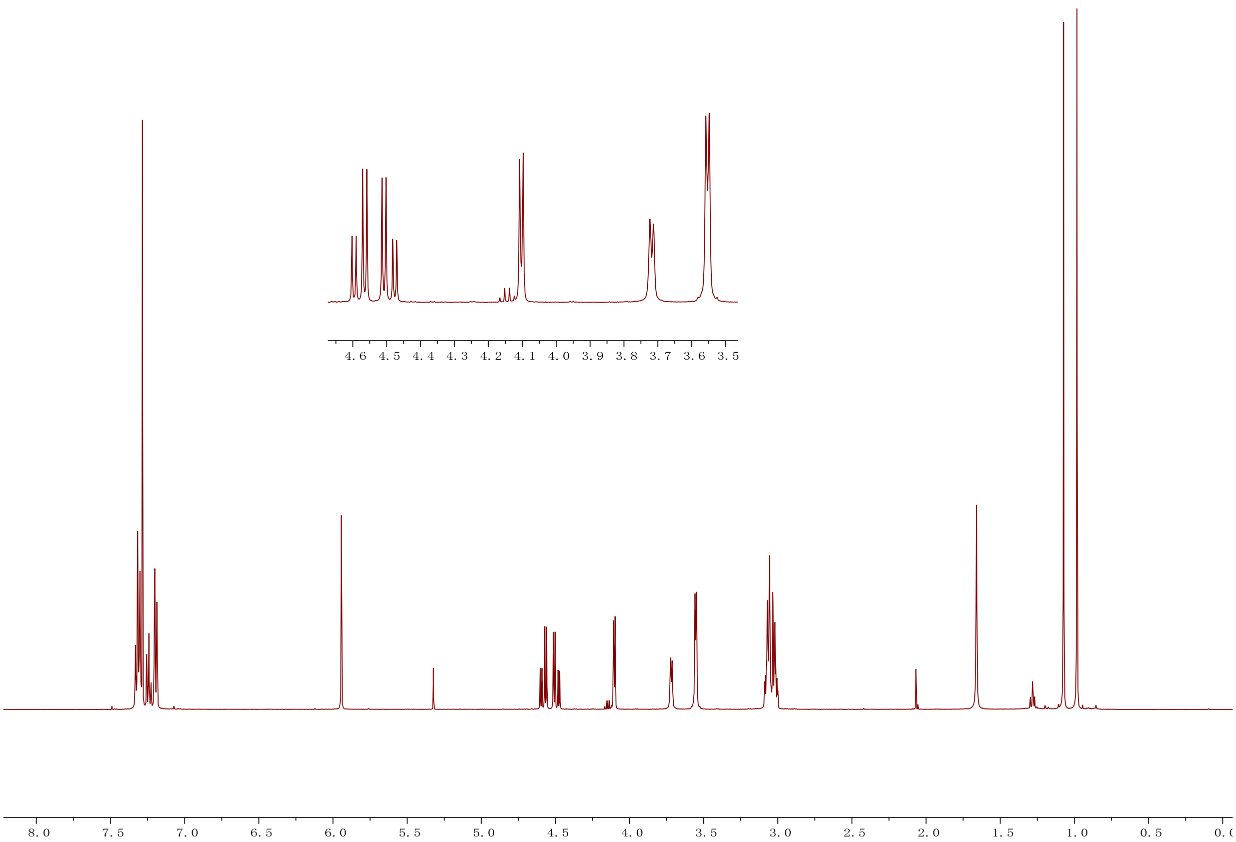


### [13]C NMR of compound 3m

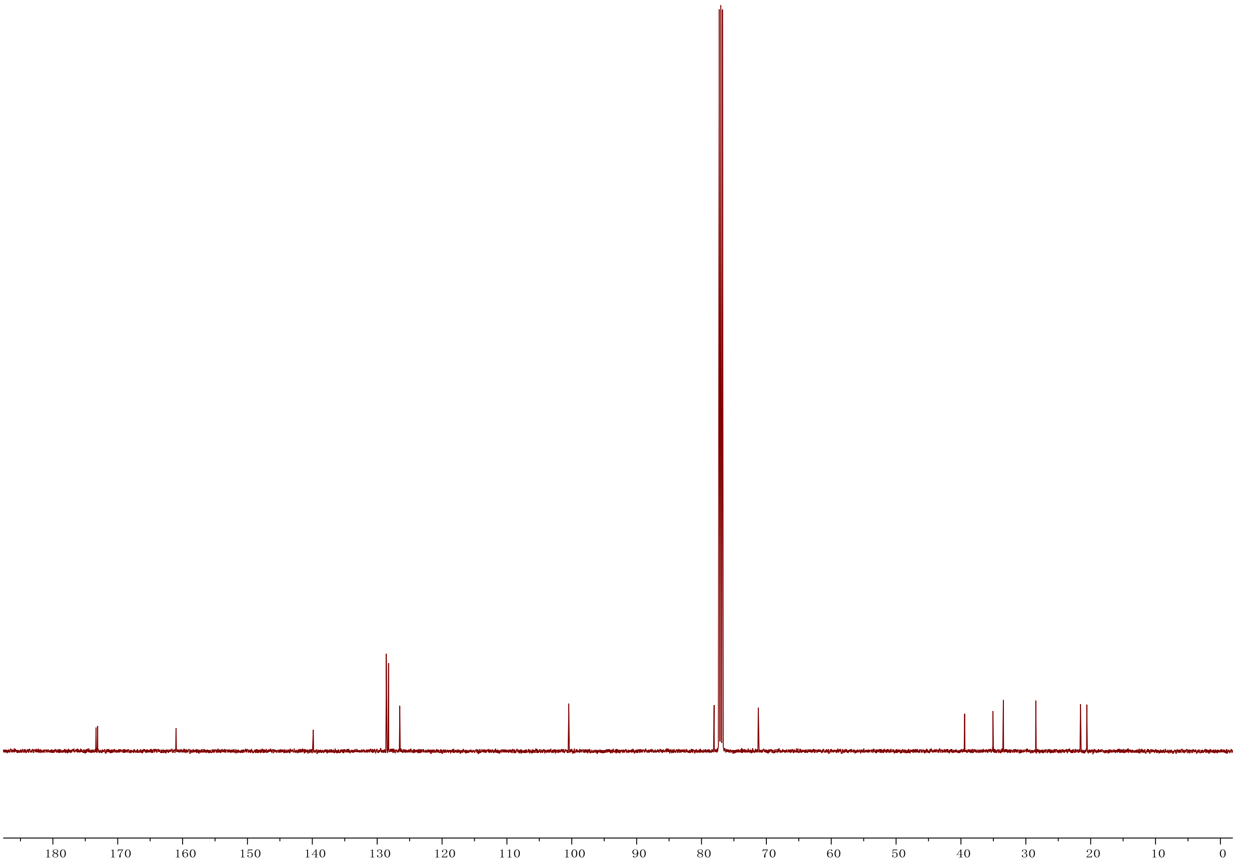

### $^{1}H$ NMR of compound 3n

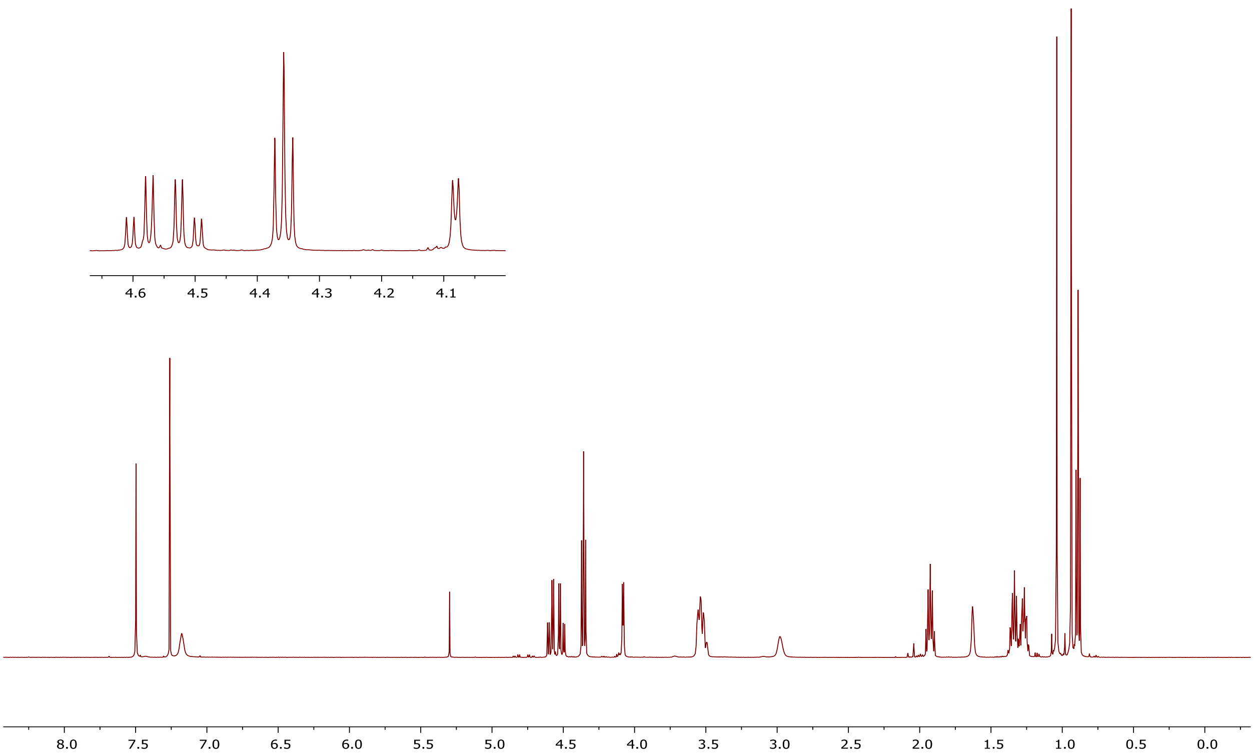


### $^{13}C$ NMR of compound 3n

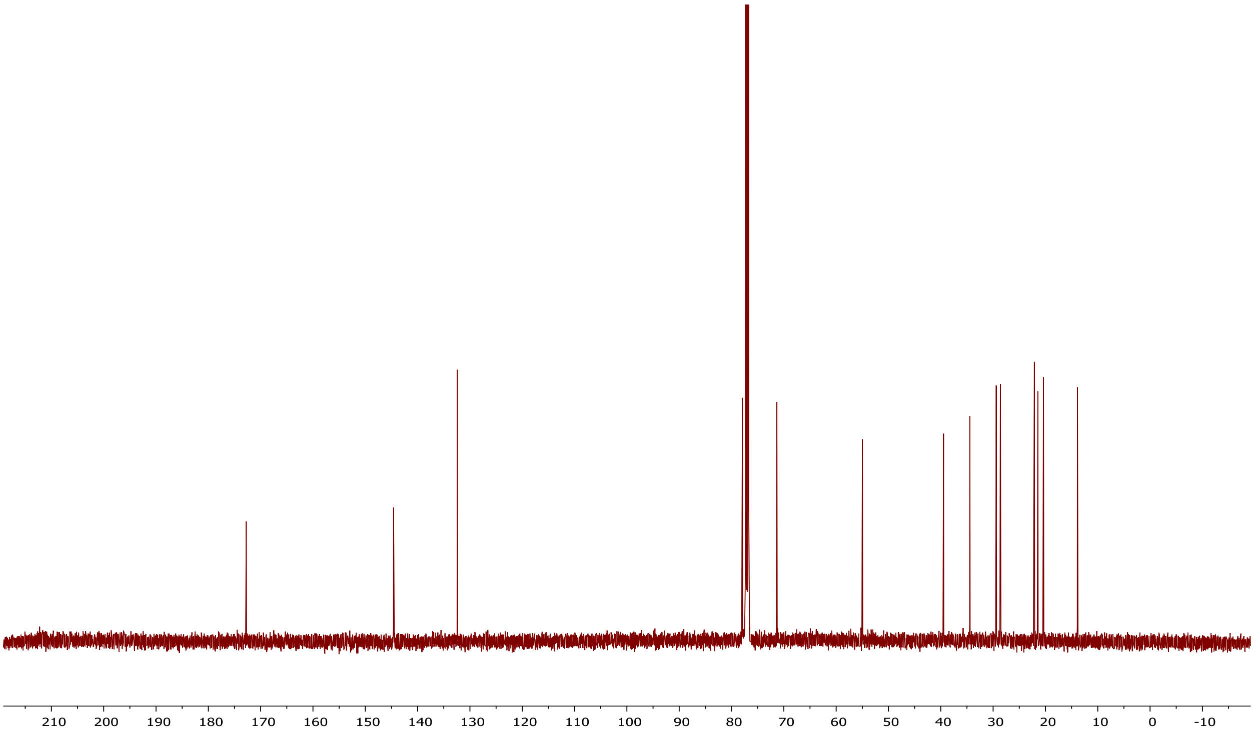

### [1]H NMR of compound 3o

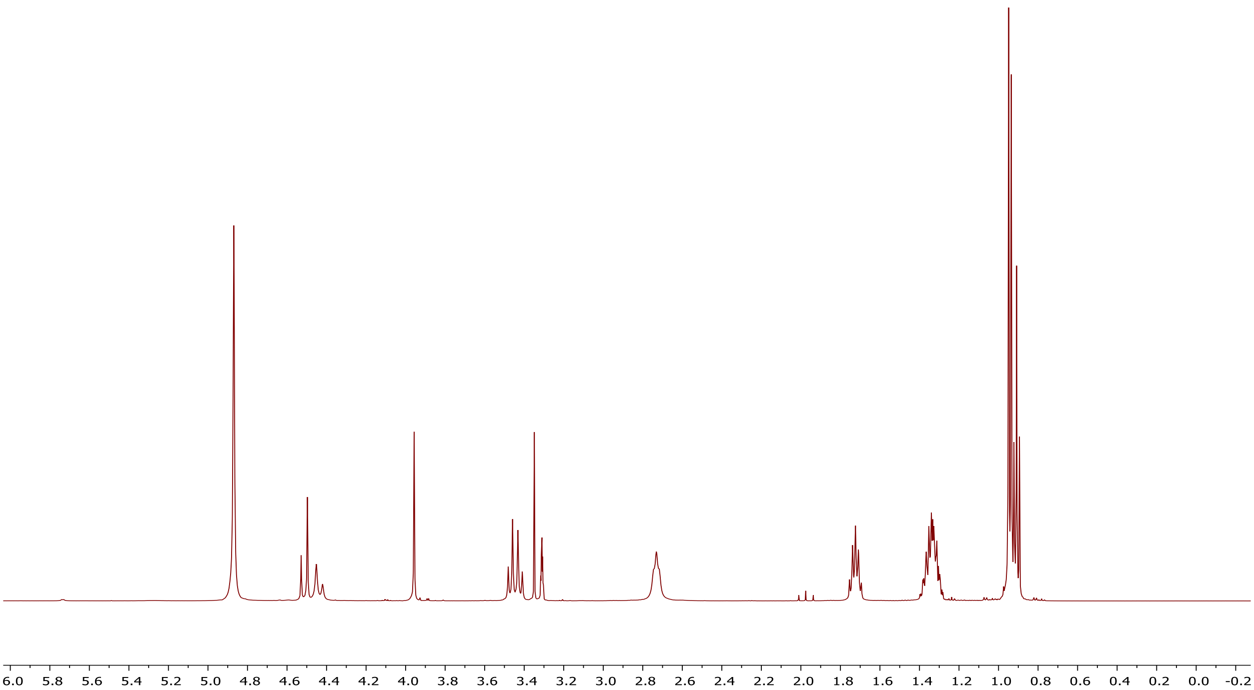


### [13]C NMR of compound 3o

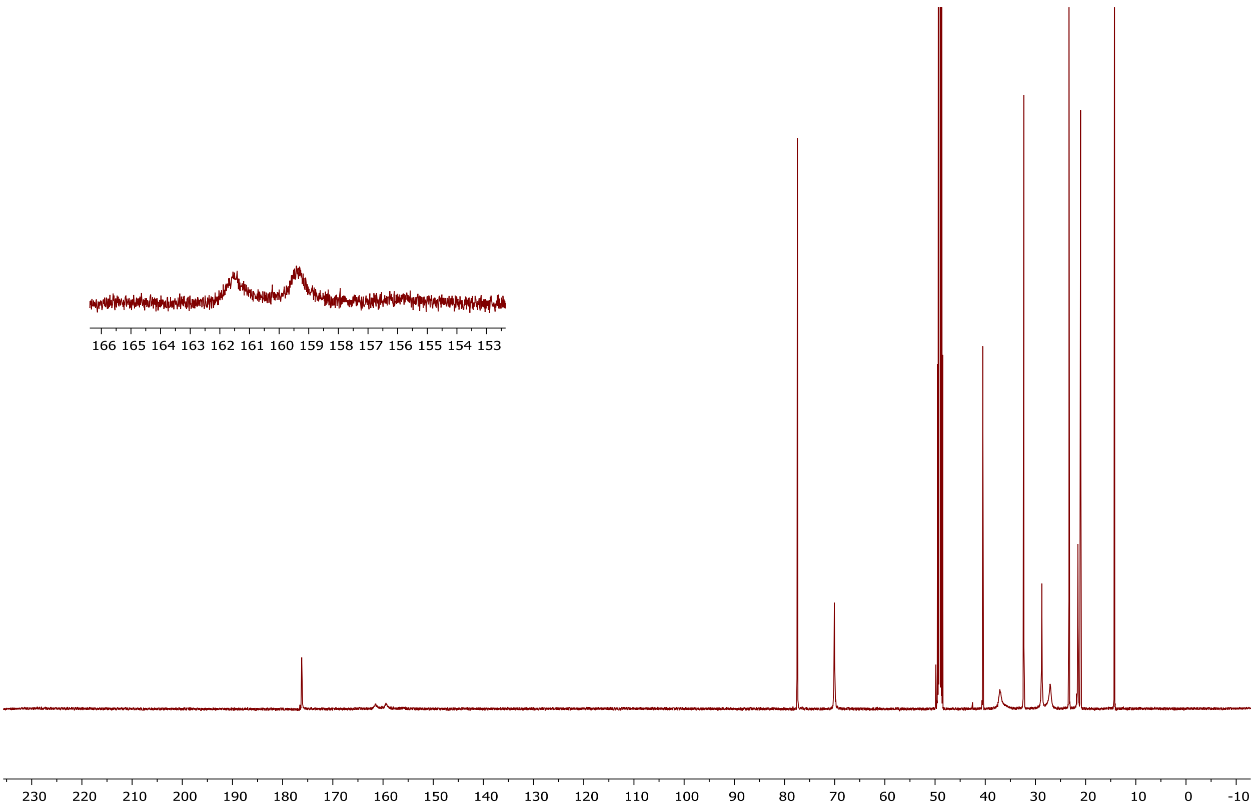

### $^{1}H$ NMR of compound 3p

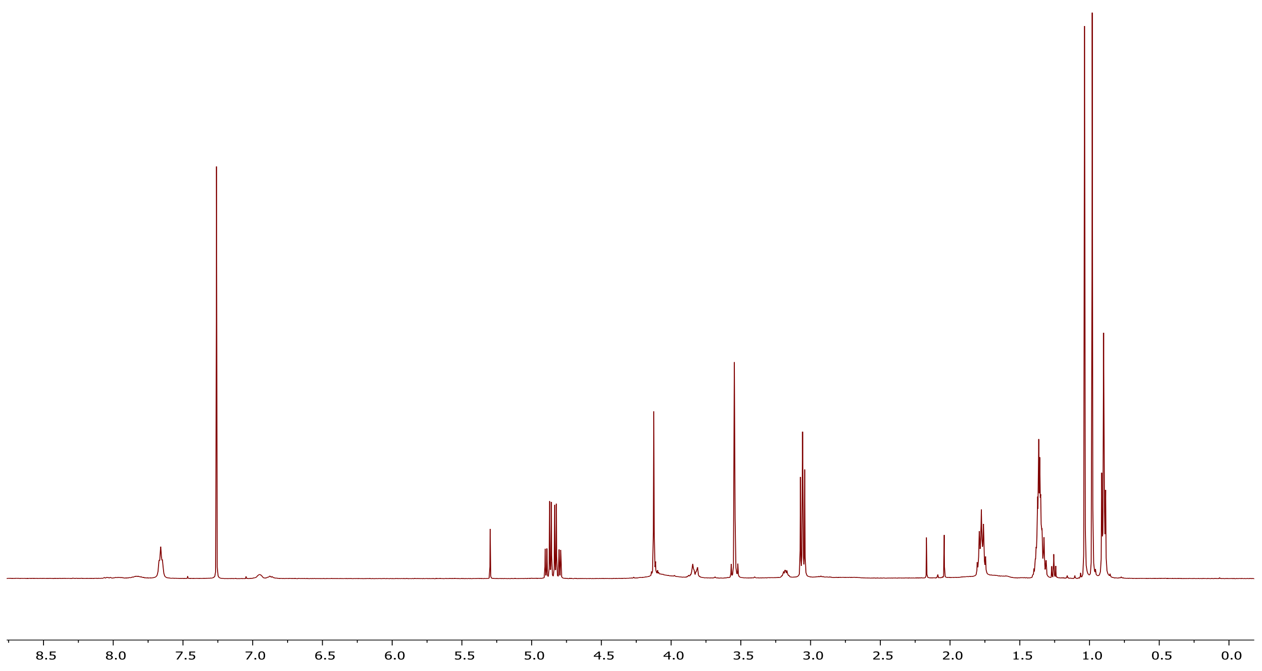


### $^{13}C$ NMR of compound 3p

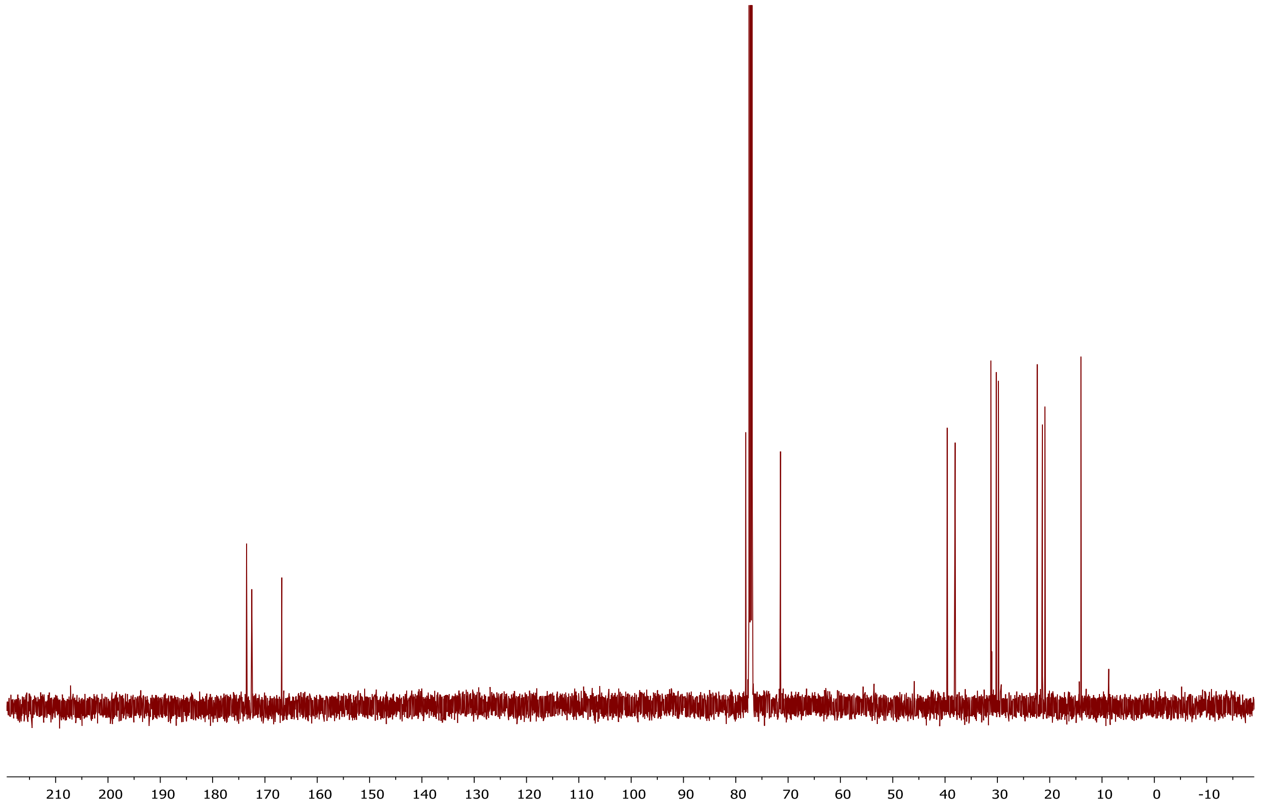

### $^{1}H$ NMR of compound 3q

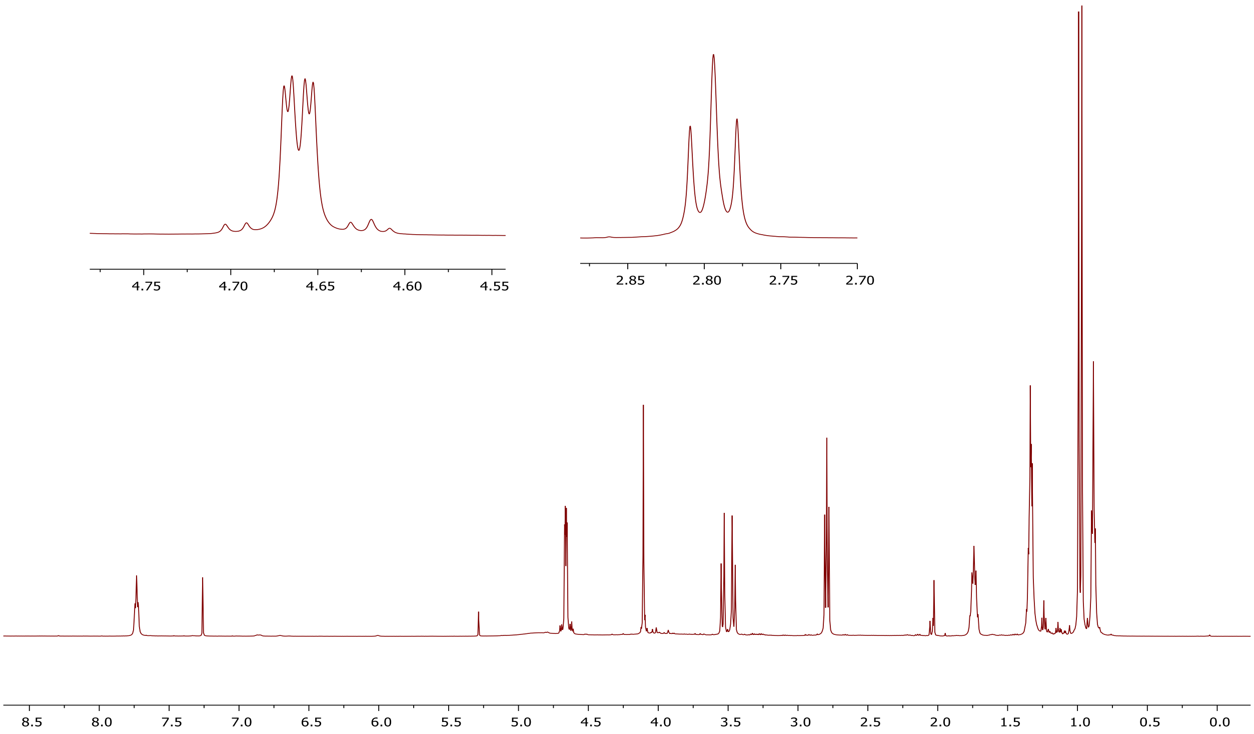


### $^{13}C$ NMR of compound 3q

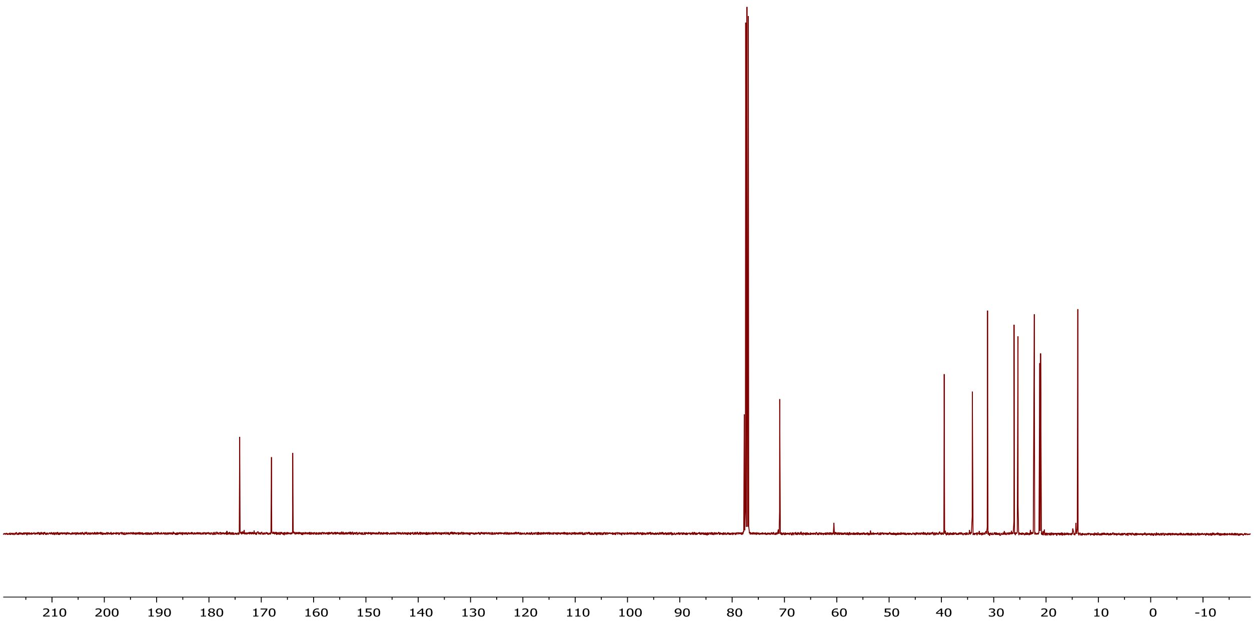

### [1]H NMR of compound 3r

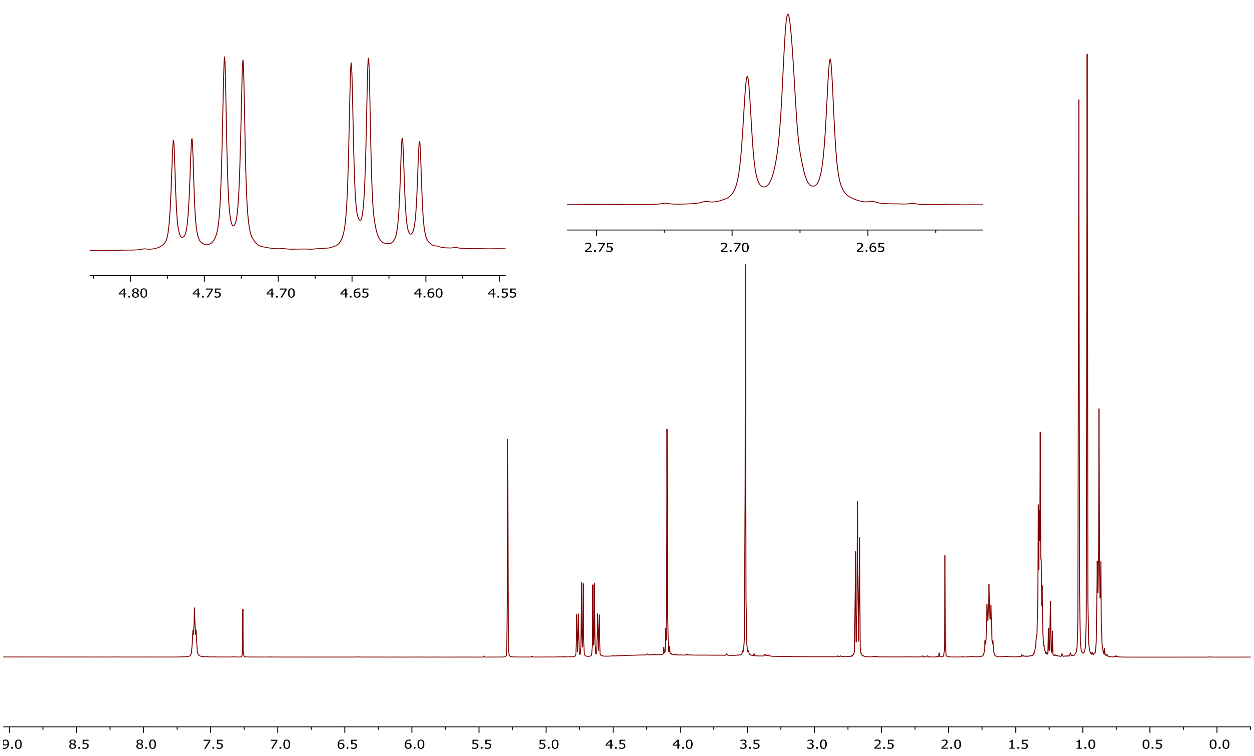


### [13]C NMR of compound 3r

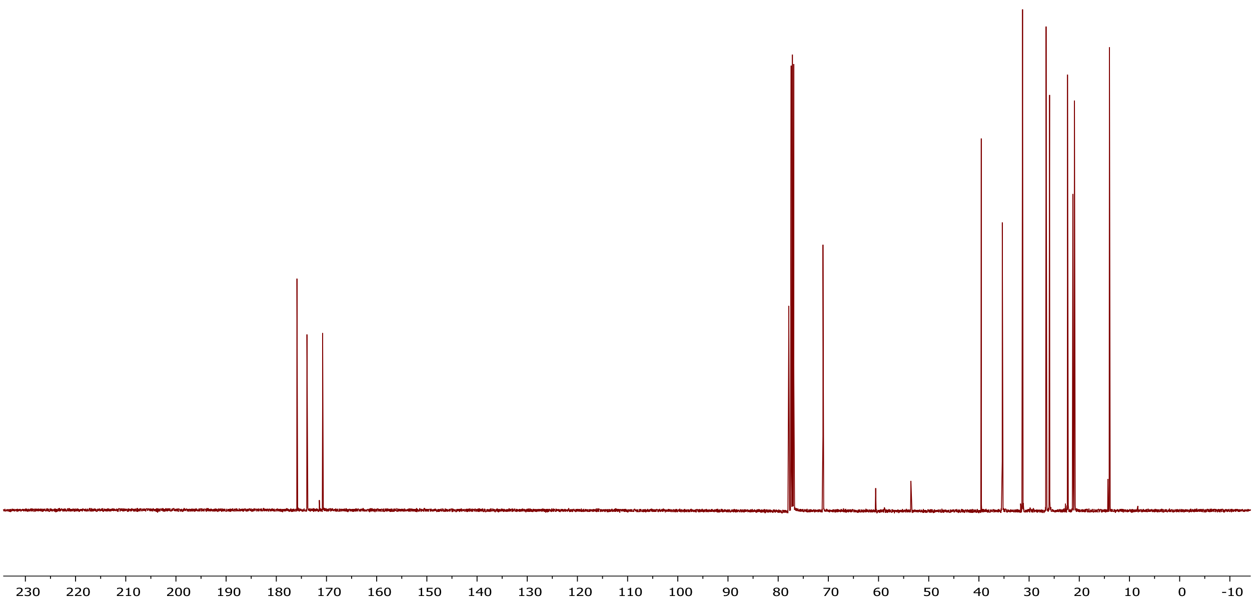

### $^{1}H$ NMR of compound 3s

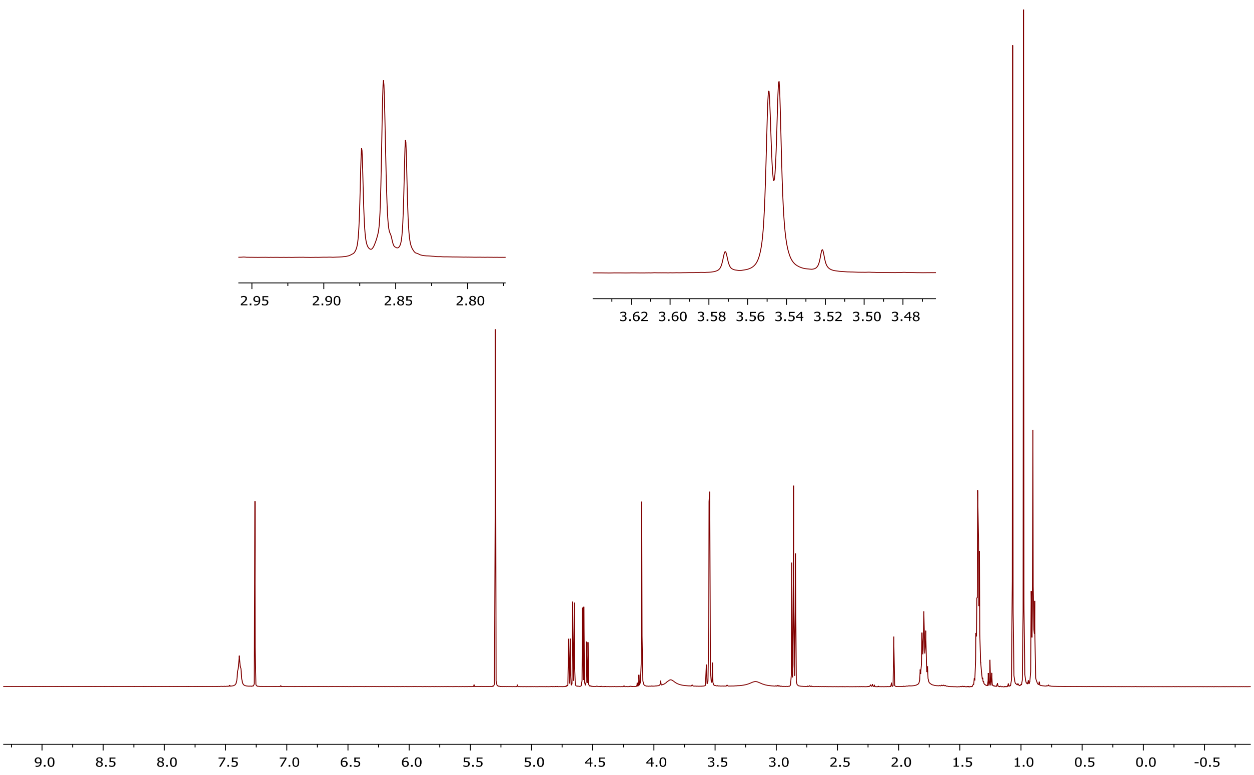


### $^{13}C$ NMR of compound 3s

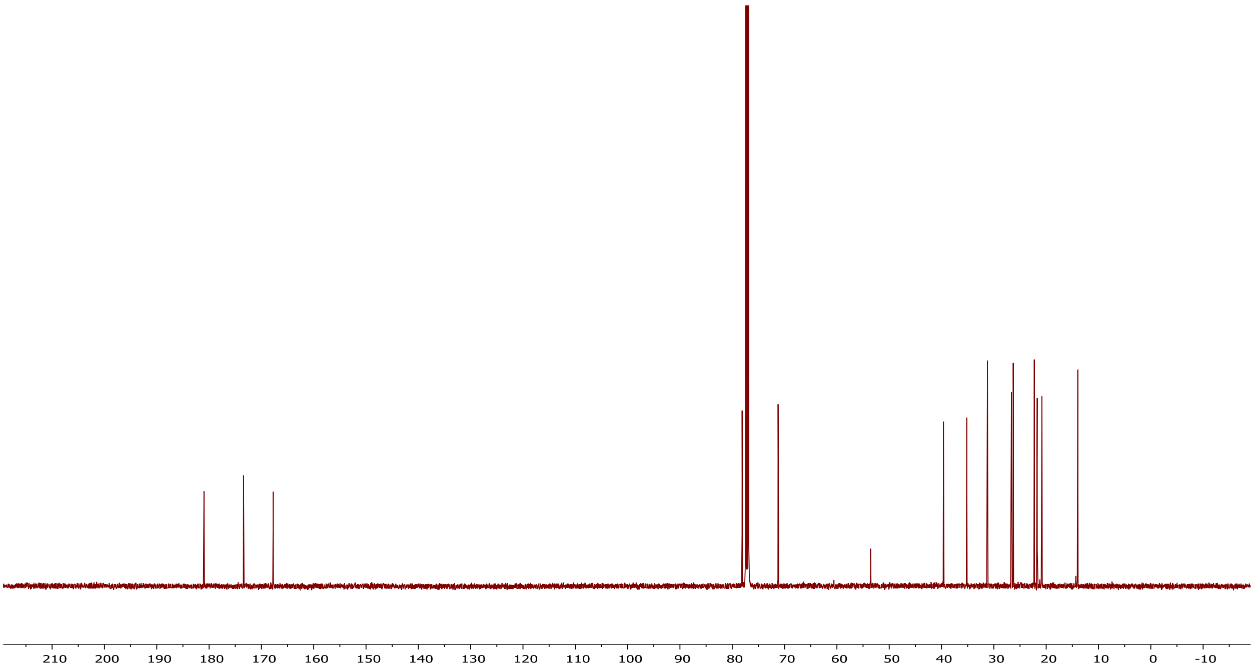

### $^{1}H$ NMR of compound 3t

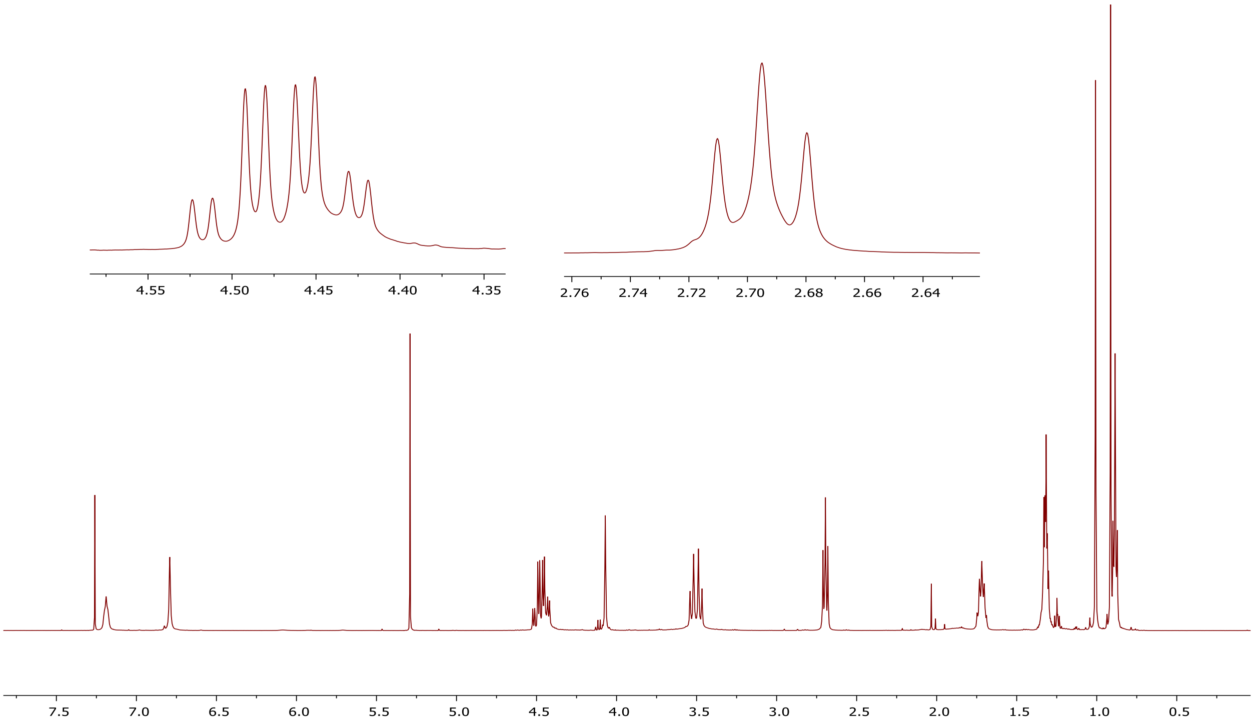


### $^{13}C$ NMR of compound 3t

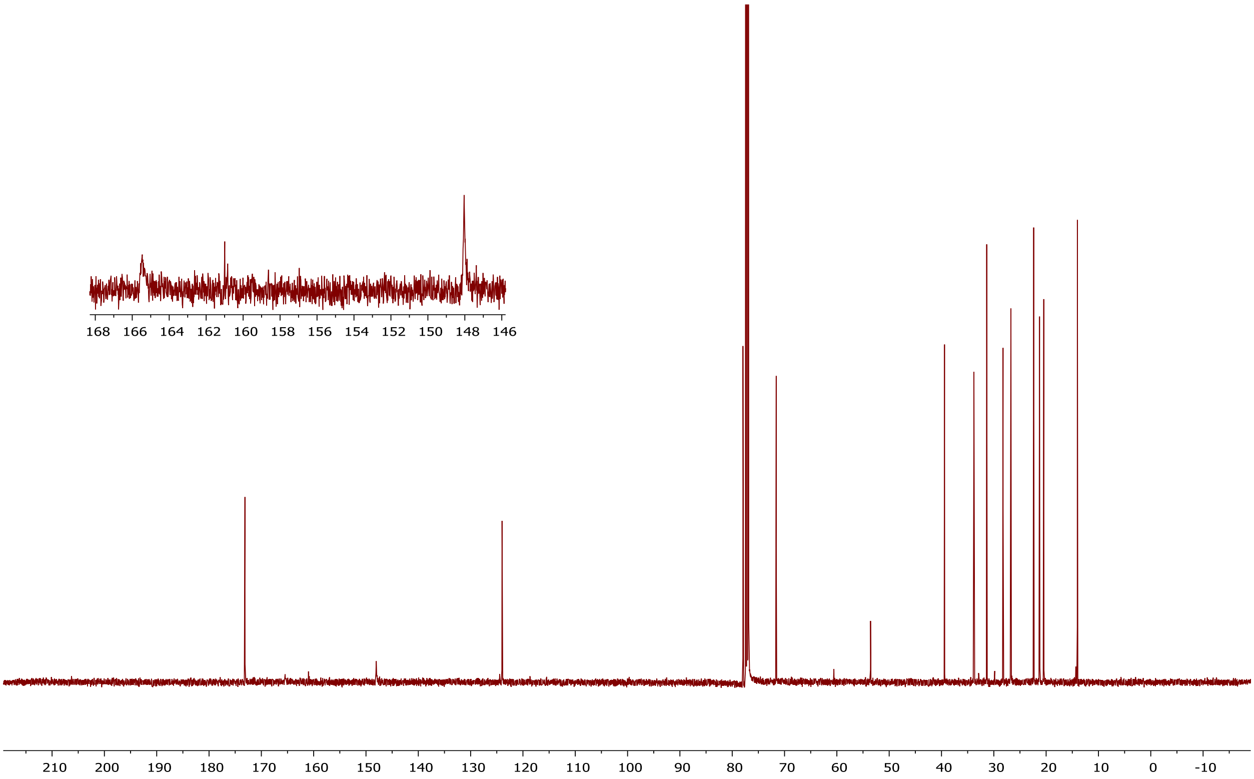

### [1]H NMR of compound 3u

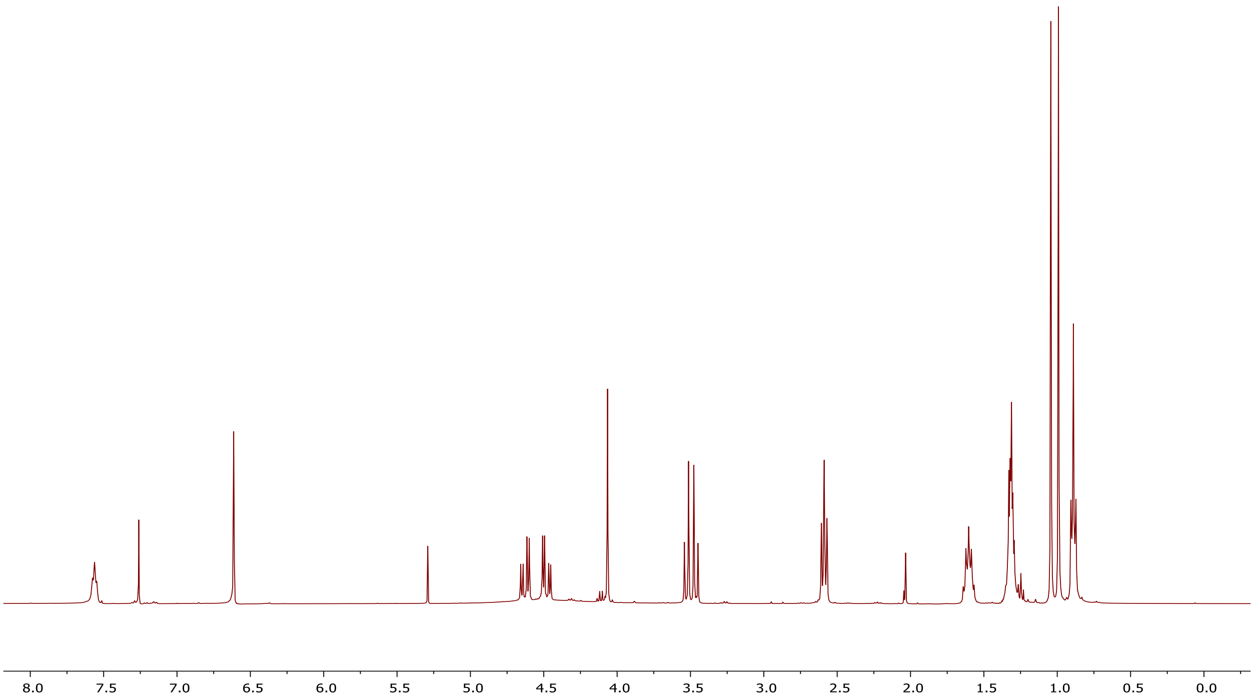


### [13]C NMR of compound 3u

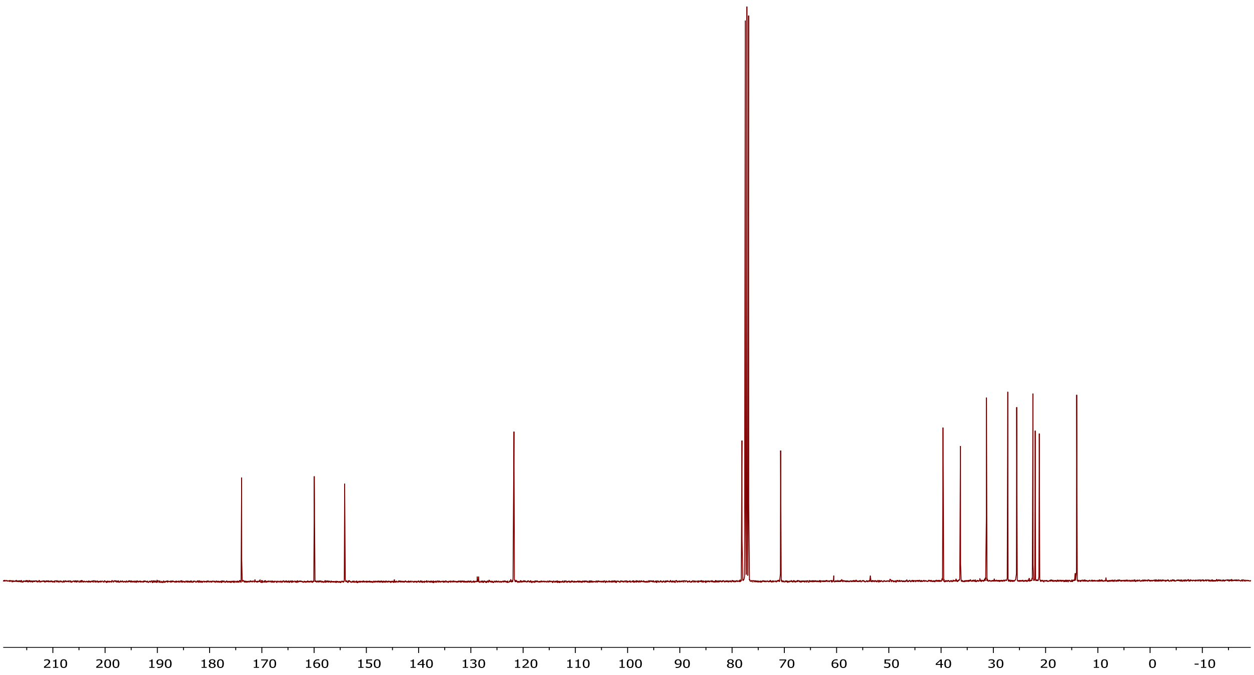

### $^{1}$H NMR of compound 3v

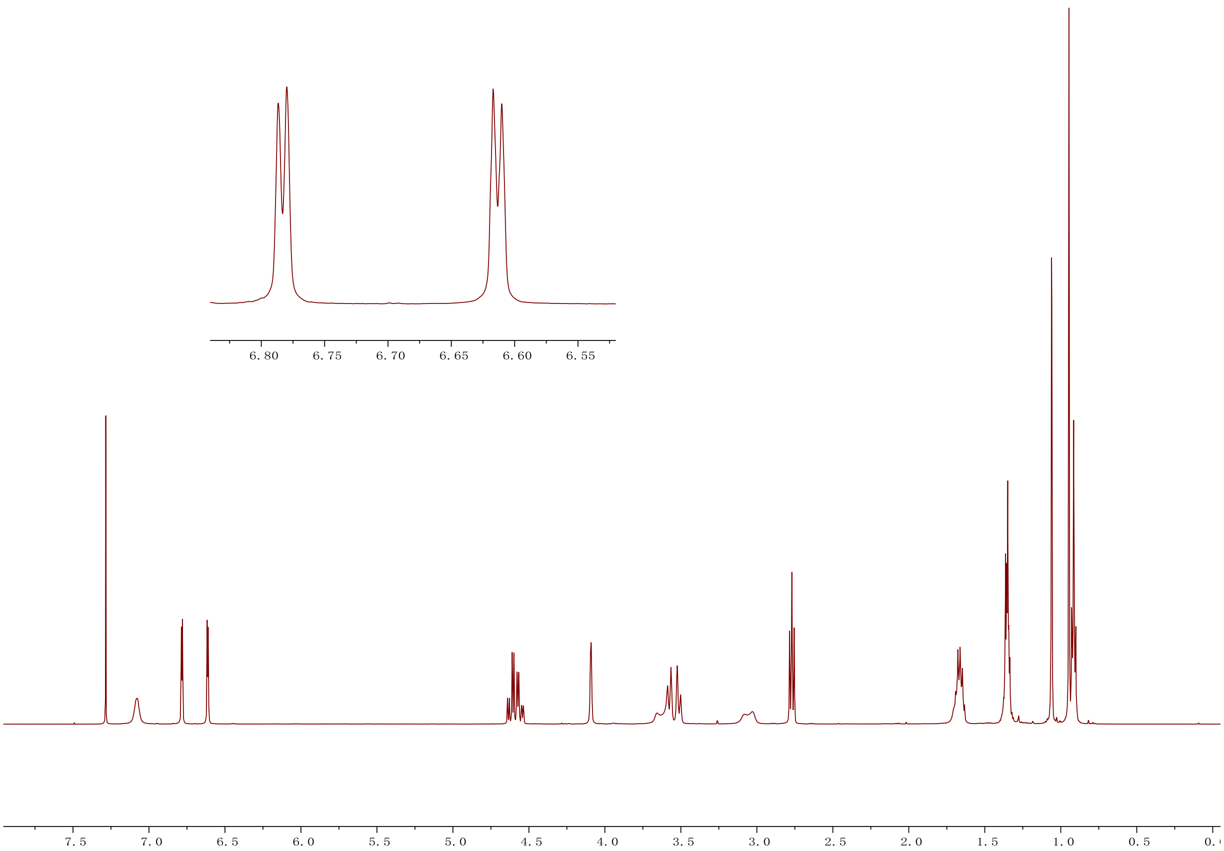


### $^{13}$C NMR of compound 3v

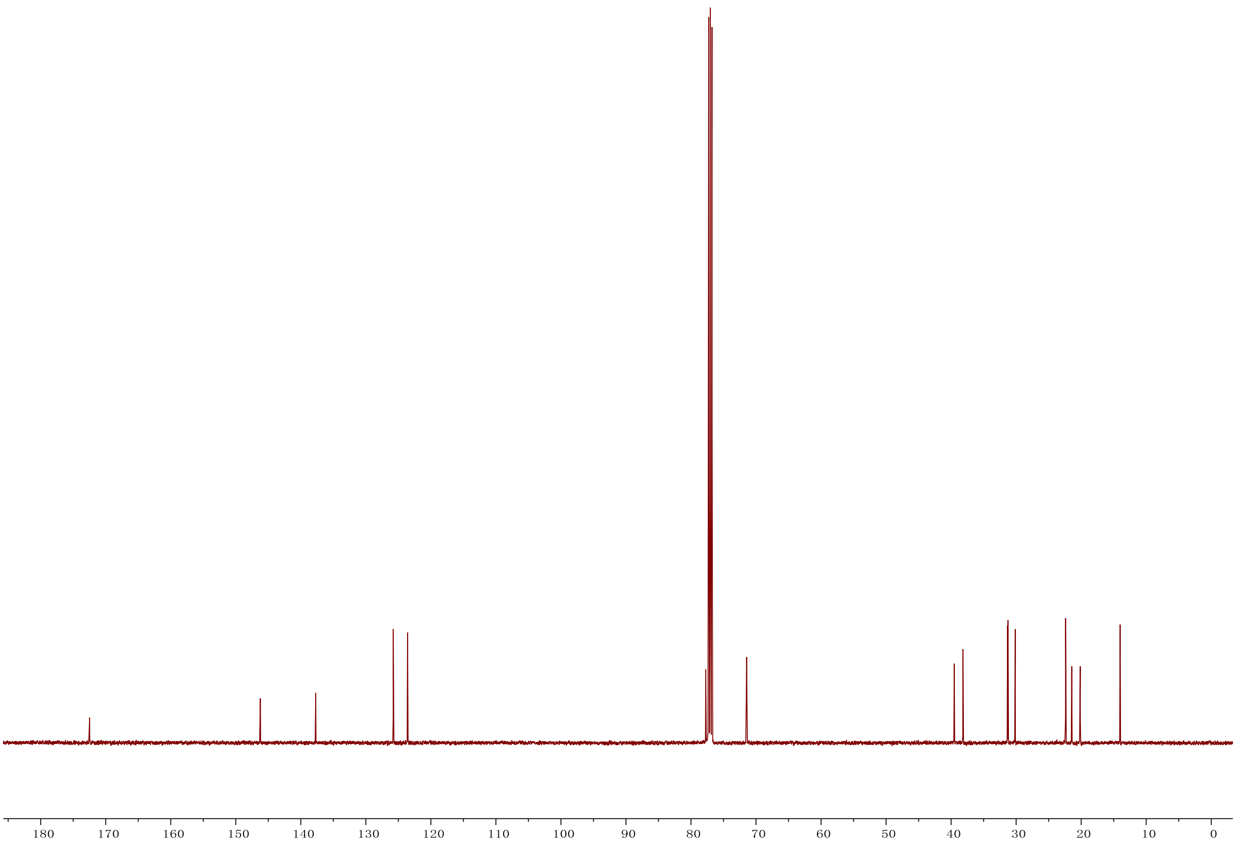

**[1]H NMR of compound 4**

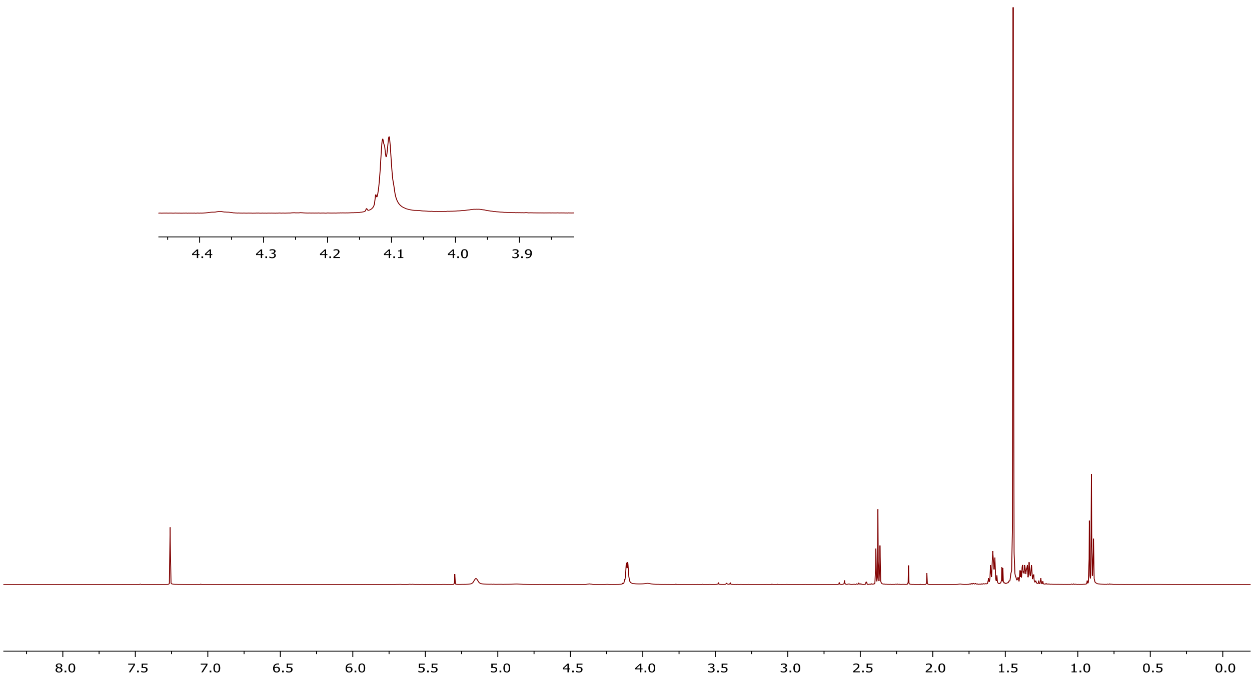


**[13]C NMR of compound 4**

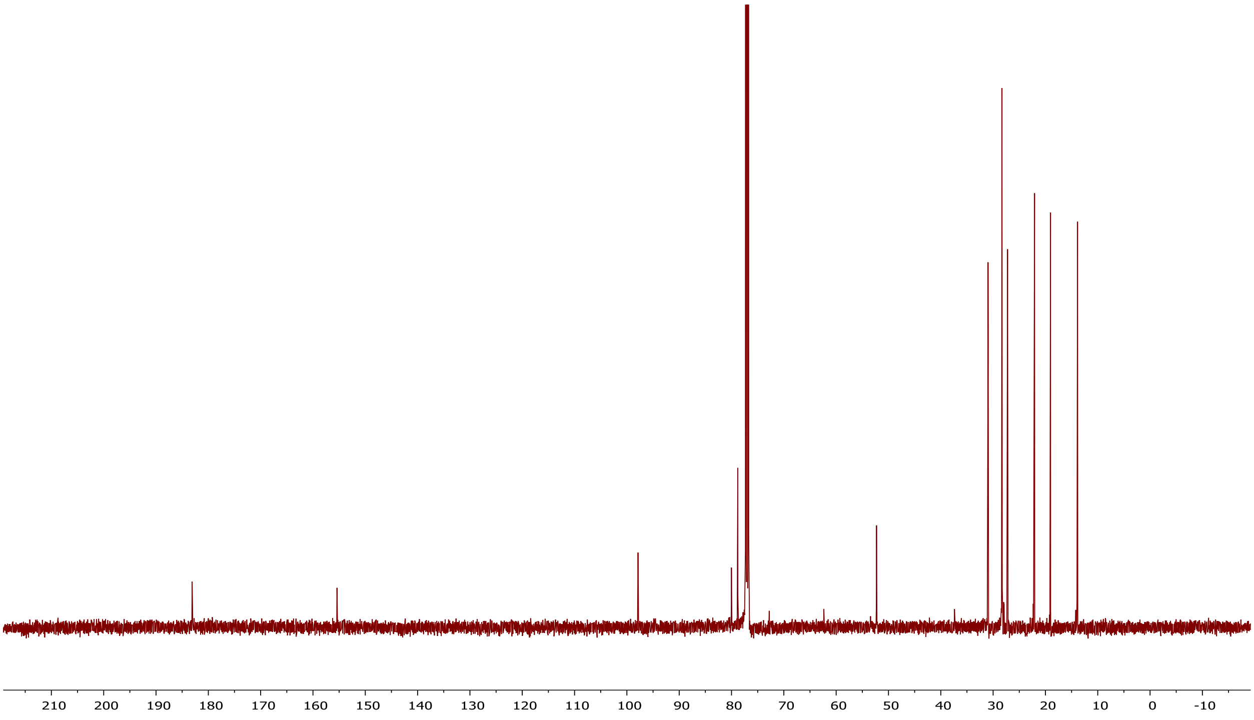

### [1]H NMR of compound 5

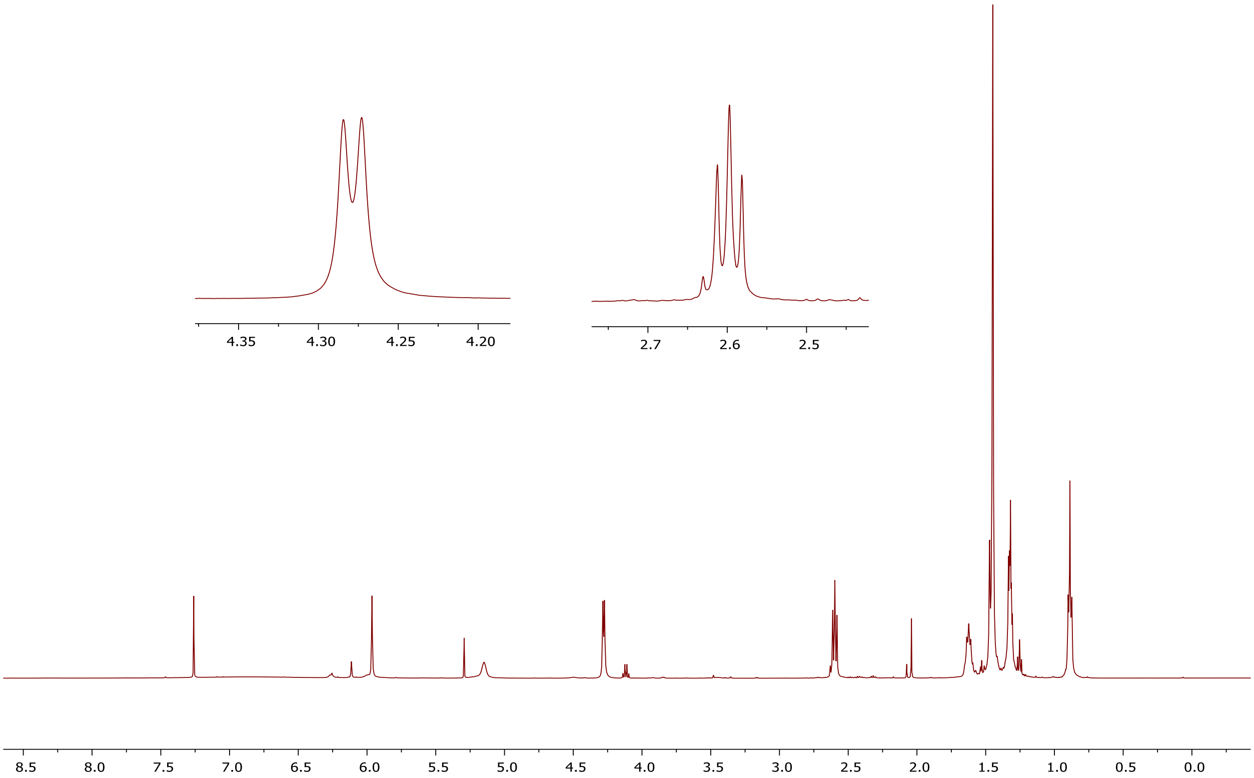


### [13]C NMR of compound 5

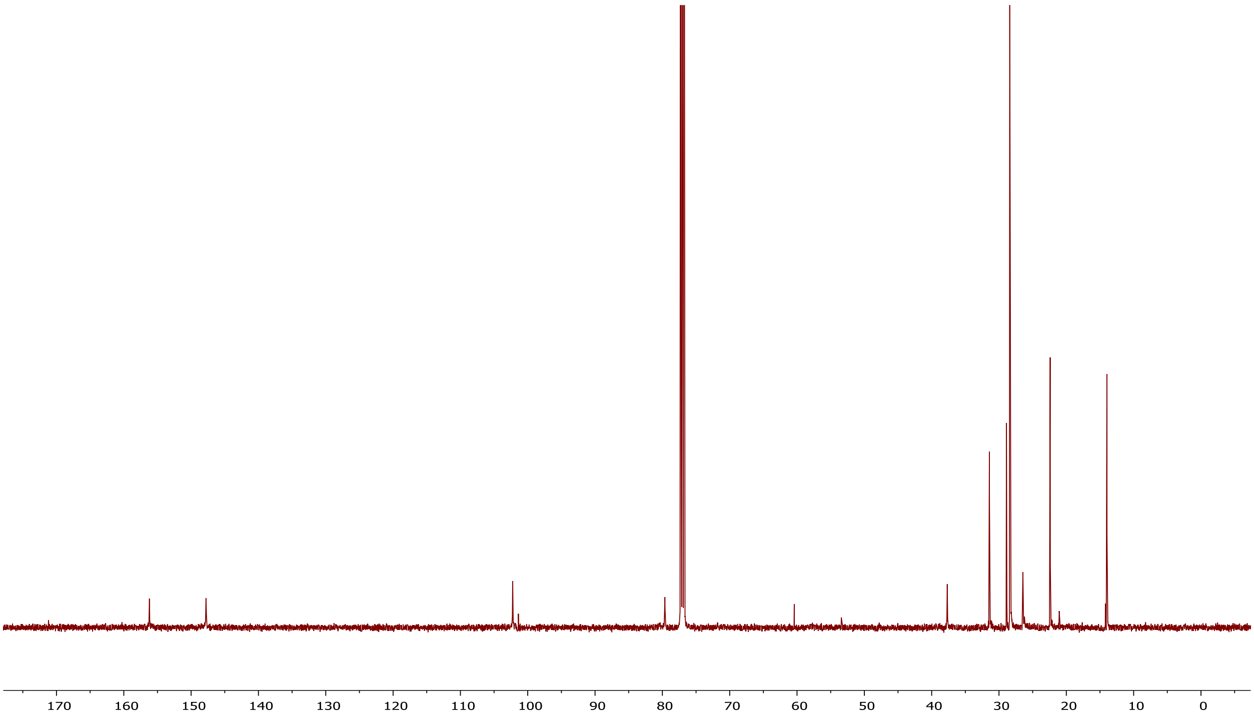

### [1]H NMR of compound 6b

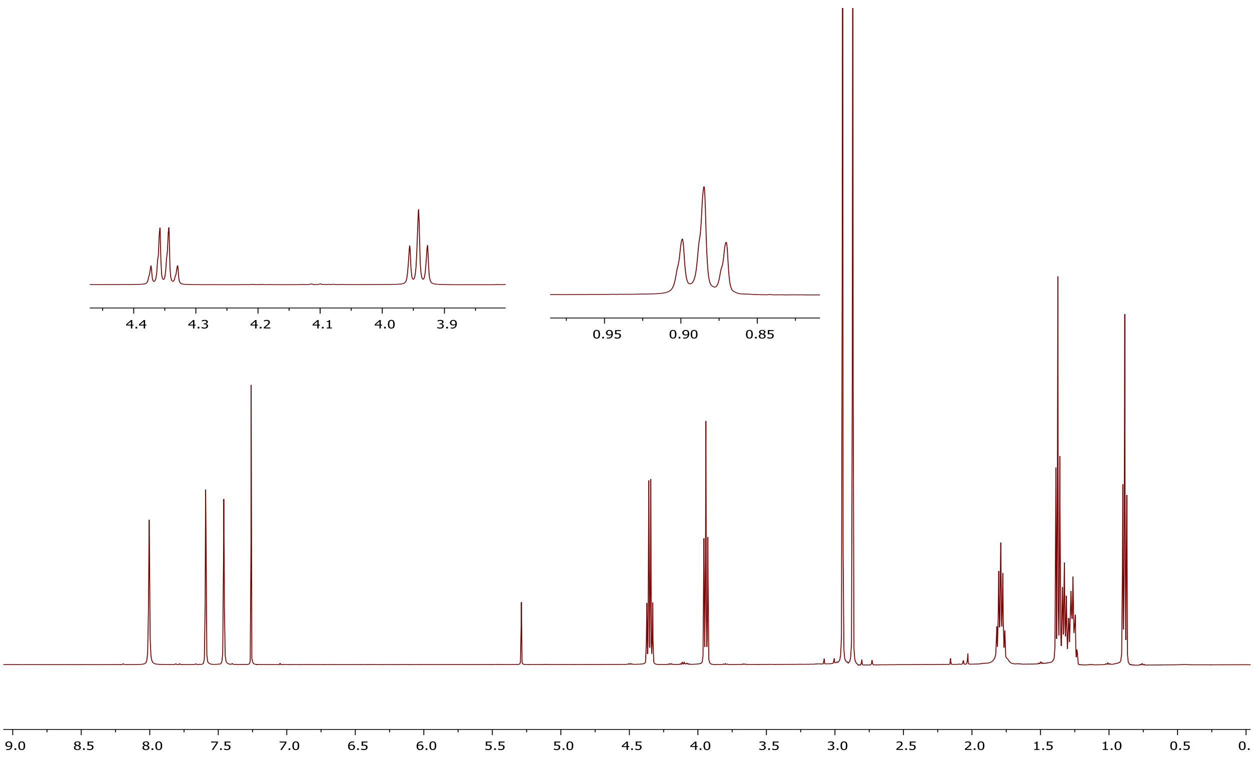


### [13]C NMR of compound 6b

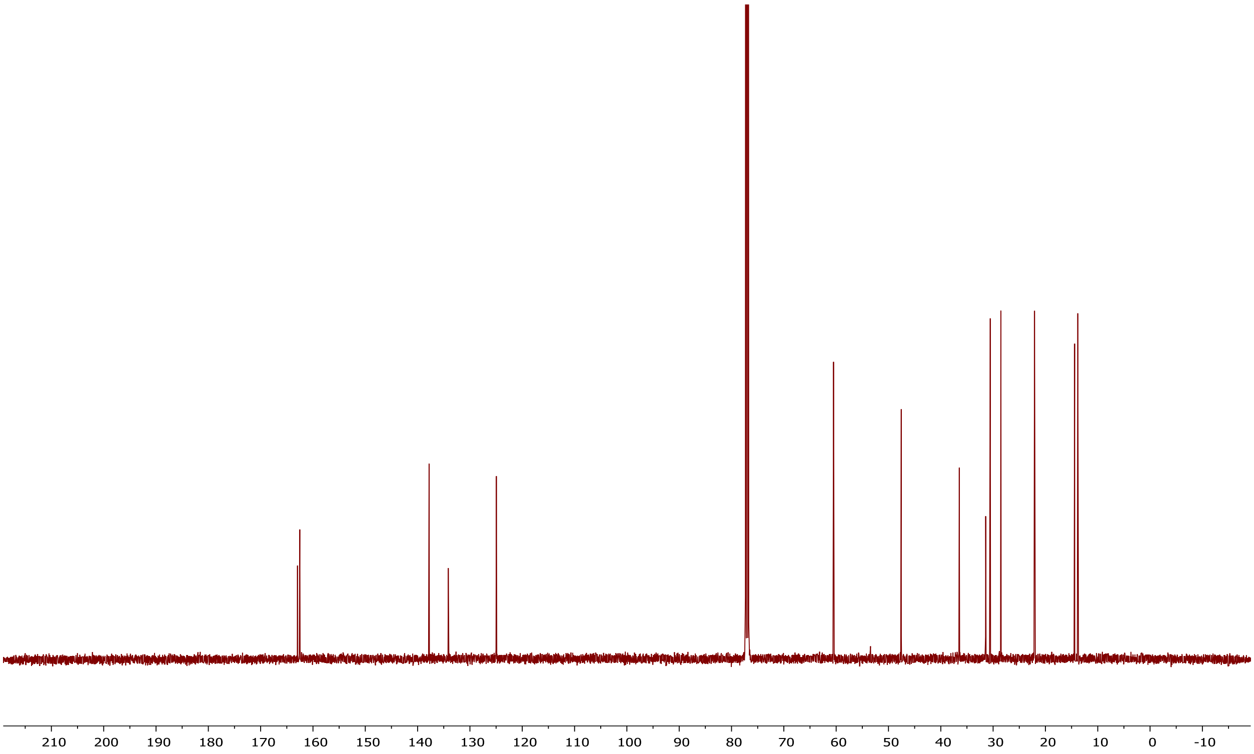

[1]H NMR of compound 6c

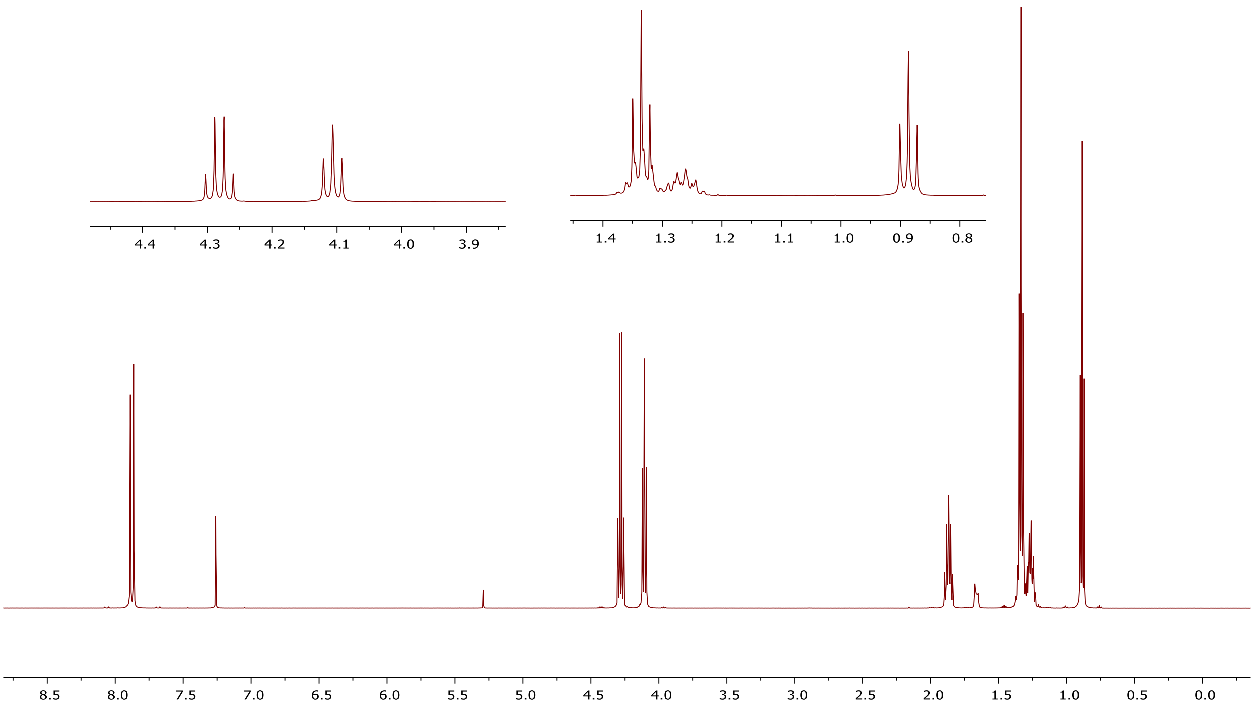


[13]C NMR of compound 6c

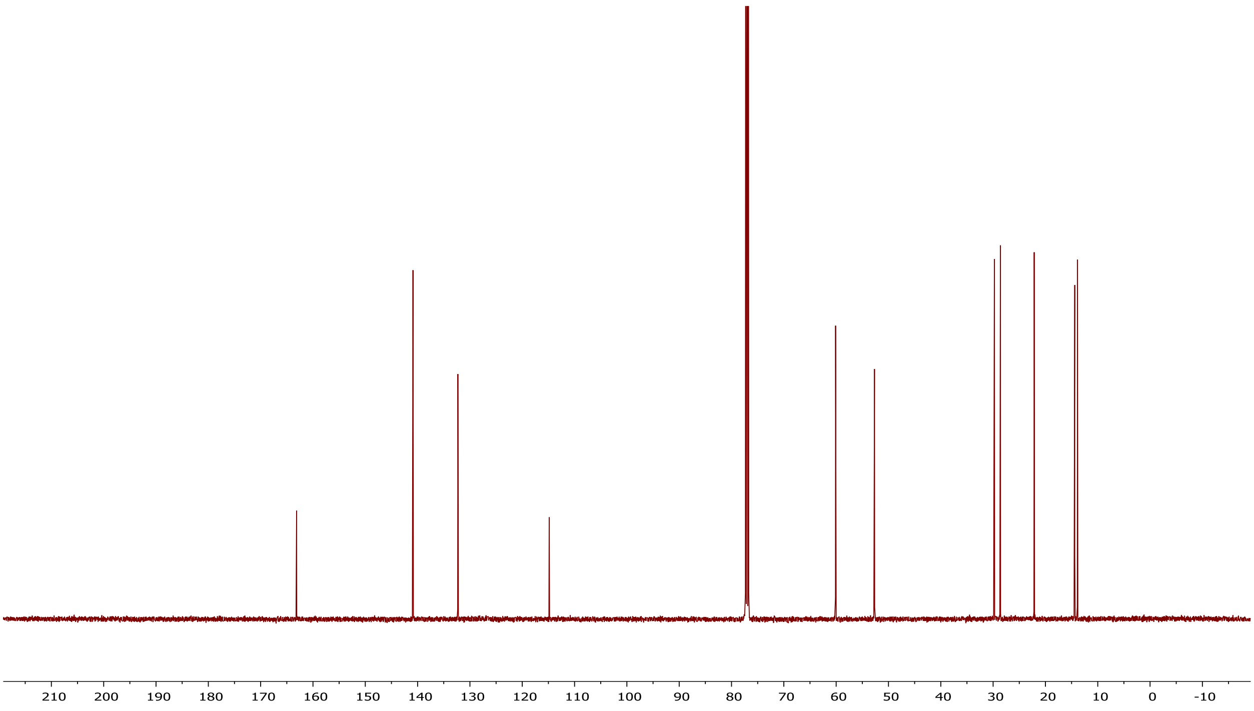

$^{1}H$ NMR of compound 6d

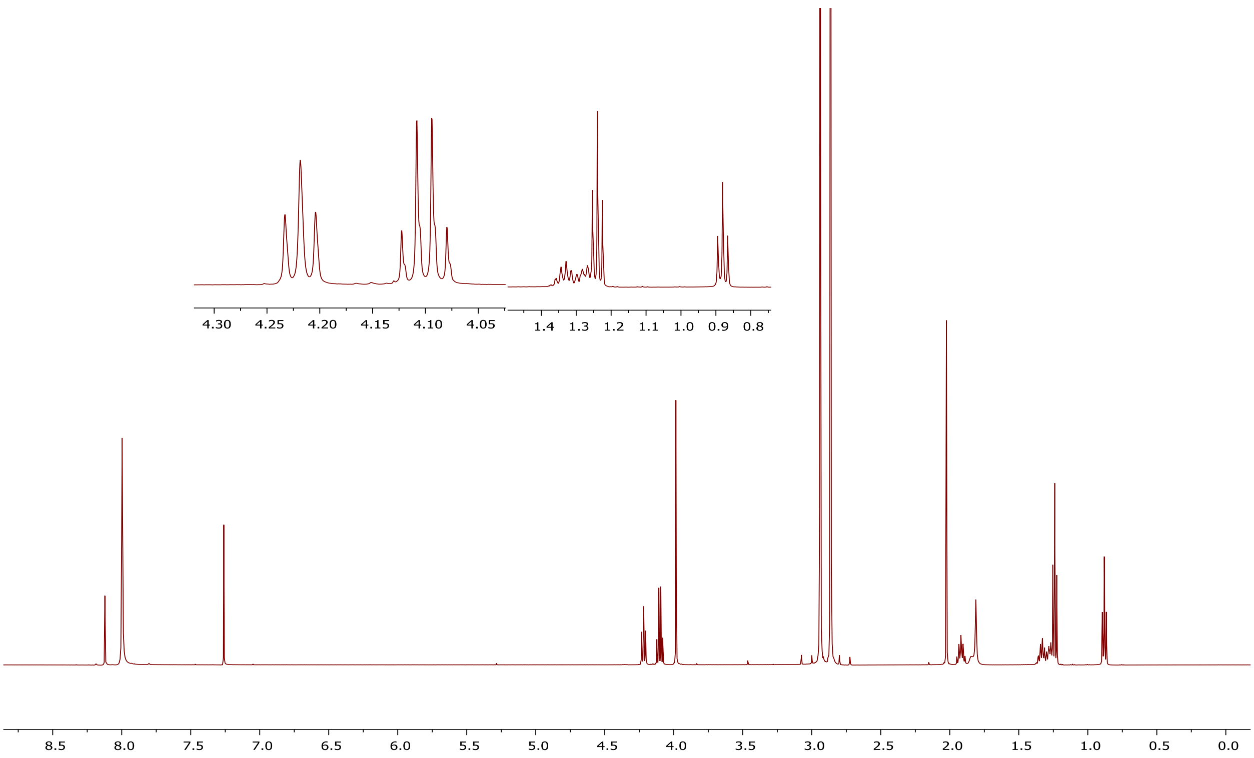


$^{13}C$ NMR of compound 6d

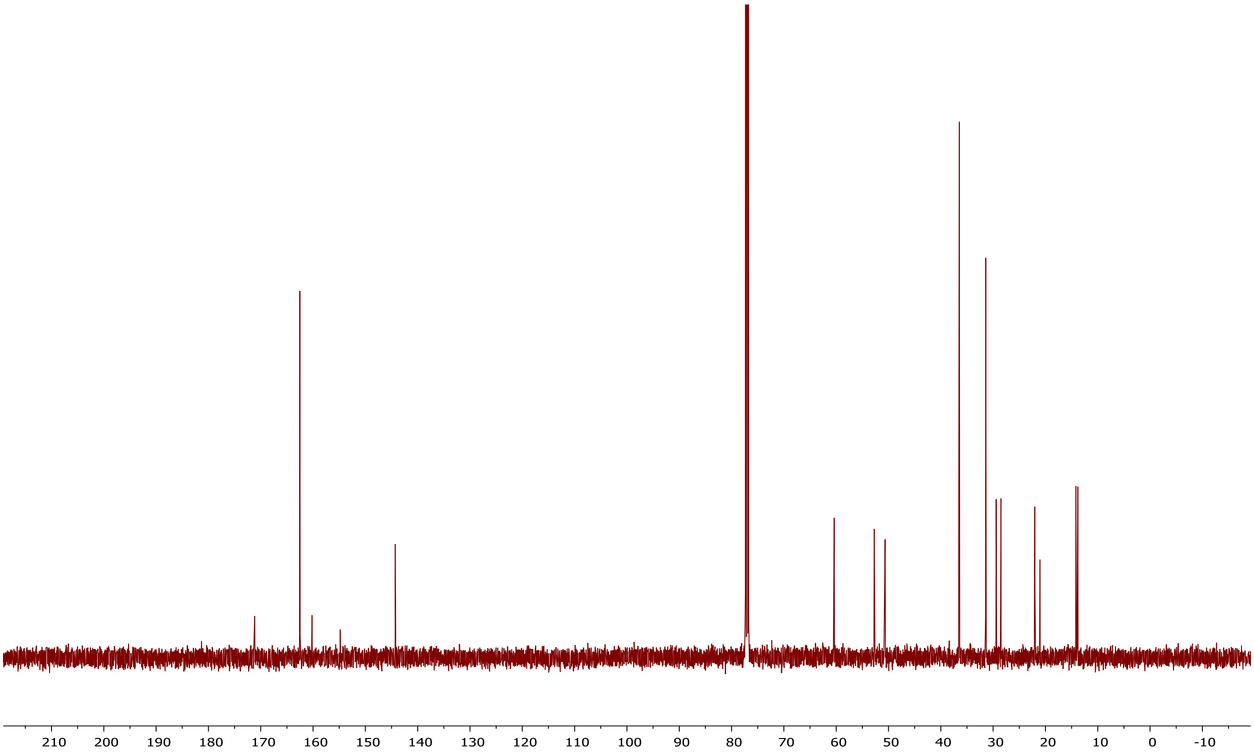

### [1]H NMR of compound 12e

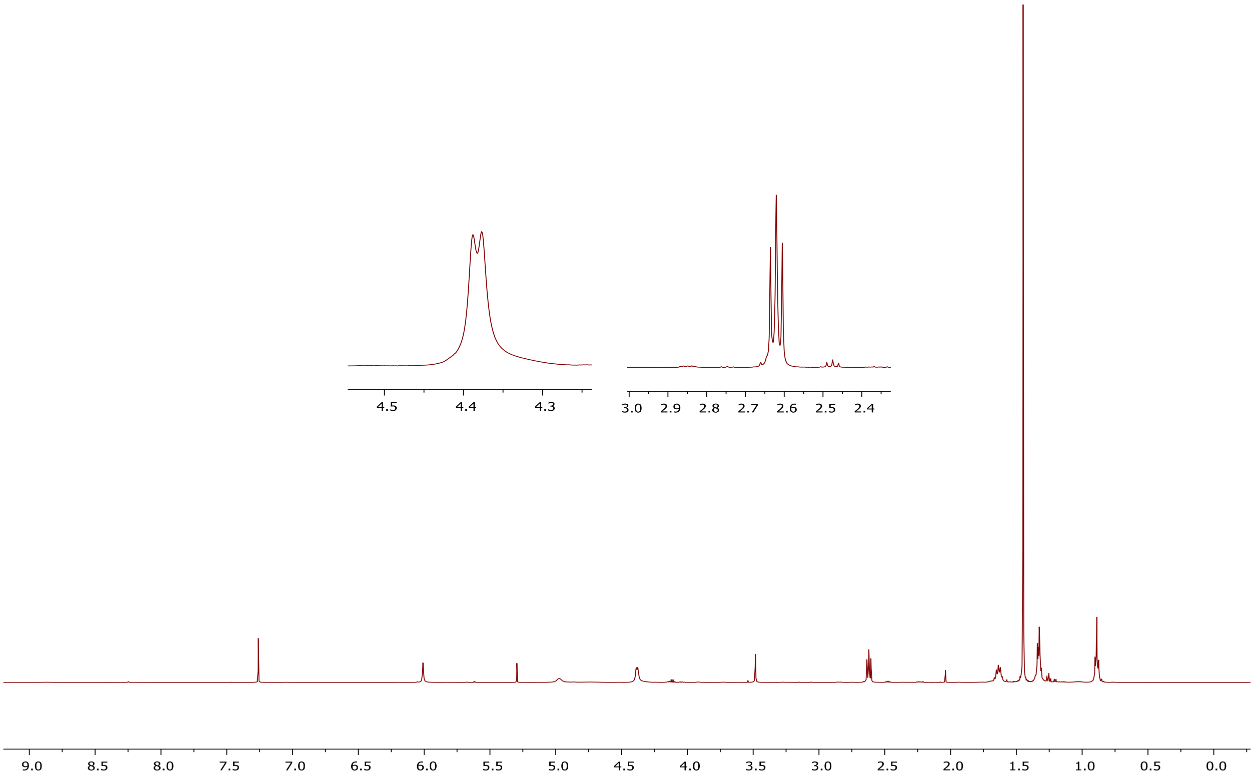


### [13]C NMR of compound 12e

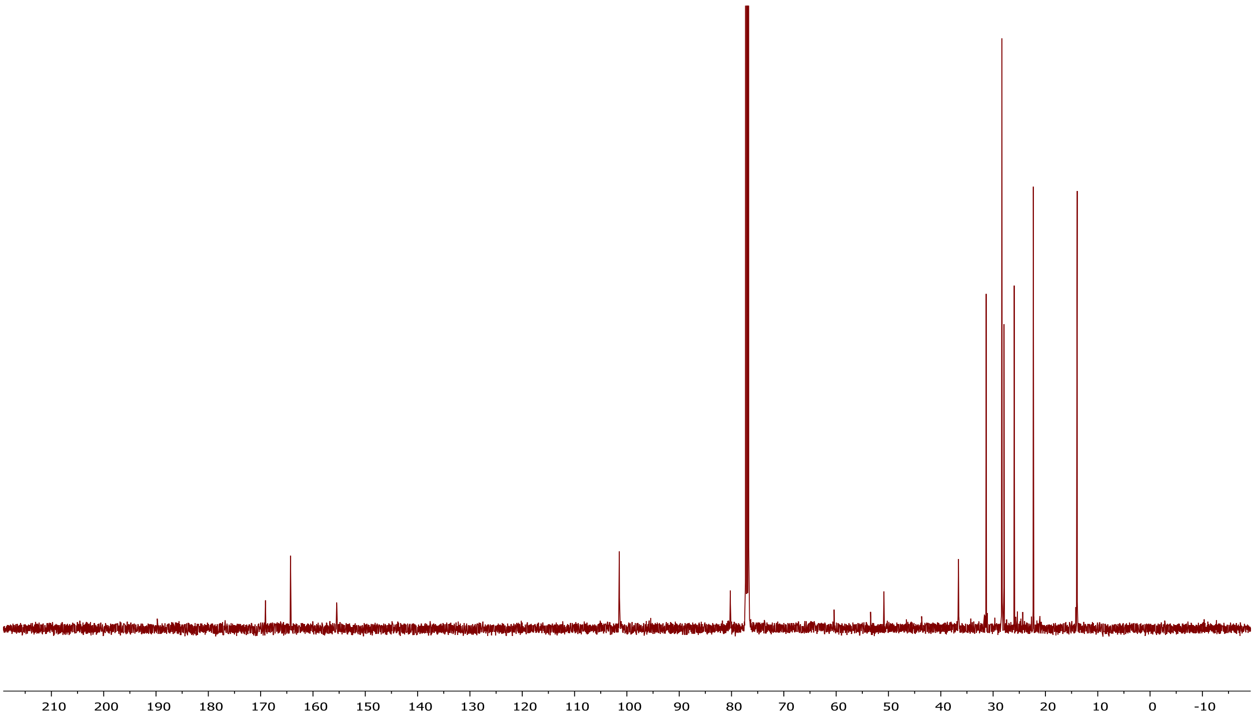

**[1]H NMR of compound 12f**

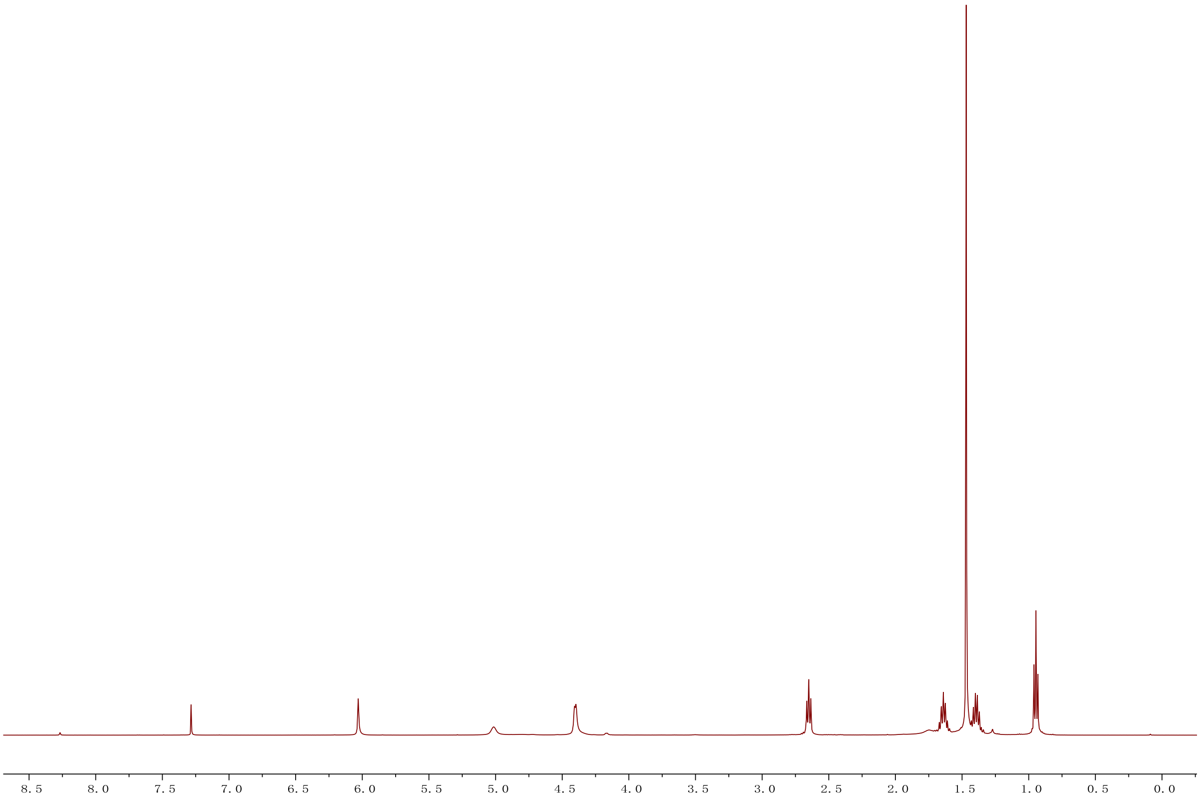


**[13]C NMR of compound 12f**

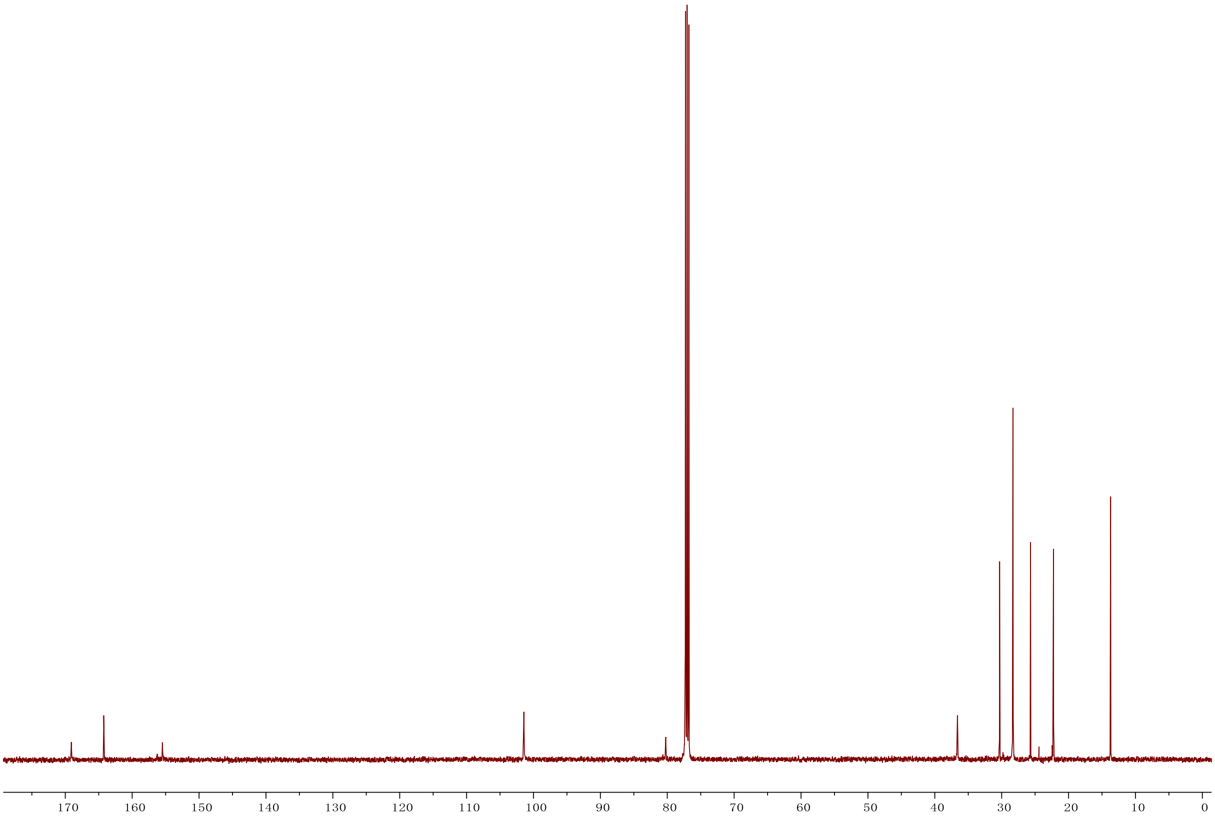

### [1]H NMR of compound 12g

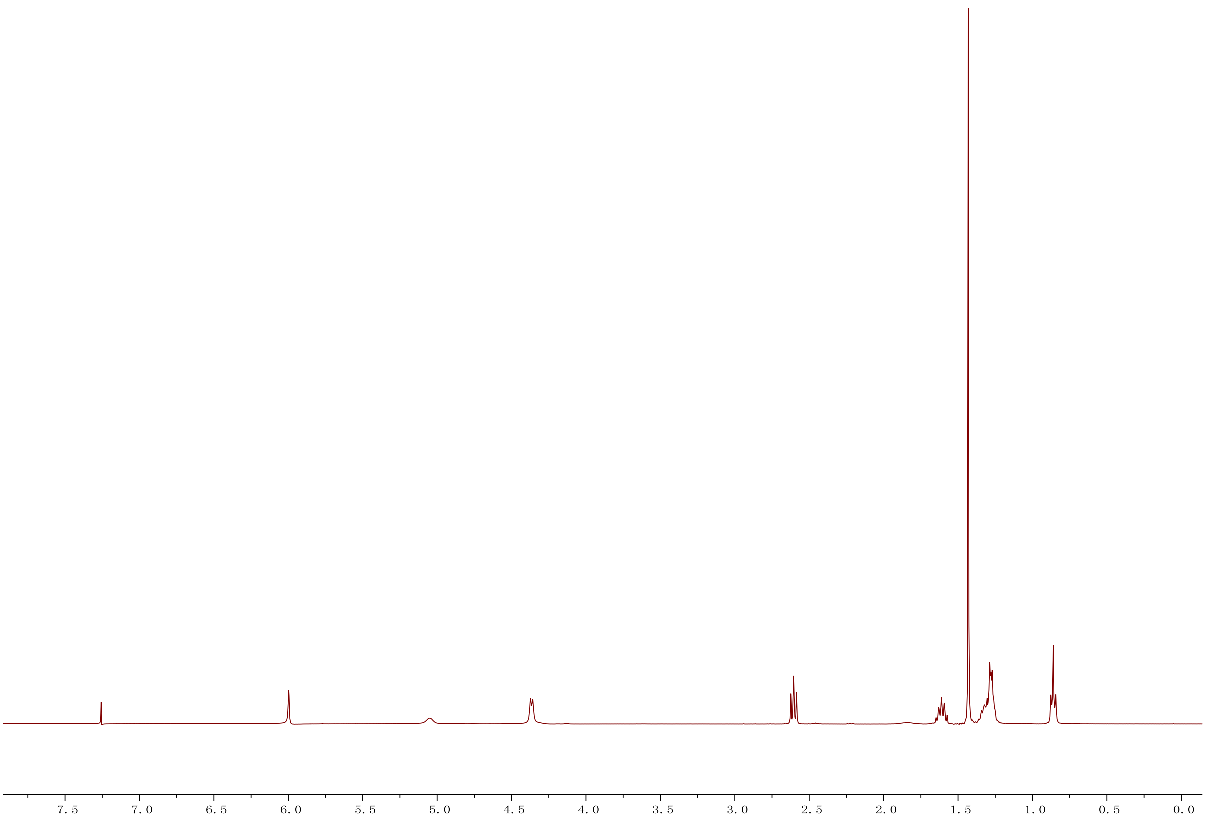


### [13]C NMR of compound 12g

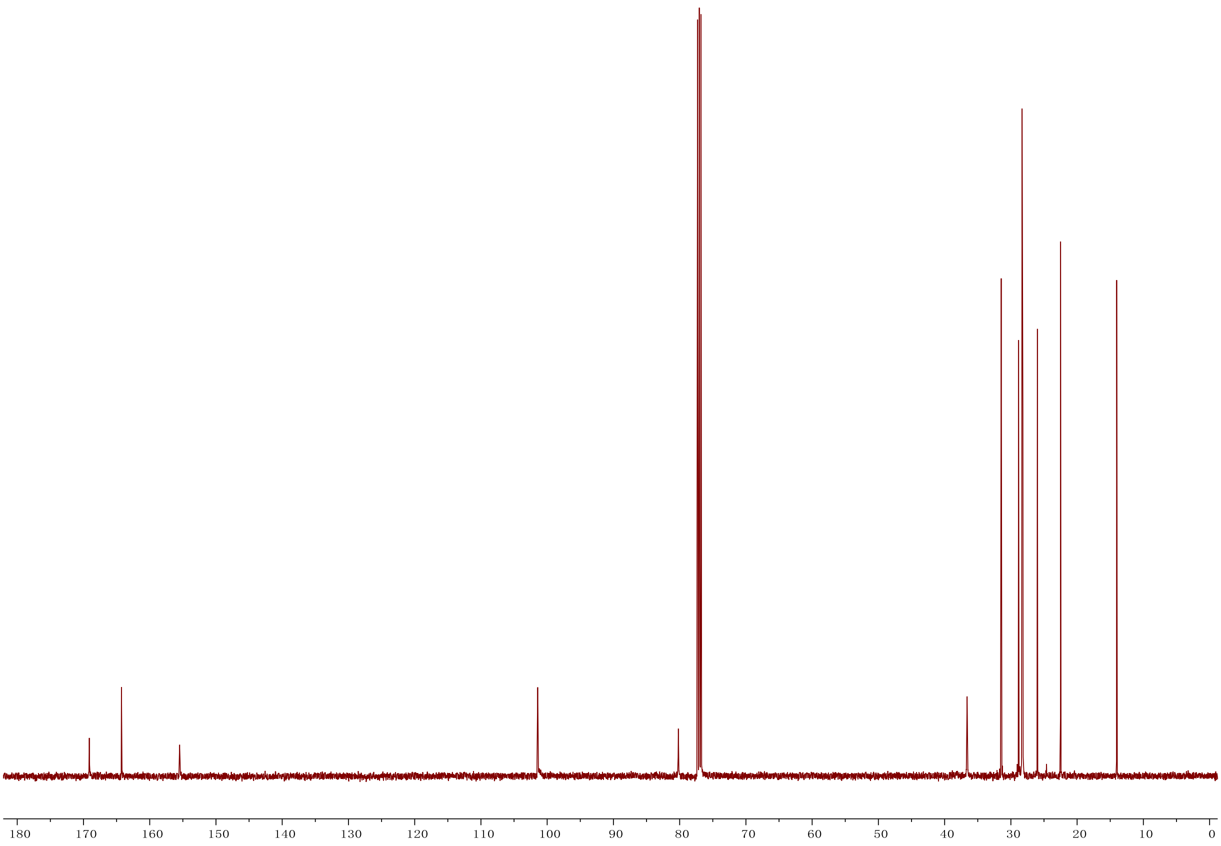

**$^{1}$H NMR of compound 12h**

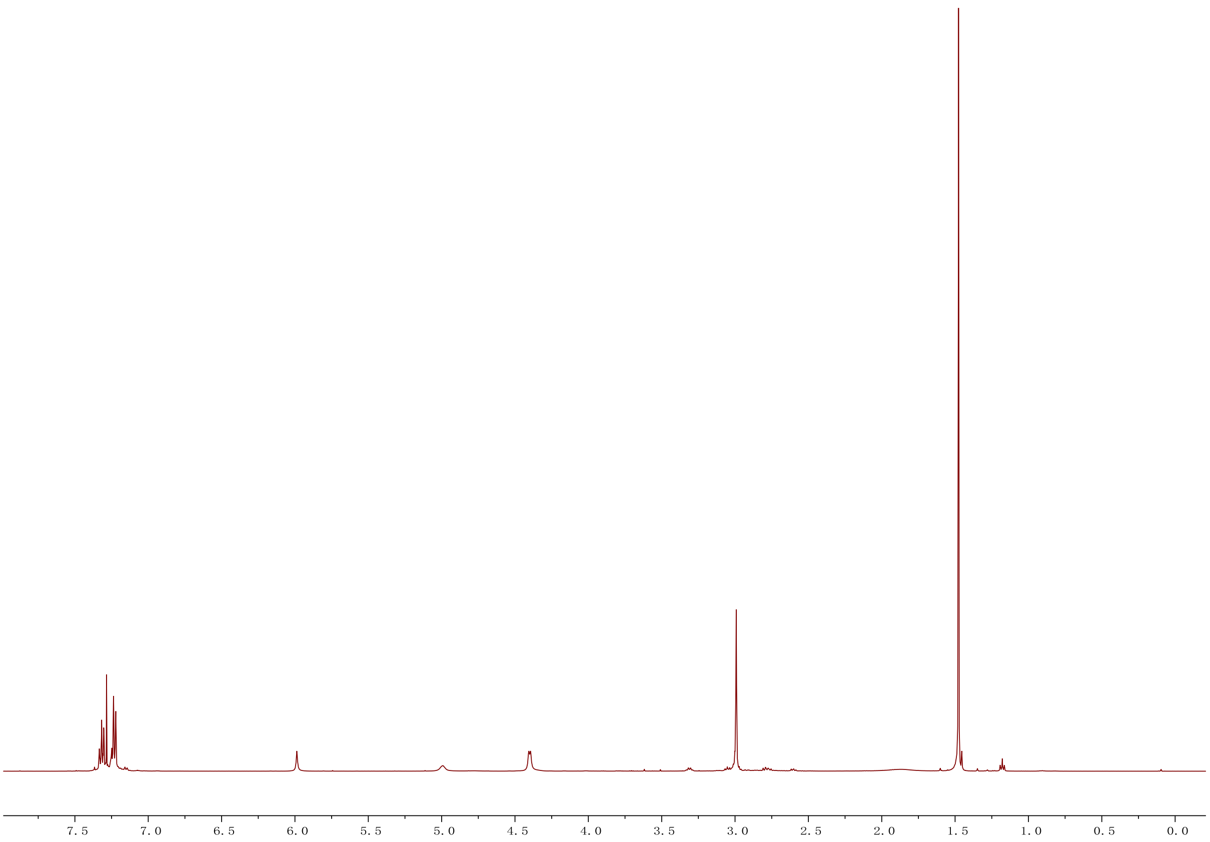


**$^{13}$C NMR of compound 12h**

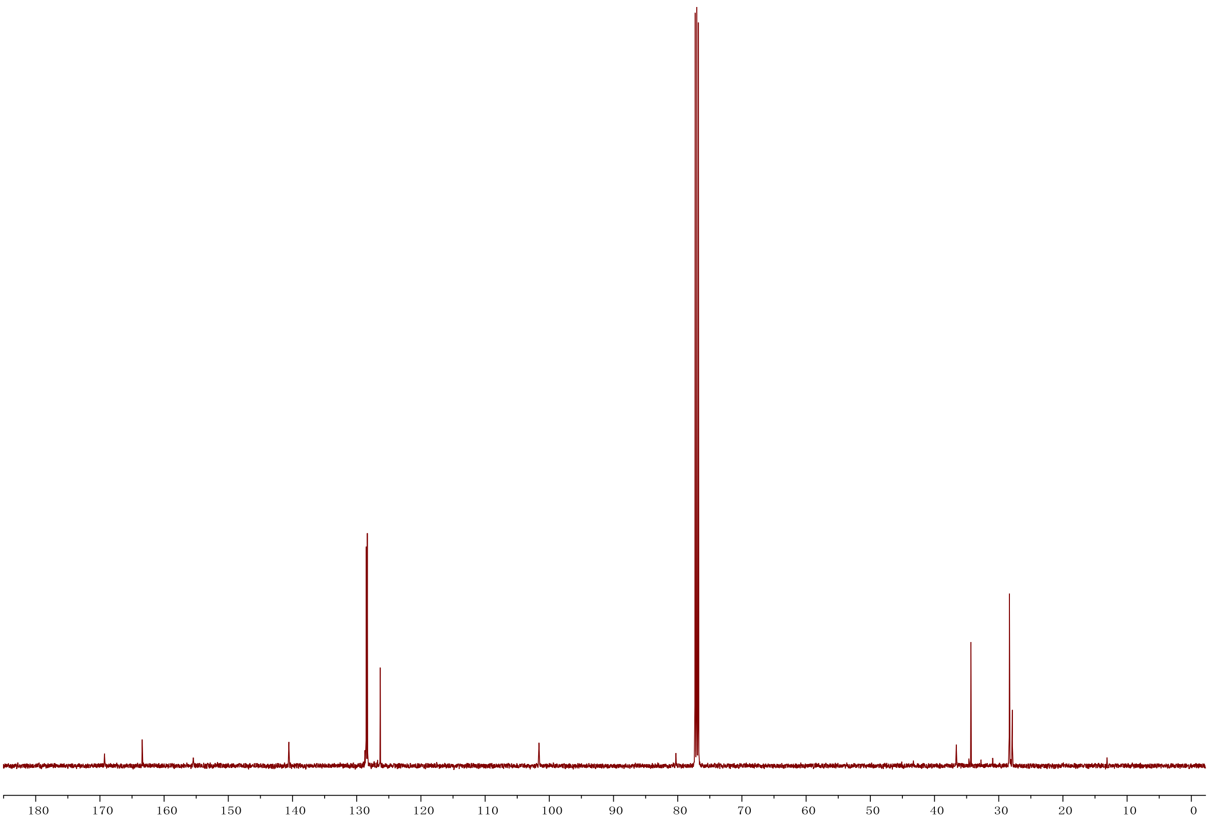

**[1]H NMR of compound 12i**

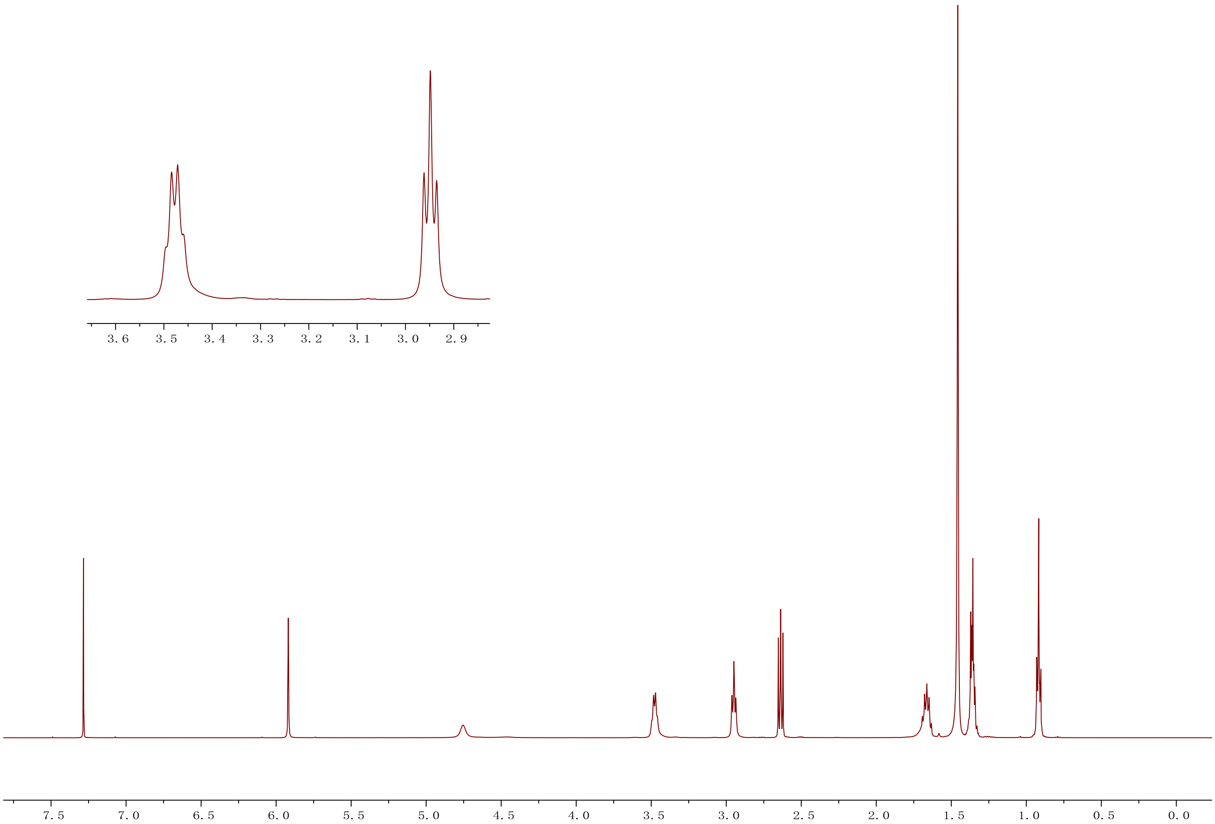


**[13]C NMR of compound 12i**

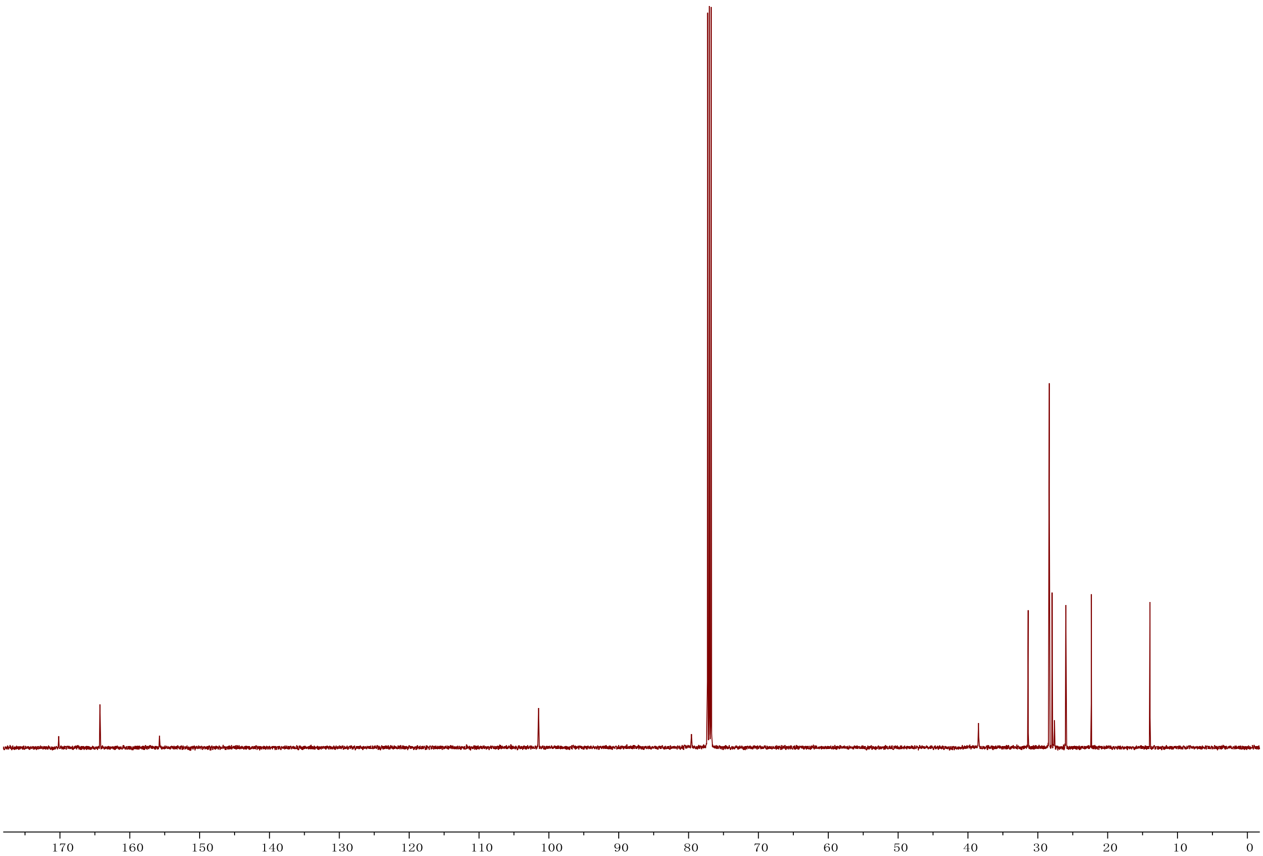

### $^{1}$H NMR of compound 15j

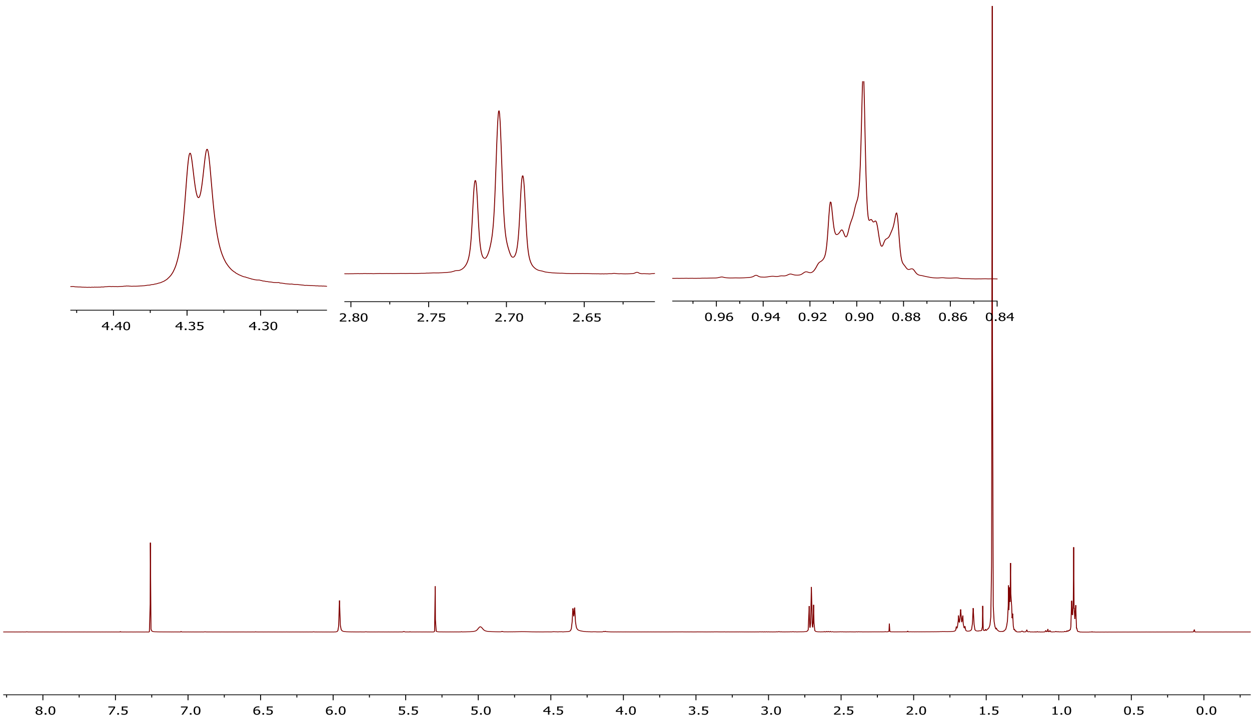


### $^{13}$C NMR of compound 15j

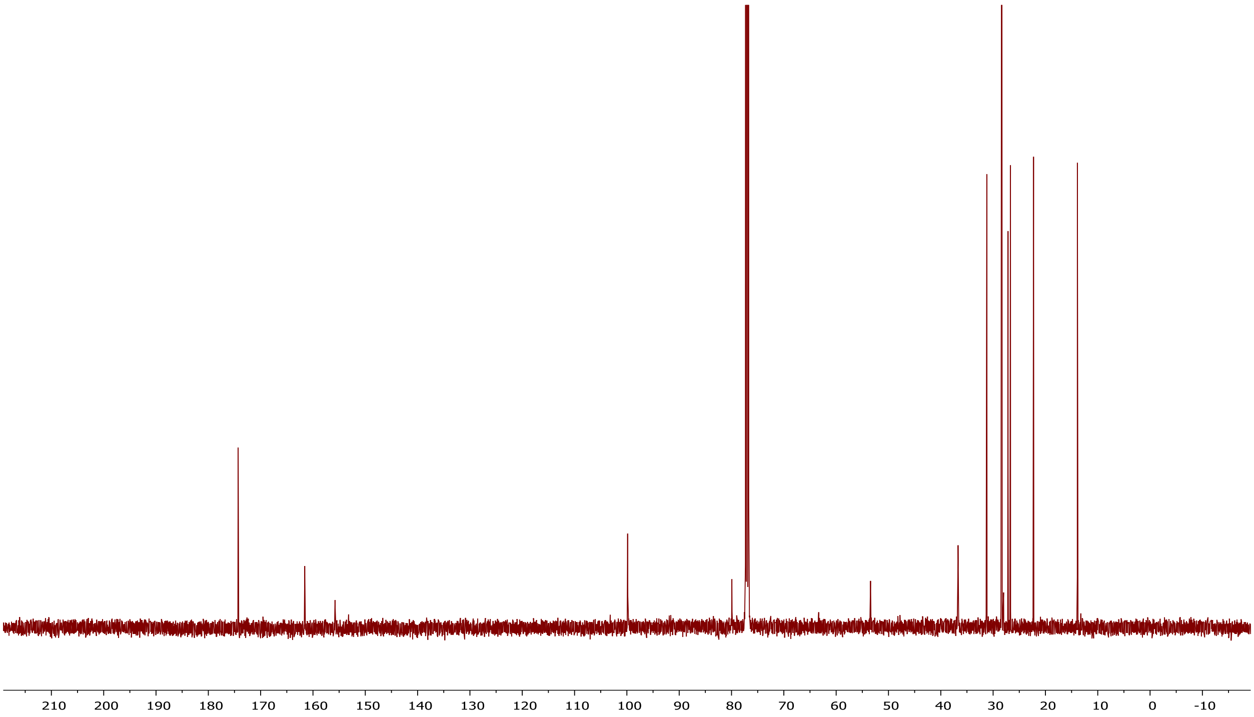

[1]H NMR of compound 15k

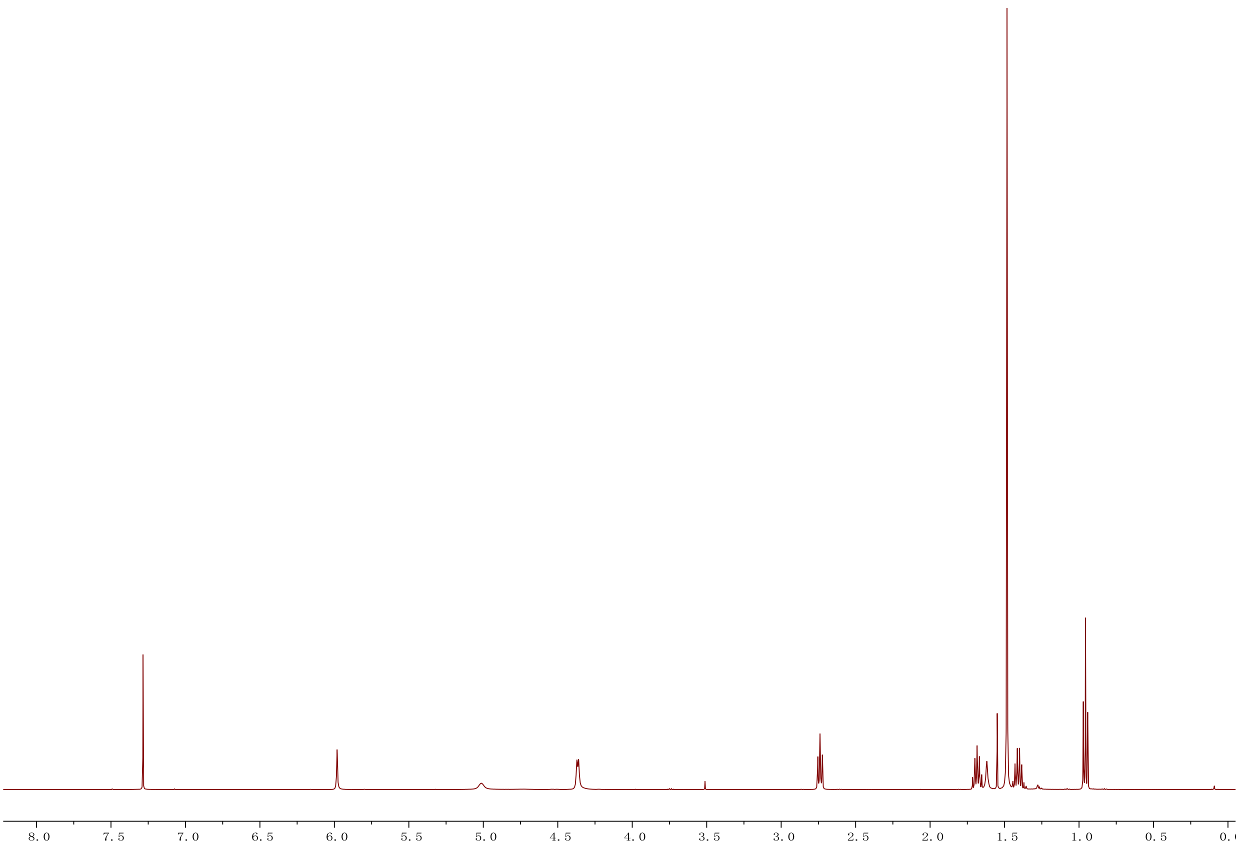


[13]C NMR of compound 15k

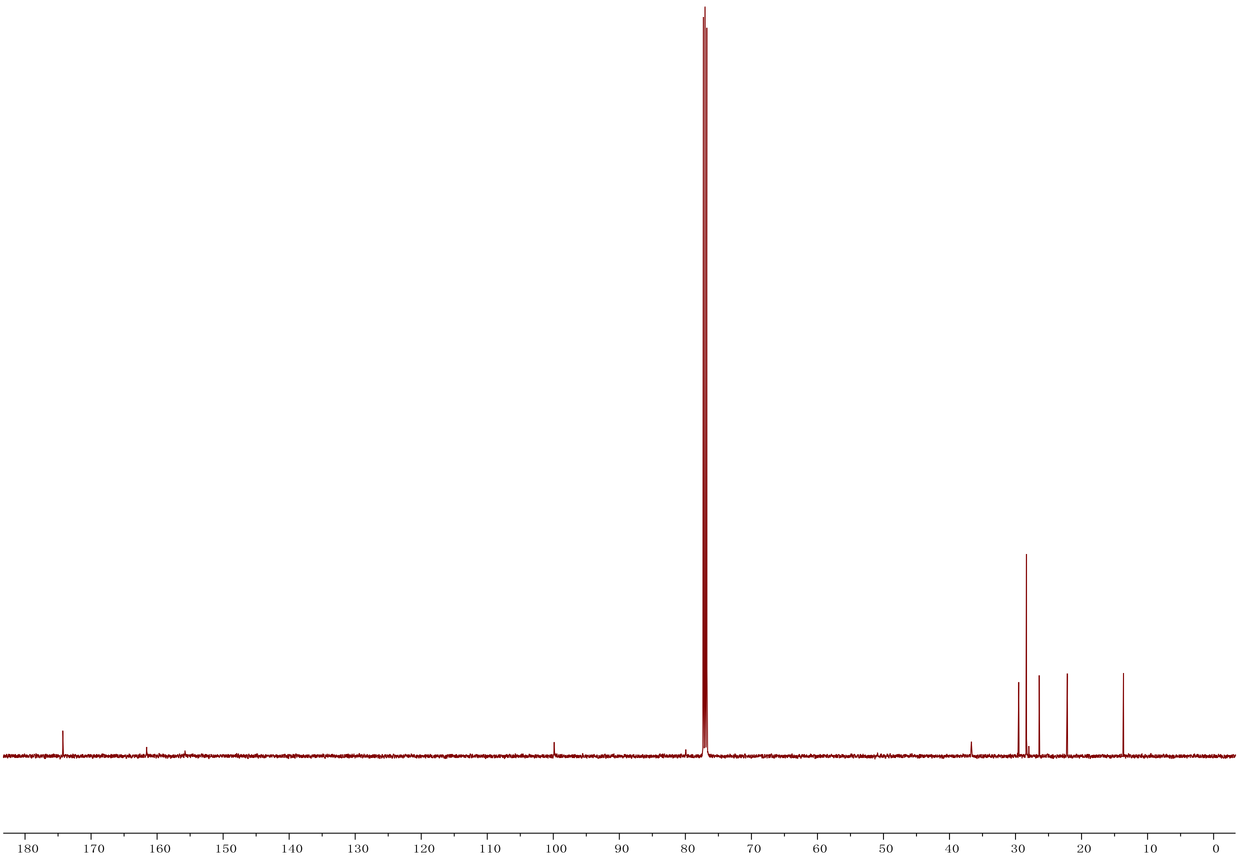

**$^{1}$H NMR of compound 15l**

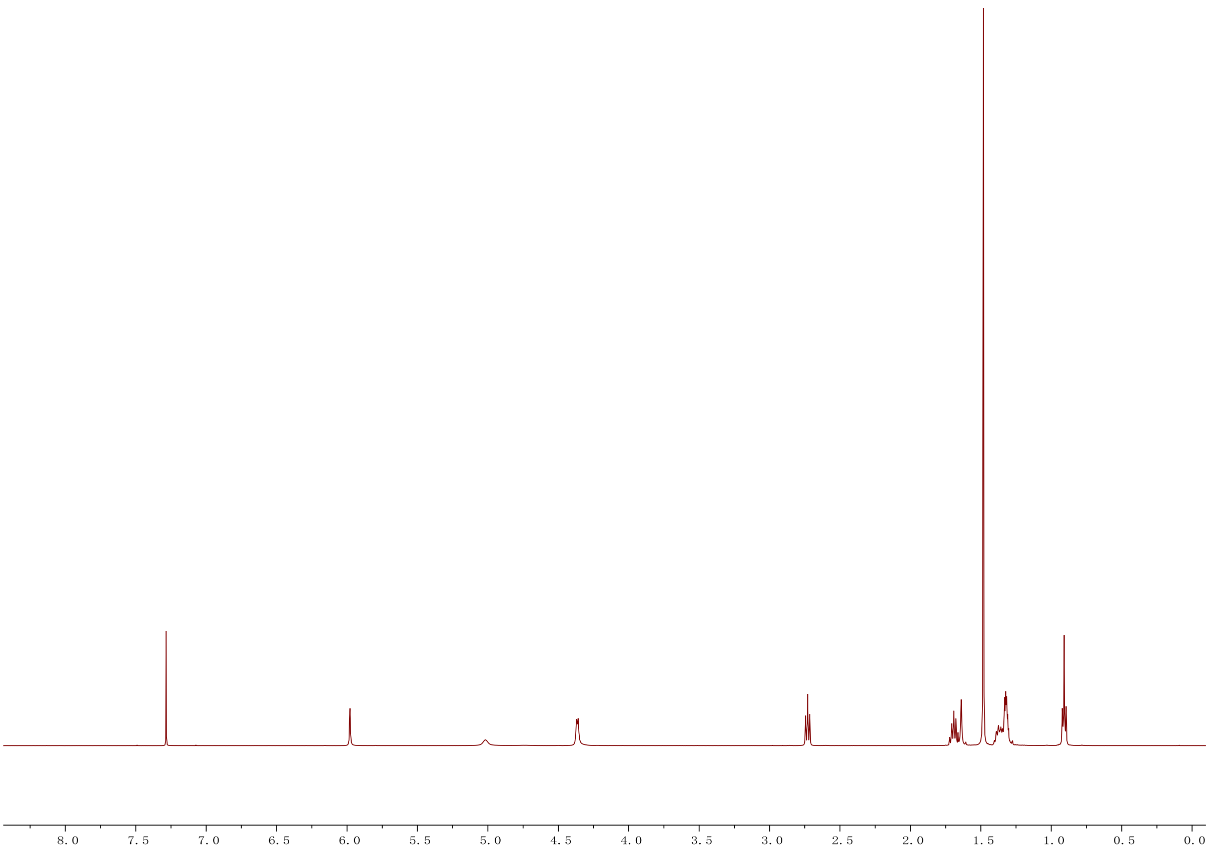


**$^{13}$C NMR of compound 15l**

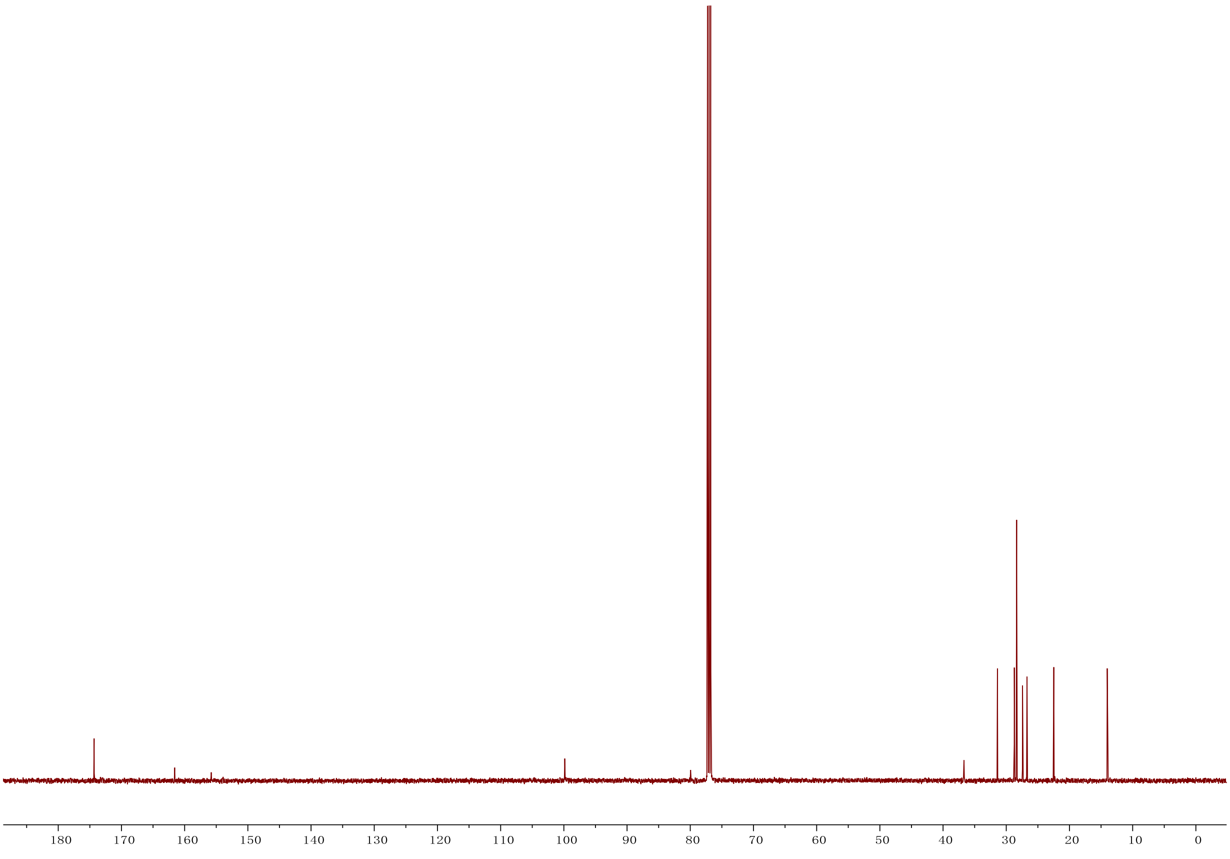

### [1]H NMR of compound 15m

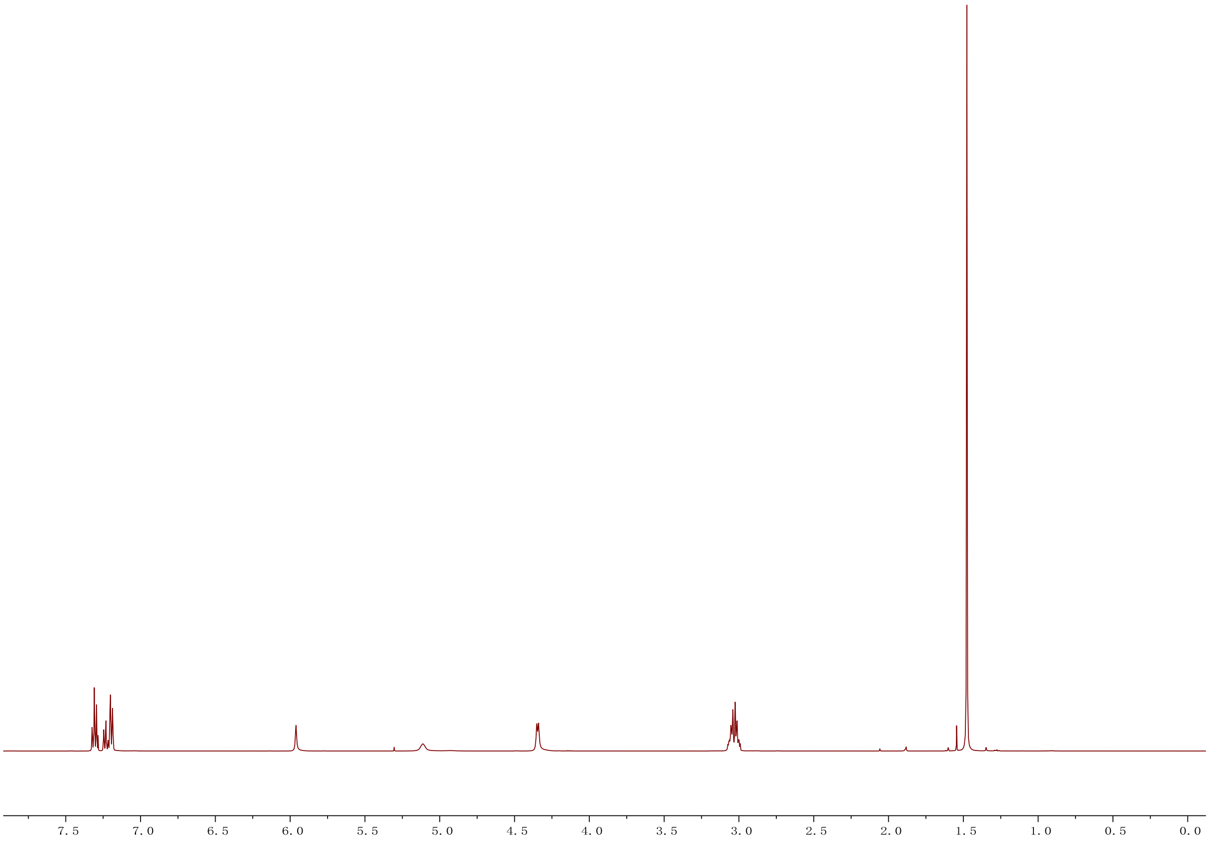


### [13]C NMR of compound 15m

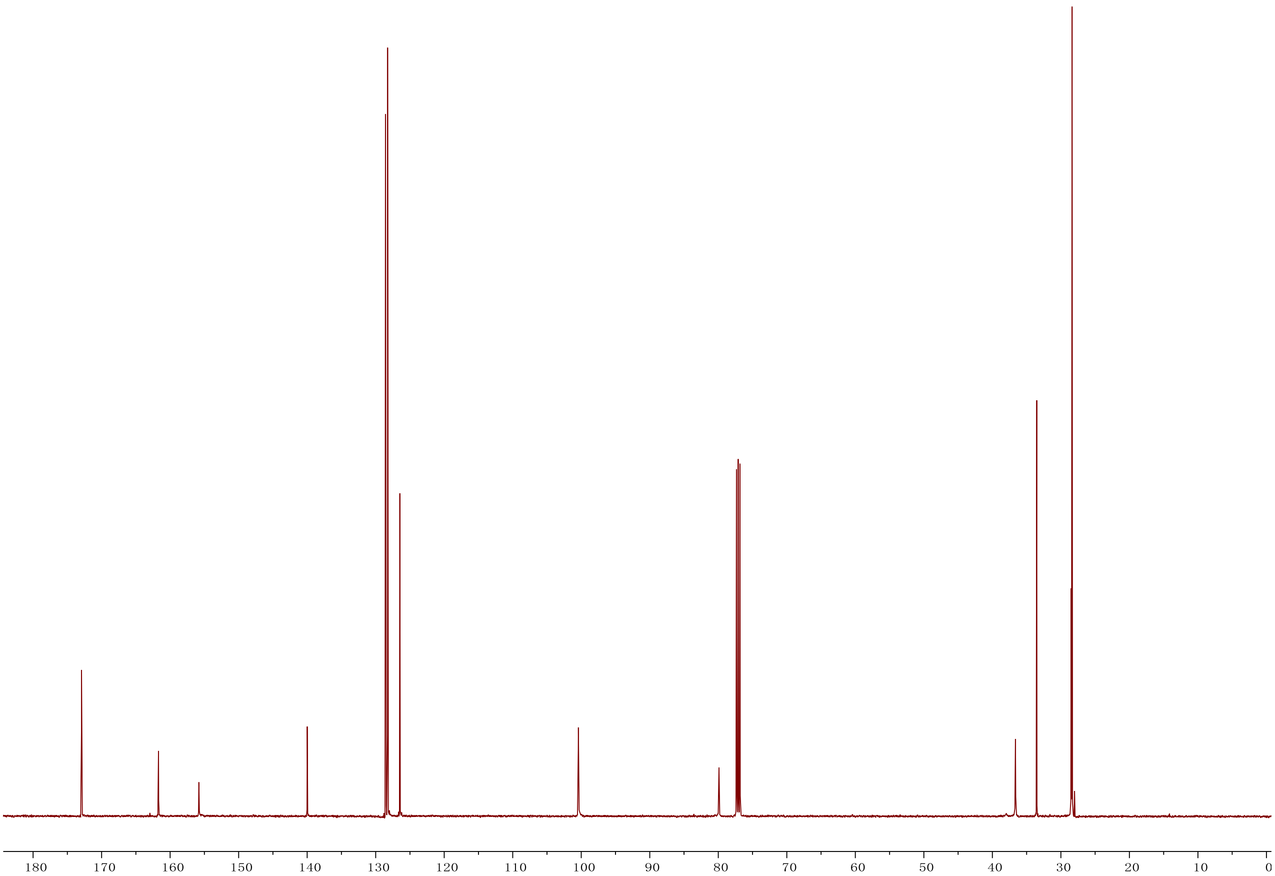

### $^{1}$H NMR of compound 17

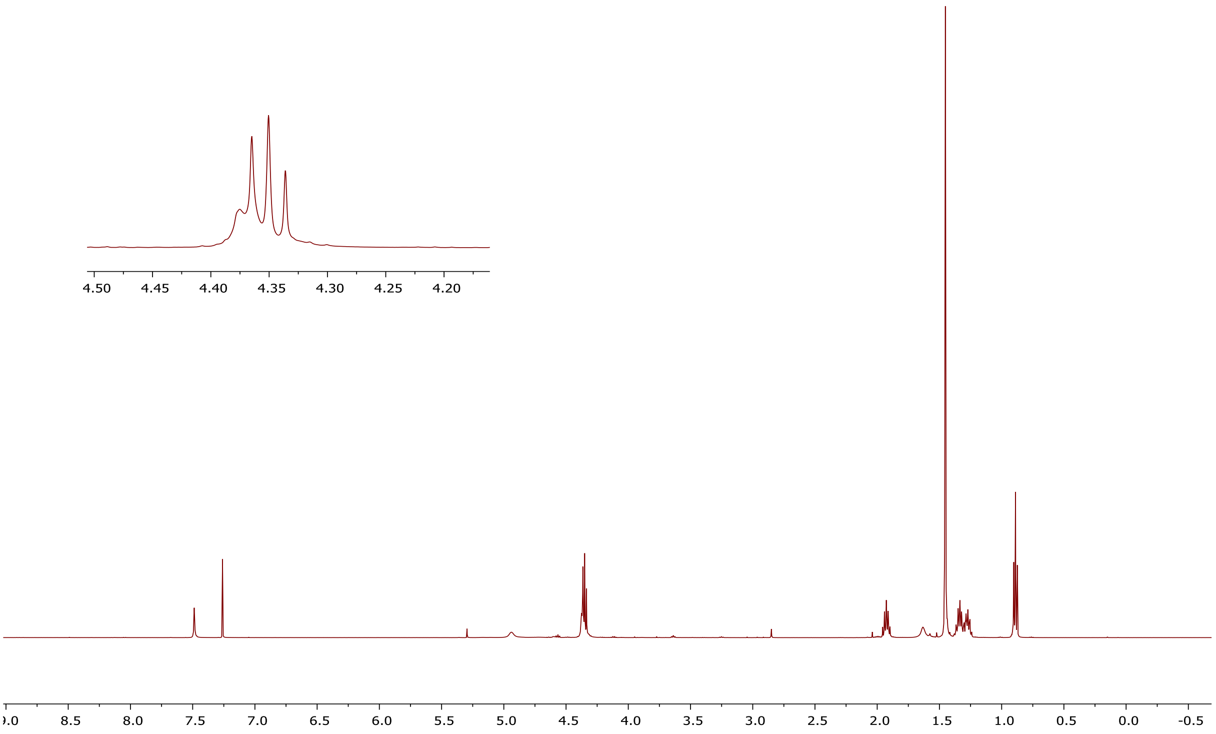


### $^{13}$C NMR of compound 17

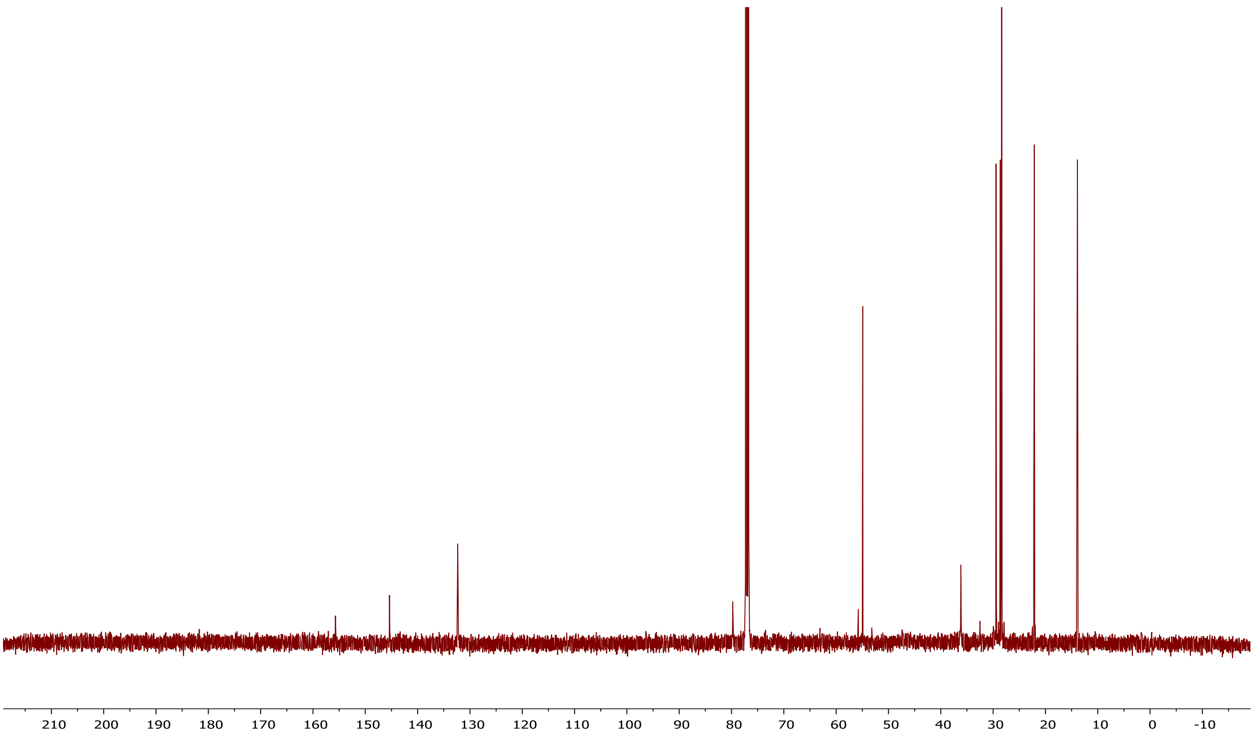

### $^{1}H$ NMR of compound 19

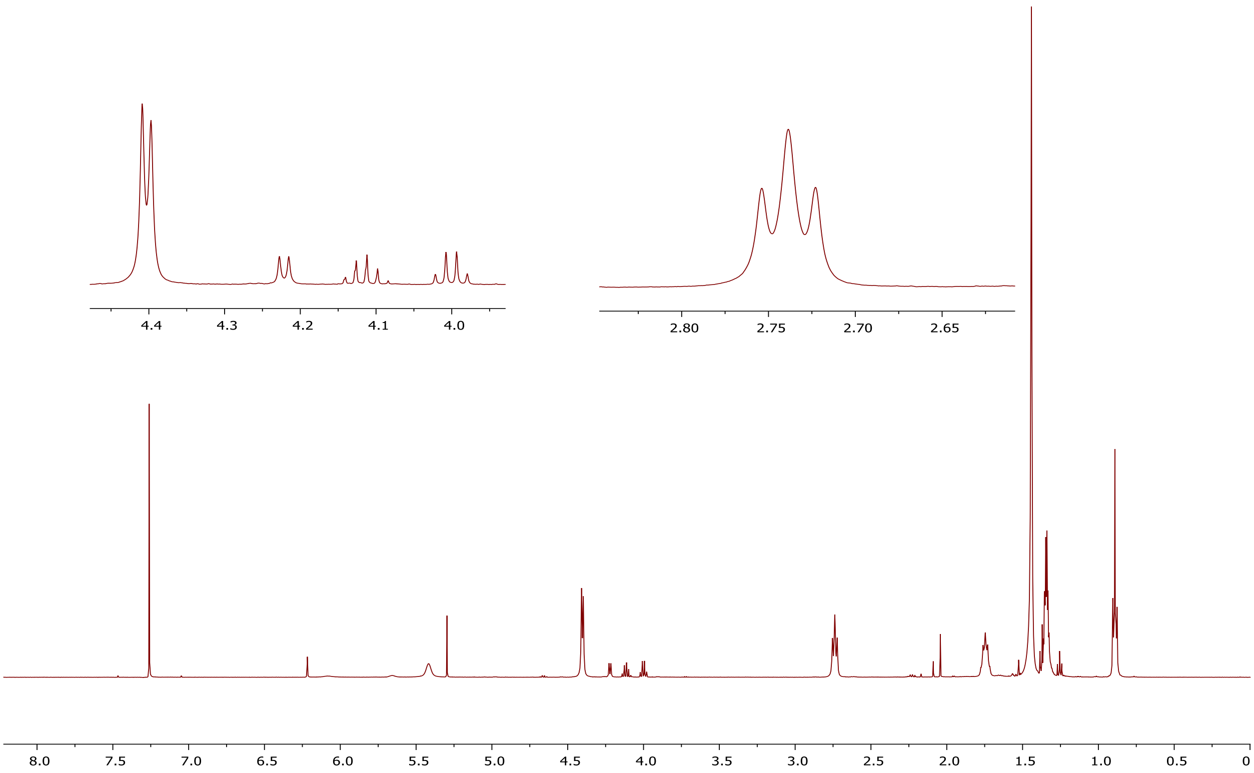


### $^{13}C$ NMR of compound 19

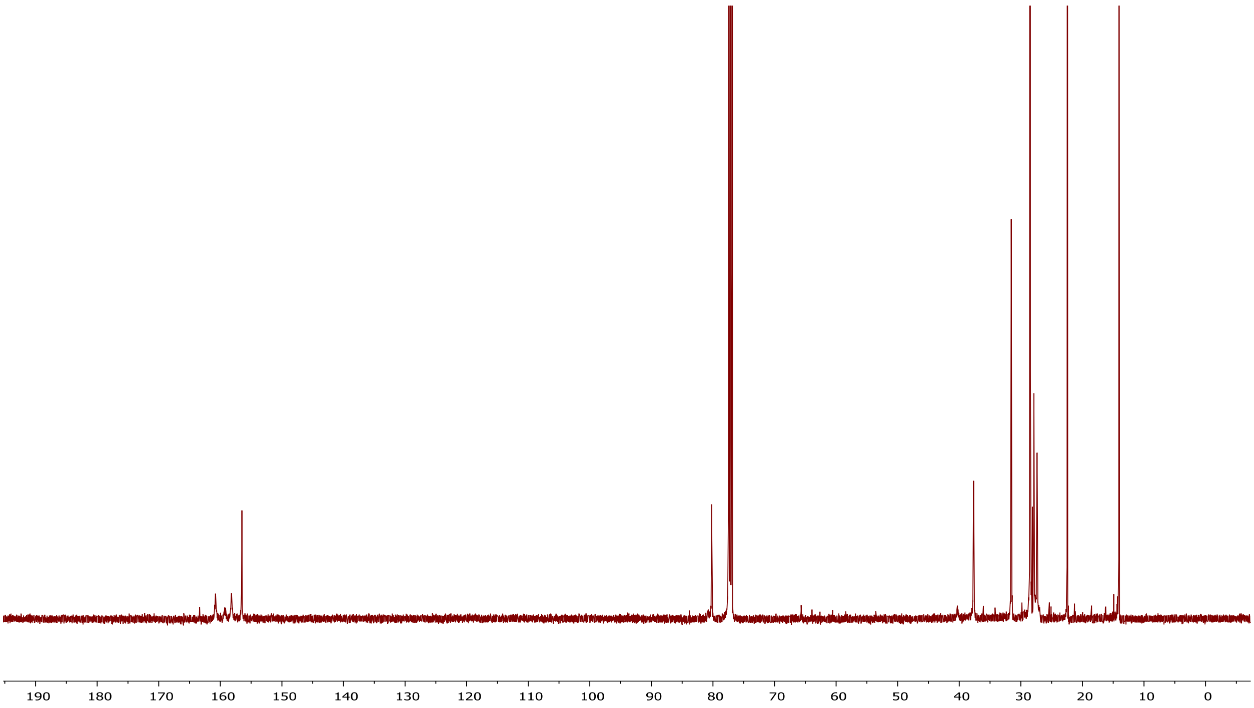

### $^{1}$H NMR of compound 21

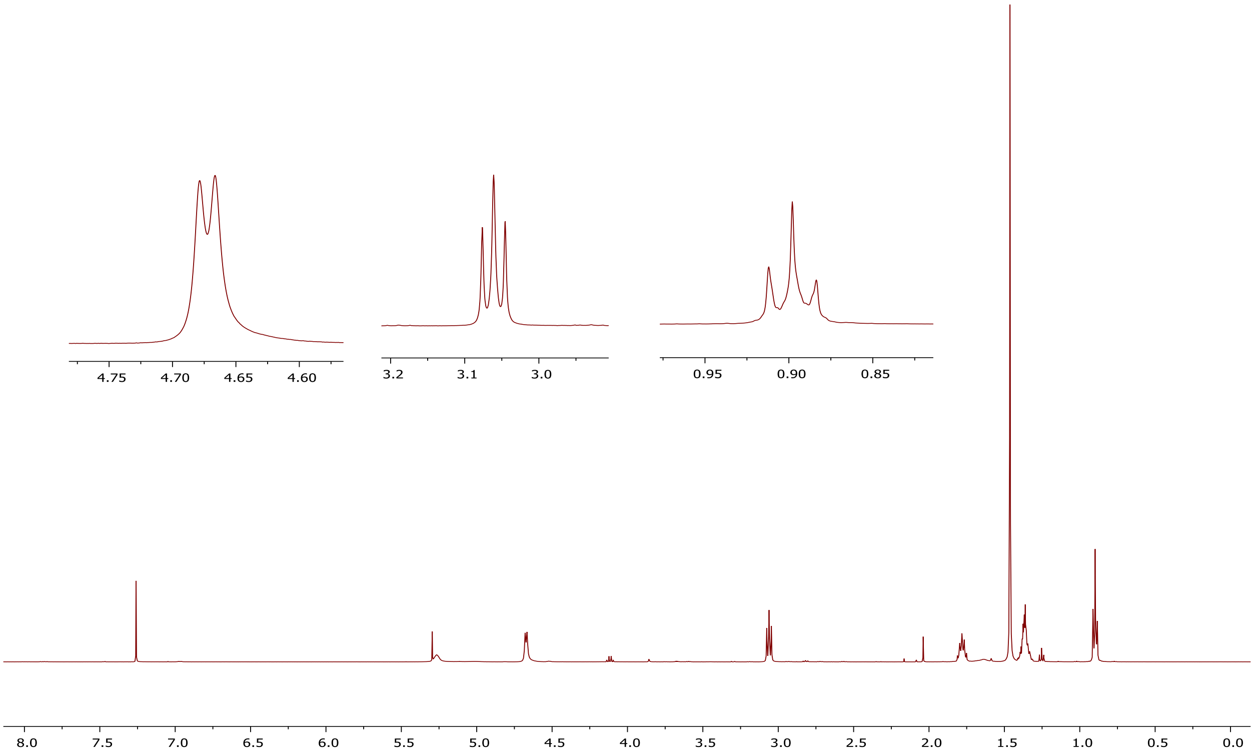


### $^{13}$C NMR of compound 21

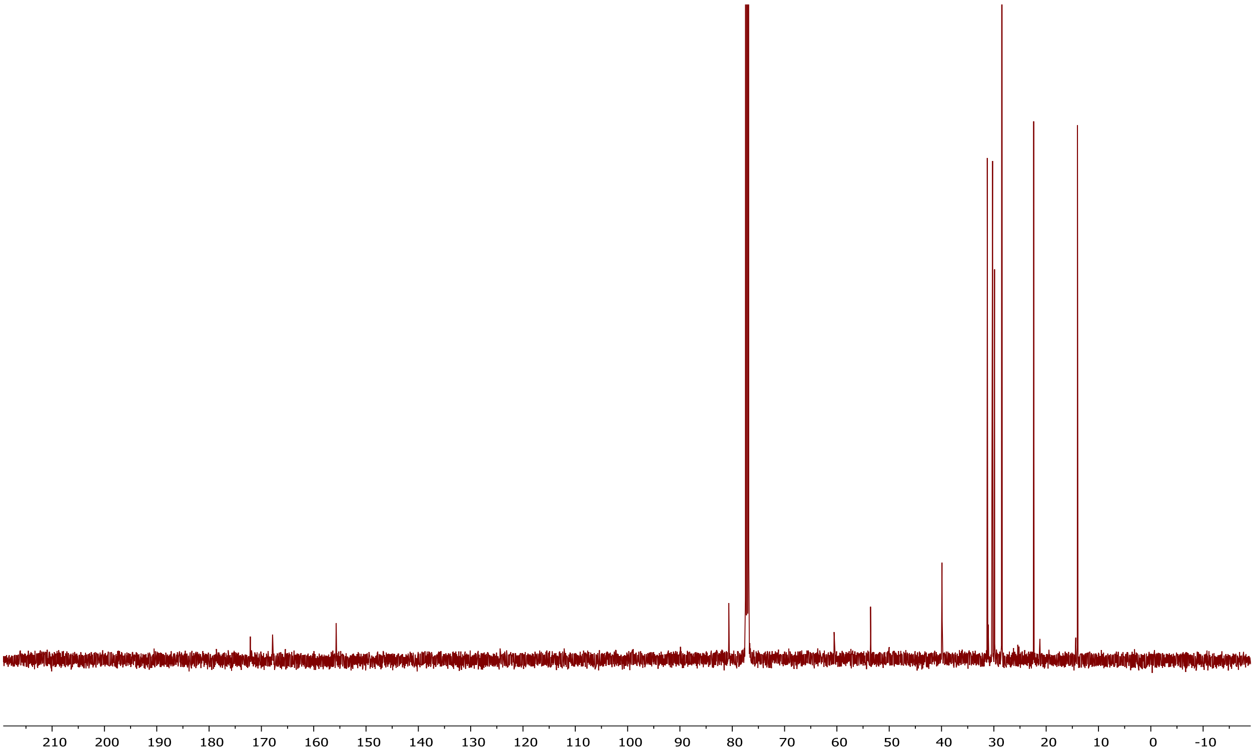

### [1]H NMR of compound 22

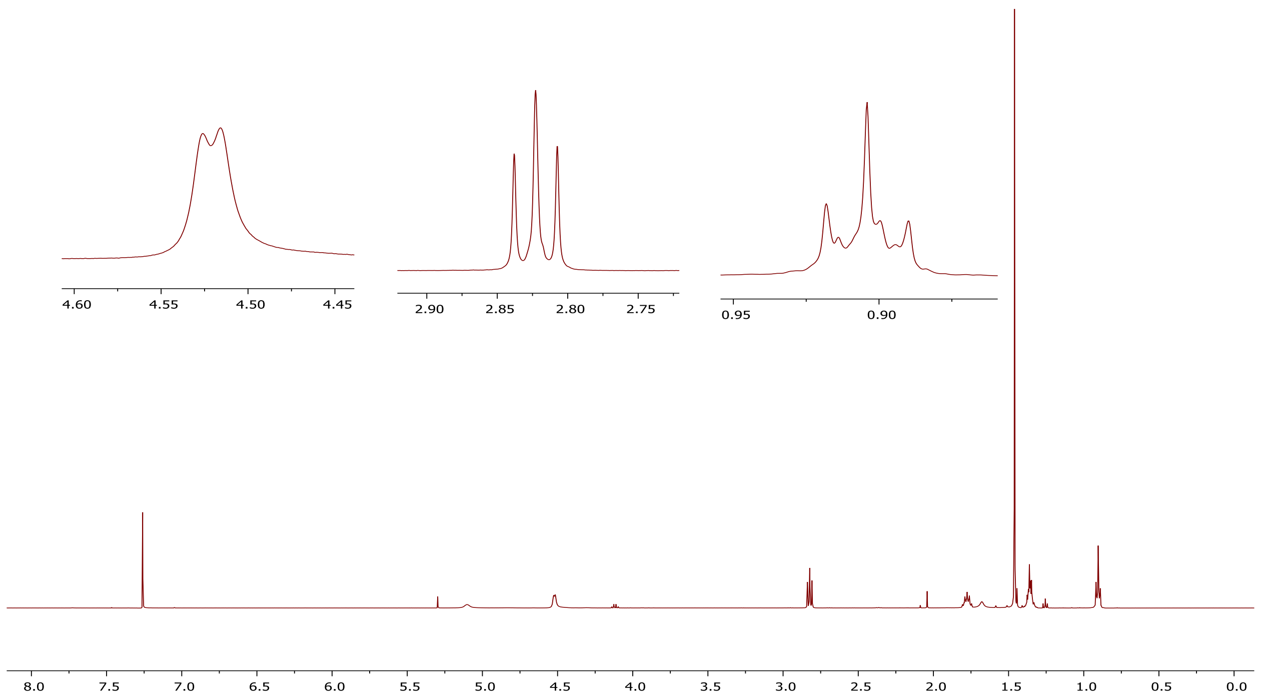


### [13]C NMR of compound 22

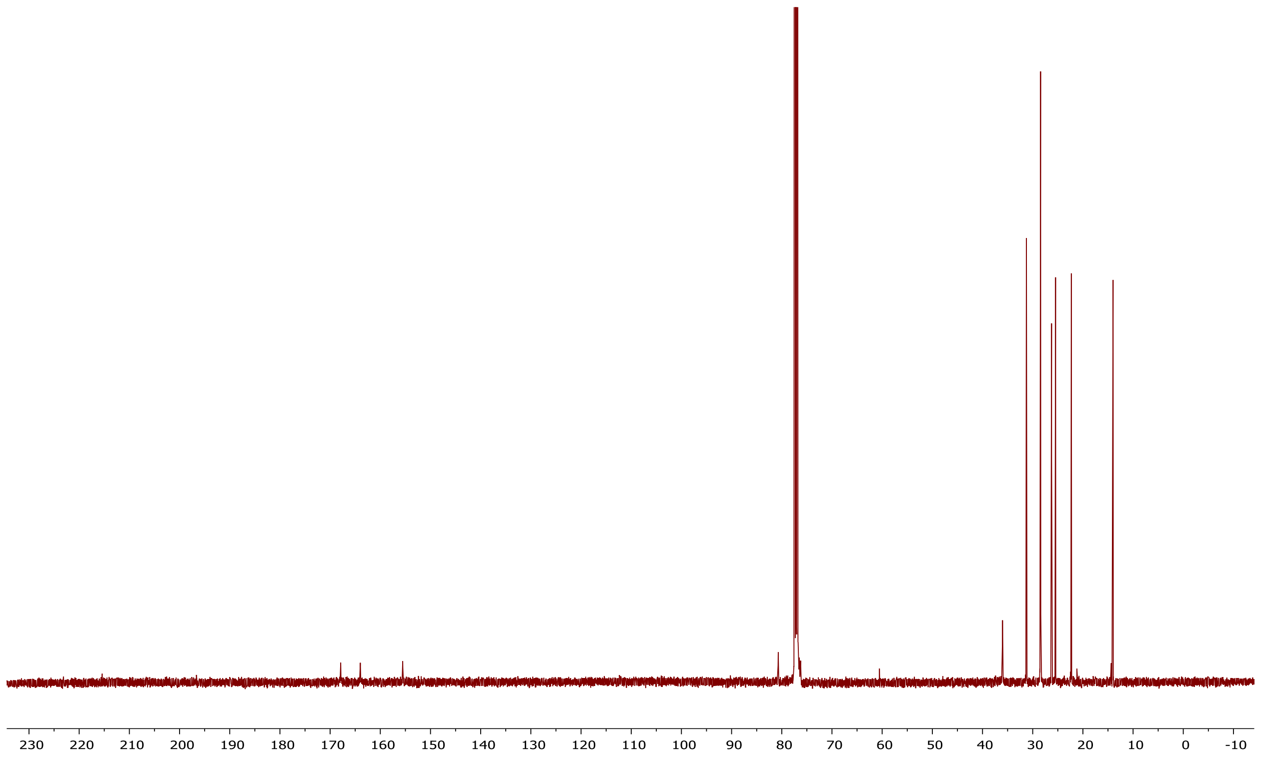

### [1]H NMR of compound 24r

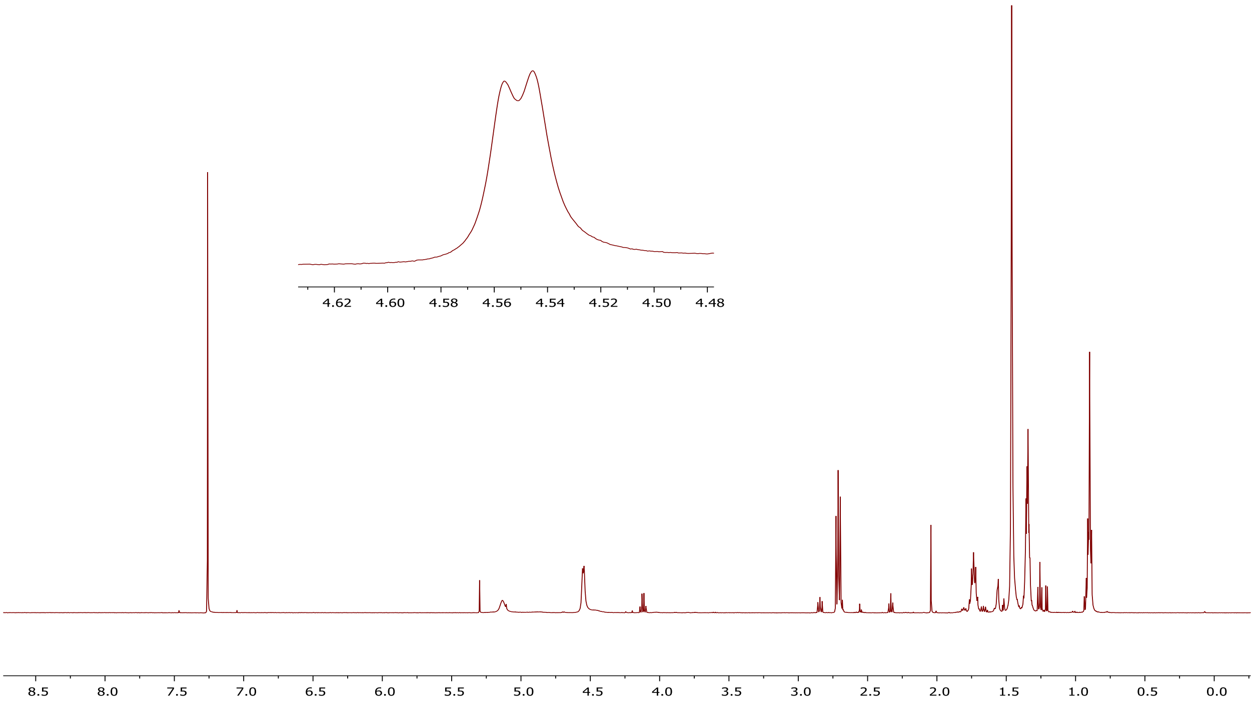


### [13]C NMR of compound 24r

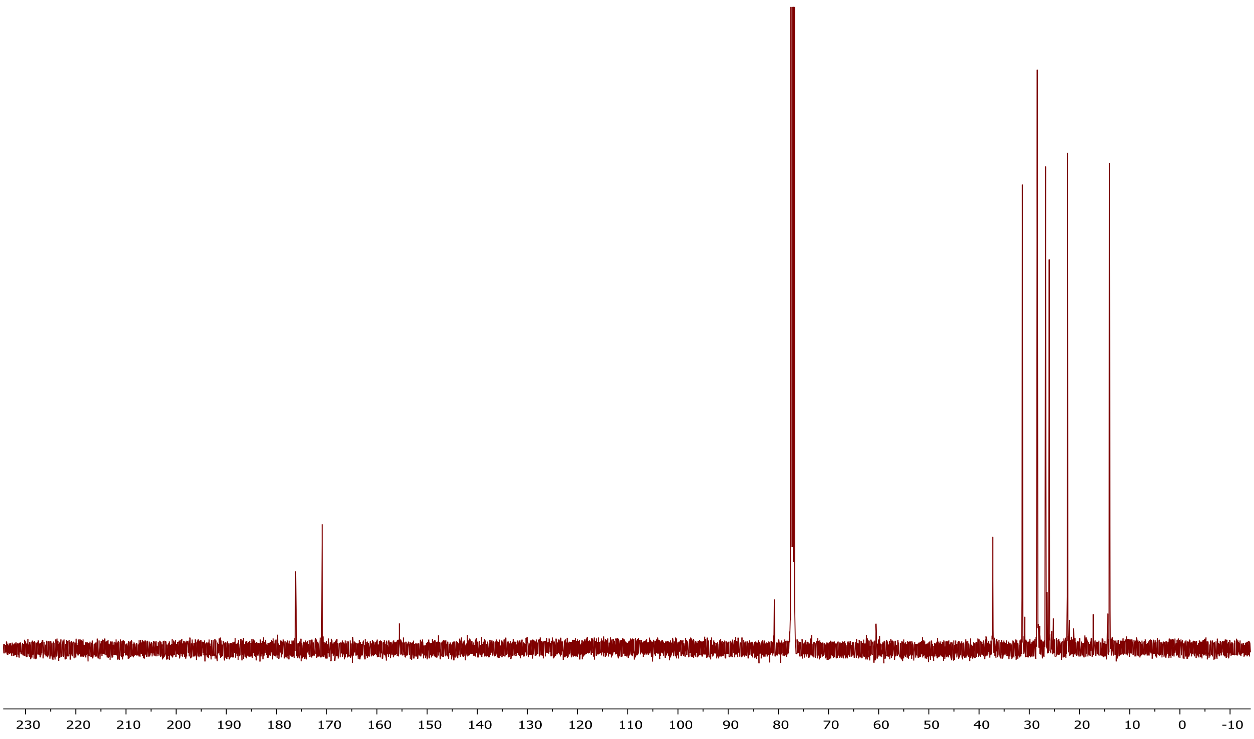

### [1]H NMR of compound 24s

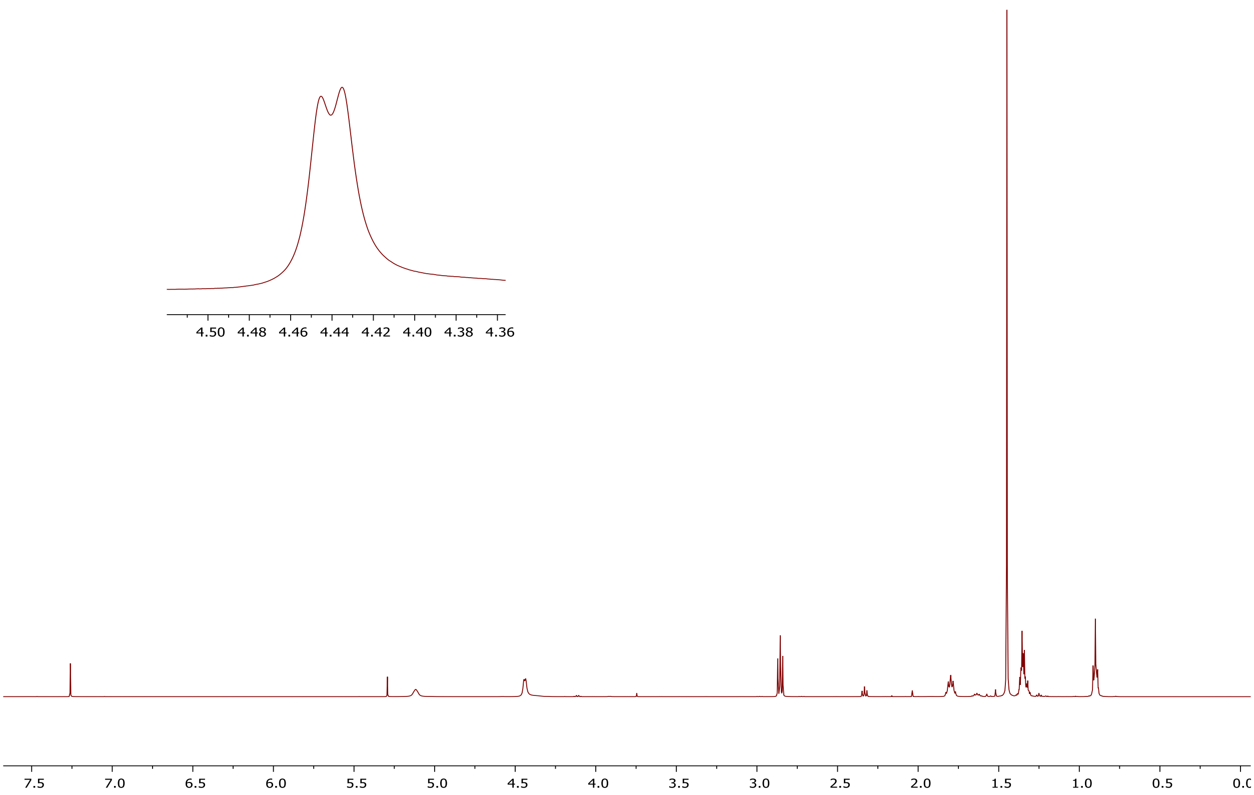


### [13]C NMR of compound 24s

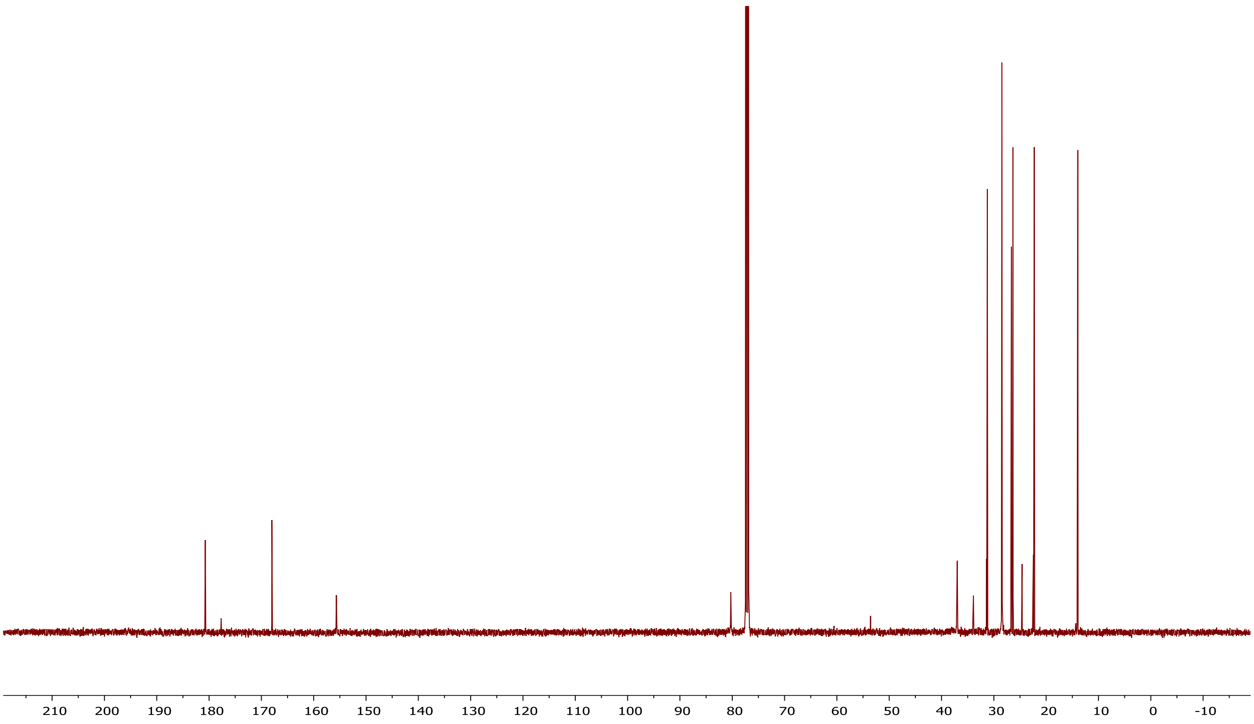

### $^{1}$H NMR of compound 25t

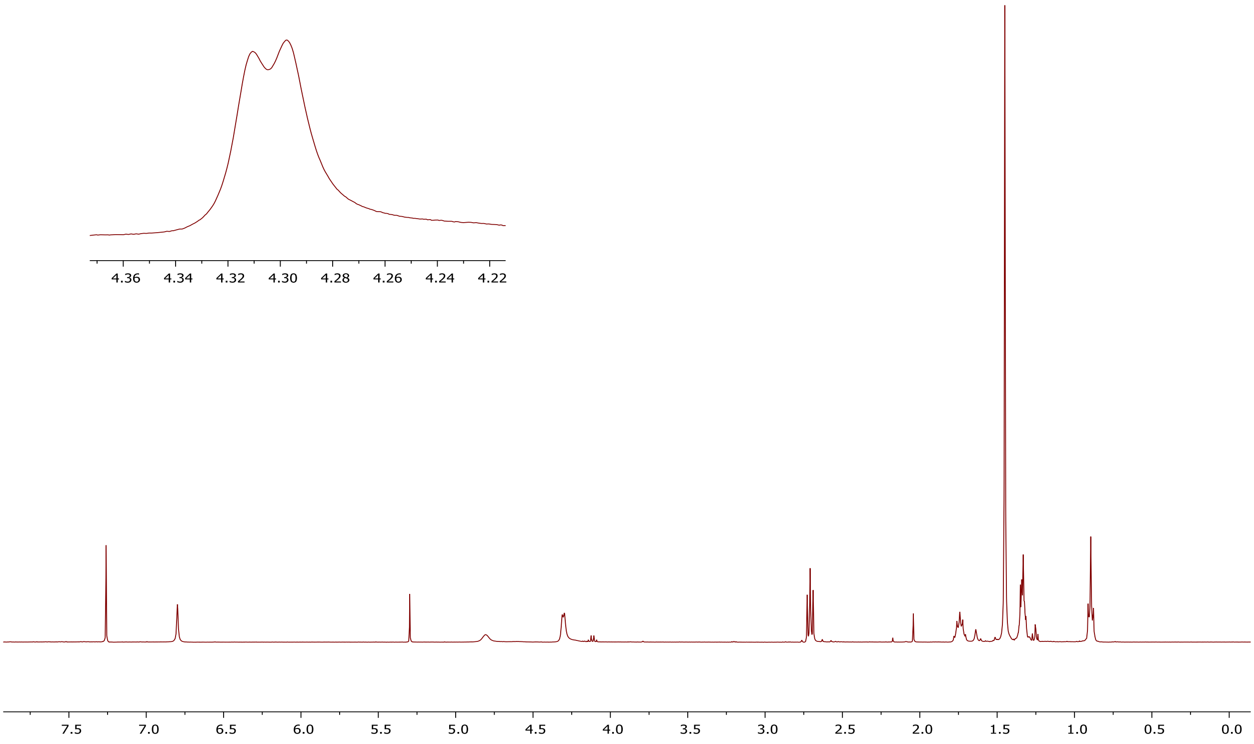


### $^{13}$C NMR of compound 25t

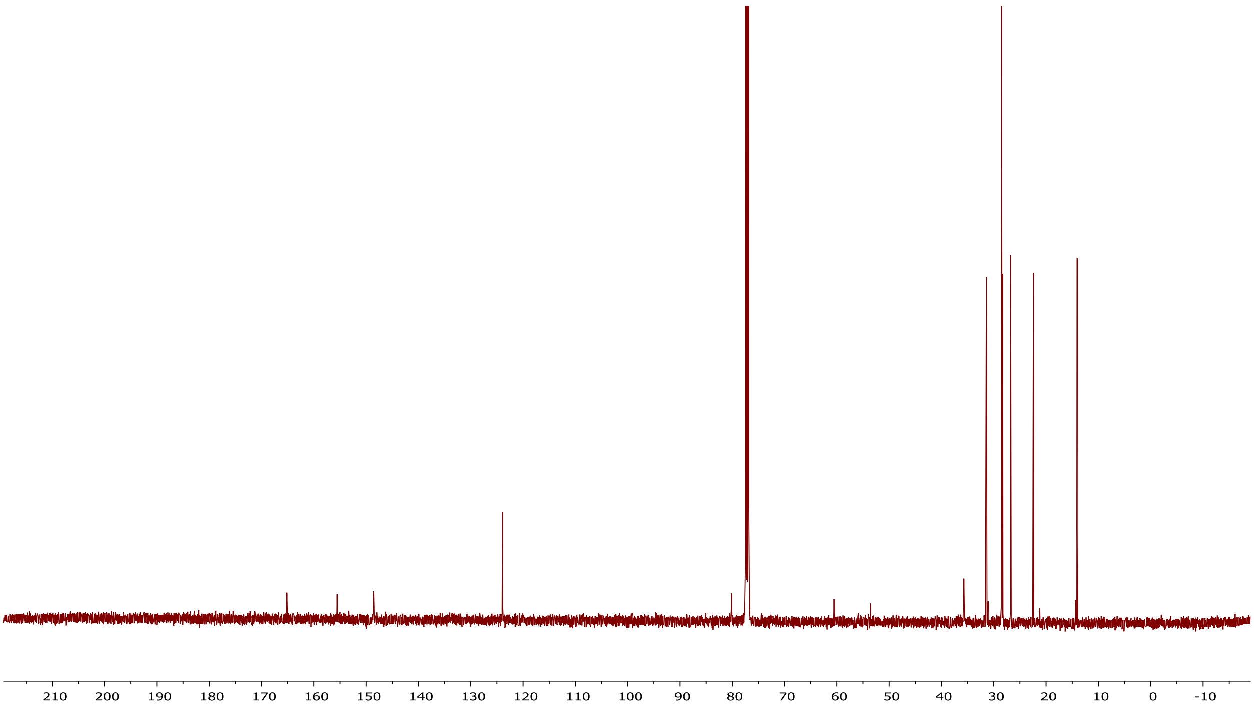

### $^{1}$H NMR of compound 25u

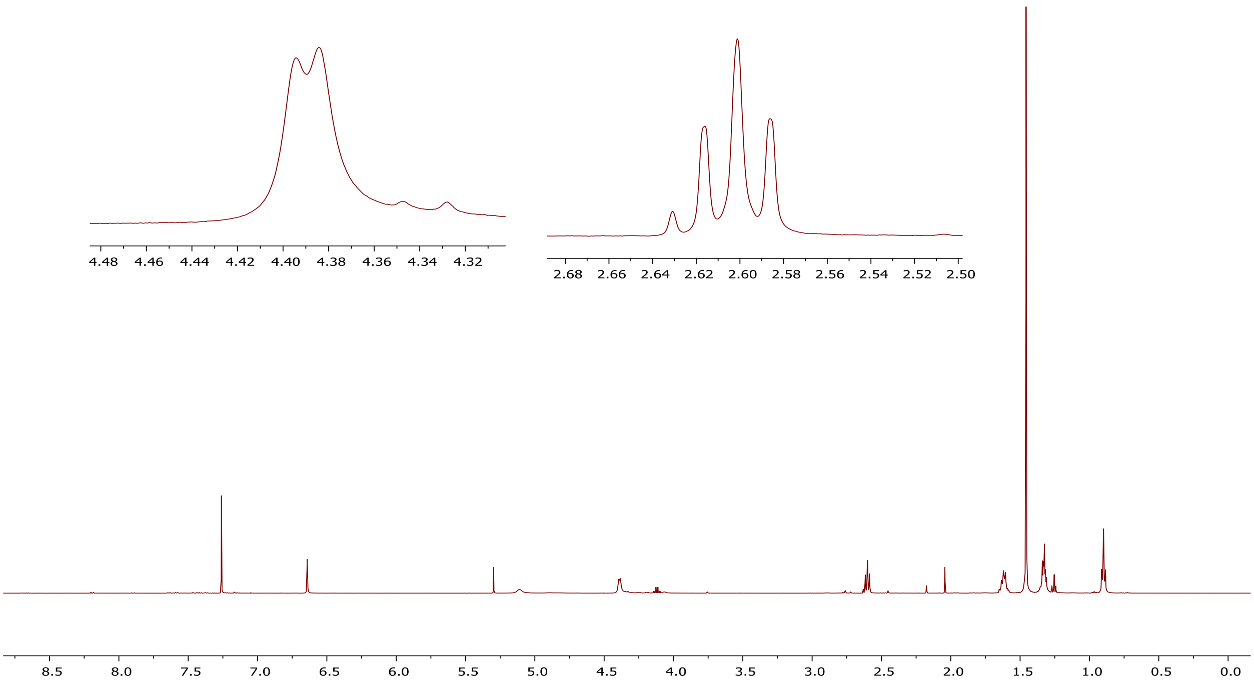


### $^{13}$C NMR of compound 25u

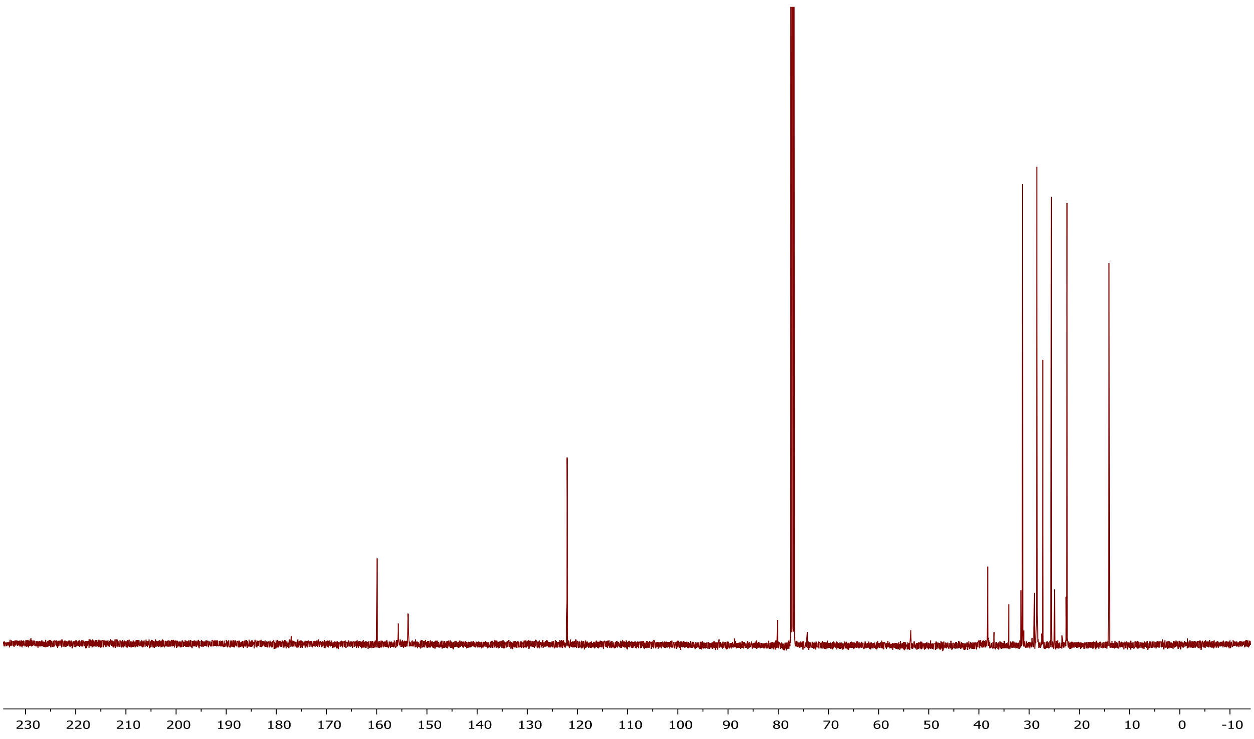

**[1]H NMR of compound 26**

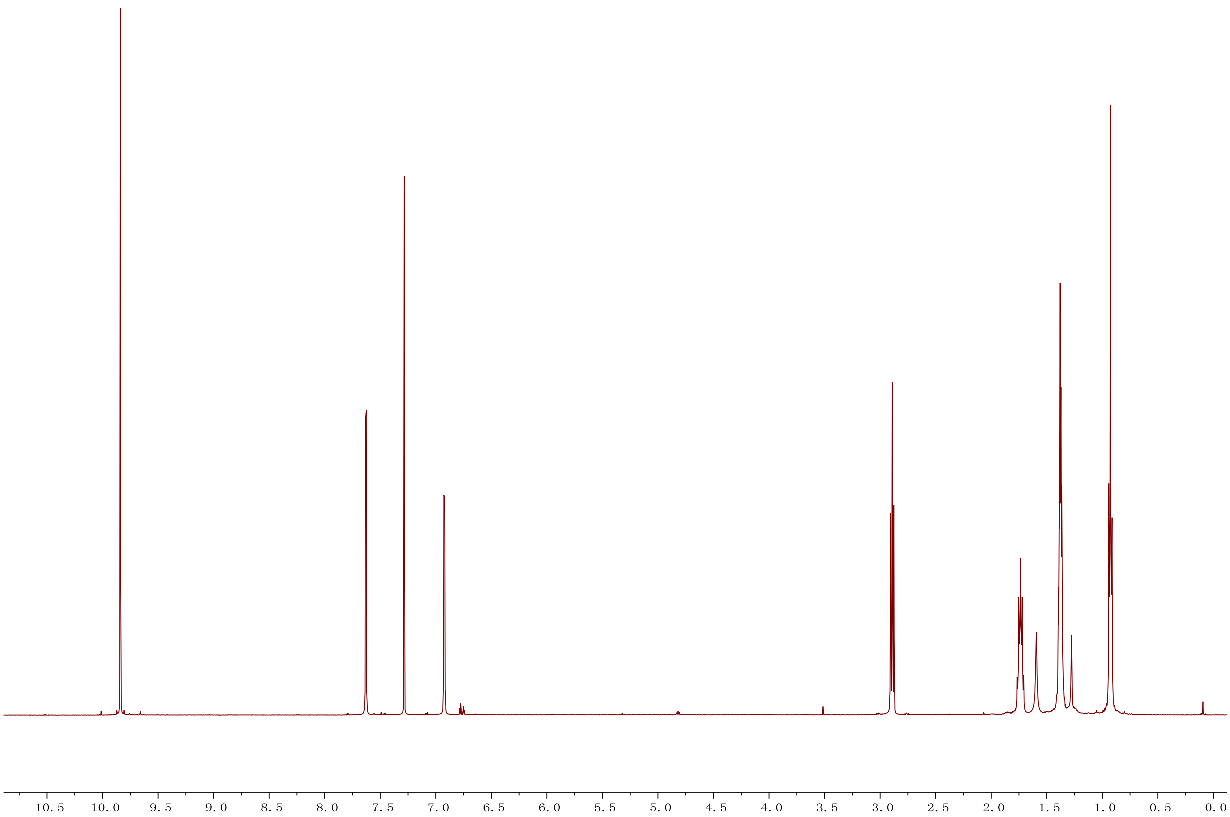


**[13]C NMR of compound 26**

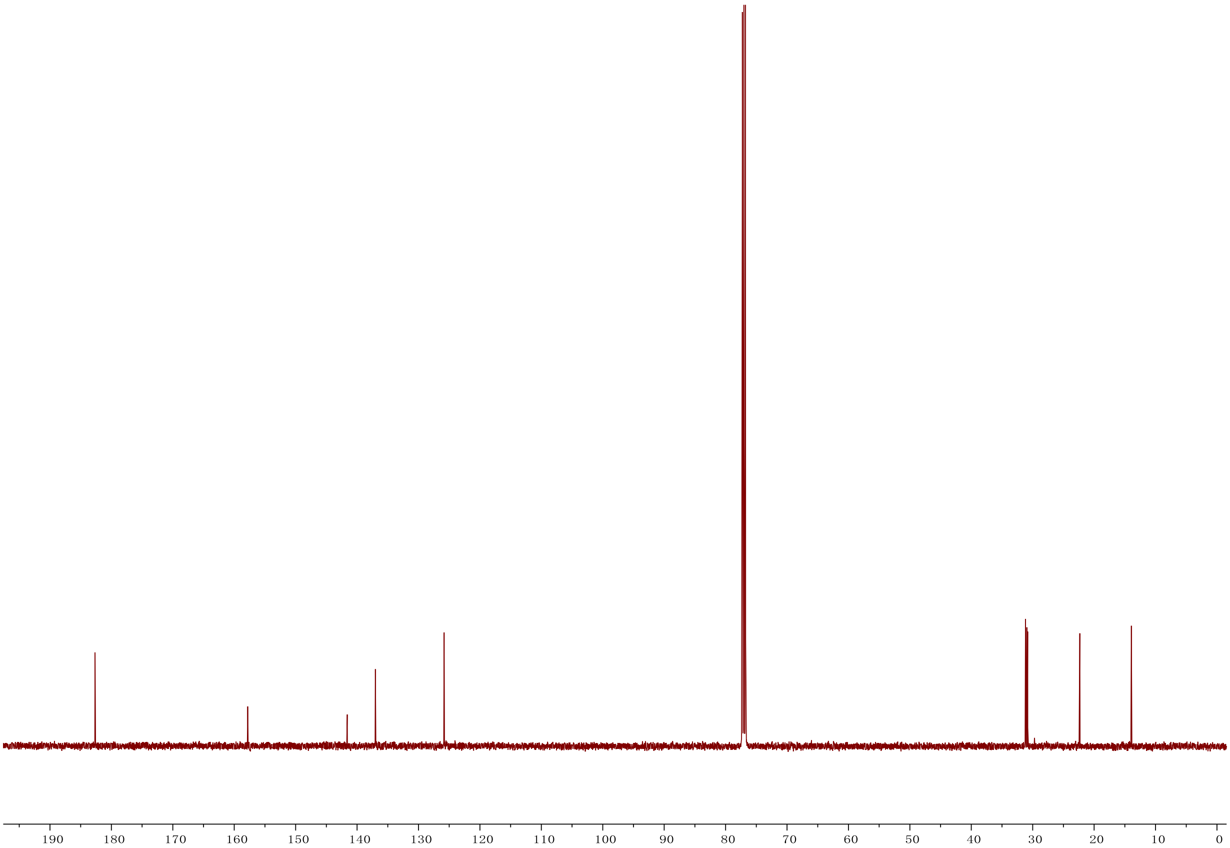

**[1]H NMR of compound 27**

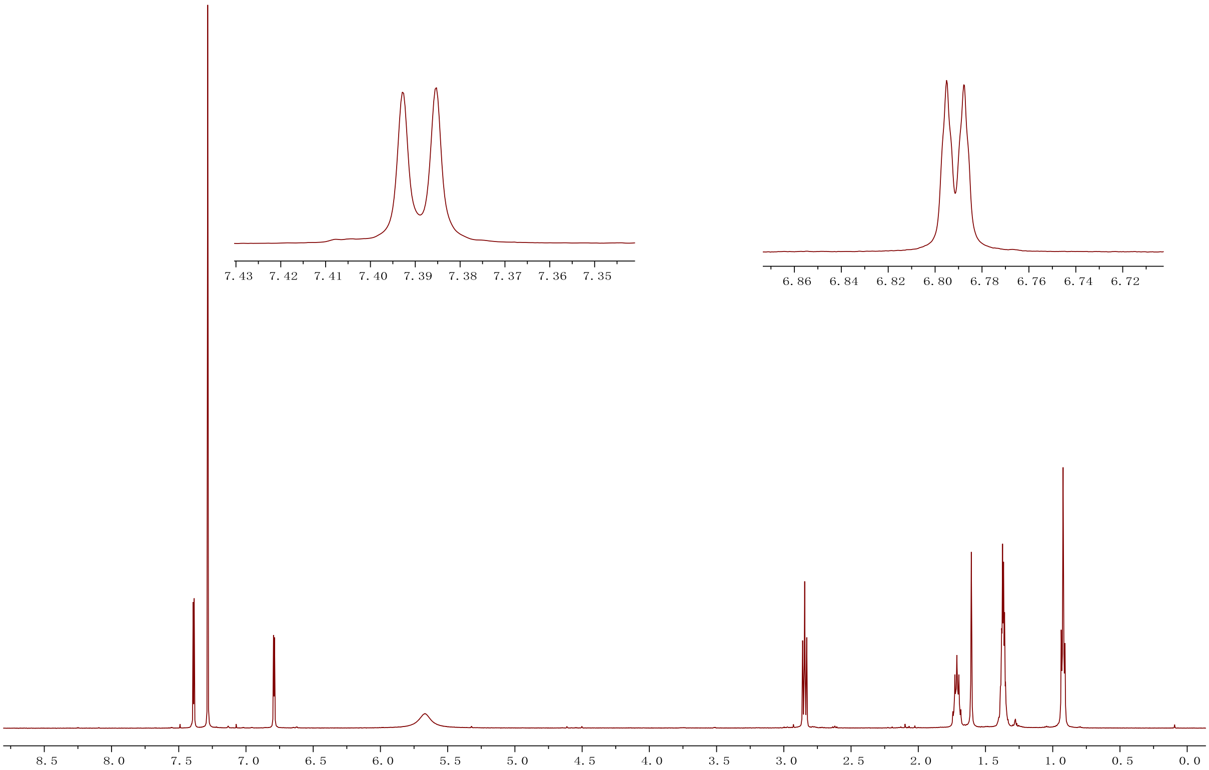


**[13]C NMR of compound 27**

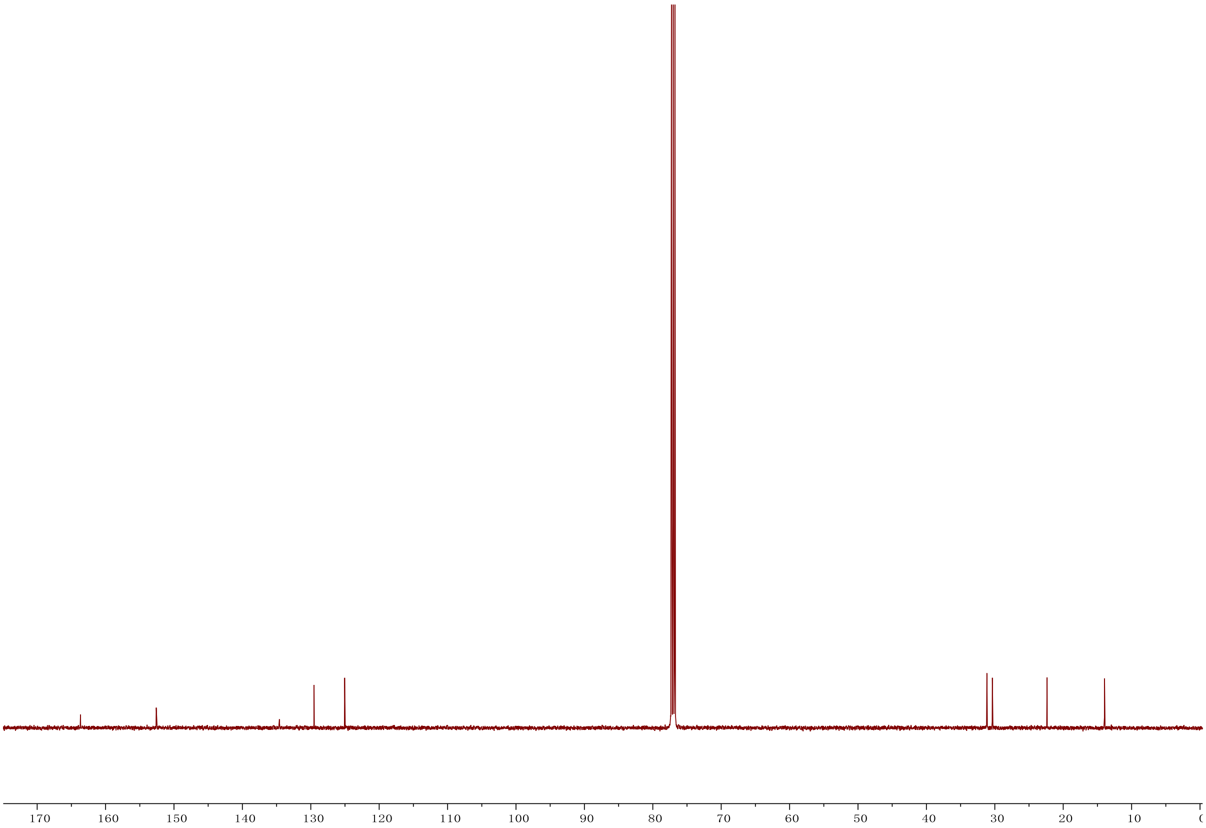
